# FilTar: Using RNA-Seq data to improve microRNA target prediction accuracy in animals

**DOI:** 10.1101/595322

**Authors:** Thomas Bradley, Simon Moxon

**Affiliations:** School of Biological Sciences, University of East Anglia, Norwich Research Park, Norwich, UK; Earlham Institute, Norwich Research Park, Norwich NR4 7UZ, UK

**Keywords:** miRNA target prediction, 3’ UTR, annotation, alternative polyadenylation

## Abstract

MicroRNAs (miRNAs) are a class of small non-coding RNA molecule, approximately 22nt in length, which guide the repression of mRNA transcripts. A number of tools have been developed to predict miRNA targets in animals which do not account for the effects of a specific cellular context on miRNA targeting. We present FilTar (**Fil**tering of predicted miRNA **Tar**gets), a method which utilises available RNA-Seq information to filter non- or lowly expressed transcripts and refine existing 3’UTR annotations for a given cellular context, to increase miRNA target prediction accuracy in animals.

The FilTar tool is available at https://github.com/TBradley27/FilTar.

## Introduction

miRNAs exert widespread post-transcriptional control over mRNA expression in most animal lineages (Bartel 2018), creating a need for the accurate identification of miRNA targets in order to better understand gene regulation. Traditional methods for providing experimental support for putative interactions include the use of reporter assays to test for a direct interaction between the miRNA and mRNA, or perturbation experiments to test for the effect of increased or decreased miRNA levels on target mRNA, or the corresponding proteins translated from these molecules (Kuhn *et al*. 2008). More recent methods allow researchers to test for direct interactions between miRNA and putative targets transcriptome-wide. These methods usually test for binding between the putative miRNA target and argonaute (AGO) (Chi *et al*. 2009; König *et al*. 2010; Van Nostrand *et al*. 2016), a key component of the miRNA-guided RISC (RNA-induced silencing) complex, and in addition some methods can also be used to determine the identity of the miRNA which is guiding AGO to the target transcript (Kudla *et al*. 2011; Helwak and Tollervey 2014).

Currently available data for these types of experiments are generally limited in number and diversity of cell types and species. Inspection of the TarBase resource (v8.0) (Karagkouni *et al*. 2017), a database of published, experimentally-supported predicted miRNA interactions, reveal that, at the time of writing, even for a widely utilised model organism such as mouse, AGO immunoprecipitation datasets are available for only three cell lines and five tissues. The problem is exacerbated when examining records for other model organisms such as rat and zebrafish, in which no data from immunoprecipitation experiments is reported. This is likely because generating data of this type is usually prohibitively expensive in terms of skills, time and material resources needed to complete sophisticated transcriptome-wide, next-generation library preparation and sequencing protocols. The limited applicability of experimental approaches therefore underlies the continuing necessity of computational approaches for predicting miRNA targets.

There are a number of existing computational tools for predicting miRNA targets in animals. Algorithms such as TargetScan use complementarity between the seed sequence of the miRNA (Lewis *et al*. 2003; Bartel 2018) and a corresponding region of the 3’UTR of its target as the basis of target prediction (Lewis *et al*. 2003; Lewis *et al*. 2005; Grimson *et al*. 2007; Friedman *et al*. 2009; Garcia *et al*. 2011; Agarwal *et al*. 2015). Alternatively, some miRNA target prediction algorithms do not require full complementarity in the miRNA seed region (Khorshid *et al*. 2013; Gumienny and Zavolan 2015; Enright *et al*. 2003; John *et al*. 2004; Wang 2016), or predict miRNA targeting to occur in the coding region of the transcript as well as the 3’UTR (Reczko *et al*. 2012). Most algorithms, in addition to considerations of seed complementarity, and the location of the target site within the transcript, also consider features such as the conservation of the miRNA target site in closely related species, the thermodynamic stability of the miRNA-mRNA duplex, and the structural accessibility of putative target sites to the miRNA-RISC complex, as variables which are also thought to influence miRNA targeting and subsequent transcript repression (Ritchie and Rasko 2014).

Although intramolecular features are often considered, current miRNA target predictions currently do not account for the broader cellular context in which miRNA targeting occurs. The clearest indication of this, is that current target prediction tools do not account for whether predicted targets are expressed within a given cell type or tissue. If the predicted target is not expressed, it cannot physically interact and be translationally inhibited or repressed by miRNA molecules. As expression profiles differ across different cell types and tissues, not incorporating expression information will then likely lead to false positive results when making miRNA target predictions.

For the prediction of miRNA targets in the 3’UTR, an additional complication is that the identity of an individual 3’UTR may not be stable across different cell types or different biological conditions due to alternative cleavage and polyadenylation (APA) (Tian and Manley 2017). APA is the process by which cellular polyadenylation machinery utilises alternative polyadenlyation sites located on precursor mRNA molecules to produce transcripts with alternative 3’UTR sequences. Differential usage of polyadenylation sites in diverse tissues or biological conditions, can result in distinct 3’UTR isoform abundance profiles existing between different cell types (Nam *et al*. 2014). One consequence of the existence of 3’UTR isoforms, is that a miRNA target site may exist for some 3’UTR isoforms of the same annotated mRNA, but not others.

As a result, APA allows the differential usage of miRNA target sites by the cell, diversifying and modifying the effect of miRNAs in different cellular contexts. For example, in cancer cells, shortening of 3’UTRs can activate oncogenes by increasing mRNA stability, partially through the reduction in the number of miRNA target sites in their 3’UTRs, decreasing the extent to which they are repressed (Mayr and Bartel 2009). In contrast, an extensive enrichment of longer 3’UTRs and hence additional miRNA target sites has been discovered in mammalian brain tissue (Miura *et al*. 2013), which has been hypothesised to serve as an extended platform for the regulation of gene expression (Wang and Yi 2014). This evidence of context-specific miRNA action underlies the utility of methods which accounts for this information in order to increase the precision and sensitivity of miRNA target predictions.

Most databases of miRNA target predictions do not incorporate information relating to APA, and instead rely on default 3’UTR annotations provided by public sequence databases such as Ensembl (Birney *et al*. 2004; Cunningham *et al*. 2019) and RefSeq (Pruitt *et al*. 2006; Pruitt *et al*. 2013), when identifying potential miRNA targets. Similarly, most prediction algorithms do not easily allow the user to generate predictions for multiple 3’UTR isoforms of the same mRNA. An exception is TargetScan (v7) (Agarwal *et al*. 2015). In this version each mRNA transcript is associated with a distinct profile of relative 3’UTR isoform abundances. From this profile, each scored target site is weighted by the abundance of the 3’UTR segment containing the predicted target site relative to all 3’UTRs of that transcript. The caveat of this analysis being that 3’UTR profiles are generated from sequencing data obtained from only four human cell lines (Nam *et al*. 2014), which is subsequently treated as being representative for all cell types. Whilst it was shown that this approach was superior to not incorporating 3’UTR profile data at all, it was sub-optimal in comparison to using 3’UTR profiles specific to each cellular context examined (Nam *et al*. 2014). Crucially, a miRNA target prediction tool which enables the user to predict miRNA targets specific to a given tissue or cell line is lacking.

Presented in this manuscript is FilTar, a tool which takes RNA-Seq data as input, and generates miRNA target predictions tailored to specific cellular contexts. Specificity of target prediction is increased by utilising information from sequencing data to both filter for abundant target transcripts and to refine 3’UTR annotations. Analysis demonstrates that predicted miRNA targets gained and lost due to 3’UTR reannotation do not substantially differ in their response to a miRNA than pre-existing miRNA targets and non-targets predictions, respectively. The cumulative effect of integrating these additional processing steps into conventional miRNA target prediction workflows is to increase prediction accuracy and to drastically alter the number of miRNA target predictions made between different cell types.

## Methods

### Workflow management and automation

All workflows are coordinated and managed by the FIlTar tool. FilTar is a command line tool for gnu-linux and macOS operating systems predominantly written in the python (v3.6.8) and R (v3.5.0) (R Core Team 2013) programming languages. Users can configure the tool to process available RNA-Seq datasets from public repositories (Leinonen *et al*. 2010a; Leinonen *et al*. 2010b and Harrison *et al*. 2018); and and also the user’s own private sequencing data. All parameters reported in this study, for given analysis and processing steps are configurable by the user. FilTar utilises Snakemake (v5.4.0) (Köster and Rahmann 2012) when managing workflows.

All of the following described analyses and data processing steps were managed within FilTar.

### Data selection, quality control, pre-processing and statistics

For analysis of miRNA transfection experiments, FASTQ sequencing data generated from RNA-Seq protocols in human or mouse cell lines with at least two biological replicates were selected for further processing. After differential expression analysis, if by inspection of cumulative plots the miRNA targets could not be observed to be downregulated relative to non-target transcripts, then the transfection experiment was considered to have failed, and relevant datasets were not used for downstream analysis (see supplementary file 1a and 1b).

For supplementary figures 3a and 3b, total reads were sampled using the seqtk tool (Li 2012).

Reads were trimmed using Trim Galore (v0.5.0) (Krueger 2015), a wrapper around Cutadapt (v1.16) (Martin 2011), using default parameters with the exception of the ‘length’ and ‘stringency’ parameters which were set to 35 and 4 respectively.

FASTQ data quality scores, GC-content, read lengths and similar statistics were generated using FASTQC (v0.11.5) (Andrews 2010). Output from FASTQC was collated with data from the log files of other processes in order to produce a summary statistics report for each used BioProject using MultiQC (v1.6) (Ewels *et al*. 2016) (see supplementary file 2).

A summary of datasets used with relevant database accessions can be found in supplementary table 5 (Tamim *et al*. 2014, Liu *et al*. 2017, Stolzenburg *et al*. 2016, Liu *et al*. 2019, Guo *et al*. 2014, Diepenbruck *et al*. 2017, Pua *et al*. 2016, Cao *et al*. 2015).

### 3’UTR reannotation

In order to build an index for the alignment of FASTQ reads to the genome, unmasked chromosomal reference genome assembly fasta files for human (GRCh38.p12) and mouse (GRCm38.p6) (Schneider *et al*. 2017) were downloaded from release 94 of Ensembl (Cunningham *et al*. 2019). All subsequent files obtained from the Ensembl resource were for this same release version. Splice-aware mapping of reads to the genome was achieved using HISAT2 (v2.1.0) (Kim *et al*. 2015): The location of exons and junction sites was determined by running the appropriate HISAT2 scripts on the relevant species-specific GTF annotation file also obtained from Ensembl. The ‘hisat2-build’ binary was executed using the ‘ss’ and ‘exon’ flags indicating splice site and exon co-ordinates built from the previous step.

The indexed genome was used for FASTQ read alignment using the ‘hisat2’ command. The ‘rna-strandness’ option was used for strand-aware alignment. The strandedness of RNA-seq datasets was predicted using the ‘quant’ command of the salmon (v0.11.3) (Patro *et al*. 2017) RNA-seq quantification tool, by setting the ‘lib-type’ option to ‘A’ for automatic inference of library type. The samtools (v1.8) (Li *et al*. 2009) ‘view’ and ‘sort’ commands were used to sort data from sam to bam format, and to sort the resultant bam files respectively.

Sorted bam files were converted to bedgraph format using the ‘genomeCoverageBed’ command of bedtools (v2.27.1) (Quinlan and Hall 2010; Quinlan 2014) using the ‘bg’,‘ibam’ and ‘split’ options. Bedgraph files representing biological replicates of the same condition were merged using bedtool’s ‘unionbedg’ command. FilTar then calculated the mean average coverage value for each record in the merged bedgraph file. Existing gene models were produced by converting Ensembl GTF annotations files into genePred format using the UCSC ‘gtfToGenePred’ binary, and then from genePred format to bed12 format using the UCSC ‘genePredToBed’ binary (Kent *et al*. 2002). APAtrap (Ye *et al*. 2018), the 3’UTR reannotation tool was used to refine 3’UTR annotations by integrating information from the bed12 file and bedgraph files using the ‘identifyDistal3UTR.pl’ perl script using default parameters.

FilTar then integrated existing 3’UTR models with new models predicted by APAtrap. Only truncations or elongations of single exon 3’UTR annotations were integrated into final 3’UTR annotations; novel 3’UTR predictions (*i.e.* prediction of 3’UTRs for transcripts without a previous 3’UTR annotation) were discarded and alterations of the 3’UTR start site were also not permitted, due to the reannotation of 3’UTR start sites by the APAtrap dependency as beginning at the start position of the final exon in standard Ensembl transcript models. No alterations to existing 3’UTR annotations spanning multiple exons were permitted, as this is not intended functionality of the APAtrap tool.

### miRNA Target Prediction

Target prediction for the analyses presented in this study was conducted using the TargetScan algorithm (v.7.01) (Agarwal *et al*. 2015). Mature miRNA sequences were obtained from release 22 of miRBase (Griffiths-Jones 2004; Kozomara *et al*. 2018). The 3’UTR sequence data required for target prediction can either be provided as multiple sequence alignments or single sequences, with the former option enabling the computation of 3’UTR branch lengths and the probability of conserved targeting (P_ct_) for putative miRNA target sites.

Multiple sequence alignments (MSA) are derived from 100-way (human reference) and 60-way (mouse reference) whole-genome alignments hosted at the UCSC genome browser (Kent *et al*. 2002) generated using the threaded blockset-aligner (Blanchette *et al*. 2004) stored in MAF (multiple alignment format) format. MAF files are indexed, and the relevant alignment regions corresponding to 3’UTR co-ordinates extracted using ‘MafIO’ functions contained within the biopython (v1.72) library (Cock *et al*. 2009). For human MSAs, during post-processing, distantly related species were removed, resulting in 84-way multiple sequence alignments (Agarwal *et al*. 2015)

If multiple sequence alignments are not used, single sequences are extracted from DNA files using relevant 3’UTR co-ordinates in bed format using the ‘getfasta’ command of bedtools with the ‘s’ option enabled. Custom scripts are used to process the output of this command in order to merge exon sequences, into a single contiguous 3’UTR sequence. Further scripting is required to convert miRNA and 3’UTR sequence and identifier information to a format which can be parsed by TargetScan algorithms.

TargetScan is executed using both Ensembl 3’UTR annotations, and updated annotations produced using FilTar for the purposes of the differential expression analysis.

The FilTar tool is also fully compatible with the miRanda (v3.3a) (Enright *et al*. 2003; John *et al*. 2004) miRNA target prediction algorithm allowing users to identify non-canonical miRNA targets i.e. predicted targets without a perfectly complementary seed match to the miRNA.

### Transcript quantification

Human and mouse cDNA files were downloaded from Ensembl. Kallisto (v0.44.0) (Bray *et al*. 2016) was used to index the cDNA data using the ‘kallisto index’ command with default parameters. Reads were pseudoaligned and relative transcript abundance quantified using the ‘kallisto quant’ executable, using the ‘bias’ option to correct for sequence-based biases. When kallisto was used with data derived from single-end RNA-sequencing experiments, 180nt and 20nt were used as required estimates of the mean average fragment length and standard deviation respectively.

### Differential expression analysis

Differential expression analysis for miRNA transfection experiments was completed within the R (v.3.5.0) statistical computing environment. Transcript-level read count data derived from RNA sequencing of miRNA mimic or negative control transfected cell lines were imported using the tximport package (v1.10.1) (Soneson *et al*. 2015). Differential expression analysis on length and library size normalised read counts was performed using DESeq2 (v1.22.2) (Love *et al*. 2014) comparing expression between negative control and miRNA mimic transfection conditions. Log_2_ fold change values were subsequently shrunken using the default DESeq2 ‘normal’ shrinkage estimator (Love *et al*. 2014) to account for the large uncertainty in predicted fold change values at low transcript expression values. For plotting, records corresponding to non-coding RNA transcripts were discarded. Transcript records were discarded when there was zero expression for all control and transfection replicates and fold change values could not be calculated. Target prediction data was used to label the remaining records as either predicted targets or non-targets of the transfected miRNA.

For all differential expression analyses, null hypothesis significance testing was performed using two-sample, one-sided Kolmogorov-Smirnov tests to test whether different fold change distributions were sampled from the same underlying distribution.

### Data Visualisation

All visualisations are produced using R’s ggplot2 package (v3.1.0) (Wickham 2016).

For figure 1, the filtered miRNA predicted target set represent protein-coding transcripts with a miRNA seed target site to the transfected miRNA mimic, which have filtered at an expression threshold of 0.1 *Transcripts per million* (TPM) (Li *et al*. 2009).

**Figure 1:**
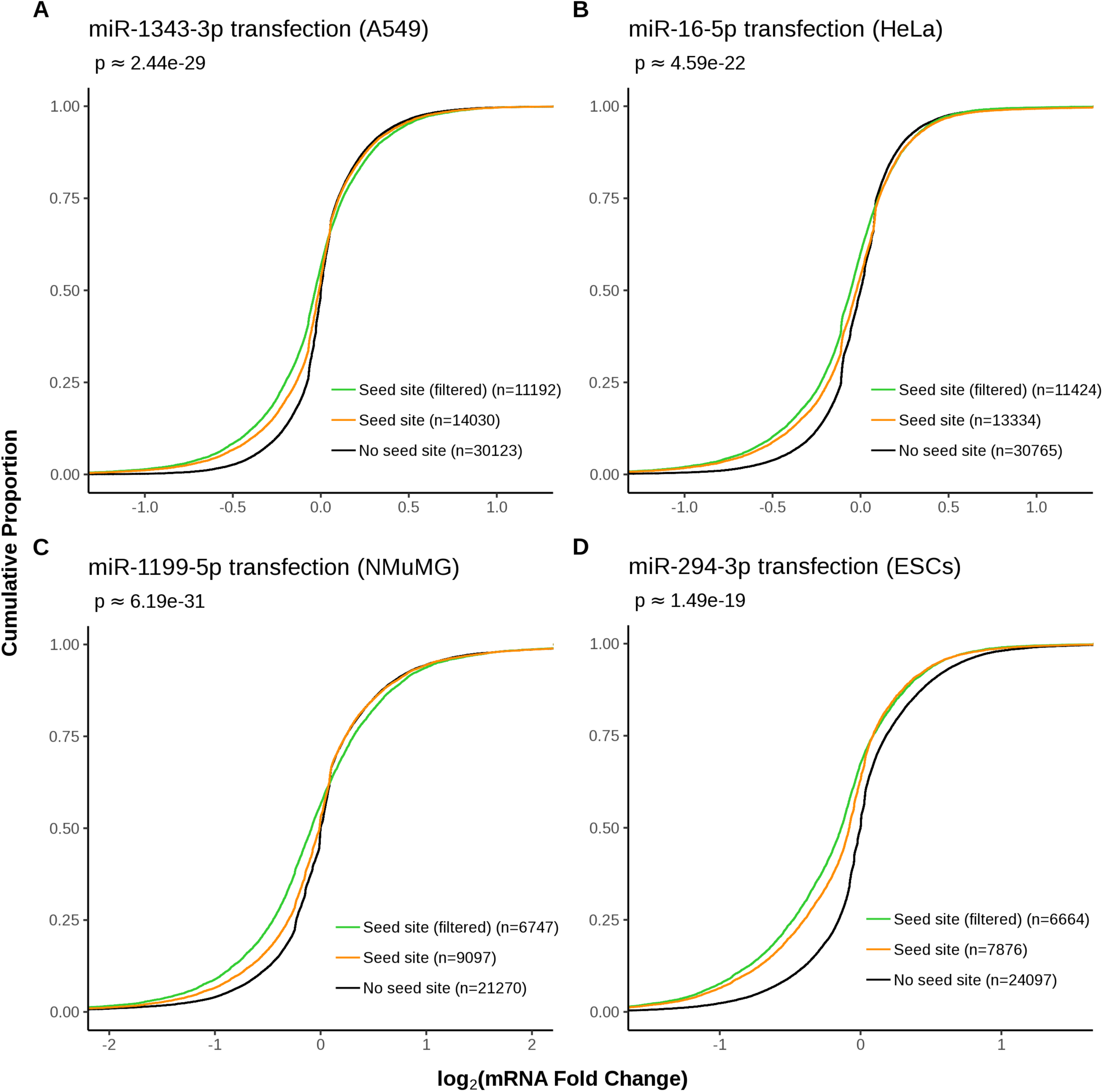
Cumulative plots demonstrating the effect of miRNA mimic transfection on expression filtered (TPM > 0.1) miRNA seed targets. Curves are plotted of the cumulative log_2_ fold change distributions of i) protein-coding non-target transcripts (black) ii) protein-coding seed target transcripts (orange) and iii) expression filtered protein-coding seed target transcripts (green). Numbers in brackets represents the number of mRNA transcripts found in each set. Approximate P-values were computed using one-sided, two-sample, Kolmogorov-Smirnov tests between full target and filtered target fold change distributions. Data presented for miRNA mimic transfection into **A)** A549 and **B)** HeLa cell lines, **C)** normal murine mammary gland (NMuMG) cells and **D)** mouse embryonic stem cells (ESCs). Results from the application of this analysis to additional datasets can be found in the supplementary file 3.

For figure 2, the ‘added seed sites’ are identified as those transcripts which had not previously been labelled as predicted miRNA targets using target prediction results derived from existing Ensembl 3’UTR annotations, but had been identified as predicted miRNA targets using target prediction results derived from 3’UTR sequences reannotated using the FilTar workflow due to 3’UTR extension.

**Figure 2:**
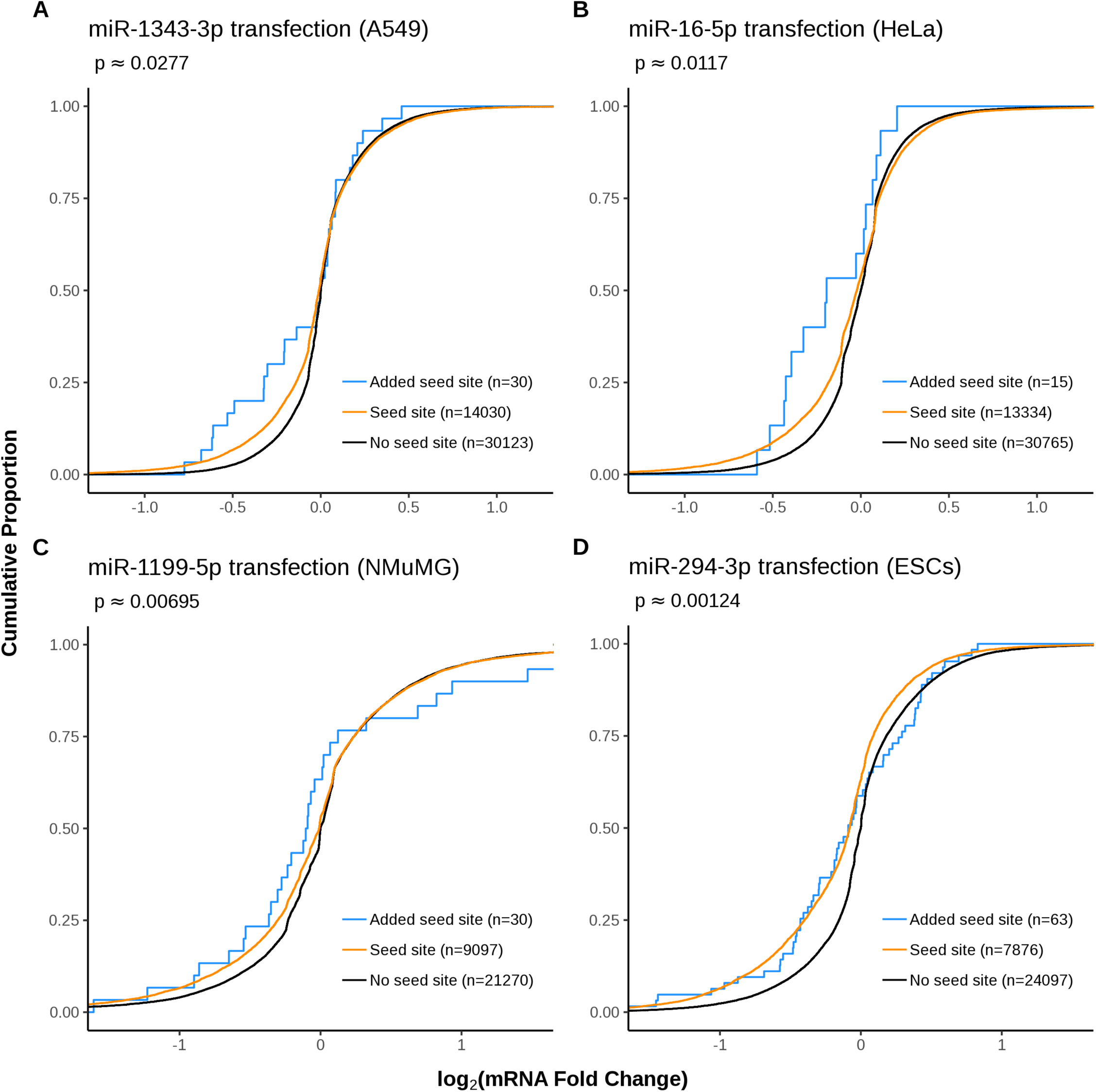
Cumulative plots demonstrating the effect of miRNA mimic transfection on predicted miRNA target transcripts newly identified by the FilTar workflow. Curves are plotted of the cumulative log_2_ fold change distributions of i) protein-coding non-target transcripts (black). ii) protein-coding seed target transcripts (orange) and iii) predicted target transcripts deriving from FilTar 3’UTR annotations but not Ensembl 3’UTR annotations (blue). Approximate P-values were computed using one-sided, two-sample, Kolmogorov-Smirnov tests between pre-existing target and newly identified target fold change distributions. Otherwise as in figure 1.

For figure 3, the ‘removed seed sites’ are identified as those transcripts which had previously been labelled as predicted miRNA targets using target prediction results derived from existing Ensembl 3’UTR annotations, but had not been identified as predicted miRNA targets using target prediction results derived from 3’UTR sequences reannotated using the FilTar workflow due to 3’UTR truncation. Filtering for all groups occurred at an expression threshold of greater than or equal to 5 TPM. This was to reduce the number of false positive 3’UTR truncations (see discussion).

**Figure 3:**
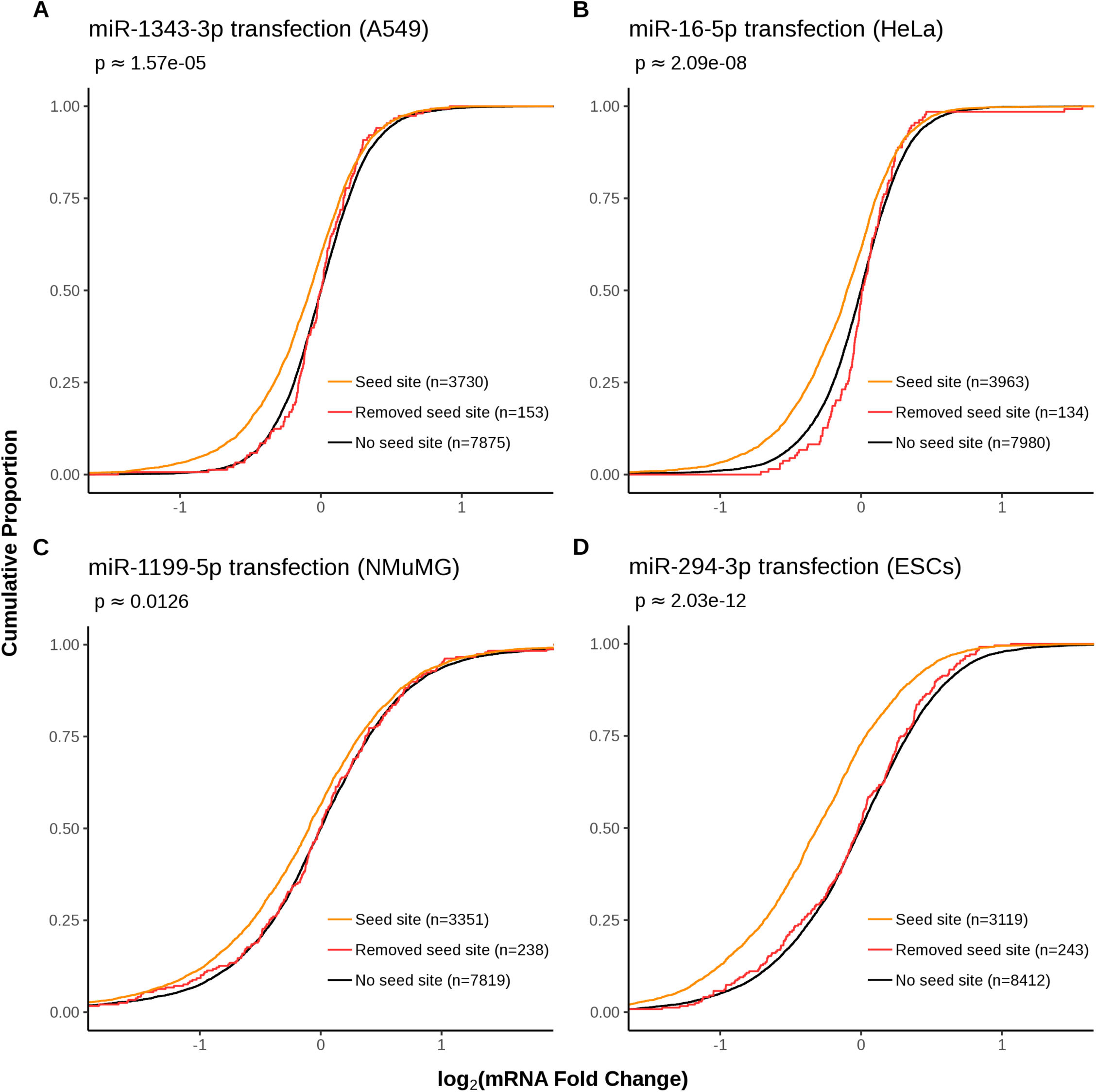
Cumulative plots demonstrating the effect of miRNA mimic transfection on previously predicted miRNA target transcripts discarded by the FilTar workflow. Curves are plotted of the cumulative log_2_ fold change distributions of expression filtered i) protein-coding non-target transcripts (black). ii) protein-coding seed target transcripts (orange) and iii) predicted target transcripts deriving from Ensembl 3’UTR annotations but not FilTar 3’UTR annotations (red). Approximate P-values were computed using one-sided, two-sample, Kolmogorov-Smirnov tests between non-target and discarded miRNA target fold change distributions. Otherwise as in figure 1.

Additional plots for remaining datasets analysed can be found in the supplementary materials (supplementary files 3, 4 and 5) with the exception of cases were there was an insufficient number of added or removed target transcripts predicted (n < 15).

## Results

Predicted miRNA targets with TPM > 0.1 as a whole, exhibited stronger repression after miRNA transfection than the full miRNA target set without expression filtering (Figure 1; upplementary file 3). Predicted miRNA targets removed for low expression generally exhibited low absolute fold change values (supplementary figure 1).

Newly gained miRNA target predictions deriving from FilTar’s refined 3’UTR annotations of protein-coding transcripts (*i.e.* miRNA targets deriving from the elongation of existing 3’UTR annotations), generally exhibited similar levels of repression to miRNA target predictions deriving from Ensembl 3’UTR annotations (Figure 2; supplementary file 4). Anomalies were results deriving from the transfection of miR-107 and miR-10a-5p miRNA mimics into HeLa cells in which newly identified miRNA target predictions did not exhibit a log fold change distribution commensurate with that exhibited by already existing miRNA target predictions (supplementary file 4).

Conversely, miRNA target transcripts that were removed as a result of FilTar truncating 3’UTR annotations relative to standard Ensembl annotations, exhibited repression similar to that of annotated non-target transcripts (figure 3; supplementary file 5). In a minority of datasets analysed, removed target transcripts exhibited significantly less repression than target transcripts, but nonetheless exhibited greater repression than annotated non-target transcripts. In these datasets, the removed target log fold change distribution tended to align with the non-target distribution at the negative extremity, but not at small negative fold change value ranges - indicating that for a minority of datasets, labelled ‘removed targets’ may be mildly repressed by targeting miRNAs. Additional analysis demonstrated that for these datasets, such targets exhibited significantly weaker repression in response to miRNA transfection than 6mer targets, which are the weakest canonical miRNA target sites (supplementary figure 2).

When the FilTar reannotation and miRNA target prediction workflow was applied transcriptome-wide, to multiple organs and cell lines, using all annotated miRBase human miRNAs, there was a mean average gain and loss of miRNA target sites corresponding to 0.18% and 1.5% of the total original miRNA target sites predicted deriving from Ensembl 3’UTR annotations (Figure 4), corresponding to a gain and loss of total miRNA seed sides in the tens and hundreds of thousands respectively (supplementary table 4). Whilst a much larger proportion of miRNA seed sites (mean average of 26.3%) are lost through expression filtering (supplementary figure 5), representing a loss of millions of miRNA seed sites (supplementary table 4). This is commensurate with the mean average of 34.0% of 3’UTR bases lost when removing lowly expressed transcripts from target predictions (supplementary table 2). When considering the combined effect of expression filtering and 3’UTR reannotation, a mean average 36.1% of 3’UTR bases are lost, affecting a mean average of 53.4% of protein-coding 3’UTRs (supplementary table 3).

**Figure 4:**
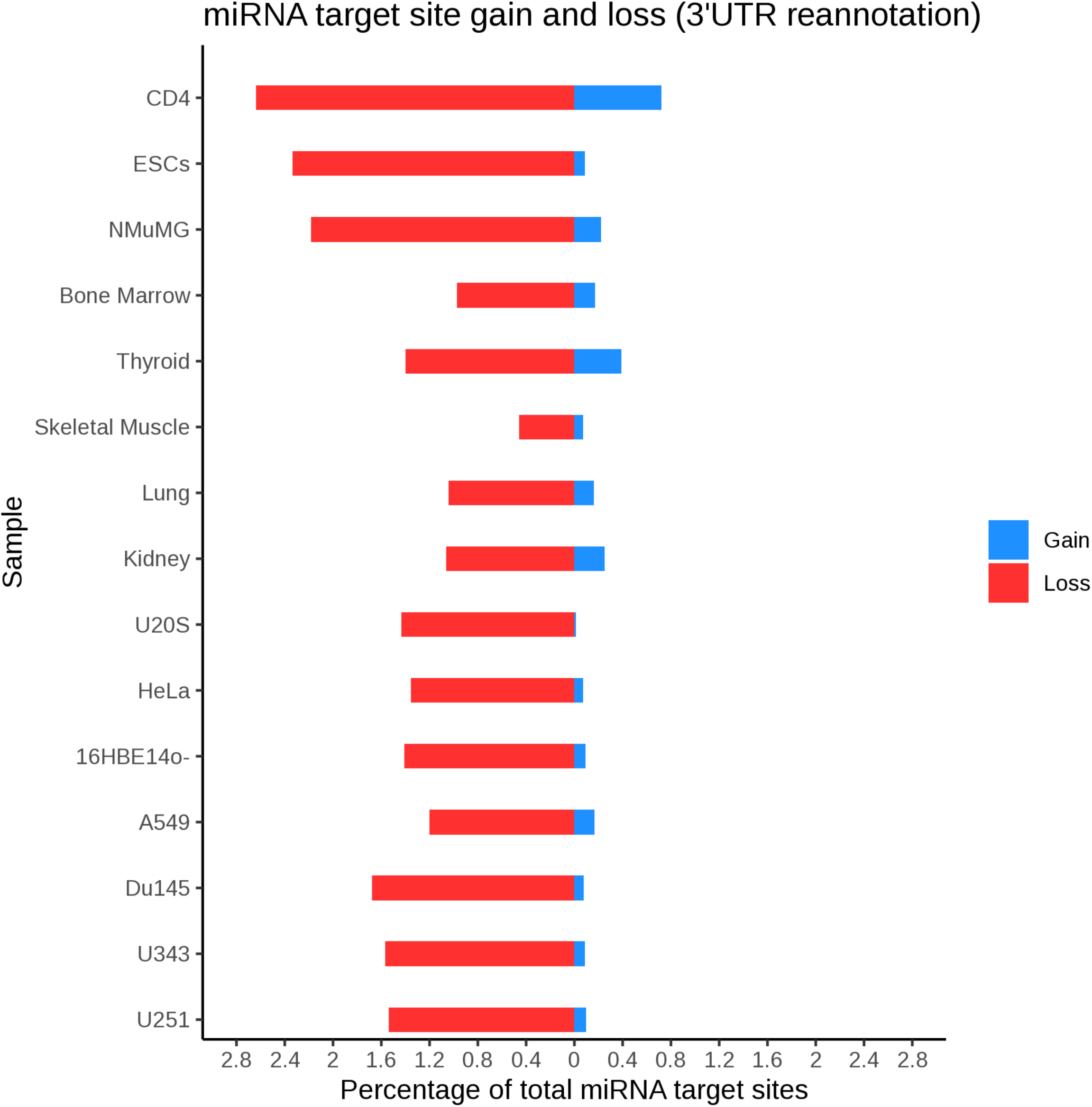
miRNA target site gain and loss across the protein-coding transcriptome when FilTar is used with all annotated human miRNAs for multiple tissues, organs and cell lines. Gained (blue) and lost (red) miRNA target sites is expressed as a percentage of the total number of target sites identified when deriving miRNA from Ensembl 3’UTR annotations

## Discussion

Results show that FilTar is successfully able to utilise RNA-Seq data to reannotate protein-coding 3’UTR sequences and filter based on expression data leading to a gain in specificity and sensitivity of target prediction evidenced through tests using experimental data.

That expression filtering target transcripts at even a modest expression threshold of 0.1 TPM leads to a loss of over a million seed sites in most datasets analysed represents a radical reduction in the number of false positive predictions associated with miRNA target prediction and is indicative of the importance of considering the biological plausibility of candidate miRNA interactions.

The number of newly predicted miRNA target sites deriving from FilTar elongated 3’UTR sequences is generally relatively low. For cell line datasets analysed, the maximum of number of newly predicted miRNA targets made for any single miRNA was 67, with the majority of datasets analysed yielding less than 15 newly predicted targets (figure 1, figure 4 and supplementary file 4). The number of newly identified target transcripts is commensurate with the universally low proportion of 3’UTRs extended, and the small proportion of bases added to the total of the 3’UTR annotation (supplementary table 1), even though this still represents a substantial increase in the number of miRNA seed target sites identified. This is in contrast to 3’UTR truncation in which the proportion of 3’UTRs truncated and bases removed from the 3’UTR annotation total are much greater. Analysis shows that there is a strong positive correlation between the number of 3’UTR bases reannotated, and the number of predicted miRNA target sites gained or lost through reannotation (supplementary Figures 6a and 6b). The bias in 3’UTR truncation as opposed to elongation can possibly be explained by either a pre-existing bias in standard Ensembl 3’UTR annotations to generate long 3’UTR models, or rather a bias in the FilTar reannotation workflow for 3’UTR truncation rather than elongation. A potential bias in the standard Ensembl annotation workflow could potentially be explained by the method of transcript annotation, in which, although transcript models are built on a tissue-specific basis, transcript models incorporated into the final Ensembl gene set typically only derive from the merging of RNA-sequencing reads from multiple different tissue samples (Aken *et al*. 2016), therefore creating a bias towards the annotation of longer 3’UTRs. This effect may be exacerbated or supplemented by the existence of 3’UTR isoforms within a given sample and transcript - creating relatively low abundance isoforms towards the distal end of the 3’UTR, making annotation difficult, and likely generating a large amount of uncertainty, biases and variability in different methods used to models used to estimate 3’UTRs.

Another possibility, is that the shortening and extension of existing 3’UTR annotations are qualitatively different problems requiring different respective sequencing depths. Within a given sample, a read sampling analysis demonstrates that there is a positive relationship, up to a point of saturation between sequencing depth and the number of bases used to elongate existing 3’UTRs (supplementary figure 3a). In addition, the saturation point for the addition of bases to 3’UTRs is still substantially less than the proportion of bases removed at 3’UTRs even at relatively low sequencing depths indicating that the discrepancy between proportion of 3’UTR bases added or subtracted from the 3’UTRs cannot be explained by insufficient sequencing depth. A similar positive relationship is observed between sequencing depth and the number of based truncated from existing 3’UTRs (supplementary figure 3b), although far less reads seem to be required for saturation to occur, indicating a weaker reliance on sequencing depth for 3’UTR truncation compared to 3’UTR elongation.

Although as mentioned previously, the sequencing depth does seem to influence the extent of 3’UTR reannotation, for a set of different biological samples, sequencing depth alone seems to have limited predictive value for this variable (supplementary figures 4a and 4b). The likely explanation being that as well as sequencing depth, the extent of 3’UTR reannotation is also determined by other key variables such as the cell type being analysed, read length used for sequencing, library preparation protocol, the use of single-end or paired-end sequencing, as well as additional researcher or lab-specific batch effects (Leek *et al*. 2010). For example, as some cell types are biased towards shorter 3’UTRs (Mayr and Bartekl 2009), whilst other towards longer 3’UTRs (Miura *et al*. 2013), generating radically different reannotation statistics irrespective of sequencing depth used.

As mentioned previously, there was generally a much larger number of miRNA target sites predicted to be removed than added during 3’UTR reannotation. This is despite FilTar permitting 3’UTR truncations only occurring on moderately-to-highly expressed transcripts after discovery that the reannotation of the 3’UTRs of lowly expressed transcripts generated a relatively large number of what seemed to be false positive predictions (supplementary Figure 7). The likely cause being that low transcript expression leads to sporadic and inconsistent coverage across the 3’UTR, in which there is insufficient information to correctly call 3’UTR truncation. The default behavior of the FilTar tool therefore is to only truncate the 3’UTRs of transcripts which are not poorly expressed (*i.e.* TPM > 5).

When examining 3’UTR truncations further, for a minority of datasets analysed, some removed miRNA predicted targets seem to be marginally effective, with some transcripts exhibiting low levels of repression upon transfection of the miRNA mimic. Further analysis indicates that these marginally repressed transcripts exhibit even weaker repression than 6-mer targeted transcripts (supplementary figure 2), one of the least effective canonical miRNA target types (Bartel 2018), indicating that the efficacy of these site types is marginal. A possible explanation for the existence of these site types is that, for some transcript annotations for which the 3’UTR was truncated, there may exist a small proportion of isoforms with longer 3’UTRs, which are too low in abundance to be detected by APAtrap, but nonetheless still confer a marginal level of repression to the transcript, and hence is detectable when analysing experimental data.

Investigations into the effect of utilising expression data when making transcriptome-wide miRNA target predictions can be extended by closer examination of not only the refinement of 3’UTR annotations across different biological contexts, and its effects on miRNA target prediction, but more precisely the definition of specific 3’UTR profiles, incorporating information about 3’UTR isoforms within a given cellular context (Agarwal *et al*. 2015). This enables the weighting of miRNA target prediction scores on the basis of sequencing data applied by the user themselves, enabling even further and extended tailoring of miRNA target prediction to the specific biological context being researched. Previous analyses indicate that the most effective target predictions occur when those predictions are weighted on the basis of 3’UTR isoform ratios (Nam *et al*. 2014). In addition, the scope of FilTar’s functionality can be increased by enabling the annotation of novel 3’UTR sequences for transcripts without a current annotated 3’UTR, and also for those 3’UTRs which themselves span multiple exons. In addition, both the configurability and precision of FilTar can be improved in the future by respectively, enabling use of additional tools for 3’UTR reannotation (Gruber et al. 2018a; Gruber et al. 2018b) and exploring the greater transcriptomic resolutions enabled by nascent single cell sequencing technologies.

## Conclusion

FilTar utilises RNA-Seq data to increase the accuracy of miRNA target predictions in animals by filtering for expressed mRNA transcripts and reannotating 3’UTRs for greater specificity to a given cellular context of interest to the researcher. FilTar’s compatibility with user-generated RNA-Seq data, confers functionality across a wide-range of potential biological contexts.

## Software Availability

The FilTar workflow can be downloaded from GitHub using the following URL: https://github.com/TBradley27/FilTar.

## Acknowledgements

We would like to thank Daniel Mapleson, Robert Davey, Tamas Dalmay and members of the Dalmay Lab for helpful comments and discussion. We would like to thank Dagnė Daškevičiūtė for help with the identification of appropriate miRNA mimic transfection datasets. This research was supported in part by the University of East Anglia high-performance computing (HPC) team, NBI Computing infrastructure for Science (CiS) group and the Earlham Institute (EI) Scientific Computing group through use of HPC and data storage resources, and assistance provided for the use of these resources.

## Supplementary Figures & Tables

**Supplementary Figure 1:**
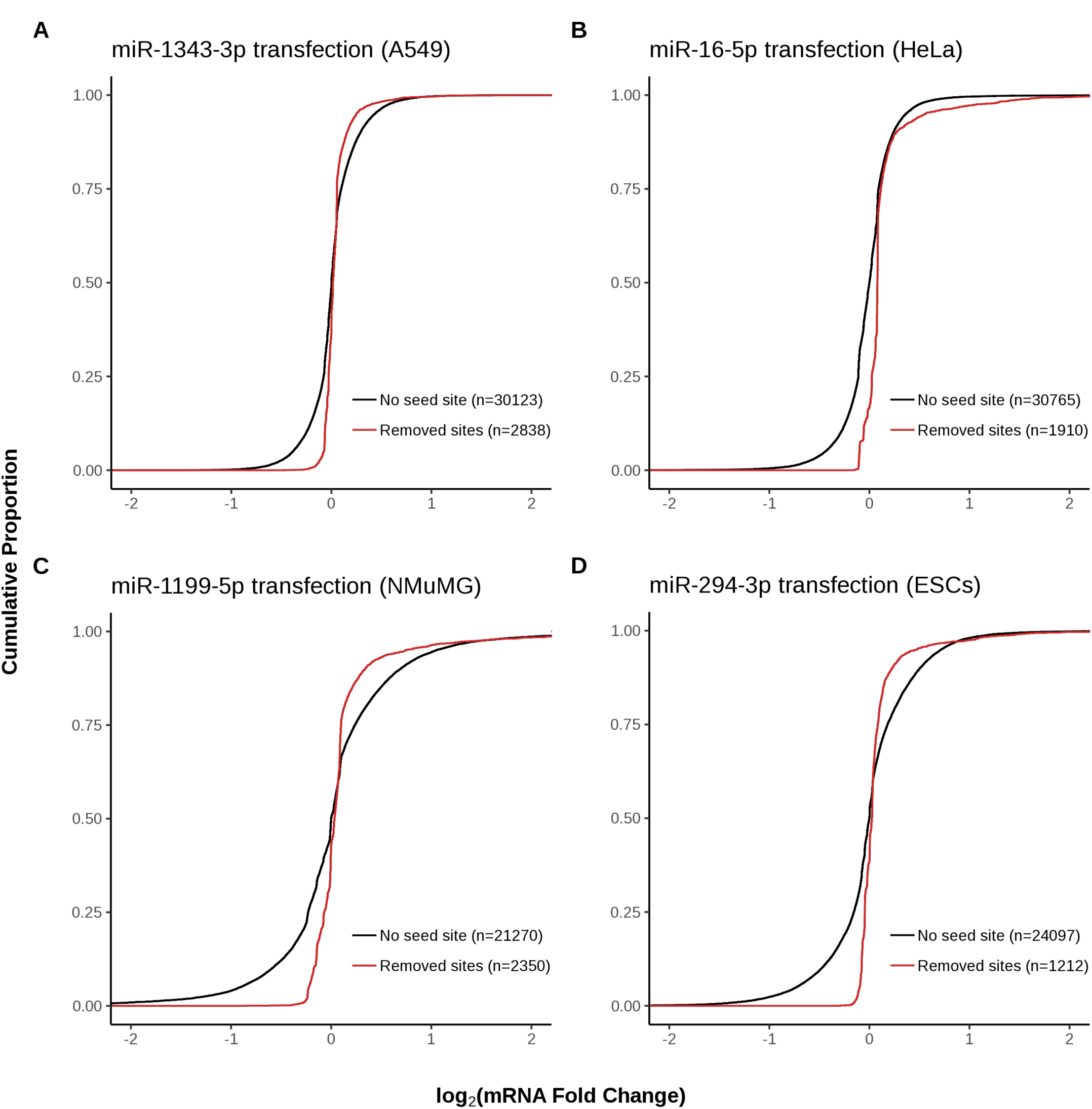
For the analysis presented in figure 1, the cumulative log_2_ fold change distributions of lowly expressed transcripts (<0.1 TPM) with canonical seeds sites (dark red), in their 3’UTRs compared against the distribution of transcripts without a canonical seed site in their 3’UTRs (black).

**Supplementary Figure 2:**
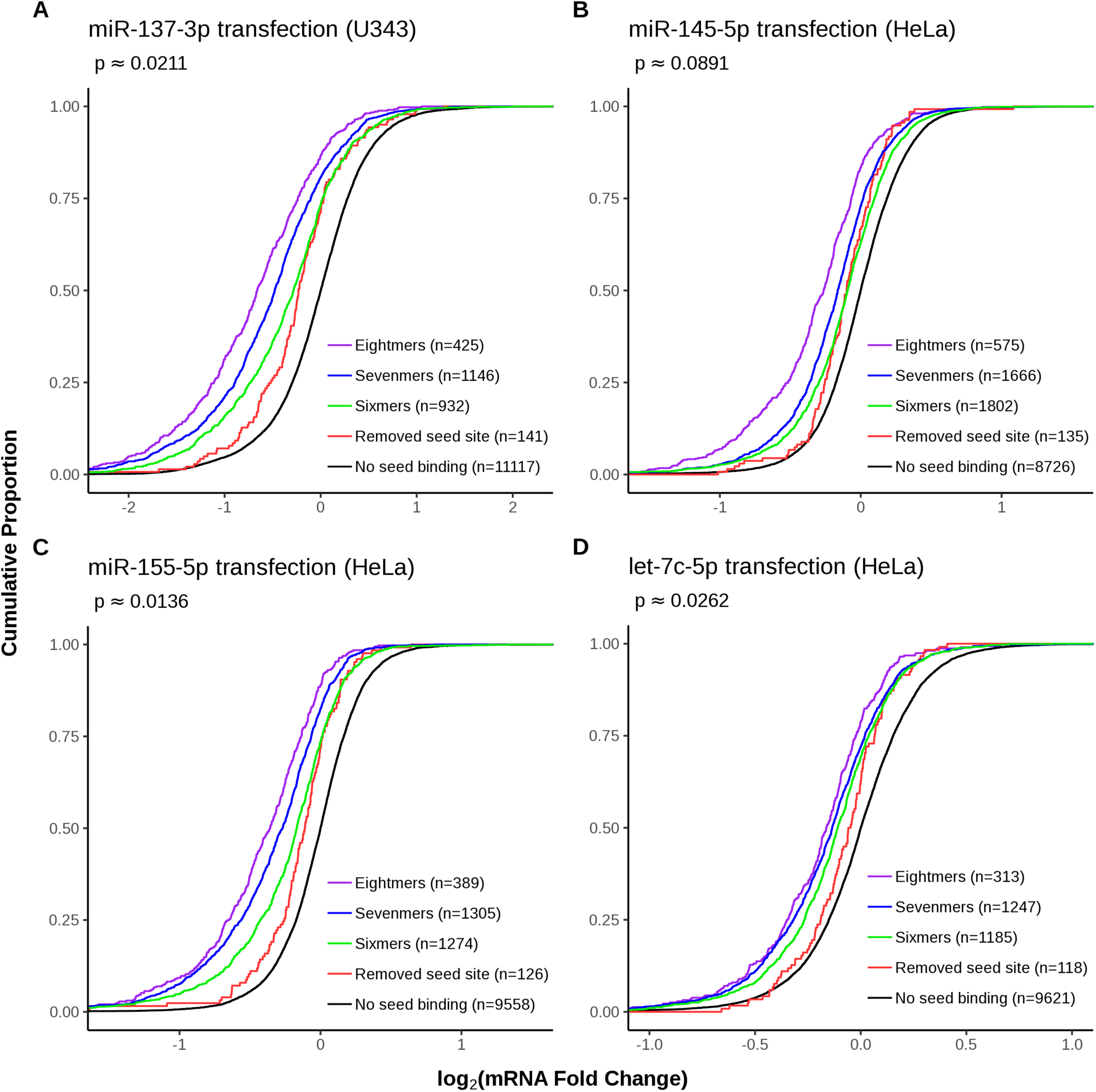
In experiments in which removed predicted target transcripts exhibit evidence of low-level repression, repression is less than that observed by transcripts targeted by marginally effective sixmer seed sequences. As in figure 3, with predicted target transcripts divided by miRNA target site type into sixmer (green), sevenmer (blue) and eightmer (purple) subsets. Approximate P-values were computed using one-sided, two-sample, Kolmogorov-Smirnov tests between discarded miRNA target and sixmer target fold change distributions.

**Supplementary Figure 3a:**
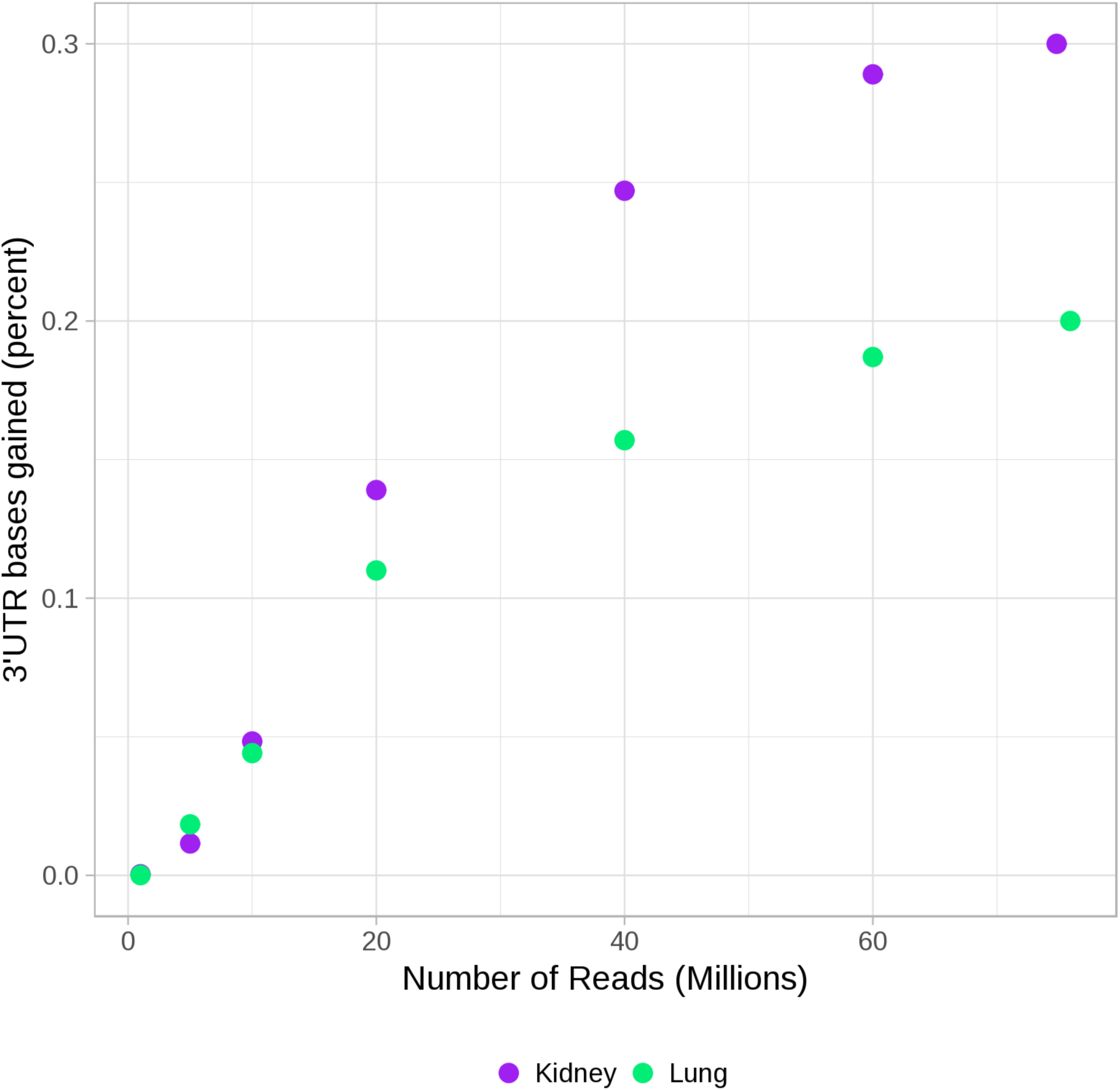
The relationship between the number of reads sequenced and the extent of 3’UTR elongation observed when using FilTar for human kidney (purple) and lung (green) datasets. Variable read counts generated by randomly sampling reads from the total.

**Supplementary Figure 3b:**
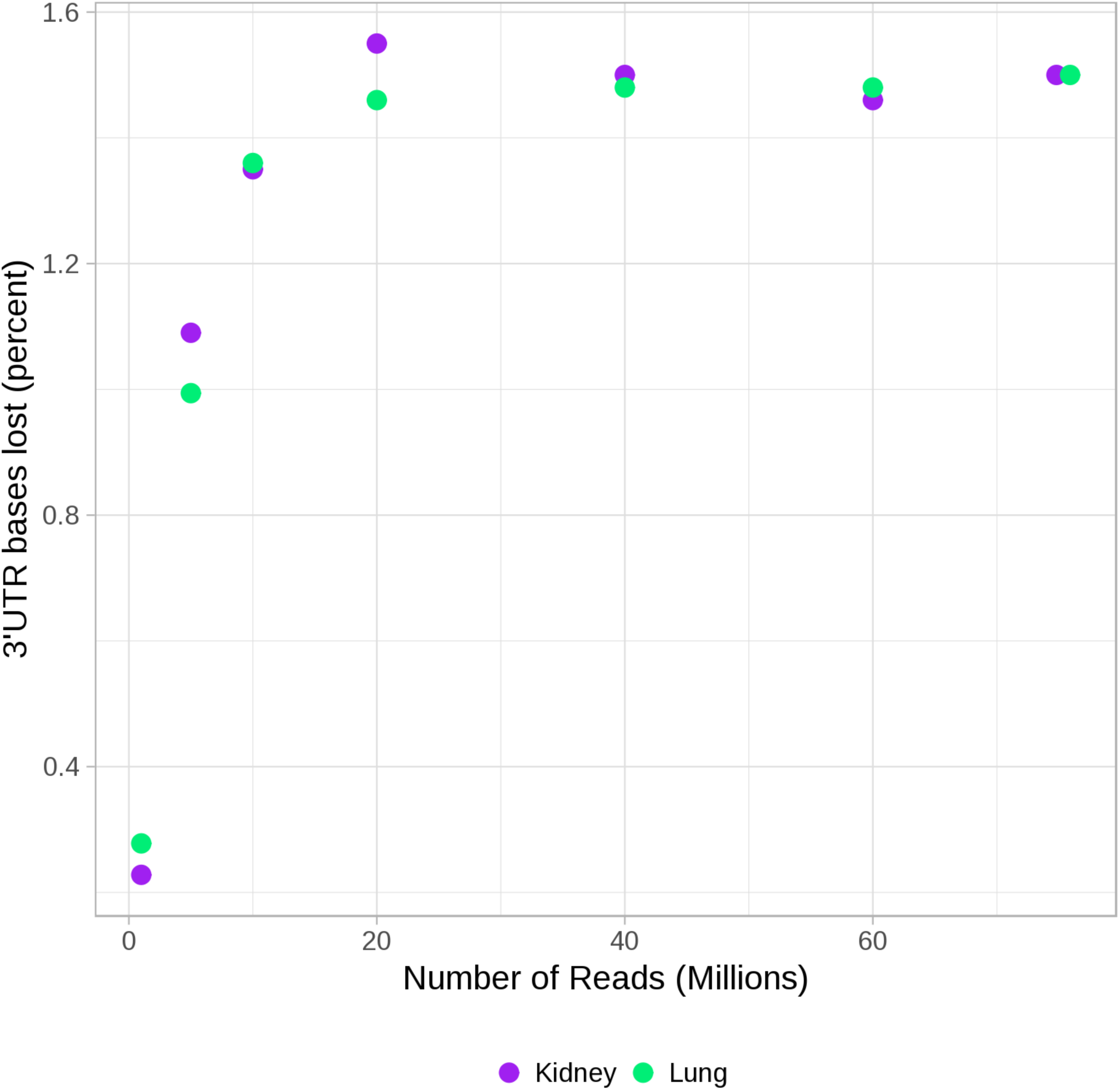
The relationship between the number of reads sequenced and the extent of 3’UTR truncation observed when using FilTar within a given sample. Otherwise as in supplementary figure 3a.

**Supplementary Figure 4a:**
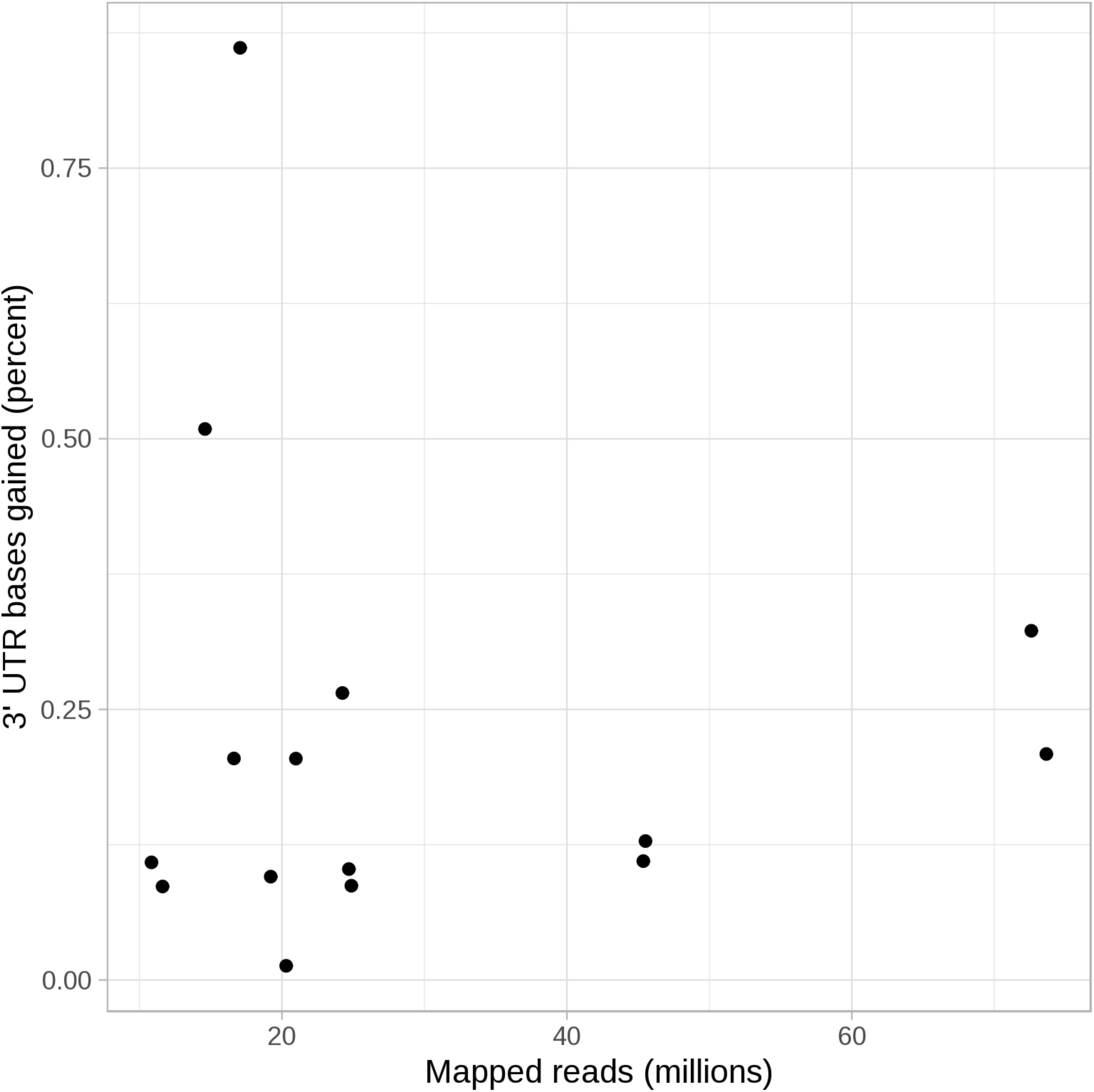
The relationship between the number of mapped reads and the extent of 3’UTR elongation observed when using FilTar. Each point represents a different dataset analysed using FilTar. Refer to suuplementary table 1 for metadata for datasets analysed.

**Supplementary Figure 4b:**
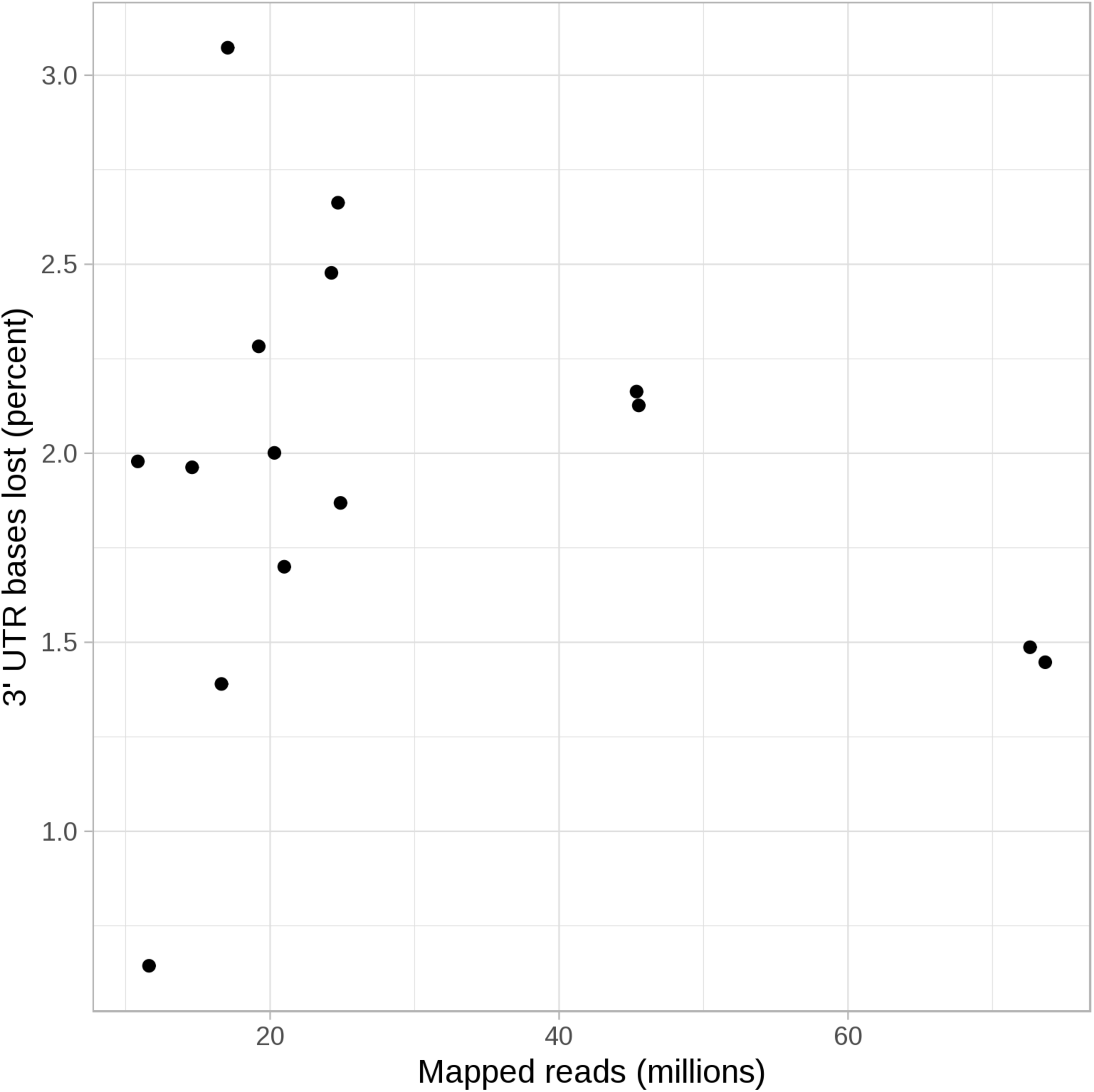
The relationship between the number of mapped reads and the extent of 3’UTR truncation observed when using FilTar. Otherwise as in supplementary figure 4a.

**Supplementary Figure 5:**
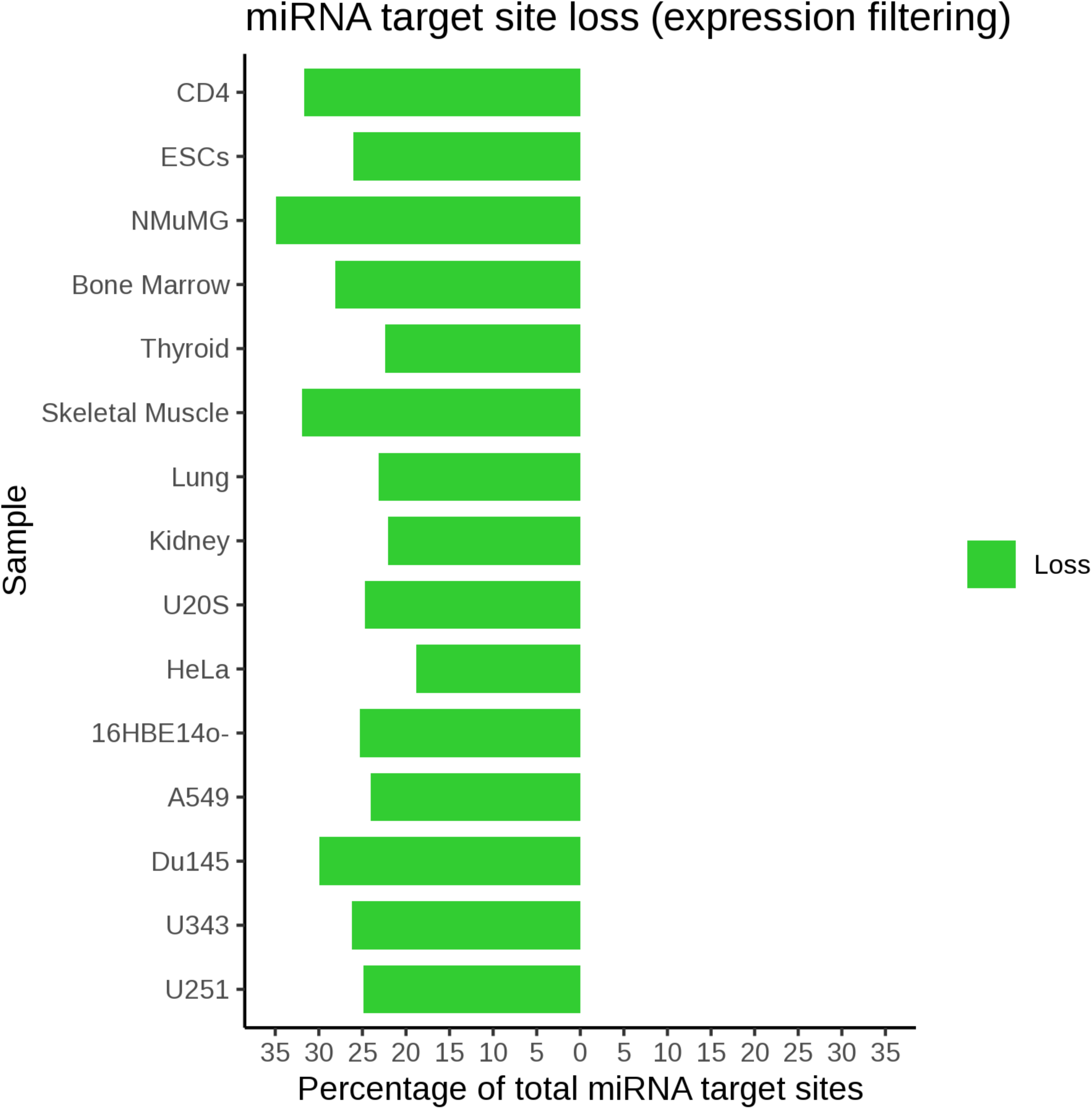
The percentage of total miRNA targets lost through expression filtering at a threshold of 0.1 TPM in a set of different cell lines and tissue types for human and mouse species.

**Supplementary Figure 6a:**
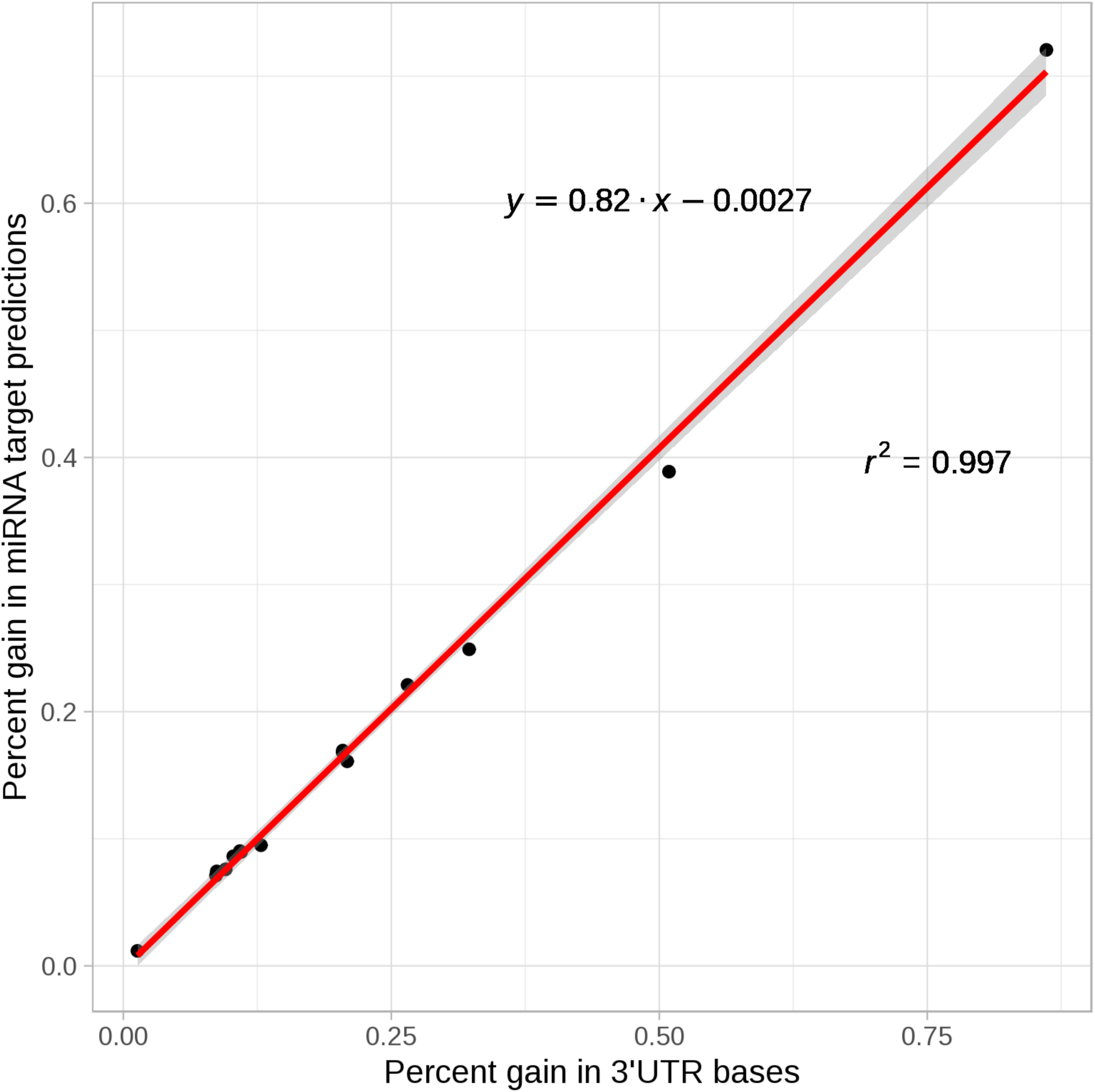
A scatter plot of the percentage gain in total miRNA target site predictions vs. percentage gain in 3’UTR bases for a number of cell lines and tissue datasets analysed (black dots). A linear regression model was fitted using the ‘lm’ function of the R stats package (red) with a 95% confidence interval (grey). R-squared is derived from the Pearson correlation coefficient.

**Supplementary Figure 6b:**
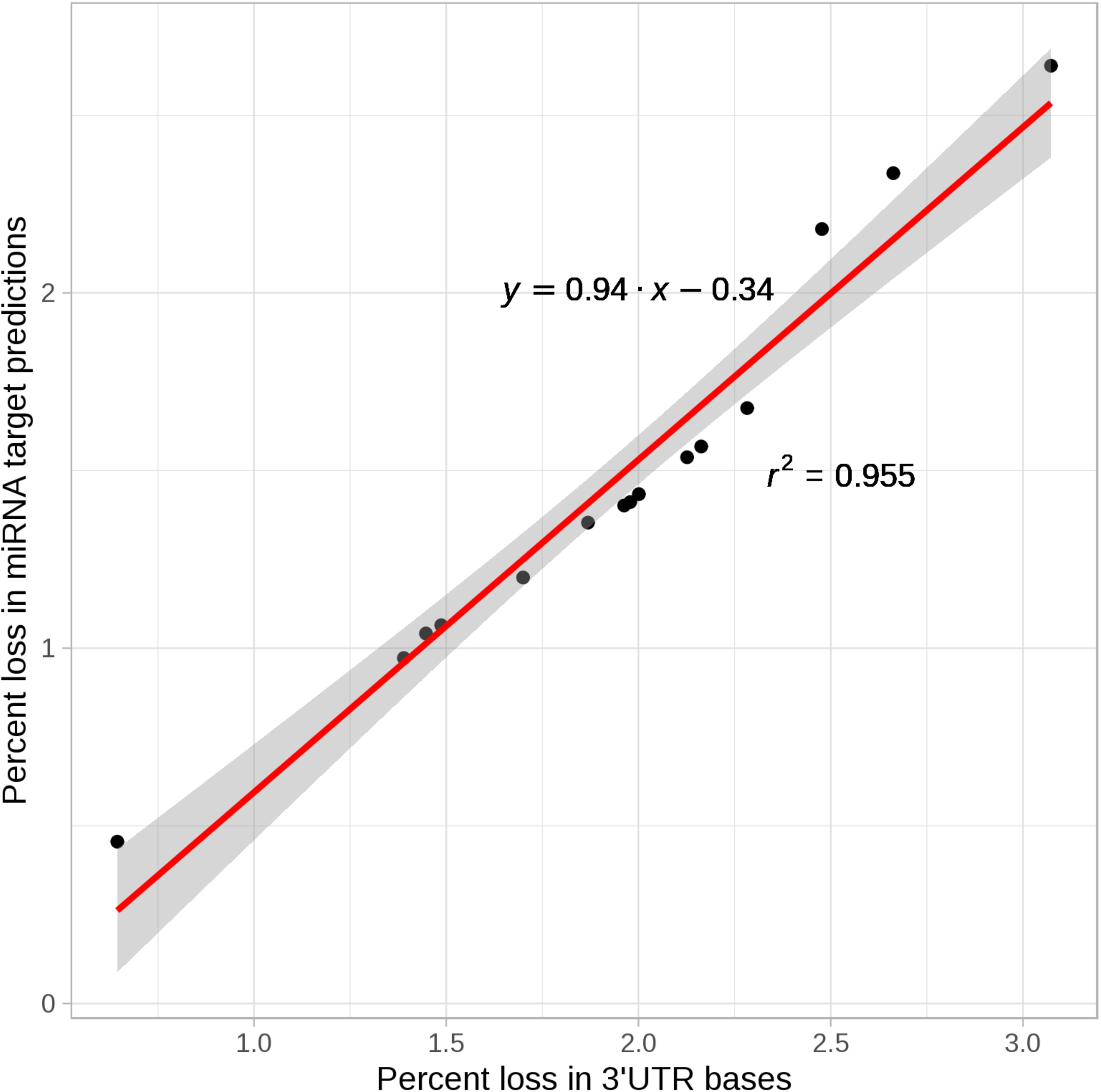
A scatter plot of the percentage loss in total miRNA target predictions vs. percentage loss in total 3’UTR bases. Otherwise as in supplementary figure 6a.

**Supplementary Figure 7:**
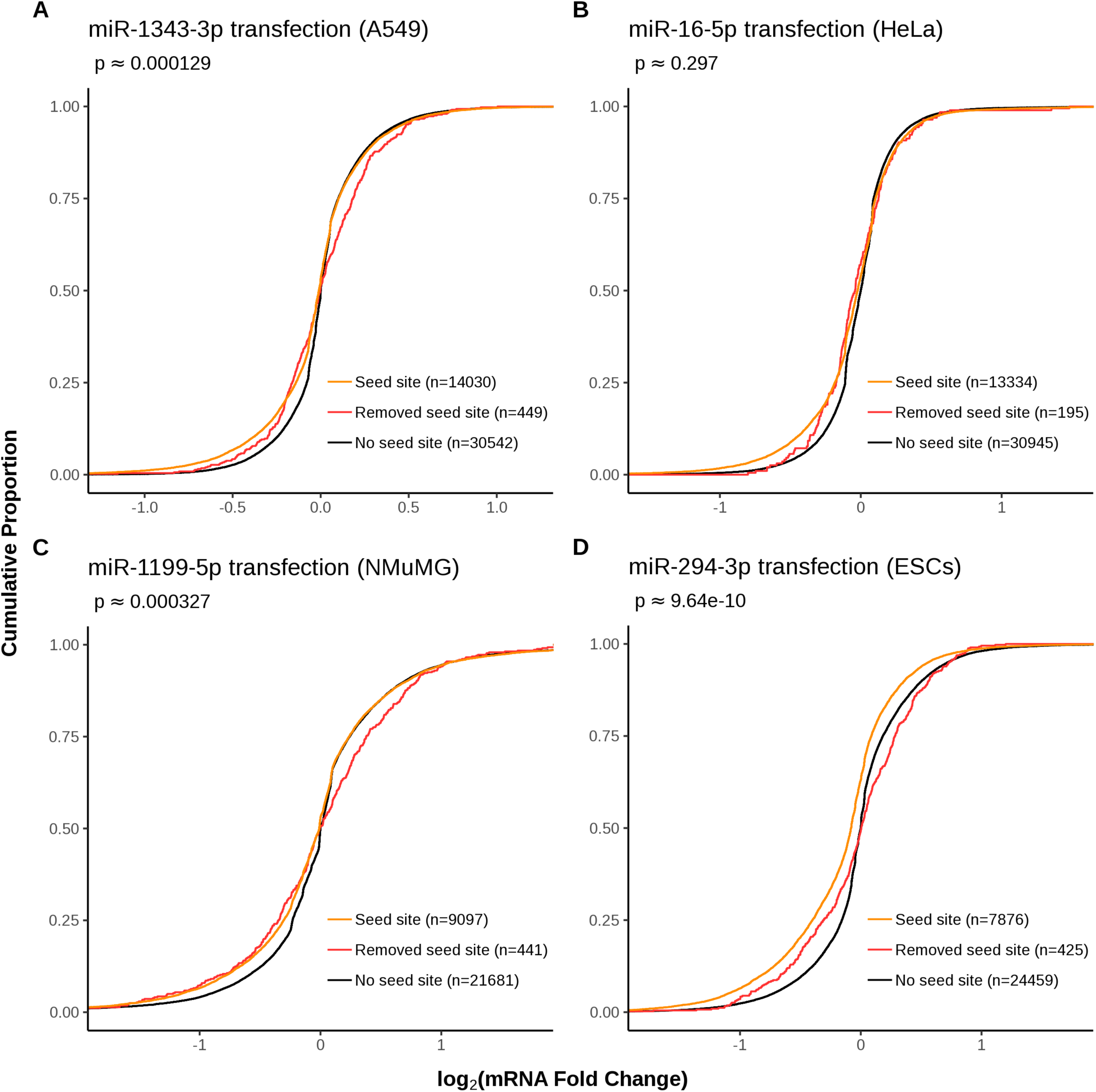
As in figure 3, with the exception that no expression threshold has been implemented.

**Supplementary Table 1:** FilTar 3’UTR reannotation summary statistics for cell line and tissue data used in this study. Statistics are the totial number or proportion of bases or transcripts gained or lost through 3’UTR reannotation respectively. All comparisons are made against a reference of Ensembl annotated 3’UTR sequences associated exclusively with protein-coding mRNA transcripts

| Species | Samples | Bases gained (Mb) | Bases gained (%) | Bases lost (Mb) | Bases lost (%) | 3' UTRs elongated | 3' UTRs elongated (%) | 3'UTRs truncated | 3' UTRs truncated (%) |
| --- | --- | --- | --- | --- | --- | --- | --- | --- | --- |
| *Homo sapiens* | U251 | 0.08 | 0.1 | 1.30 | 2.1 | 352 | 0.7 | 5730 | 10.6 |
|  | U343 | 0.07 | 0.1 | 1.32 | 2.2 | 296 | 0.5 | 7395 | 13.7 |
|  | Du145 | 0.06 | 0.1 | 1.40 | 2.3 | 453 | 0.8 | 5342 | 9.9 |
|  | A549 | 0.13 | 0.2 | 1.04 | 1.7 | 281 | 0.5 | 6774 | 12.5 |
|  | 16HBE14o- | 0.07 | 0.1 | 1.21 | 2.0 | 213 | 0.4 | 6600 | 12.2 |
|  | HeLa | 0.05 | 0.1 | 1.14 | 1.9 | 289 | 0.5 | 4087 | 7.6 |
|  | U20S | 0.01 | 0.0 | 1.23 | 2.0 | 120 | 0.2 | 3614 | 6.7 |
|  | Kidney | 0.20 | 0.3 | 0.91 | 1.5 | 708 | 1.3 | 5738 | 10.6 |
|  | Lung | 0.13 | 0.2 | 0.89 | 1.4 | 538 | 1.0 | 5686 | 10.5 |
|  | Skeletal muscle | 0.05 | 0.1 | 0.39 | 0.6 | 136 | 0.3 | 3018 | 5.6 |
|  | Thyroid | 0.31 | 0.5 | 1.20 | 2.0 | 460 | 0.9 | 7356 | 13.6 |
|  | Bone marrow | 0.13 | 0.2 | 0.85 | 1.4 | 292 | 0.5 | 5444 | 10.1 |
| *Mus musculus* | NMuMG | 0.13 | 0.3 | 1.18 | 2.5 | 454 | 1.1 | 6440 | 15.8 |
|  | CD4+ | 0.05 | 0.1 | 1.27 | 2.7 | 345 | 0.8 | 2447 | 6.0 |
|  | ESCs | 0.41 | 0.9 | 1.46 | 3.1 | 493 | 1.2 | 7502 | 18.4 |

**Supplementary Table 2:** Summary statistics of the effects of filtering protein-coding transcripts at an expression threshold of 0.1 TPM. Statistics are for the total number and proportion of bases and transcripts removed as a result of expression filtering.

| **Species** | **Samples** | **Bases**<br>**lost**<br>**(Mb)** | **Bases**<br>**lost**<br>**(%)** | **3'UTRs**<br>**Removed** | **3' UTRs**<br>**Removed**<br>**(%)** |
| --- | --- | --- | --- | --- | --- |
| *Homo sapiens* | U251 | 19.56 | 32.0 | 22653 | 42.0 |
|  | U343 | 21.07 | 34.4 | 21929 | 40.6 |
|  | Du145 | 24.06 | 39.3 | 25494 | 47.2 |
|  | A549 | 19.50 | 31.9 | 20783 | 38.5 |
|  | 16HBE14o- | 20.73 | 33.9 | 21221 | 39.3 |
|  | HeLa | 15.09 | 24.6 | 18907 | 35.0 |
|  | U20S | 19.96 | 32.6 | 22548 | 41.8 |
|  | Kidney | 17.78 | 29.0 | 22476 | 41.6 |
|  | Lung | 18.68 | 30.5 | 22647 | 42.0 |
|  | Skeletal muscle | 25.86 | 42.2 | 28148 | 52.1 |
|  | Thyroid | 17.84 | 29.1 | 21529 | 39.9 |
|  | Bone marrow | 22.78 | 37.2 | 23040 | 42.7 |
| *Mus musculus* | NMuMG | 20.66 | 43.2 | 19592 | 47.7 |
|  | CD4+ | 18.61 | 38.9 | 19862 | 48.5 |
|  | ESCs | 15.61 | 32.6 | 15505 | 37.9 |

**Supplementary Table 3:** Summed statistics from supplementary table 1 and supplementary table 2 relating to total combined 3’UTR bases and 3’UTRs affected by expression filtering and 3’UTR truncation

| **Species** | **Samples** | **Bases lost (Mb)** | **Bases lost (%)** | **3'UTRs Affected** | **3' UTRs Affected (%)** |
| --- | --- | --- | --- | --- | --- |
| *Homo sapiens* | U251 | 20.86 | 34.1 | 28383 | 52.6 |
|  | U343 | 22.39 | 36.6 | 29324 | 54.3 |
|  | Du145 | 25.45 | 41.6 | 30836 | 57.1 |
|  | A549 | 20.54 | 33.5 | 27557 | 51.1 |
|  | 16HBE14o- | 21.94 | 35.8 | 27821 | 51.5 |
|  | HeLa | 16.23 | 26.5 | 22994 | 42.6 |
|  | U20S | 21.19 | 34.6 | 26162 | 48.5 |
|  | Kidney | 18.69 | 30.5 | 28214 | 52.3 |
|  | Lung | 19.57 | 32.0 | 28333 | 52.5 |
|  | Skeletal muscle | 26.26 | 42.9 | 31166 | 57.7 |
|  | Thyroid | 19.04 | 31.1 | 28885 | 53.5 |
|  | Bone marrow | 23.63 | 38.6 | 28484 | 52.8 |
| *Mus musculus* | NMuMG | 21.84 | 45.7 | 25969 | 63.5 |
|  | CD4+ | 19.87 | 41.6 | 22309 | 54.5 |
|  | ESCs | 17.07 | 35.7 | 23007 | 56.3 |

**Supplementary table 4:** The total number of miRNA seed sites lost through expression filtering of transcripts at TPM > 0.1 or gained and lost through 3’UTR reannotation. Total miRNA seed sites for human: 52084138 and mouse: 28216437

| **Species** | **Samples** | **Seed sites gained**<br>**( 3'UTR**<br>**reannotation)** | **Seed sites lost**<br>**(3'UTR**<br>**reannotation)** | **Seed sites lost**<br>**(expression**<br>**filtering)** |
| --- | --- | --- | --- | --- |
| *Homo sapiens* | U251 | 49345 | 800764 | 12942294 |
|  | U343 | 46701 | 816545 | 13657488 |
|  | Du145 | 39571 | 872804 | 12508511 |
|  | A549 | 87549 | 624503 | 15578814 |
|  | 16HBE14o- | 47031 | 735041 | 13193677 |
|  | HeLa | 38712 | 704948 | 9792951 |
|  | U20S | 6146 | 746686 | 12879630 |
|  | Kidney | 129715 | 554534 | 11476758 |
|  | Lung | 83821 | 542432 | 12057289 |
|  | Skeletal muscle | 37028 | 237223 | 16615464 |
|  | Thyroid | 202504 | 730038 | 11682705 |
|  | Bone marrow | 88212 | 506415 | 14632213 |
| *Mus musculus* | NMuMG | 62367 | 615046 | 9858668 |
|  | CD4+ | 203359 | 744867 | 7358255 |
|  | ESCs | 24318 | 659420 | 8947356 |

**Supplementary Table 5:** A summary of all datasets used in the analyses reported in this study

| Species | BioProject Accession | Source/Study | Sample | Run Accessions |
| --- | --- | --- | --- | --- |
| *Homo sapiens* | PRJNA231155 | Tamim *et al.* 2014 | U251 | SRR1047622,SRR1047623,SRR1047624,SRR1047625 |
|  |  |  | U343 | SRR1047630,SRR1047631,SRR1047632,SRR1047633 |
|  | PRJNA292016 | Liu *et al.* 2017 | Du145 | SRR2146408,SRR2146409,SRR2146410,SRR2146411 |
|  | PRJNA304643 | Stolzenburg *et al.* 2016 | A549 | SRR2968576,SRR2968577,SRR2968578,SRR2968579<br>SRR2968580,SRR2968581,SRR2968582,SRR2968583 |
|  |  |  | 16HBE14o- | SRR2968584,SRR2968586,SRR2968588,SRR2968590<br>SRR2968592,SRR2968594,SRR2968596,SRR2968598 |
|  | PRJNA512378 | Liu *et al.* 2019 | HeLa | SRR8382192,SRR8382193,SRR8382194,SRR8382195<br>SRR8382196,SRR8382197,SRR8382198,SRR8382199<br>SRR8382200,SRR8382201,SRR8382202,SRR8382203<br>SRR8382204,SRR8382205,SRR8382206,SRR8382207<br>SRR8382208,SRR8382209,SRR8382210,SRR8382211<br>SRR8382212,SRR8382213,SRR8382214,SRR8382215<br>SRR8382216,SRR8382217,SRR8382218,SRR8382219<br>SRR8382220,SRR8382221,SRR8382222,SRR8382223<br>SRR8382224,SRR8382225,SRR8382226,SRR8382227<br>SRR8382228,SRR8382229,SRR8382230,SRR8382231<br>SRR8382232,SRR8382233,SRR8382234,SRR8382235<br>SRR8382236,SRR8382237,SRR8382238,SRR8382239<br>SRR8382240,SRR8382241,SRR8382242,SRR8382243 |
|  | PRJNA223608 | Guo *et al.* 2014 | U20S | SRR1598955,SRR1598970,SRR1598976,SRR1598977<br>SRR1598972,SRR1598973 |
|  | PRJEB2445 | Illumina BodyMap2 transcriptome | Kidney | ERR030885,ERR030893 |
|  |  |  | Lung | ERR030879,ERR030896 |
|  | PRJEB6971 | Science for Life Laboratory, Stockholm | Skeletal Muscle | ERR579142,ERR579143 |
|  |  |  | Thyroid | ERR315358,ERR315422 |
|  |  |  | Bone Marrow | ERR315404,ERR315406 |
| *Mus Musculus* | PRJNA340017 | Diepenbruck *et al.* 2017 | NMuMG | SRR4054984,SRR4054985,SRR4054992,SRR4054995<br>SRR4054996,SRR4054999,SRR4055002,SRR4055005 |
|  | PRJNA309441 | Pua *et al.* 2016 | CD4+ | SRR3112249,SRR3112250,SRR3112251,SRR3112252<br>SRR3112245,SRR3112246,SRR3112247,SRR3112248<br>SRR3112237,SRR3112238,SRR3112239,SRR3112240<br>SRR3112241,SRR3112242,SRR3112243,SRR3112244 |
|  | PRJNA270999 | Cao *et al.* 2015 | ESCs | SRR1734389,SRR1734391,SRR1734393,SRR1734395 |

## Supplementary Files

- **Supplementary File 1a:** Cumulative plots of predicted targets and non-targets for datasets which did not pass the QC stage of the study.
- **Supplementary File 1b:** A table of metadata for Supplementary File 1a
- **Supplementary File 2:** A compressed and archived folder of MultiQC Reports
- **Supplementary File 3:** The analysis presented in figure 1 as applied to all datasets
- **Supplementary File 4:** The analysis presented in figure 2 as applied to all datasets
- **Supplementary File 5:** The analysis presented in figure 3 as applied to all available datasets (datasets with insufficiently high number of ‘added targets’ were discarded).

## References

Agarwal, Vikram, George W Bell, Jin-Wu Nam, and David P Bartel. 2015. “Predicting Effective microRNA Target Sites in Mammalian mRNAs.” Journal Article. eLife 4. doi:10.7554/eLife.05005.

Aken, Bronwen L, Sarah Ayling, Daniel Barrell, Laura Clarke, Valery Curwen, Susan Fairley, Julio Fernandez Banet, Konstantinos Billis, Carlos García Girón, and Thibaut Hourlier. 2016. “The Ensembl Gene Annotation System.” Journal Article. Database 2016.

Andrews, S. (2010). FastQC: a quality control tool for high throughput sequence data.

Bartel, David P. 2018. “Metazoan Micrornas.” Journal Article. Cell 173 (1): 20–51.

Birney, E., Andrews, T. D., Bevan, P., Caccamo, M., Chen, Y., Clarke, L., … & Down, T. (2004). An overview of Ensembl. Genome research, 14(5), 925–928.

Blanchette, Mathieu, W James Kent, Cathy Riemer, Laura Elnitski, Arian FA Smit, Krishna M Roskin, Robert Baertsch, Kate Rosenbloom, Hiram Clawson, and Eric D Green. 2004. “Aligning Multiple Genomic Sequences with the Threaded Blockset Aligner.” Journal Article. Genome Research 14 (4): 708–15.

Bray, Nicolas L, Harold Pimentel, Páll Melsted, and Lior Pachter. 2016. “Near-Optimal Probabilistic Rna-Seq Quantification.” Journal Article. Nature Biotechnology 34 (5): 525.

Cao, Y., Guo, W. T., Tian, S., He, X., Wang, X. W., Liu, X., … & Cai, Y. (2015). miR-290/371-Mbd2-Myc circuit regulates glycolytic metabolism to promote pluripotency. The EMBO journal, 34(5), 609–623.

Chi, Sung Wook, Julie B Zang, Aldo Mele, and Robert B Darnell. 2009. “Argonaute Hits-Clip Decodes microRNA–mRNA Interaction Maps.” Journal Article. Nature 460 (7254): 479–86. http://www.nature.com/nature/journal/v460/n7254/pdf/nature08170.pdf.

Cock, Peter JA, Tiago Antao, Jeffrey T Chang, Brad A Chapman, Cymon J Cox, Andrew Dalke, Iddo Friedberg, Thomas Hamelryck, Frank Kauff, and Bartek Wilczynski. 2009. “Biopython: Freely Available Python Tools for Computational Molecular Biology and Bioinformatics.” Journal Article. Bioinformatics 25 (11): 1422–3.

Cunningham, Fiona, Premanand Achuthan, Wasiu Akanni, James Allen, M Ridwan Amode, Irina M Armean, Ruth Bennett, Jyothish Bhai, Konstantinos Billis, and Sanjay Boddu. 2018. “Ensembl 2019.” Journal Article. Nucleic Acids Research 47 (D1): D745–D751.

Diepenbruck, M., Tiede, S., Saxena, M., Ivanek, R., Kalathur, R. K. R., Lüönd, F., … & Christofori, G. (2017). miR-1199-5p and Zeb1 function in a double-negative feedback loop potentially coordinating EMT and tumour metastasis. Nature communications, 8(1), 1168.

Enright, A. J., B. John, U. Gaul, T. Tuschl, C. Sander, and D. S. Marks. 2003. “MicroRNA Targets in Drosophila.” Journal Article. Genome Biol 5 (1): R1. doi:10.1186/gb-2003-5-1-r1.

Ewels, Philip, Måns Magnusson, Sverker Lundin, and Max Käller. 2016. “MultiQC: Summarize Analysis Results for Multiple Tools and Samples in a Single Report.” Journal Article. Bioinformatics 32 (19): 3047–8.

Friedman, Robin C., Kyle Kai-How Farh, Christopher B. Burge, and David P. Bartel. 2009. “Most Mammalian mRNAs Are Conserved Targets of microRNAs.” Journal Article. Genome Research 19 (1): 92–105. doi:10.1101/gr.082701.108.

Garcia, David M, Daehyun Baek, Chanseok Shin, George W Bell, Andrew Grimson, and David P Bartel. 2011. “Weak Seed-Pairing Stability and High Target-Site Abundance Decrease the Proficiency of Lsy-6 and Other microRNAs.” Journal Article. Nature Structural & Molecular Biology 18 (10): 1139–46.

Griffiths-Jones, S. (2004). The microRNA registry. Nucleic acids research, 32(suppl_1), D109–D111.

Grimson, Andrew, Kyle Kai-How Farh, Wendy K Johnston, Philip Garrett-Engele, Lee P Lim, and David P Bartel. 2007. “MicroRNA Targeting Specificity in Mammals: Determinants Beyond Seed Pairing.” Journal Article. Molecular Cell 27 (1): 91–105.

Gruber, A. J., Schmidt, R., Ghosh, S., Martin, G., Gruber, A. R., van Nimwegen, E., & Zavolan, M. (2018). Discovery of physiological and cancer-related regulators of 3′ UTR processing with KAPAC. Genome biology, 19(1), 44.

Gruber, A. J., Gypas, F., Riba, A., Schmidt, R., & Zavolan, M. (2018). Terminal exon characterization with TECtool reveals an abundance of cell-specific isoforms. Nature methods, 15(10), 832.

Gumienny, Rafal, and Mihaela Zavolan. 2015. “Accurate Transcriptome-Wide Prediction of microRNA Targets and Small Interfering Rna Off-Targets with Mirza-G.” Journal Article. Nucleic Acids Research 43 (3): 1380–91. doi:10.1093/nar/gkv050.

Guo, J. U., Agarwal, V., Guo, H., & Bartel, D. P. (2014). Expanded identification and characterization of mammalian circular RNAs. Genome biology, 15(7), 409.

Harrison, P. W., Alako, B., Amid, C., Cerdeño-Tárraga, A., Cleland, I., Holt, S., … & Leinonen, R. (2018). The European Nucleotide Archive in 2018. Nucleic acids research, 47(D1), D84–D88.

Helwak, A., & Tollervey, D. (2014). Mapping the miRNA interactome by cross-linking ligation and sequencing of hybrids (CLASH). Nature protocols, 9(3), 711.

John, Bino, Anton J Enright, Alexei Aravin, Thomas Tuschl, Chris Sander, and Debora S Marks. 2004. “Human microRNA Targets.” Journal Article. PLoS Biol 2 (11): e363. http://www.ncbi.nlm.nih.gov/pmc/articles/PMC521178/pdf/pbio.0020363.pdf.

Karagkouni, D., Paraskevopoulou, M. D., Chatzopoulos, S., Vlachos, I. S., Tastsoglou, S., Kanellos, I., … & Vergoulis, T. (2017). DIANA-TarBase v8: a decade-long collection of experimentally supported miRNA–gene interactions. Nucleic acids research, 46(D1), D239–D245.

Kent, W James, Charles W Sugnet, Terrence S Furey, Krishna M Roskin, Tom H Pringle, Alan M Zahler, and David Haussler. 2002. “The Human Genome Browser at Ucsc.” Journal Article. Genome Research 12 (6): 996–1006.

Khorshid, Mohsen, Jean Hausser, Mihaela Zavolan, and Erik van Nimwegen. 2013. “A Biophysical miRNA-mRNA Interaction Model Infers Canonical and Noncanonical Targets.” Journal Article. Nature Methods 10 (3): 253–55. http://www.nature.com/nmeth/journal/v10/n3/pdf/nmeth.2341.pdf.

Kim, Daehwan, Ben Langmead, and Steven L Salzberg. 2015. “HISAT: A Fast Spliced Aligner with Low Memory Requirements.” Journal Article. Nature Methods 12 (4): 357.

Kiriakidou, Marianthi, Peter T Nelson, Andrei Kouranov, Petko Fitziev, Costas Bouyioukos, Zissimos Mourelatos, and Artemis Hatzigeorgiou. 2004. “A Combined Computational-Experimental Approach Predicts Human microRNA Targets.” Journal Article. Genes & Development 18 (10): 1165–78.

Kozomara, Ana, Maria Birgaoanu, and Sam Griffiths-Jones. 2018. “MiRBase: From microRNA Sequences to Function.” Journal Article. Nucleic Acids Research 47 (D1): D155–D162. doi:10.1093/nar/gky1141.

König, Julian, Kathi Zarnack, Gregor Rot, Tomaž Curk, Melis Kayikci, Blaž Zupan, Daniel J Turner, Nicholas M Luscombe, and Jernej Ule. 2010. “ICLIP Reveals the Function of hnRNP Particles in Splicing at Individual Nucleotide Resolution.” Journal Article. Nature Structural & Molecular Biology 17 (7): 909–15.

Köster, Johannes, and Sven Rahmann. 2012. “Snakemake—a Scalable Bioinformatics Workflow Engine.” Journal Article. Bioinformatics 28 (19): 2520–2.

Krueger, Felix. 2015. “Trim Galore.” Journal Article. A Wrapper Tool Around Cutadapt and FastQC to Consistently Apply Quality and Adapter Trimming to FastQ Files.

Kudla, G., Granneman, S., Hahn, D., Beggs, J. D., & Tollervey, D. (2011). Cross-linking, ligation, and sequencing of hybrids reveals RNA–RNA interactions in yeast. Proceedings of the National Academy of Sciences, 108(24), 10010–10015.

Kuhn, Donald E, Mickey M Martin, David S Feldman, Alvin V Terry Jr, Gerard J Nuovo, and Terry S Elton. 2008. “Experimental Validation of miRNA Targets.” Journal Article. Methods 44 (1): 47–54.

Leek, J. T., Scharpf, R. B., Bravo, H. C., Simcha, D., Langmead, B., Johnson, W. E., … & Irizarry, A. (2010). Tackling the widespread and critical impact of batch effects in high-throughput data. Nature Reviews Genetics, 11(10), 733.

Leinonen, R., Akhtar, R., Birney, E., Bower, L., Cerdeno-Tárraga, A., Cheng, Y., … & Hoad, G. (2010). The European nucleotide archive. Nucleic acids research, 39(suppl_1), D28–D31.

Leinonen, R., Sugawara, H., Shumway, M., & International Nucleotide Sequence Database Collaboration. (2010). The sequence read archive. Nucleic acids research, 39(suppl_1), D19–D21.

Lewis, Benjamin P, I-hung Shih, Matthew W Jones-Rhoades, David P Bartel, and Christopher B Burge. 2003. “Prediction of Mammalian microRNA Targets.” Journal Article. Cell 115 (7): 787–98.

Lewis, B. P., C. B. Burge, and D. P. Bartel. 2005. “Conserved Seed Pairing, Often Flanked by Adenosines, Indicates That Thousands of Human Genes Are microRNA Targets.” Journal Article. Cell 120 (1): 15–20. doi:10.1016/j.cell.2004.12.035.

Li, B., Ruotti, V., Stewart, R. M., Thomson, J. A., & Dewey, C. N. (2009). RNA-Seq gene expression estimation with read mapping uncertainty. Bioinformatics, 26(4), 493–500.

Li, Heng, Bob Handsaker, Alec Wysoker, Tim Fennell, Jue Ruan, Nils Homer, Gabor Marth, Goncalo Abecasis, and Richard Durbin. 2009. “The Sequence Alignment/Map Format and Samtools.” Journal Article. Bioinformatics 25 (16): 2078–9.

Li, H. (2012). seqtk Toolkit for processing sequences in FASTA/Q formats.

Liu, C., Liu, R., Zhang, D., Deng, Q., Liu, B., Chao, H. P., … & Zhong, Y. (2017). MicroRNA-141 suppresses prostate cancer stem cells and metastasis by targeting a cohort of pro-metastasis genes. Nature communications, 8, 14270.

Liu, W., & Wang, X. (2019). Prediction of functional microRNA targets by integrative modeling of microRNA binding and target expression data. Genome biology, 20(1), 18.

Love, Michael I, Wolfgang Huber, and Simon Anders. 2014. “Moderated Estimation of Fold Change and Dispersion for Rna-Seq Data with Deseq2.” Journal Article. Genome Biology 15 (12): 550.

Martin, Marcel. 2011. “Cutadapt Removes Adapter Sequences from High-Throughput Sequencing Reads.” Journal Article. EMBnet. Journal 17 (1): 10–12.

Mayr, Christine, and David P Bartel. 2009. “Widespread Shortening of 3′ Utrs by Alternative Cleavage and Polyadenylation Activates Oncogenes in Cancer Cells.” Journal Article. Cell 138 (4): 673–84.

Miura, Pedro, Sol Shenker, Celia Andreu-Agullo, Jakub O Westholm, and Eric C Lai. 2013. “Widespread and Extensive Lengthening of 3′ Utrs in the Mammalian Brain.” Journal Article. Genome Research 23 (5): 812–25.

Nam, J. W., O. S. Rissland, D. Koppstein, C. Abreu-Goodger, C. H. Jan, V. Agarwal, M. A. Yildirim, A. Rodriguez, and D. P. Bartel. 2014. “Global Analyses of the Effect of Different Cellular Contexts on microRNA Targeting.” Journal Article. Mol Cell 53 (6): 1031–43. doi:10.1016/j.molcel.2014.02.013.

Patro, Rob, Geet Duggal, Michael I Love, Rafael A Irizarry, and Carl Kingsford. 2017. “Salmon Provides Fast and Bias-Aware Quantification of Transcript Expression.” Journal Article. Nature Methods 14 (4): 417.

Pruitt, Kim D, Tatiana Tatusova, and Donna R Maglott. 2006. “NCBI Reference Sequences (Refseq): A Curated Non-Redundant Sequence Database of Genomes, Transcripts and Proteins.” Journal Article. Nucleic Acids Research 35 (suppl_1): D61–D65.

Pruitt, K. D., Brown, G. R., Hiatt, S. M., Thibaud-Nissen, F., Astashyn, A., Ermolaeva, O., … & Murphy, M. R. (2013). RefSeq: an update on mammalian reference sequences. Nucleic acids research, 42(D1), D756–D763.

Pua, H. H., Steiner, D. F., Patel, S., Gonzalez, J. R., Ortiz-Carpena, J. F., Kageyama, R., … & McManus, M. T. (2016). MicroRNAs 24 and 27 suppress allergic inflammation and target a network of regulators of T helper 2 cell-associated cytokine production. Immunity, 44(4), 821–832.

Quinlan, Aaron R, and Ira M Hall. 2010. “BEDTools: A Flexible Suite of Utilities for Comparing Genomic Features.” Journal Article. Bioinformatics 26 (6): 841–42.

Quinlan, Aaron R. 2014. “BEDTools: The Swiss-army Tool for Genome Feature Analysis.” Journal Article. Current Protocols in Bioinformatics 47 (1): 11.12. 1–11.12. 34.

Reczko, Martin, Manolis Maragkakis, Panagiotis Alexiou, Ivo Grosse, and Artemis G Hatzigeorgiou. 2012. “Functional microRNA Targets in Protein Coding Sequences.” Journal Article. Bioinformatics 28 (6): 771–76. http://bioinformatics.oxfordjournals.org/content/28/6/771.full.pdf.

Ritchie, W., and J. E. Rasko. 2014. “Refining microRNA Target Predictions: Sorting the Wheat from the Chaff.” Journal Article. Biochem Biophys Res Commun 445 (4): 780–4. doi:10.1016/j.bbrc.2014.01.181.

Schneider, V. A., Graves-Lindsay, T., Howe, K., Bouk, N., Chen, H. C., Kitts, P. A., … & Fulton, R. (2017). Evaluation of GRCh38 and de novo haploid genome assemblies demonstrates the enduring quality of the reference assembly. Genome research, 27(5), 849–864.

Soneson, Charlotte, Michael I Love, and Mark D Robinson. 2015. “Differential Analyses for Rna-Seq: Transcript-Level Estimates Improve Gene-Level Inferences.” Journal Article. F1000Research 4.

Stolzenburg, L. R., Wachtel, S., Dang, H., & Harris, A. (2016). miR-1343 attenuates pathways of fibrosis by targeting the TGF-β receptors. Biochemical Journal, 473(3), 245–256.

Team, R Core. 2013. “R: A Language and Environment for Statistical Computing.” Journal Article.

Tian, Bin, and James L Manley. 2017. “Alternative Polyadenylation of mRNA Precursors.” Journal Article. Nature Reviews Molecular Cell Biology 18 (1): 18.

Tamim, S., Vo, D. T., Uren, P. J., Qiao, M., Bindewald, E., Kasprzak, W. K., … & Nakano, I. (2014). Genomic analyses reveal broad impact of miR-137 on genes associated with malignant transformation and neuronal differentiation in glioblastoma cells. PloS one, 9(1), e85591.

Van Nostrand, Eric L, Gabriel A Pratt, Alexander A Shishkin, Chelsea Gelboin-Burkhart, Mark Y Fang, Balaji Sundararaman, Steven M Blue, Thai B Nguyen, Christine Surka, and Keri Elkins. 2016. “Robust Transcriptome-Wide Discovery of Rna-Binding Protein Binding Sites with Enhanced Clip (eCLIP).” Journal Article. Nature Methods.

Wang, Li, and Rui Yi. 2014. “3′ Utrs Take a Long Shot in the Brain.” Journal Article. Bioessays 36 (1): 39–45.

Wang, Xiaowei. 2016. “Improving microRNA Target Prediction by Modeling with Unambiguously Identified microRNA-Target Pairs from Clip-Ligation Studies.” Journal Article. Bioinformatics, btw002.

Wickham, H. (2016). ggplot2: elegant graphics for data analysis. Springer.

Ye, C., Long, Y., Ji, G., Li, Q. Q., & Wu, X. (2018). APAtrap: identification and quantification of alternative polyadenylation sites from RNA-seq data. Bioinformatics, 34(11), 1841–1849.

