## Supplementary File 1a for "FilTar: Using RNA-Seq data to improve microRNA target prediction accuracy in animals"

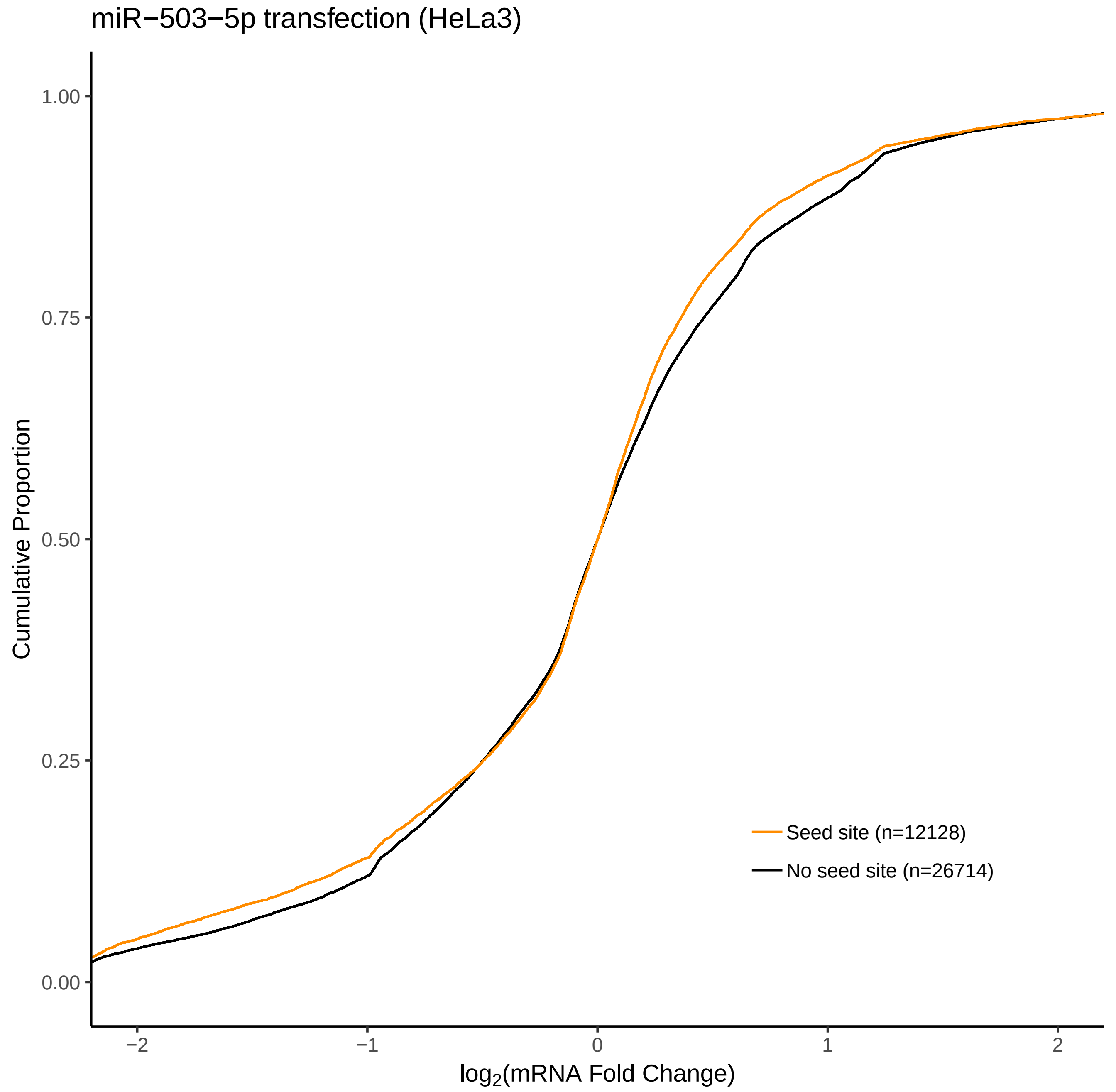

miR−503−5p transfection (HeLa3)
Cumulative Proportion
1.00
0.75
0.50
0.25
0.00
−2
−1
0
1
2
log2(mRNA Fold Change)
Seed site (n=12128)
No seed site (n=26714)

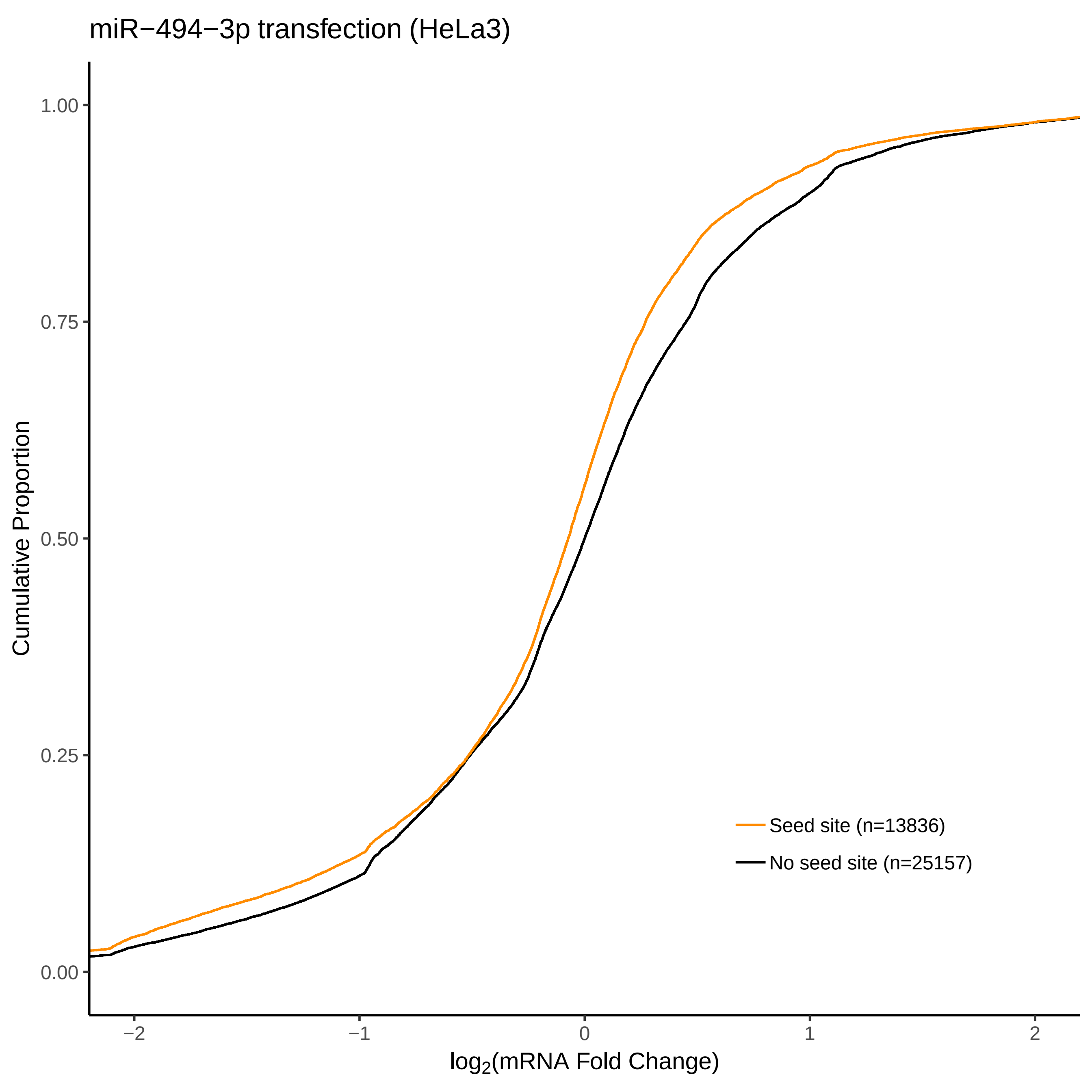

miR−494−3p transfection (HeLa3)
Cumulative Proportion
1.00
0.75
0.50
0.25
0.00
−2
−1
0
1
2
log2(mRNA Fold Change)
Seed site (n=13836)
No seed site (n=25157)

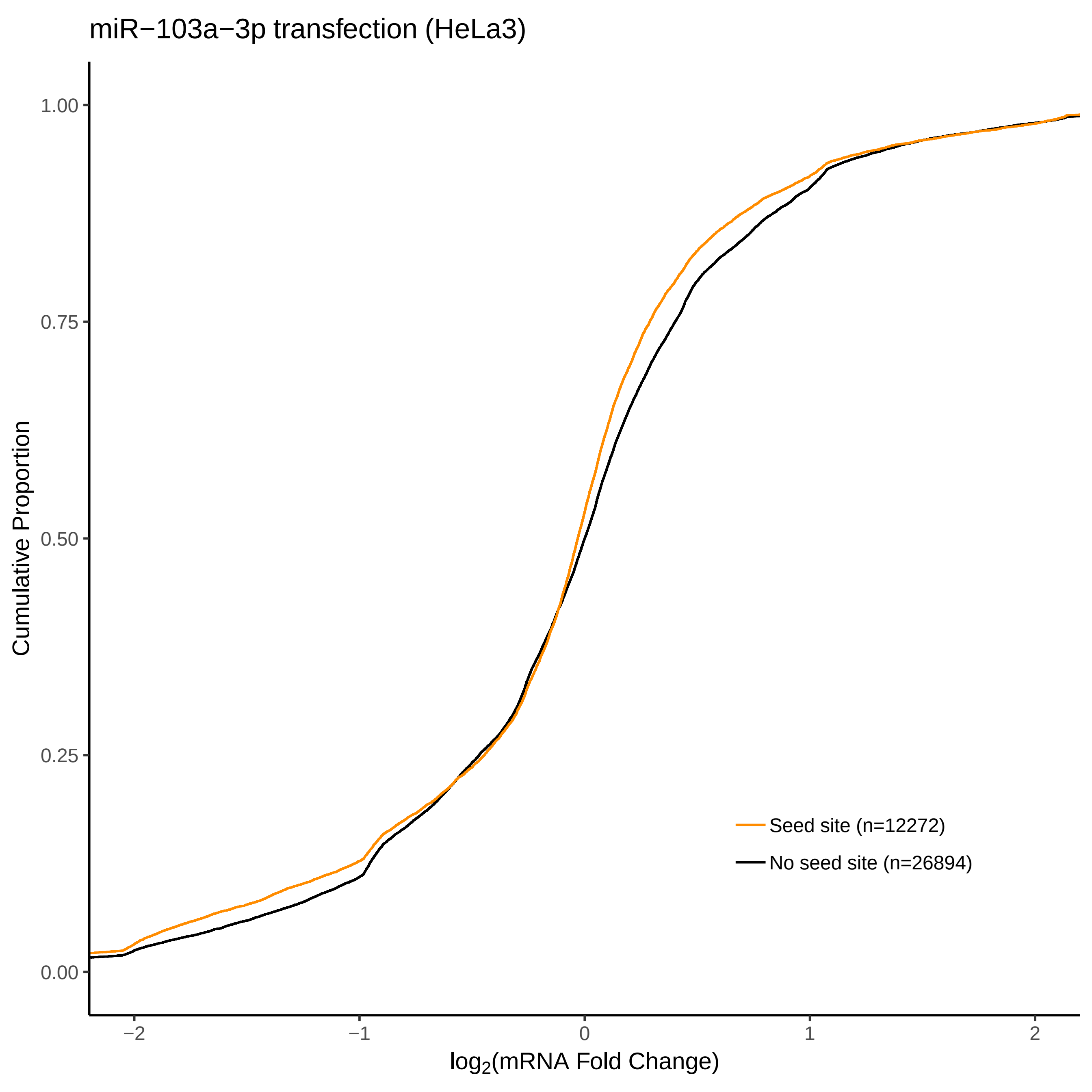

miR−103a−3p transfection (HeLa3)
Cumulative Proportion
1.00
0.75
0.50
0.25
0.00
−2
−1
0
1
2
log2(mRNA Fold Change)
Seed site (n=12272)
No seed site (n=26894)

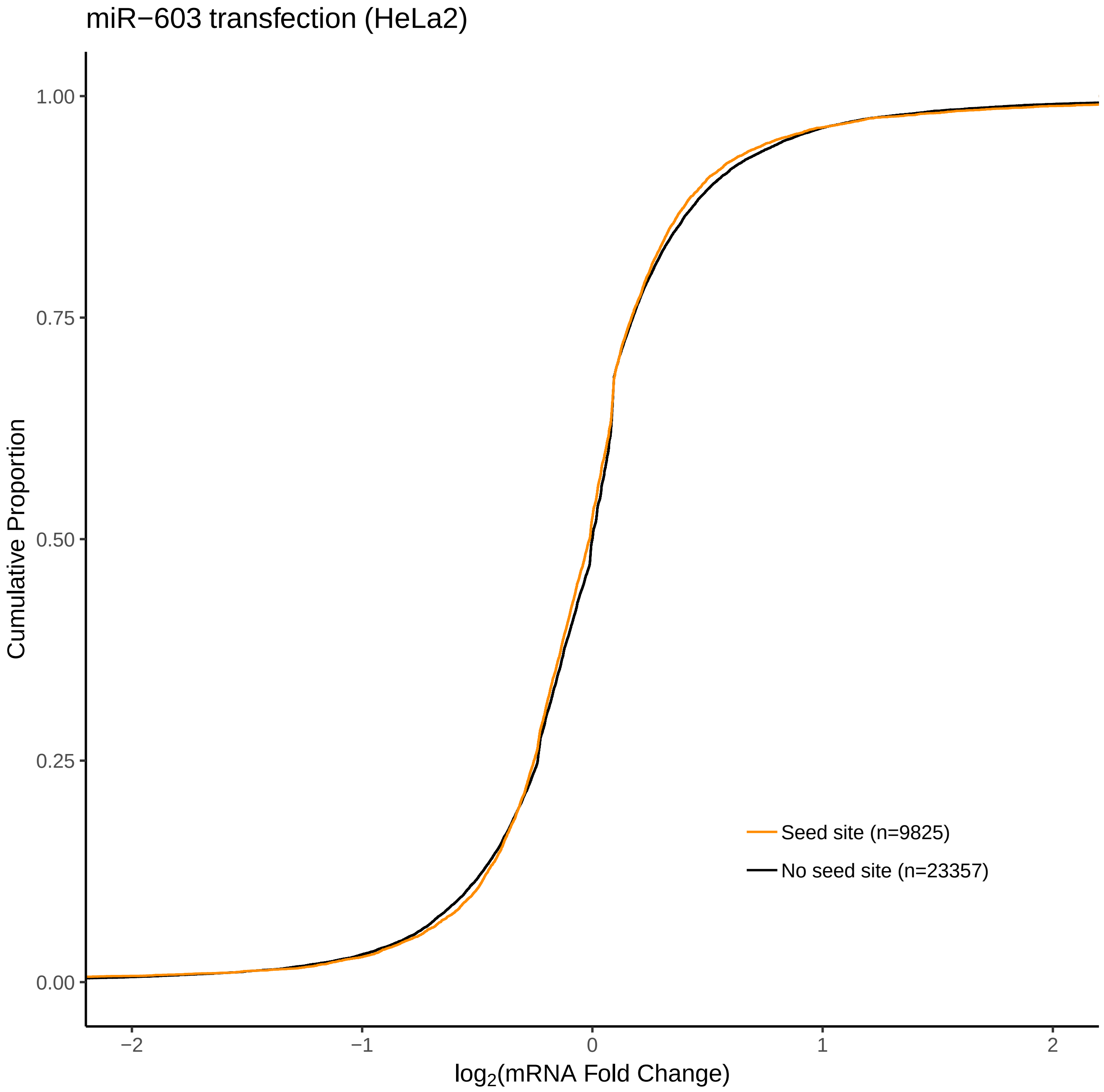
miR−603 transfection (HeLa2)
Cumulative Proportion
1.00
0.75
0.50
0.25
0.00
−2
−1
0
1
2
log2(mRNA Fold Change)
Seed site (n=9825)
No seed site (n=23357)

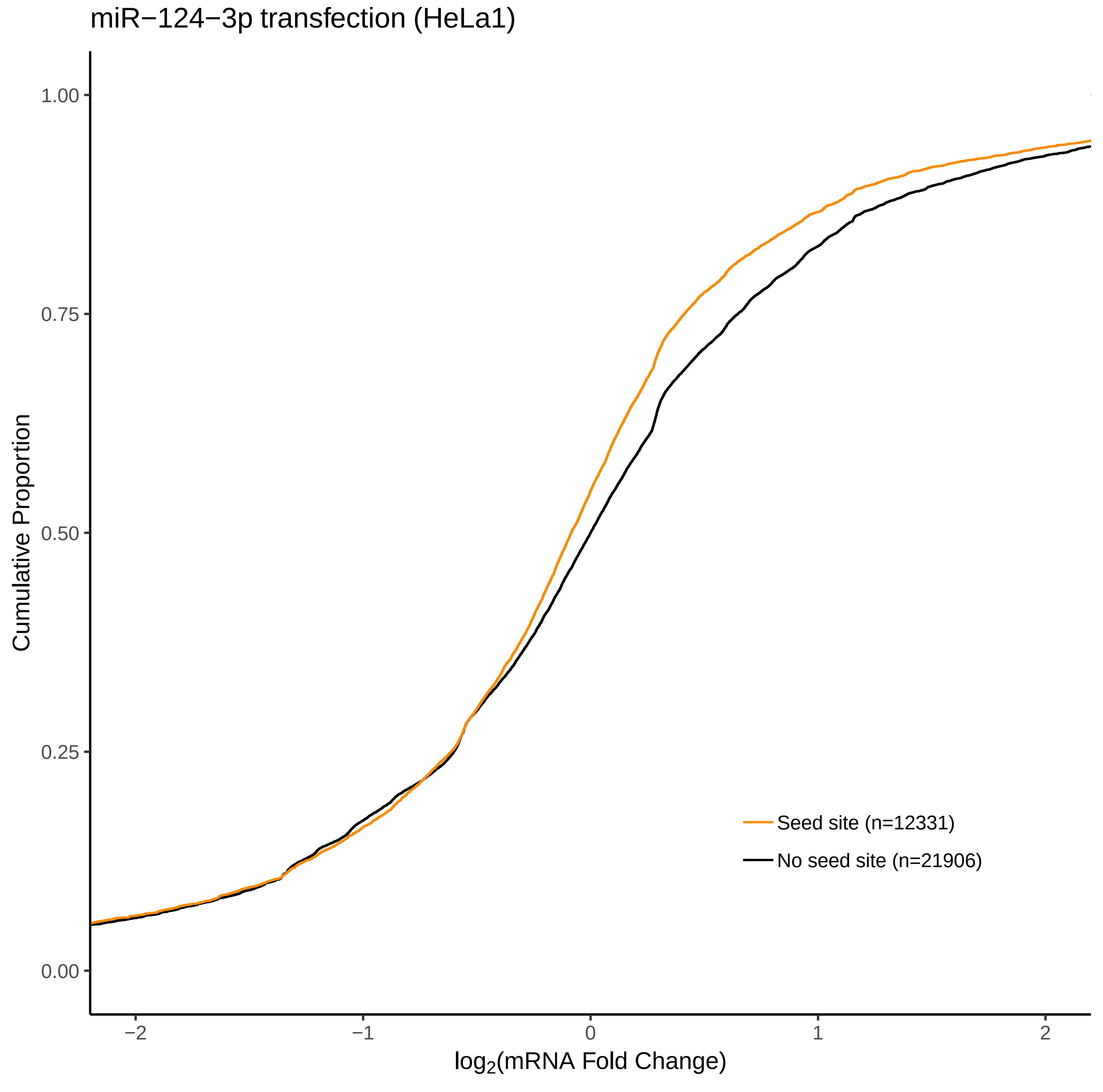

miR−124−3p transfection (HeLa1)
Cumulative Proportion
1.00
0.75
0.50
0.25
0.00
−2
−1
0
1
2
$\log_2$(mRNA Fold Change)
Seed site (n=12331)
No seed site (n=21906)

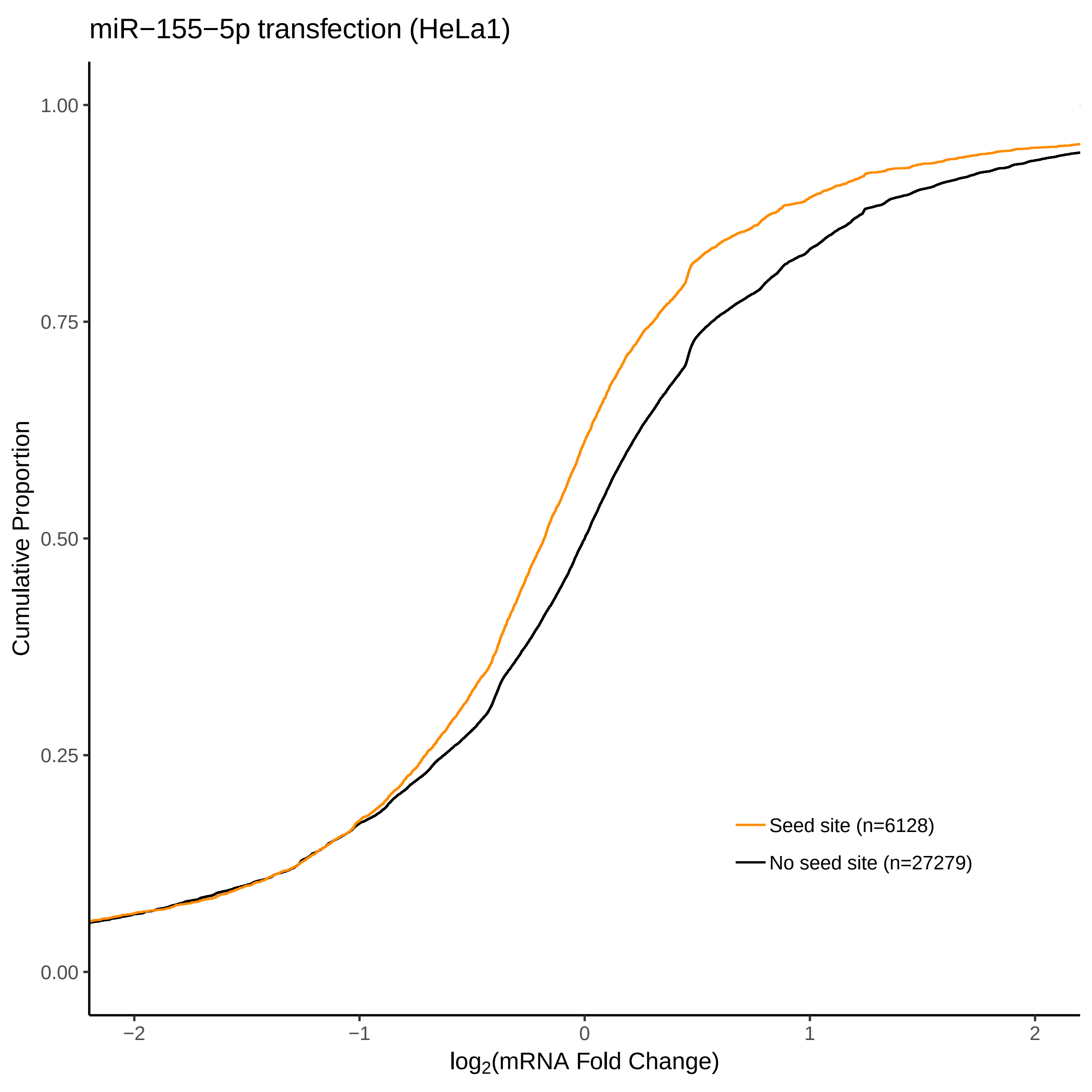
miR−155−5p transfection (HeLa1)
Cumulative Proportion
1.00
0.75
0.50
0.25
0.00
−2
−1
0
1
2
log2(mRNA Fold Change)
Seed site (n=6128)
No seed site (n=27279)

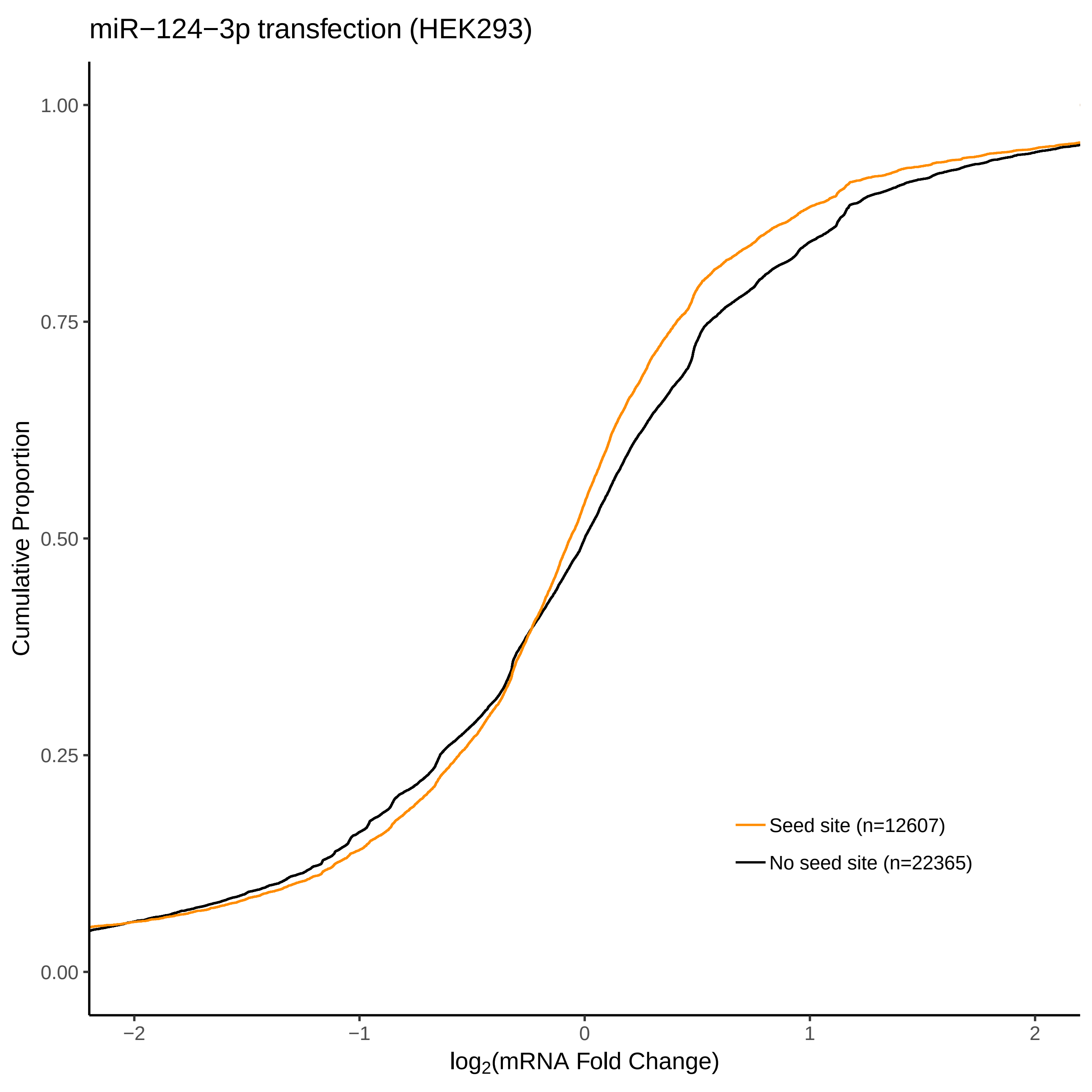
miR−124−3p transfection (HEK293)
Cumulative Proportion
1.00
0.75
0.50
0.25
0.00
−2
−1
0
1
2
log2(mRNA Fold Change)
Seed site (n=12607)
No seed site (n=22365)

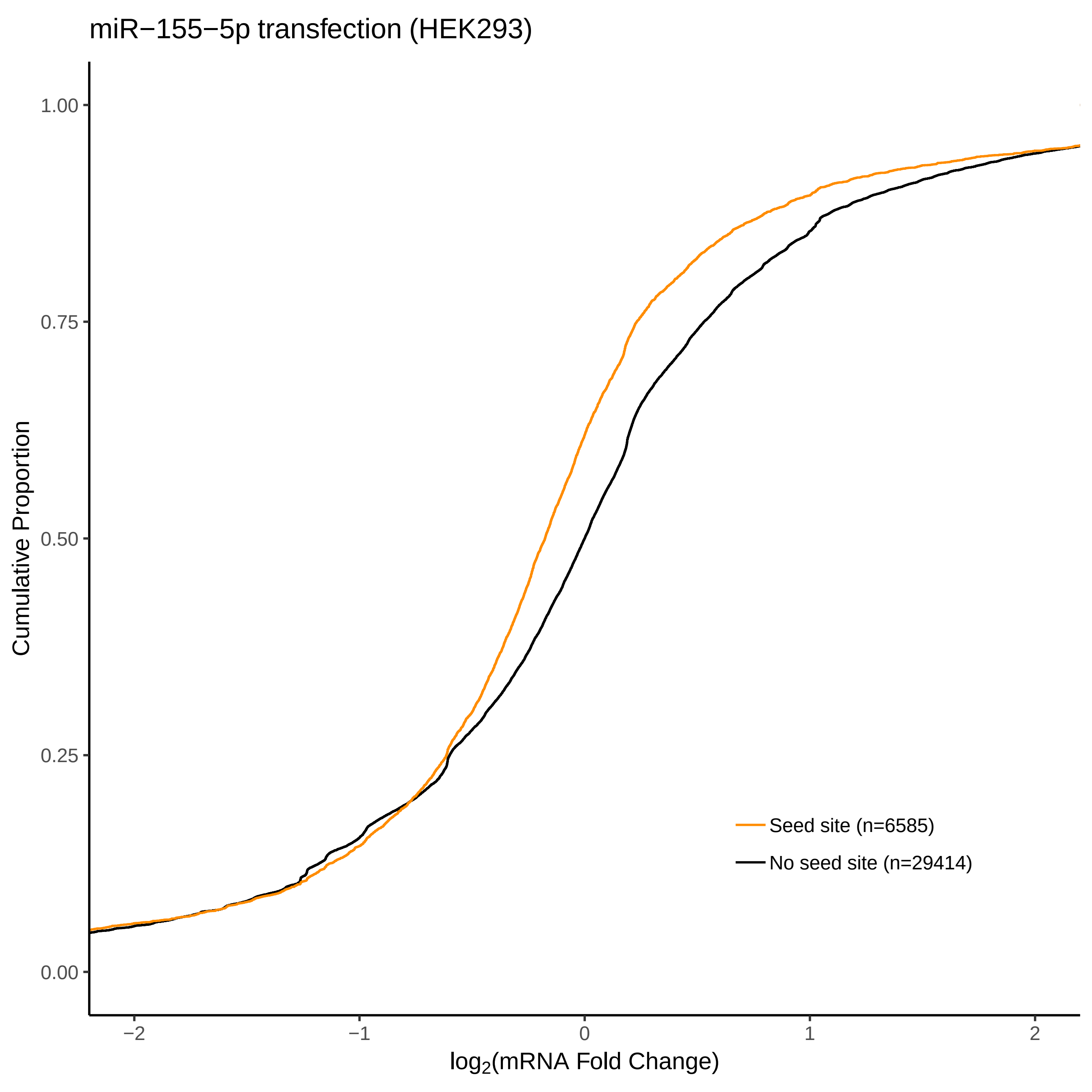
miR−155−5p transfection (HEK293)
Cumulative Proportion
1.00
0.75
0.50
0.25
0.00
−2
−1
0
1
2
$\log_2$(mRNA Fold Change)
Seed site (n=6585)
No seed site (n=29414)

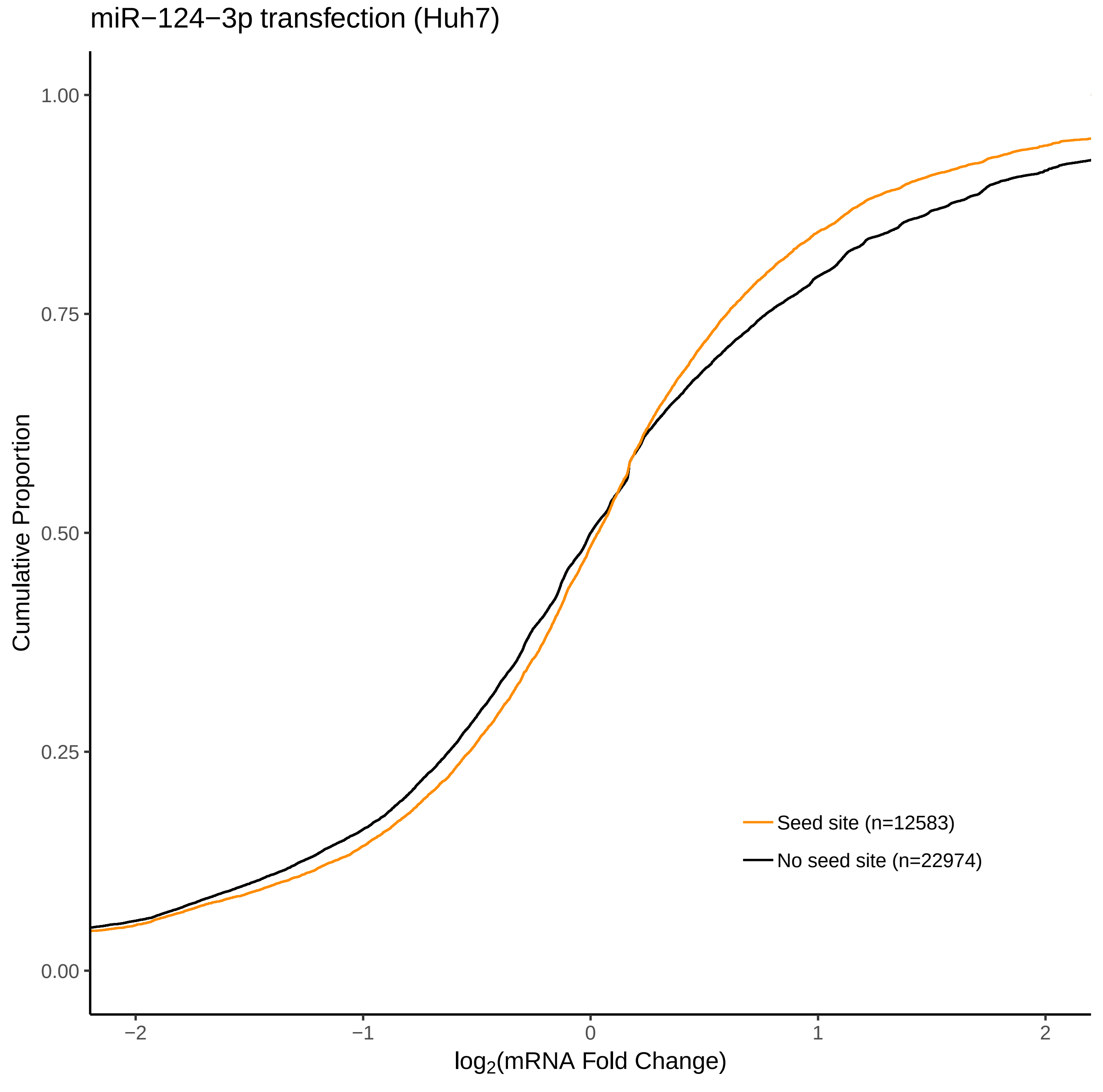
miR−124−3p transfection (Huh7)
Cumulative Proportion
1.00
0.75
0.50
0.25
0.00
−2
−1
0
1
2
$\log_2$(mRNA Fold Change)
Seed site (n=12583)
No seed site (n=22974)

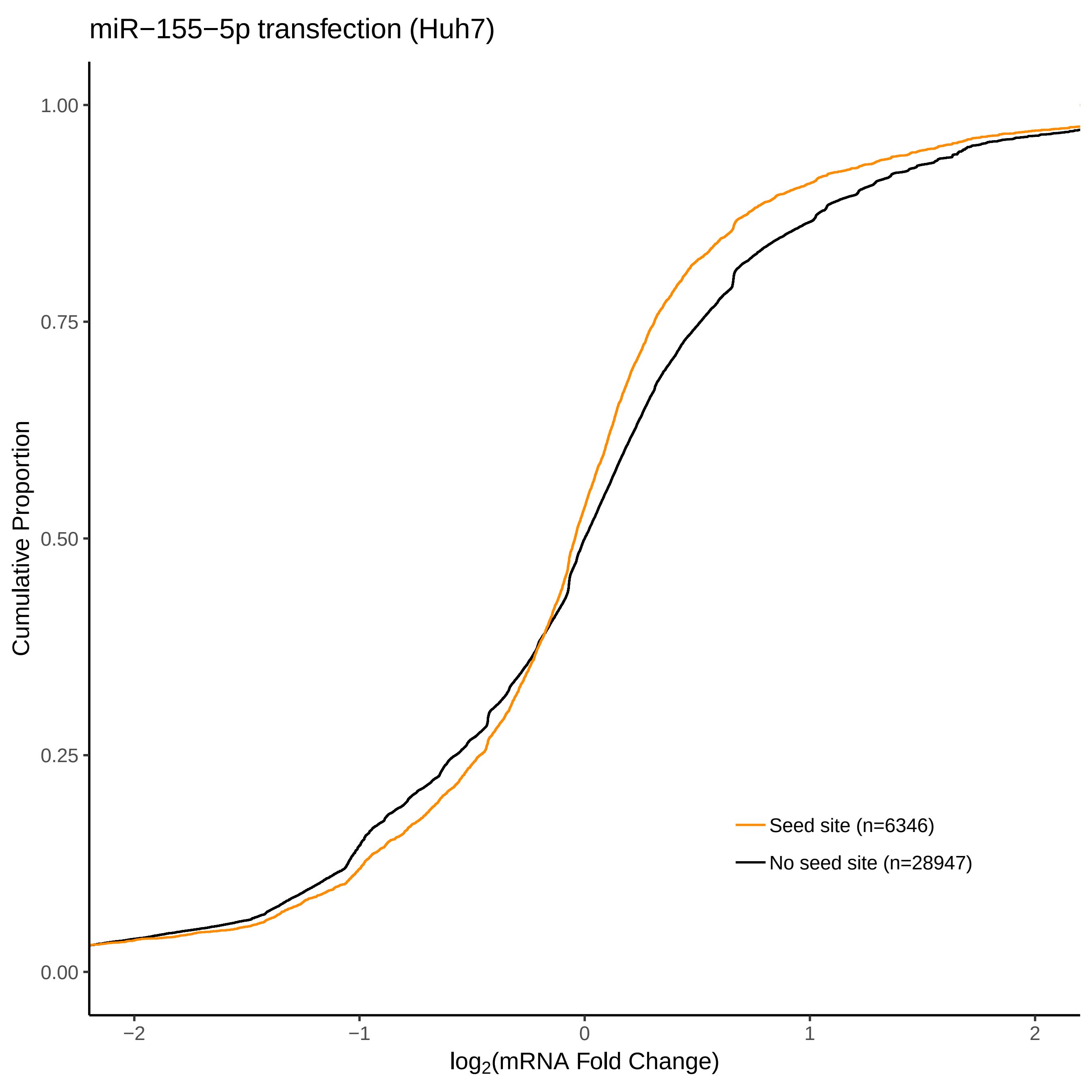

miR−155−5p transfection (Huh7)
Cumulative Proportion
1.00
0.75
0.50
0.25
0.00
−2
−1
0
1
2
log2(mRNA Fold Change)
Seed site (n=6346)
No seed site (n=28947)

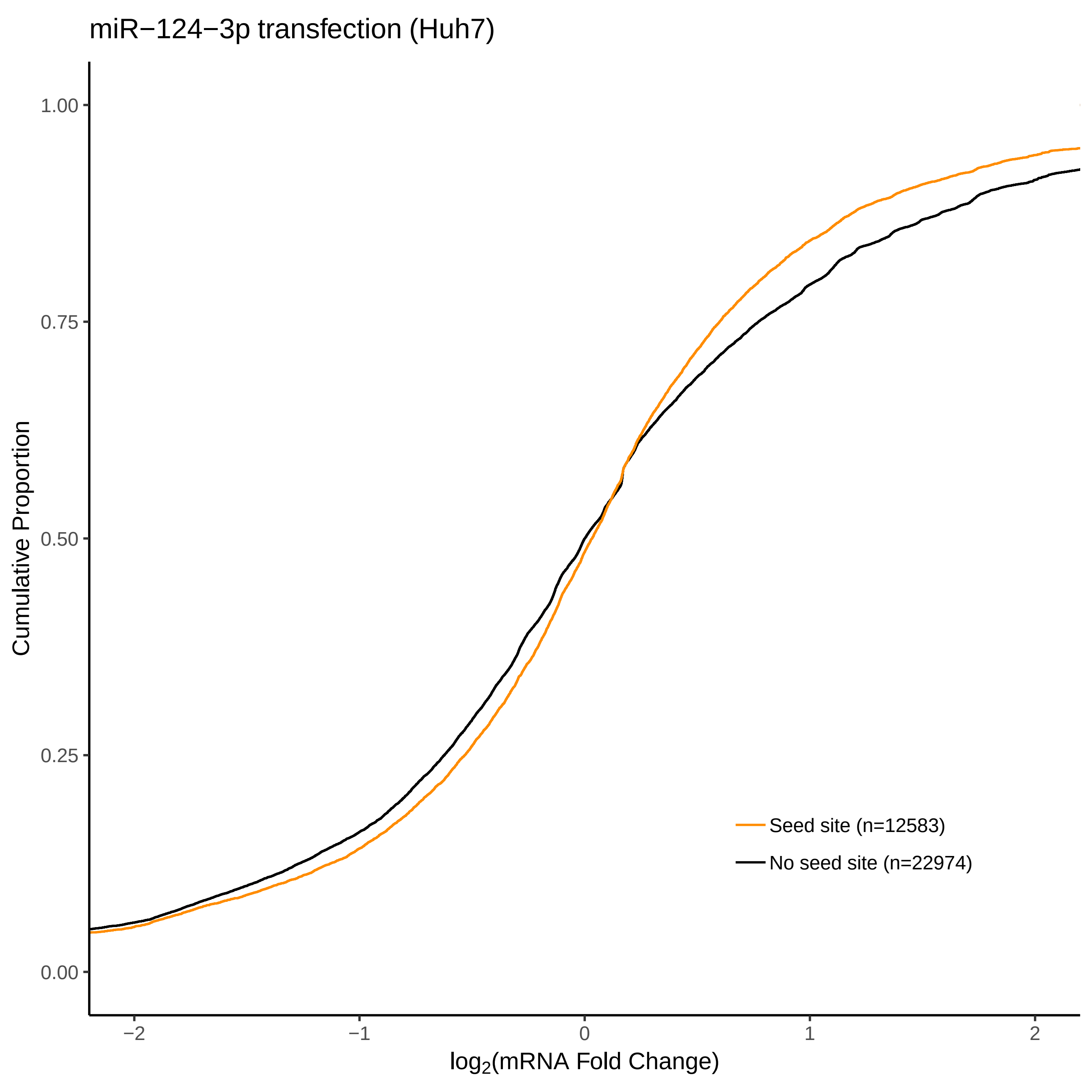

miR−124−3p transfection (Huh7)
Cumulative Proportion
1.00
0.75
0.50
0.25
0.00
−2
−1
0
1
2
log2(mRNA Fold Change)
Seed site (n=12583)
No seed site (n=22974)
