## Supplementary File 1b for "FilTar: Using RNA-Seq data to improve microRNA target prediction accuracy in animals"

### Known RNA-Seq microRNA transfection datasets not used in the analysis presented in this study

| Species | BioProject Accession | Source/Study | Sample | Run Accessions |
|---|---|---|---|---|
| *Homo sapiens* | PRJNA229375 | Nam *et al.* 2014 | HeLa | SRR1032873,SRR1032874, SRR1032875,SRR1032876, SRR1032877,SRR1032878, |
| | | | HEK293 | SRR1032879,SRR1032880, SRR1032881,SRR1032882 SRR1032883,SRR1032884 |
| | | | Huh7 | SRR1032885,SRR1032886, SRR1032887,SRR1032888 SRR1032890,SRR1032891, SRR1032892 |
| | | | IMR90 | SRR1032893,SRR1032894, SRR1032895,SRR1032896 |
| | PRJNA284262 | Zhang *et al.* 2016 | HeLa | SRR2031925,SRR2031926, SRR2031927,SRR2031928 |
| | PRJNA271411 | Iyer *et al.* 2015 | HeLa | SRR1737410,SRR1737413, SRR1737415,SRR1737416, SRR1737420,SRR1737421, SRR1737429,SRR1737430 |

**References**


Nam, J. W., Rissland, O. S., Koppstein, D., Abreu-Goodger, C., Jan, C. H., Agarwal, V., ... & Bartel, D. P. (2014). Global analyses of the effect of different cellular contexts on microRNA targeting. *Molecular cell*, *53*(6), 1031-1043.

Zhang, C., Lu, J., Liu, B., Cui, Q., & Wang, Y. (2016). Primate-specific miR-603 is implicated in the risk and pathogenesis of Alzheimer's disease. *Aging (Albany NY)*, *8*(2), 272.

Polioudakis, D., Abell, N. S., & Iyer, V. R. (2015). miR-503 represses human cell proliferation and directly targets the oncogene DDHD2 by non-canonical target pairing. *BMC genomics*, *16*(1), 40.
