## Supplementary File 2 for "FilTar: Using RNA-Seq data to improve microRNA target prediction accuracy in animals": PRJNA223608.html

MultiQC Report


### Toggle navigation v1.6 (2c9b9aa)

Loading report..

- General Stats
- Salmon
- Kallisto
- HISAT2
- Cutadapt
- FastQC
  - Sequence Counts
  - Sequence Quality Histograms
  - Per Sequence Quality Scores
  - Per Base Sequence Content
  - Per Sequence GC Content
  - Per Base N Content
  - Sequence Length Distribution
  - Sequence Duplication Levels
  - Overrepresented sequences
  - Adapter Content

Toolbox

##### MultiQC Toolbox

###### Apply Highlight Samples

+

Regex mode off
help
 Clear

###### Apply Rename Samples

+

Click here for bulk input.

Paste two columns of a tab-delimited table here (eg. from Excel).

First column should be the old name, second column the new name.

Add

Regex mode off
help
 Clear

###### Apply Show / Hide Samples

Hide matching samples

Show only matching samples

+

Regex mode off
help
 Clear

###### Export Plots

- Images
- Data

px

px

Aspect ratio

PNG
JPEG
SVG

Plot scaling

X

Download the raw data used to create the plots in this report below:

Format:

Tab-separated
Comma-separated
JSON

Note that additional data was saved in `reports/PRJNA223608_data` when this report was generated.

---

###### Choose Plots

 All
 None


---


   Download Plot Images

If you use plots from MultiQC in a publication or presentation, please cite:

> **MultiQC: Summarize analysis results for multiple tools and samples in a single report**  
> *Philip Ewels, Måns Magnusson, Sverker Lundin and Max Käller*  
> Bioinformatics (2016)  
> doi: 10.1093/bioinformatics/btw354  
> PMID: 27312411

###### Save Settings

You can save the toolbox settings for this report to the browser.

 Save


---

###### Load Settings

Choose a saved report profile from the dropdown box below:

[ select ]

Load
 Delete
 Set default
 Clear default

###### About MultiQC

This report was generated using MultiQC, version 1.6 (2c9b9aa)

You can see a YouTube video describing how to use MultiQC reports here:
https://youtu.be/qPbIlO\_KWN0

For more information about MultiQC, including other videos and
extensive documentation, please visit http://multiqc.info

You can report bugs, suggest improvements and find the source code for MultiQC on GitHub:
https://github.com/ewels/MultiQC

MultiQC is published in Bioinformatics:

> **MultiQC: Summarize analysis results for multiple tools and samples in a single report**  
> *Philip Ewels, Måns Magnusson, Sverker Lundin and Max Käller*  
> Bioinformatics (2016)  
> doi: 10.1093/bioinformatics/btw354  
> PMID: 27312411

A modular tool to aggregate results from bioinformatics analyses across many samples into a single report.

###### JavaScript Disabled

MultiQC reports use JavaScript for plots and toolbox functions. It looks like
you have JavaScript disabled in your web browser. Please note that many of the report
functions will not work as intended.

Loading report..

Report generated on 2019-01-30, 12:25 based on data in:

- `/gpfs/afm/moxon/thomas2/APAtrap/reports/SRR1598955_trimmed_fastqc.zip`
- `/gpfs/afm/moxon/thomas2/APAtrap/reports/SRR1598970_trimmed_fastqc.zip`
- `/gpfs/afm/moxon/thomas2/APAtrap/reports/SRR1598976_trimmed_fastqc.zip`
- `/gpfs/afm/moxon/thomas2/APAtrap/reports/SRR1598977_trimmed_fastqc.zip`
- `/gpfs/afm/moxon/thomas2/APAtrap/reports/SRR1598972_trimmed_fastqc.zip`
- `/gpfs/afm/moxon/thomas2/APAtrap/reports/SRR1598973_trimmed_fastqc.zip`
- `/gpfs/afm/moxon/thomas2/APAtrap/results/trimmed_fastq/SRR1598955.fastq.gz_trimming_report.txt`
- `/gpfs/afm/moxon/thomas2/APAtrap/results/trimmed_fastq/SRR1598970.fastq.gz_trimming_report.txt`
- `/gpfs/afm/moxon/thomas2/APAtrap/results/trimmed_fastq/SRR1598976.fastq.gz_trimming_report.txt`
- `/gpfs/afm/moxon/thomas2/APAtrap/results/trimmed_fastq/SRR1598977.fastq.gz_trimming_report.txt`
- `/gpfs/afm/moxon/thomas2/APAtrap/results/trimmed_fastq/SRR1598972.fastq.gz_trimming_report.txt`
- `/gpfs/afm/moxon/thomas2/APAtrap/results/trimmed_fastq/SRR1598973.fastq.gz_trimming_report.txt`
- `/gpfs/afm/moxon/thomas2/APAtrap/reports/hisat2/SRR1598955.txt`
- `/gpfs/afm/moxon/thomas2/APAtrap/reports/hisat2/SRR1598970.txt`
- `/gpfs/afm/moxon/thomas2/APAtrap/reports/hisat2/SRR1598976.txt`
- `/gpfs/afm/moxon/thomas2/APAtrap/reports/hisat2/SRR1598977.txt`
- `/gpfs/afm/moxon/thomas2/APAtrap/reports/hisat2/SRR1598972.txt`
- `/gpfs/afm/moxon/thomas2/APAtrap/reports/hisat2/SRR1598973.txt`
- `/gpfs/afm/moxon/thomas2/APAtrap/logs/SRR1598955_kallisto.out`
- `/gpfs/afm/moxon/thomas2/APAtrap/logs/SRR1598970_kallisto.out`
- `/gpfs/afm/moxon/thomas2/APAtrap/logs/SRR1598976_kallisto.out`
- `/gpfs/afm/moxon/thomas2/APAtrap/logs/SRR1598977_kallisto.out`
- `/gpfs/afm/moxon/thomas2/APAtrap/logs/SRR1598972_kallisto.out`
- `/gpfs/afm/moxon/thomas2/APAtrap/logs/SRR1598973_kallisto.out`
- `/gpfs/afm/moxon/thomas2/APAtrap/results/salmon/runs/hsa/SRR1598955`
- `/gpfs/afm/moxon/thomas2/APAtrap/results/salmon/runs/hsa/SRR1598970`
- `/gpfs/afm/moxon/thomas2/APAtrap/results/salmon/runs/hsa/SRR1598976`
- `/gpfs/afm/moxon/thomas2/APAtrap/results/salmon/runs/hsa/SRR1598977`
- `/gpfs/afm/moxon/thomas2/APAtrap/results/salmon/runs/hsa/SRR1598972`
- `/gpfs/afm/moxon/thomas2/APAtrap/results/salmon/runs/hsa/SRR1598973`

---

×
don't show again

**Welcome!** Not sure where to start?  
Watch a tutorial video
  *(6:06)*

#### General Statistics

 Copy table

 Configure Columns

 Sort by highlight

 Plot
Showing ^6^/_6_ rows and ^10^/_11_ columns.

| Sample Name | % Aligned | M Aligned | % Aligned | M Aligned | % Aligned | % Trimmed | % Dups | % GC | Length | % Failed | M Seqs |
| --- | --- | --- | --- | --- | --- | --- | --- | --- | --- | --- | --- |
| SRR1598955 | 55.1% | 10.7 | 54.3% | 10.6 | 89.2% | 10.0% | 44.4% | 48% | 36 bp | 8% | 19.5 |
| SRR1598970 | 38.3% | 10.1 | 37.9% | 10.0 | 87.9% | 17.9% | 77.8% | 49% | 36 bp | 25% | 26.4 |
| SRR1598972 | 54.4% | 11.3 | 53.7% | 11.1 | 89.7% | 10.9% | 47.9% | 48% | 36 bp | 17% | 20.7 |
| SRR1598973 | 35.1% | 26.3 | 35.3% | 26.4 | 38.9% | 23.2% | 79.2% | 50% | 49 bp | 25% | 74.8 |
| SRR1598976 | 52.9% | 10.5 | 52.2% | 10.4 | 91.0% | 9.9% | 59.2% | 48% | 36 bp | 17% | 19.9 |
| SRR1598977 | 34.5% | 9.0 | 34.6% | 9.0 | 87.6% | 16.9% | 79.7% | 49% | 36 bp | 17% | 26.1 |

×

###### General Statistics: Columns

Uncheck the tick box to hide columns. Click and drag the handle on the left to change order.

Show All
Show None

| Sort | Visible | Group | Column | Description | ID | Scale |
| --- | --- | --- | --- | --- | --- | --- |
| \|\| |  | Salmon | % Aligned | % Mapped reads | `percent_mapped` | None |
| \|\| |  | Salmon | M Aligned | Mapped reads (millions) | `num_mapped` | read\_count |
| \|\| |  | Kallisto | % Aligned | % processed reads that were pseudoaligned | `percent_aligned` | None |
| \|\| |  | Kallisto | M Aligned | Pseudoaligned reads (millions) | `pseudoaligned_reads` | read\_count |
| \|\| |  | HISAT2 | % Aligned | overall alignment rate | `overall_alignment_rate` | None |
| \|\| |  | Cutadapt | % Trimmed | % Total Base Pairs trimmed | `percent_trimmed` | None |
| \|\| |  | FastQC | % Dups | % Duplicate Reads | `percent_duplicates` | None |
| \|\| |  | FastQC | % GC | Average % GC Content | `percent_gc` | None |
| \|\| |  | FastQC | Length | Average Sequence Length (bp) | `avg_sequence_length` | None |
| \|\| |  | FastQC | % Failed | Percentage of modules failed in FastQC report (includes those not plotted here) | `percent_fails` | None |
| \|\| |  | FastQC | M Seqs | Total Sequences (millions) | `total_sequences` | read\_count |

Close

#### Salmon

Salmon is a tool for quantifying the expression of transcripts using RNA-seq data.

loading..

---

#### Kallisto

Kallisto is a program for quantifying abundances of transcripts from RNA-Seq data.

Number of Reads
Percentages

loading..

---

#### HISAT2

HISAT2 is a fast and sensitive alignment program for mapping NGS reads (both DNA and RNA) against a reference genome or population of reference genomes.

Number of Reads
Percentages

loading..

---

#### Cutadapt

Close
