## Supplementary File 2 for "FilTar: Using RNA-Seq data to improve microRNA target prediction accuracy in animals": PRJNA304643.html

MultiQC Report


### Toggle navigation v1.6 (2c9b9aa)

Loading report..

- General Stats
- Salmon
- Kallisto
- HISAT2
- Cutadapt
- FastQC
  - Sequence Counts
  - Sequence Quality Histograms
  - Per Sequence Quality Scores
  - Per Base Sequence Content
  - Per Sequence GC Content
  - Per Base N Content
  - Sequence Length Distribution
  - Sequence Duplication Levels
  - Overrepresented sequences
  - Adapter Content

Toolbox

##### MultiQC Toolbox

###### Apply Highlight Samples

+

Regex mode off
help
 Clear

###### Apply Rename Samples

+

Click here for bulk input.

Paste two columns of a tab-delimited table here (eg. from Excel).

First column should be the old name, second column the new name.

Add

Regex mode off
help
 Clear

###### Apply Show / Hide Samples

Hide matching samples

Show only matching samples

+

Regex mode off
help
 Clear

###### Export Plots

- Images
- Data

px

px

Aspect ratio

PNG
JPEG
SVG

Plot scaling

X

Download the raw data used to create the plots in this report below:

Format:

Tab-separated
Comma-separated
JSON

Note that additional data was saved in `reports/paired_end/PRJNA304643_data` when this report was generated.

---

###### Choose Plots

 All
 None


---


   Download Plot Images

If you use plots from MultiQC in a publication or presentation, please cite:

> **MultiQC: Summarize analysis results for multiple tools and samples in a single report**  
> *Philip Ewels, Måns Magnusson, Sverker Lundin and Max Käller*  
> Bioinformatics (2016)  
> doi: 10.1093/bioinformatics/btw354  
> PMID: 27312411

###### Save Settings

You can save the toolbox settings for this report to the browser.

 Save


---

###### Load Settings

Choose a saved report profile from the dropdown box below:

[ select ]

Load
 Delete
 Set default
 Clear default

###### About MultiQC

This report was generated using MultiQC, version 1.6 (2c9b9aa)

You can see a YouTube video describing how to use MultiQC reports here:
https://youtu.be/qPbIlO\_KWN0

For more information about MultiQC, including other videos and
extensive documentation, please visit http://multiqc.info

You can report bugs, suggest improvements and find the source code for MultiQC on GitHub:
https://github.com/ewels/MultiQC

MultiQC is published in Bioinformatics:

> **MultiQC: Summarize analysis results for multiple tools and samples in a single report**  
> *Philip Ewels, Måns Magnusson, Sverker Lundin and Max Käller*  
> Bioinformatics (2016)  
> doi: 10.1093/bioinformatics/btw354  
> PMID: 27312411

A modular tool to aggregate results from bioinformatics analyses across many samples into a single report.

###### JavaScript Disabled

MultiQC reports use JavaScript for plots and toolbox functions. It looks like
you have JavaScript disabled in your web browser. Please note that many of the report
functions will not work as intended.

Loading report..

Report generated on 2019-01-30, 15:09 based on data in:

- `/gpfs/afm/moxon/thomas2/APAtrap/results/trimmed_fastq/SRR2968576_1.fastq.gz_trimming_report.txt`
- `/gpfs/afm/moxon/thomas2/APAtrap/results/trimmed_fastq/SRR2968576_2.fastq.gz_trimming_report.txt`
- `/gpfs/afm/moxon/thomas2/APAtrap/results/trimmed_fastq/SRR2968577_1.fastq.gz_trimming_report.txt`
- `/gpfs/afm/moxon/thomas2/APAtrap/results/trimmed_fastq/SRR2968577_2.fastq.gz_trimming_report.txt`
- `/gpfs/afm/moxon/thomas2/APAtrap/results/trimmed_fastq/SRR2968578_1.fastq.gz_trimming_report.txt`
- `/gpfs/afm/moxon/thomas2/APAtrap/results/trimmed_fastq/SRR2968578_2.fastq.gz_trimming_report.txt`
- `/gpfs/afm/moxon/thomas2/APAtrap/results/trimmed_fastq/SRR2968579_1.fastq.gz_trimming_report.txt`
- `/gpfs/afm/moxon/thomas2/APAtrap/results/trimmed_fastq/SRR2968579_2.fastq.gz_trimming_report.txt`
- `/gpfs/afm/moxon/thomas2/APAtrap/results/trimmed_fastq/SRR2968580_1.fastq.gz_trimming_report.txt`
- `/gpfs/afm/moxon/thomas2/APAtrap/results/trimmed_fastq/SRR2968580_2.fastq.gz_trimming_report.txt`
- `/gpfs/afm/moxon/thomas2/APAtrap/results/trimmed_fastq/SRR2968581_1.fastq.gz_trimming_report.txt`
- `/gpfs/afm/moxon/thomas2/APAtrap/results/trimmed_fastq/SRR2968581_2.fastq.gz_trimming_report.txt`
- `/gpfs/afm/moxon/thomas2/APAtrap/results/trimmed_fastq/SRR2968582_1.fastq.gz_trimming_report.txt`
- `/gpfs/afm/moxon/thomas2/APAtrap/results/trimmed_fastq/SRR2968582_2.fastq.gz_trimming_report.txt`
- `/gpfs/afm/moxon/thomas2/APAtrap/results/trimmed_fastq/SRR2968583_1.fastq.gz_trimming_report.txt`
- `/gpfs/afm/moxon/thomas2/APAtrap/results/trimmed_fastq/SRR2968583_2.fastq.gz_trimming_report.txt`
- `/gpfs/afm/moxon/thomas2/APAtrap/results/trimmed_fastq/SRR2968584_1.fastq.gz_trimming_report.txt`
- `/gpfs/afm/moxon/thomas2/APAtrap/results/trimmed_fastq/SRR2968584_2.fastq.gz_trimming_report.txt`
- `/gpfs/afm/moxon/thomas2/APAtrap/results/trimmed_fastq/SRR2968585_1.fastq.gz_trimming_report.txt`
- `/gpfs/afm/moxon/thomas2/APAtrap/results/trimmed_fastq/SRR2968585_2.fastq.gz_trimming_report.txt`
- `/gpfs/afm/moxon/thomas2/APAtrap/results/trimmed_fastq/SRR2968586_1.fastq.gz_trimming_report.txt`
- `/gpfs/afm/moxon/thomas2/APAtrap/results/trimmed_fastq/SRR2968586_2.fastq.gz_trimming_report.txt`
- `/gpfs/afm/moxon/thomas2/APAtrap/results/trimmed_fastq/SRR2968587_1.fastq.gz_trimming_report.txt`
- `/gpfs/afm/moxon/thomas2/APAtrap/results/trimmed_fastq/SRR2968587_2.fastq.gz_trimming_report.txt`
- `/gpfs/afm/moxon/thomas2/APAtrap/results/trimmed_fastq/SRR2968588_1.fastq.gz_trimming_report.txt`
- `/gpfs/afm/moxon/thomas2/APAtrap/results/trimmed_fastq/SRR2968588_2.fastq.gz_trimming_report.txt`
- `/gpfs/afm/moxon/thomas2/APAtrap/results/trimmed_fastq/SRR2968589_1.fastq.gz_trimming_report.txt`
- `/gpfs/afm/moxon/thomas2/APAtrap/results/trimmed_fastq/SRR2968589_2.fastq.gz_trimming_report.txt`
- `/gpfs/afm/moxon/thomas2/APAtrap/results/trimmed_fastq/SRR2968590_1.fastq.gz_trimming_report.txt`
- `/gpfs/afm/moxon/thomas2/APAtrap/results/trimmed_fastq/SRR2968590_2.fastq.gz_trimming_report.txt`
- `/gpfs/afm/moxon/thomas2/APAtrap/results/trimmed_fastq/SRR2968591_1.fastq.gz_trimming_report.txt`
- `/gpfs/afm/moxon/thomas2/APAtrap/results/trimmed_fastq/SRR2968591_2.fastq.gz_trimming_report.txt`
- `/gpfs/afm/moxon/thomas2/APAtrap/results/trimmed_fastq/SRR2968592_1.fastq.gz_trimming_report.txt`
- `/gpfs/afm/moxon/thomas2/APAtrap/results/trimmed_fastq/SRR2968592_2.fastq.gz_trimming_report.txt`
- `/gpfs/afm/moxon/thomas2/APAtrap/results/trimmed_fastq/SRR2968593_1.fastq.gz_trimming_report.txt`
- `/gpfs/afm/moxon/thomas2/APAtrap/results/trimmed_fastq/SRR2968593_2.fastq.gz_trimming_report.txt`
- `/gpfs/afm/moxon/thomas2/APAtrap/results/trimmed_fastq/SRR2968594_1.fastq.gz_trimming_report.txt`
- `/gpfs/afm/moxon/thomas2/APAtrap/results/trimmed_fastq/SRR2968594_2.fastq.gz_trimming_report.txt`
- `/gpfs/afm/moxon/thomas2/APAtrap/results/trimmed_fastq/SRR2968595_1.fastq.gz_trimming_report.txt`
- `/gpfs/afm/moxon/thomas2/APAtrap/results/trimmed_fastq/SRR2968595_2.fastq.gz_trimming_report.txt`
- `/gpfs/afm/moxon/thomas2/APAtrap/results/trimmed_fastq/SRR2968596_1.fastq.gz_trimming_report.txt`
- `/gpfs/afm/moxon/thomas2/APAtrap/results/trimmed_fastq/SRR2968596_2.fastq.gz_trimming_report.txt`
- `/gpfs/afm/moxon/thomas2/APAtrap/results/trimmed_fastq/SRR2968597_1.fastq.gz_trimming_report.txt`
- `/gpfs/afm/moxon/thomas2/APAtrap/results/trimmed_fastq/SRR2968597_2.fastq.gz_trimming_report.txt`
- `/gpfs/afm/moxon/thomas2/APAtrap/results/trimmed_fastq/SRR2968598_1.fastq.gz_trimming_report.txt`
- `/gpfs/afm/moxon/thomas2/APAtrap/results/trimmed_fastq/SRR2968598_2.fastq.gz_trimming_report.txt`
- `/gpfs/afm/moxon/thomas2/APAtrap/results/trimmed_fastq/SRR2968599_1.fastq.gz_trimming_report.txt`
- `/gpfs/afm/moxon/thomas2/APAtrap/results/trimmed_fastq/SRR2968599_2.fastq.gz_trimming_report.txt`
- `/gpfs/afm/moxon/thomas2/APAtrap/reports/SRR2968576_1_val_1_fastqc.zip`
- `/gpfs/afm/moxon/thomas2/APAtrap/reports/SRR2968576_2_val_2_fastqc.zip`
- `/gpfs/afm/moxon/thomas2/APAtrap/reports/SRR2968577_1_val_1_fastqc.zip`
- `/gpfs/afm/moxon/thomas2/APAtrap/reports/SRR2968577_2_val_2_fastqc.zip`
- `/gpfs/afm/moxon/thomas2/APAtrap/reports/SRR2968578_1_val_1_fastqc.zip`
- `/gpfs/afm/moxon/thomas2/APAtrap/reports/SRR2968578_2_val_2_fastqc.zip`
- `/gpfs/afm/moxon/thomas2/APAtrap/reports/SRR2968579_1_val_1_fastqc.zip`
- `/gpfs/afm/moxon/thomas2/APAtrap/reports/SRR2968579_2_val_2_fastqc.zip`
- `/gpfs/afm/moxon/thomas2/APAtrap/reports/SRR2968580_1_val_1_fastqc.zip`
- `/gpfs/afm/moxon/thomas2/APAtrap/reports/SRR2968580_2_val_2_fastqc.zip`
- `/gpfs/afm/moxon/thomas2/APAtrap/reports/SRR2968581_1_val_1_fastqc.zip`
- `/gpfs/afm/moxon/thomas2/APAtrap/reports/SRR2968581_2_val_2_fastqc.zip`
- `/gpfs/afm/moxon/thomas2/APAtrap/reports/SRR2968582_1_val_1_fastqc.zip`
- `/gpfs/afm/moxon/thomas2/APAtrap/reports/SRR2968582_2_val_2_fastqc.zip`
- `/gpfs/afm/moxon/thomas2/APAtrap/reports/SRR2968583_1_val_1_fastqc.zip`
- `/gpfs/afm/moxon/thomas2/APAtrap/reports/SRR2968583_2_val_2_fastqc.zip`
- `/gpfs/afm/moxon/thomas2/APAtrap/reports/SRR2968584_1_val_1_fastqc.zip`
- `/gpfs/afm/moxon/thomas2/APAtrap/reports/SRR2968584_2_val_2_fastqc.zip`
- `/gpfs/afm/moxon/thomas2/APAtrap/reports/SRR2968585_1_val_1_fastqc.zip`
- `/gpfs/afm/moxon/thomas2/APAtrap/reports/SRR2968585_2_val_2_fastqc.zip`
- `/gpfs/afm/moxon/thomas2/APAtrap/reports/SRR2968586_1_val_1_fastqc.zip`
- `/gpfs/afm/moxon/thomas2/APAtrap/reports/SRR2968586_2_val_2_fastqc.zip`
- `/gpfs/afm/moxon/thomas2/APAtrap/reports/SRR2968587_1_val_1_fastqc.zip`
- `/gpfs/afm/moxon/thomas2/APAtrap/reports/SRR2968587_2_val_2_fastqc.zip`
- `/gpfs/afm/moxon/thomas2/APAtrap/reports/SRR2968588_1_val_1_fastqc.zip`
- `/gpfs/afm/moxon/thomas2/APAtrap/reports/SRR2968588_2_val_2_fastqc.zip`
- `/gpfs/afm/moxon/thomas2/APAtrap/reports/SRR2968589_1_val_1_fastqc.zip`
- `/gpfs/afm/moxon/thomas2/APAtrap/reports/SRR2968589_2_val_2_fastqc.zip`
- `/gpfs/afm/moxon/thomas2/APAtrap/reports/SRR2968590_1_val_1_fastqc.zip`
- `/gpfs/afm/moxon/thomas2/APAtrap/reports/SRR2968590_2_val_2_fastqc.zip`
- `/gpfs/afm/moxon/thomas2/APAtrap/reports/SRR2968591_1_val_1_fastqc.zip`
- `/gpfs/afm/moxon/thomas2/APAtrap/reports/SRR2968591_2_val_2_fastqc.zip`
- `/gpfs/afm/moxon/thomas2/APAtrap/reports/SRR2968592_1_val_1_fastqc.zip`
- `/gpfs/afm/moxon/thomas2/APAtrap/reports/SRR2968592_2_val_2_fastqc.zip`
- `/gpfs/afm/moxon/thomas2/APAtrap/reports/SRR2968593_1_val_1_fastqc.zip`
- `/gpfs/afm/moxon/thomas2/APAtrap/reports/SRR2968593_2_val_2_fastqc.zip`
- `/gpfs/afm/moxon/thomas2/APAtrap/reports/SRR2968594_1_val_1_fastqc.zip`
- `/gpfs/afm/moxon/thomas2/APAtrap/reports/SRR2968594_2_val_2_fastqc.zip`
- `/gpfs/afm/moxon/thomas2/APAtrap/reports/SRR2968595_1_val_1_fastqc.zip`
- `/gpfs/afm/moxon/thomas2/APAtrap/reports/SRR2968595_2_val_2_fastqc.zip`
- `/gpfs/afm/moxon/thomas2/APAtrap/reports/SRR2968596_1_val_1_fastqc.zip`
- `/gpfs/afm/moxon/thomas2/APAtrap/reports/SRR2968596_2_val_2_fastqc.zip`
- `/gpfs/afm/moxon/thomas2/APAtrap/reports/SRR2968597_1_val_1_fastqc.zip`
- `/gpfs/afm/moxon/thomas2/APAtrap/reports/SRR2968597_2_val_2_fastqc.zip`
- `/gpfs/afm/moxon/thomas2/APAtrap/reports/SRR2968598_1_val_1_fastqc.zip`
- `/gpfs/afm/moxon/thomas2/APAtrap/reports/SRR2968598_2_val_2_fastqc.zip`
- `/gpfs/afm/moxon/thomas2/APAtrap/reports/SRR2968599_1_val_1_fastqc.zip`
- `/gpfs/afm/moxon/thomas2/APAtrap/reports/SRR2968599_2_val_2_fastqc.zip`
- `/gpfs/afm/moxon/thomas2/APAtrap/reports/hisat2/SRR2968576.txt`
- `/gpfs/afm/moxon/thomas2/APAtrap/reports/hisat2/SRR2968577.txt`
- `/gpfs/afm/moxon/thomas2/APAtrap/reports/hisat2/SRR2968578.txt`
- `/gpfs/afm/moxon/thomas2/APAtrap/reports/hisat2/SRR2968579.txt`
- `/gpfs/afm/moxon/thomas2/APAtrap/reports/hisat2/SRR2968580.txt`
- `/gpfs/afm/moxon/thomas2/APAtrap/reports/hisat2/SRR2968581.txt`
- `/gpfs/afm/moxon/thomas2/APAtrap/reports/hisat2/SRR2968582.txt`
- `/gpfs/afm/moxon/thomas2/APAtrap/reports/hisat2/SRR2968583.txt`
- `/gpfs/afm/moxon/thomas2/APAtrap/reports/hisat2/SRR2968584.txt`
- `/gpfs/afm/moxon/thomas2/APAtrap/reports/hisat2/SRR2968585.txt`
- `/gpfs/afm/moxon/thomas2/APAtrap/reports/hisat2/SRR2968586.txt`
- `/gpfs/afm/moxon/thomas2/APAtrap/reports/hisat2/SRR2968587.txt`
- `/gpfs/afm/moxon/thomas2/APAtrap/reports/hisat2/SRR2968588.txt`
- `/gpfs/afm/moxon/thomas2/APAtrap/reports/hisat2/SRR2968589.txt`
- `/gpfs/afm/moxon/thomas2/APAtrap/reports/hisat2/SRR2968590.txt`
- `/gpfs/afm/moxon/thomas2/APAtrap/reports/hisat2/SRR2968591.txt`
- `/gpfs/afm/moxon/thomas2/APAtrap/reports/hisat2/SRR2968592.txt`
- `/gpfs/afm/moxon/thomas2/APAtrap/reports/hisat2/SRR2968593.txt`
- `/gpfs/afm/moxon/thomas2/APAtrap/reports/hisat2/SRR2968594.txt`
- `/gpfs/afm/moxon/thomas2/APAtrap/reports/hisat2/SRR2968595.txt`
- `/gpfs/afm/moxon/thomas2/APAtrap/reports/hisat2/SRR2968596.txt`
- `/gpfs/afm/moxon/thomas2/APAtrap/reports/hisat2/SRR2968597.txt`
- `/gpfs/afm/moxon/thomas2/APAtrap/reports/hisat2/SRR2968598.txt`
- `/gpfs/afm/moxon/thomas2/APAtrap/reports/hisat2/SRR2968599.txt`
- `/gpfs/afm/moxon/thomas2/APAtrap/logs/SRR2968576_kallisto.out`
- `/gpfs/afm/moxon/thomas2/APAtrap/logs/SRR2968577_kallisto.out`
- `/gpfs/afm/moxon/thomas2/APAtrap/logs/SRR2968578_kallisto.out`
- `/gpfs/afm/moxon/thomas2/APAtrap/logs/SRR2968579_kallisto.out`
- `/gpfs/afm/moxon/thomas2/APAtrap/logs/SRR2968580_kallisto.out`
- `/gpfs/afm/moxon/thomas2/APAtrap/logs/SRR2968581_kallisto.out`
- `/gpfs/afm/moxon/thomas2/APAtrap/logs/SRR2968582_kallisto.out`
- `/gpfs/afm/moxon/thomas2/APAtrap/logs/SRR2968583_kallisto.out`
- `/gpfs/afm/moxon/thomas2/APAtrap/logs/SRR2968584_kallisto.out`
- `/gpfs/afm/moxon/thomas2/APAtrap/logs/SRR2968585_kallisto.out`
- `/gpfs/afm/moxon/thomas2/APAtrap/logs/SRR2968586_kallisto.out`
- `/gpfs/afm/moxon/thomas2/APAtrap/logs/SRR2968587_kallisto.out`
- `/gpfs/afm/moxon/thomas2/APAtrap/logs/SRR2968588_kallisto.out`
- `/gpfs/afm/moxon/thomas2/APAtrap/logs/SRR2968589_kallisto.out`
- `/gpfs/afm/moxon/thomas2/APAtrap/logs/SRR2968590_kallisto.out`
- `/gpfs/afm/moxon/thomas2/APAtrap/logs/SRR2968591_kallisto.out`
- `/gpfs/afm/moxon/thomas2/APAtrap/logs/SRR2968592_kallisto.out`
- `/gpfs/afm/moxon/thomas2/APAtrap/logs/SRR2968593_kallisto.out`
- `/gpfs/afm/moxon/thomas2/APAtrap/logs/SRR2968594_kallisto.out`
- `/gpfs/afm/moxon/thomas2/APAtrap/logs/SRR2968595_kallisto.out`
- `/gpfs/afm/moxon/thomas2/APAtrap/logs/SRR2968596_kallisto.out`
- `/gpfs/afm/moxon/thomas2/APAtrap/logs/SRR2968597_kallisto.out`
- `/gpfs/afm/moxon/thomas2/APAtrap/logs/SRR2968598_kallisto.out`
- `/gpfs/afm/moxon/thomas2/APAtrap/logs/SRR2968599_kallisto.out`
- `/gpfs/afm/moxon/thomas2/APAtrap/results/salmon/runs/hsa/SRR2968576`
- `/gpfs/afm/moxon/thomas2/APAtrap/results/salmon/runs/hsa/SRR2968577`
- `/gpfs/afm/moxon/thomas2/APAtrap/results/salmon/runs/hsa/SRR2968578`
- `/gpfs/afm/moxon/thomas2/APAtrap/results/salmon/runs/hsa/SRR2968579`
- `/gpfs/afm/moxon/thomas2/APAtrap/results/salmon/runs/hsa/SRR2968580`
- `/gpfs/afm/moxon/thomas2/APAtrap/results/salmon/runs/hsa/SRR2968581`
- `/gpfs/afm/moxon/thomas2/APAtrap/results/salmon/runs/hsa/SRR2968582`
- `/gpfs/afm/moxon/thomas2/APAtrap/results/salmon/runs/hsa/SRR2968583`
- `/gpfs/afm/moxon/thomas2/APAtrap/results/salmon/runs/hsa/SRR2968584`
- `/gpfs/afm/moxon/thomas2/APAtrap/results/salmon/runs/hsa/SRR2968585`
- `/gpfs/afm/moxon/thomas2/APAtrap/results/salmon/runs/hsa/SRR2968586`
- `/gpfs/afm/moxon/thomas2/APAtrap/results/salmon/runs/hsa/SRR2968587`
- `/gpfs/afm/moxon/thomas2/APAtrap/results/salmon/runs/hsa/SRR2968588`
- `/gpfs/afm/moxon/thomas2/APAtrap/results/salmon/runs/hsa/SRR2968589`
- `/gpfs/afm/moxon/thomas2/APAtrap/results/salmon/runs/hsa/SRR2968590`
- `/gpfs/afm/moxon/thomas2/APAtrap/results/salmon/runs/hsa/SRR2968591`
- `/gpfs/afm/moxon/thomas2/APAtrap/results/salmon/runs/hsa/SRR2968592`
- `/gpfs/afm/moxon/thomas2/APAtrap/results/salmon/runs/hsa/SRR2968593`
- `/gpfs/afm/moxon/thomas2/APAtrap/results/salmon/runs/hsa/SRR2968594`
- `/gpfs/afm/moxon/thomas2/APAtrap/results/salmon/runs/hsa/SRR2968595`
- `/gpfs/afm/moxon/thomas2/APAtrap/results/salmon/runs/hsa/SRR2968596`
- `/gpfs/afm/moxon/thomas2/APAtrap/results/salmon/runs/hsa/SRR2968597`
- `/gpfs/afm/moxon/thomas2/APAtrap/results/salmon/runs/hsa/SRR2968598`
- `/gpfs/afm/moxon/thomas2/APAtrap/results/salmon/runs/hsa/SRR2968599`

---

×
don't show again

**Welcome!** Not sure where to start?  
Watch a tutorial video
  *(6:06)*

#### General Statistics

 Copy table

 Configure Columns

 Sort by highlight

 Plot
Showing ^120^/_120_ rows and ^10^/_12_ columns.

| Sample Name | % Aligned | M Aligned | Frag Length | % Aligned | M Aligned | % Aligned | % Trimmed | % Dups | % GC | Length | % Failed | M Seqs |
| --- | --- | --- | --- | --- | --- | --- | --- | --- | --- | --- | --- | --- |
| SRR2968576 | 87.3% | 20.9 |  |  |  | 98.3% |  |  |  |  |  |  |
| SRR2968576\_1 |  |  |  |  |  |  | 0.8% |  |  |  |  |  |
| SRR2968576\_1\_val\_1 |  |  | 179.5bp | 87.6% | 21.0 |  |  | 46.6% | 49% | 48 bp | 18% | 24.0 |
| SRR2968576\_2 |  |  |  |  |  |  | 2.3% |  |  |  |  |  |
| SRR2968576\_2\_val\_2 |  |  |  |  |  |  |  | 46.2% | 49% | 48 bp | 18% | 24.0 |
| SRR2968577 | 87.8% | 17.5 |  |  |  | 98.3% |  |  |  |  |  |  |
| SRR2968577\_1 |  |  |  |  |  |  | 1.0% |  |  |  |  |  |
| SRR2968577\_1\_val\_1 |  |  | 173.6bp | 88.1% | 17.6 |  |  | 44.8% | 49% | 48 bp | 18% | 20.0 |
| SRR2968577\_2 |  |  |  |  |  |  | 2.7% |  |  |  |  |  |
| SRR2968577\_2\_val\_2 |  |  |  |  |  |  |  | 44.6% | 49% | 48 bp | 18% | 20.0 |
| SRR2968578 | 88.0% | 15.0 |  |  |  | 98.3% |  |  |  |  |  |  |
| SRR2968578\_1 |  |  |  |  |  |  | 0.9% |  |  |  |  |  |
| SRR2968578\_1\_val\_1 |  |  | 171.1bp | 88.2% | 15.0 |  |  | 42.7% | 49% | 48 bp | 18% | 17.1 |
| SRR2968578\_2 |  |  |  |  |  |  | 2.5% |  |  |  |  |  |
| SRR2968578\_2\_val\_2 |  |  |  |  |  |  |  | 42.6% | 49% | 48 bp | 18% | 17.1 |
| SRR2968579 | 77.6% | 17.2 |  |  |  | 98.3% |  |  |  |  |  |  |
| SRR2968579\_1 |  |  |  |  |  |  | 0.8% |  |  |  |  |  |
| SRR2968579\_1\_val\_1 |  |  | 182.4bp | 77.9% | 17.3 |  |  | 50.4% | 50% | 48 bp | 27% | 22.2 |
| SRR2968579\_2 |  |  |  |  |  |  | 2.5% |  |  |  |  |  |
| SRR2968579\_2\_val\_2 |  |  |  |  |  |  |  | 50.0% | 50% | 48 bp | 18% | 22.2 |
| SRR2968580 | 85.8% | 26.6 |  |  |  | 98.4% |  |  |  |  |  |  |
| SRR2968580\_1 |  |  |  |  |  |  | 0.6% |  |  |  |  |  |
| SRR2968580\_1\_val\_1 |  |  | 180.7bp | 86.0% | 26.6 |  |  | 50.7% | 49% | 48 bp | 27% | 31.0 |
| SRR2968580\_2 |  |  |  |  |  |  | 2.0% |  |  |  |  |  |
| SRR2968580\_2\_val\_2 |  |  |  |  |  |  |  | 50.6% | 49% | 48 bp | 27% | 31.0 |
| SRR2968581 | 88.1% | 28.8 |  |  |  | 98.3% |  |  |  |  |  |  |
| SRR2968581\_1 |  |  |  |  |  |  | 0.7% |  |  |  |  |  |
| SRR2968581\_1\_val\_1 |  |  | 178.9bp | 88.3% | 28.9 |  |  | 49.1% | 49% | 48 bp | 18% | 32.7 |
| SRR2968581\_2 |  |  |  |  |  |  | 2.1% |  |  |  |  |  |
| SRR2968581\_2\_val\_2 |  |  |  |  |  |  |  | 48.9% | 49% | 48 bp | 18% | 32.7 |
| SRR2968582 | 88.5% | 18.9 |  |  |  | 98.3% |  |  |  |  |  |  |
| SRR2968582\_1 |  |  |  |  |  |  | 0.7% |  |  |  |  |  |
| SRR2968582\_1\_val\_1 |  |  | 184.7bp | 88.7% | 19.0 |  |  | 44.3% | 49% | 48 bp | 18% | 21.4 |
| SRR2968582\_2 |  |  |  |  |  |  | 2.1% |  |  |  |  |  |
| SRR2968582\_2\_val\_2 |  |  |  |  |  |  |  | 43.8% | 49% | 48 bp | 18% | 21.4 |
| SRR2968583 | 87.6% | 15.0 |  |  |  | 98.2% |  |  |  |  |  |  |
| SRR2968583\_1 |  |  |  |  |  |  | 1.6% |  |  |  |  |  |
| SRR2968583\_1\_val\_1 |  |  | 184.9bp | 87.9% | 15.0 |  |  | 40.5% | 48% | 48 bp | 18% | 17.1 |
| SRR2968583\_2 |  |  |  |  |  |  | 3.0% |  |  |  |  |  |
| SRR2968583\_2\_val\_2 |  |  |  |  |  |  |  | 40.3% | 48% | 48 bp | 9% | 17.1 |
| SRR2968584 | 85.4% | 10.9 |  |  |  | 90.0% |  |  |  |  |  |  |
| SRR2968584\_1 |  |  |  |  |  |  | 1.3% |  |  |  |  |  |
| SRR2968584\_1\_val\_1 |  |  | 225.0bp | 85.7% | 10.9 |  |  | 33.1% | 49% | 48 bp | 18% | 12.7 |
| SRR2968584\_2 |  |  |  |  |  |  | 15.8% |  |  |  |  |  |
| SRR2968584\_2\_val\_2 |  |  |  |  |  |  |  | 25.4% | 49% | 47 bp | 18% | 12.7 |
| SRR2968585 | 85.5% | 11.3 |  |  |  | 89.8% |  |  |  |  |  |  |
| SRR2968585\_1 |  |  |  |  |  |  | 1.4% |  |  |  |  |  |
| SRR2968585\_1\_val\_1 |  |  | 224.3bp | 85.8% | 11.4 |  |  | 33.7% | 49% | 48 bp | 18% | 13.3 |
| SRR2968585\_2 |  |  |  |  |  |  | 13.0% |  |  |  |  |  |
| SRR2968585\_2\_val\_2 |  |  |  |  |  |  |  | 21.8% | 49% | 47 bp | 18% | 13.3 |
| SRR2968586 | 85.8% | 10.2 |  |  |  | 90.1% |  |  |  |  |  |  |
| SRR2968586\_1 |  |  |  |  |  |  | 1.5% |  |  |  |  |  |
| SRR2968586\_1\_val\_1 |  |  | 213.4bp | 86.1% | 10.2 |  |  | 32.7% | 49% | 48 bp | 18% | 11.9 |
| SRR2968586\_2 |  |  |  |  |  |  | 16.4% |  |  |  |  |  |
| SRR2968586\_2\_val\_2 |  |  |  |  |  |  |  | 25.6% | 49% | 47 bp | 18% | 11.9 |
| SRR2968587 | 85.9% | 10.6 |  |  |  | 90.0% |  |  |  |  |  |  |
| SRR2968587\_1 |  |  |  |  |  |  | 1.6% |  |  |  |  |  |
| SRR2968587\_1\_val\_1 |  |  | 212.6bp | 86.2% | 10.7 |  |  | 33.3% | 49% | 48 bp | 18% | 12.4 |
| SRR2968587\_2 |  |  |  |  |  |  | 12.8% |  |  |  |  |  |
| SRR2968587\_2\_val\_2 |  |  |  |  |  |  |  | 22.3% | 49% | 47 bp | 18% | 12.4 |
| SRR2968588 | 86.0% | 8.5 |  |  |  | 90.3% |  |  |  |  |  |  |
| SRR2968588\_1 |  |  |  |  |  |  | 1.3% |  |  |  |  |  |
| SRR2968588\_1\_val\_1 |  |  | 206.5bp | 86.3% | 8.5 |  |  | 30.6% | 49% | 48 bp | 18% | 9.9 |
| SRR2968588\_2 |  |  |  |  |  |  | 14.2% |  |  |  |  |  |
| SRR2968588\_2\_val\_2 |  |  |  |  |  |  |  | 23.7% | 49% | 47 bp | 18% | 9.9 |
| SRR2968589 | 86.1% | 8.9 |  |  |  | 90.2% |  |  |  |  |  |  |
| SRR2968589\_1 |  |  |  |  |  |  | 1.4% |  |  |  |  |  |
| SRR2968589\_1\_val\_1 |  |  | 206.6bp | 86.4% | 8.9 |  |  | 31.3% | 49% | 48 bp | 18% | 10.3 |
| SRR2968589\_2 |  |  |  |  |  |  | 12.0% |  |  |  |  |  |
| SRR2968589\_2\_val\_2 |  |  |  |  |  |  |  | 20.7% | 49% | 47 bp | 18% | 10.3 |
| SRR2968590 | 84.9% | 11.6 |  |  |  | 90.0% |  |  |  |  |  |  |
| SRR2968590\_1 |  |  |  |  |  |  | 1.4% |  |  |  |  |  |
| SRR2968590\_1\_val\_1 |  |  | 195.7bp | 85.2% | 11.6 |  |  | 34.8% | 49% | 48 bp | 18% | 13.7 |
| SRR2968590\_2 |  |  |  |  |  |  | 15.8% |  |  |  |  |  |
| SRR2968590\_2\_val\_2 |  |  |  |  |  |  |  | 26.9% | 49% | 47 bp | 18% | 13.7 |
| SRR2968591 | 85.0% | 12.1 |  |  |  | 89.8% |  |  |  |  |  |  |
| SRR2968591\_1 |  |  |  |  |  |  | 1.5% |  |  |  |  |  |
| SRR2968591\_1\_val\_1 |  |  | 195.7bp | 85.2% | 12.1 |  |  | 35.5% | 49% | 48 bp | 18% | 14.2 |
| SRR2968591\_2 |  |  |  |  |  |  | 12.9% |  |  |  |  |  |
| SRR2968591\_2\_val\_2 |  |  |  |  |  |  |  | 23.7% | 49% | 47 bp | 18% | 14.2 |
| SRR2968592 | 82.6% | 9.7 |  |  |  | 90.1% |  |  |  |  |  |  |
| SRR2968592\_1 |  |  |  |  |  |  | 1.5% |  |  |  |  |  |
| SRR2968592\_1\_val\_1 |  |  | 202.5bp | 82.8% | 9.7 |  |  | 32.9% | 50% | 48 bp | 18% | 11.7 |
| SRR2968592\_2 |  |  |  |  |  |  | 16.0% |  |  |  |  |  |
| SRR2968592\_2\_val\_2 |  |  |  |  |  |  |  | 25.6% | 50% | 47 bp | 18% | 11.7 |
| SRR2968593 | 82.6% | 10.1 |  |  |  | 89.9% |  |  |  |  |  |  |
| SRR2968593\_1 |  |  |  |  |  |  | 1.7% |  |  |  |  |  |
| SRR2968593\_1\_val\_1 |  |  | 202.9bp | 82.9% | 10.1 |  |  | 33.5% | 49% | 48 bp | 9% | 12.2 |
| SRR2968593\_2 |  |  |  |  |  |  | 12.9% |  |  |  |  |  |
| SRR2968593\_2\_val\_2 |  |  |  |  |  |  |  | 22.6% | 50% | 47 bp | 18% | 12.2 |
| SRR2968594 | 85.0% | 13.0 |  |  |  | 90.0% |  |  |  |  |  |  |
| SRR2968594\_1 |  |  |  |  |  |  | 1.5% |  |  |  |  |  |
| SRR2968594\_1\_val\_1 |  |  | 200.9bp | 85.3% | 13.0 |  |  | 34.0% | 49% | 48 bp | 18% | 15.2 |
| SRR2968594\_2 |  |  |  |  |  |  | 15.6% |  |  |  |  |  |
| SRR2968594\_2\_val\_2 |  |  |  |  |  |  |  | 27.1% | 49% | 47 bp | 18% | 15.2 |
| SRR2968595 | 85.1% | 13.5 |  |  |  | 89.8% |  |  |  |  |  |  |
| SRR2968595\_1 |  |  |  |  |  |  | 1.7% |  |  |  |  |  |
| SRR2968595\_1\_val\_1 |  |  | 200.9bp | 85.4% | 13.6 |  |  | 34.9% | 49% | 48 bp | 18% | 15.9 |
| SRR2968595\_2 |  |  |  |  |  |  | 12.7% |  |  |  |  |  |
| SRR2968595\_2\_val\_2 |  |  |  |  |  |  |  | 23.8% | 49% | 47 bp | 18% | 15.9 |
| SRR2968596 | 85.1% | 12.0 |  |  |  | 89.8% |  |  |  |  |  |  |
| SRR2968596\_1 |  |  |  |  |  |  | 1.5% |  |  |  |  |  |
| SRR2968596\_1\_val\_1 |  |  | 209.6bp | 85.3% | 12.0 |  |  | 32.8% | 49% | 48 bp | 18% | 14.1 |
| SRR2968596\_2 |  |  |  |  |  |  | 16.5% |  |  |  |  |  |
| SRR2968596\_2\_val\_2 |  |  |  |  |  |  |  | 25.5% | 49% | 47 bp | 18% | 14.1 |
| SRR2968597 | 85.1% | 12.5 |  |  |  | 89.7% |  |  |  |  |  |  |
| SRR2968597\_1 |  |  |  |  |  |  | 1.7% |  |  |  |  |  |
| SRR2968597\_1\_val\_1 |  |  | 209.5bp | 85.4% | 12.6 |  |  | 33.6% | 49% | 48 bp | 18% | 14.7 |
| SRR2968597\_2 |  |  |  |  |  |  | 13.7% |  |  |  |  |  |
| SRR2968597\_2\_val\_2 |  |  |  |  |  |  |  | 22.3% | 49% | 47 bp | 18% | 14.7 |
| SRR2968598 | 83.9% | 11.5 |  |  |  | 89.9% |  |  |  |  |  |  |
| SRR2968598\_1 |  |  |  |  |  |  | 1.5% |  |  |  |  |  |
| SRR2968598\_1\_val\_1 |  |  | 191.9bp | 84.2% | 11.6 |  |  | 33.8% | 49% | 48 bp | 18% | 13.7 |
| SRR2968598\_2 |  |  |  |  |  |  | 17.1% |  |  |  |  |  |
| SRR2968598\_2\_val\_2 |  |  |  |  |  |  |  | 26.9% | 49% | 47 bp | 18% | 13.7 |
| SRR2968599 | 83.9% | 12.0 |  |  |  | 89.8% |  |  |  |  |  |  |
| SRR2968599\_1 |  |  |  |  |  |  | 1.7% |  |  |  |  |  |
| SRR2968599\_1\_val\_1 |  |  | 192.2bp | 84.2% | 12.0 |  |  | 34.6% | 49% | 48 bp | 18% | 14.3 |
| SRR2968599\_2 |  |  |  |  |  |  | 13.2% |  |  |  |  |  |
| SRR2968599\_2\_val\_2 |  |  |  |  |  |  |  | 23.6% | 49% | 47 bp | 18% | 14.3 |

Close
