## Supplementary File 3 for "FilTar: Using RNA-Seq data to improve microRNA target prediction accuracy in animals"

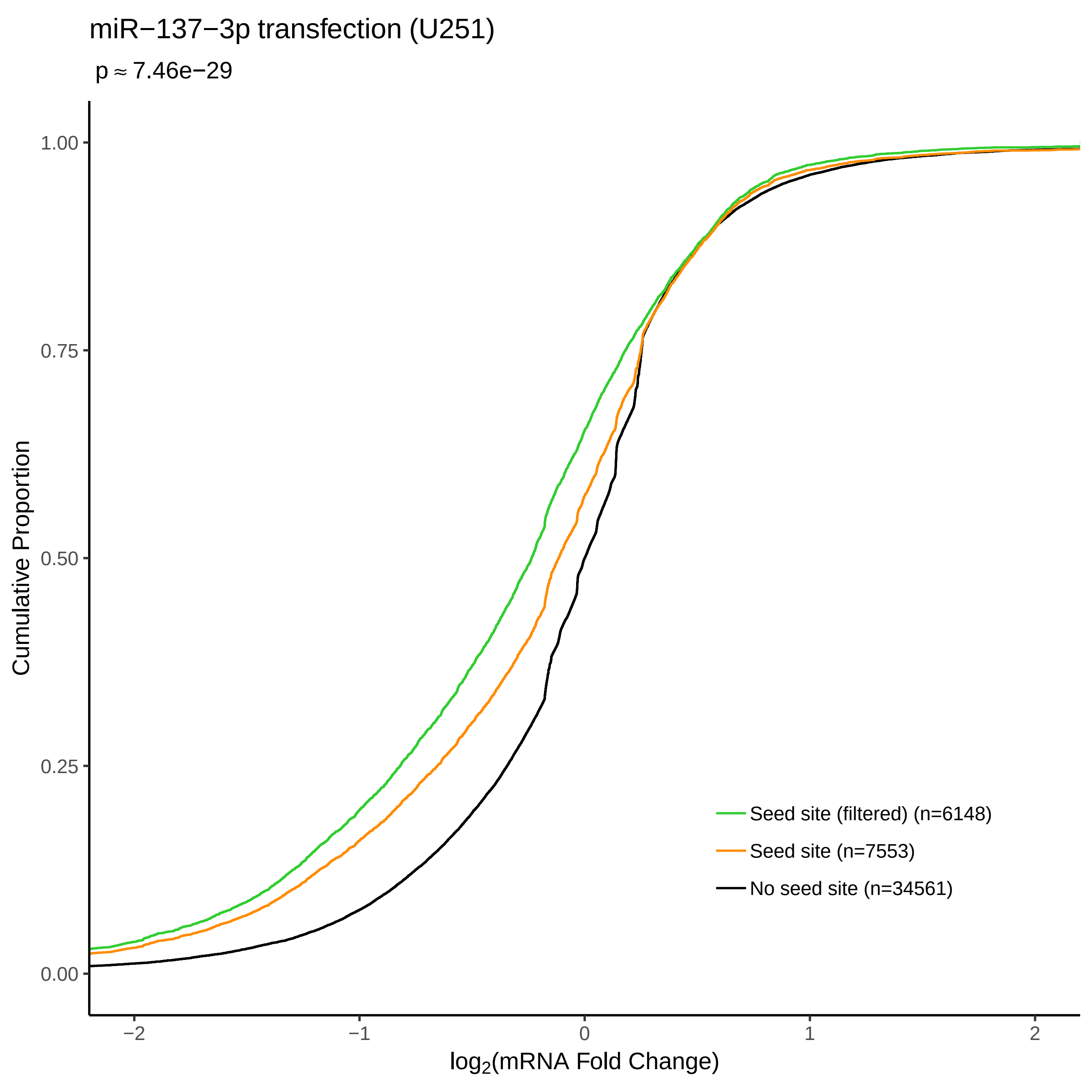

miR−137−3p transfection (U251)
p ≈ 7.46e−29
Cumulative Proportion
1.00
0.75
0.50
0.25
0.00
−2
−1
0
1
2
log2(mRNA Fold Change)
Seed site (filtered) (n=6148)
Seed site (n=7553)
No seed site (n=34561)

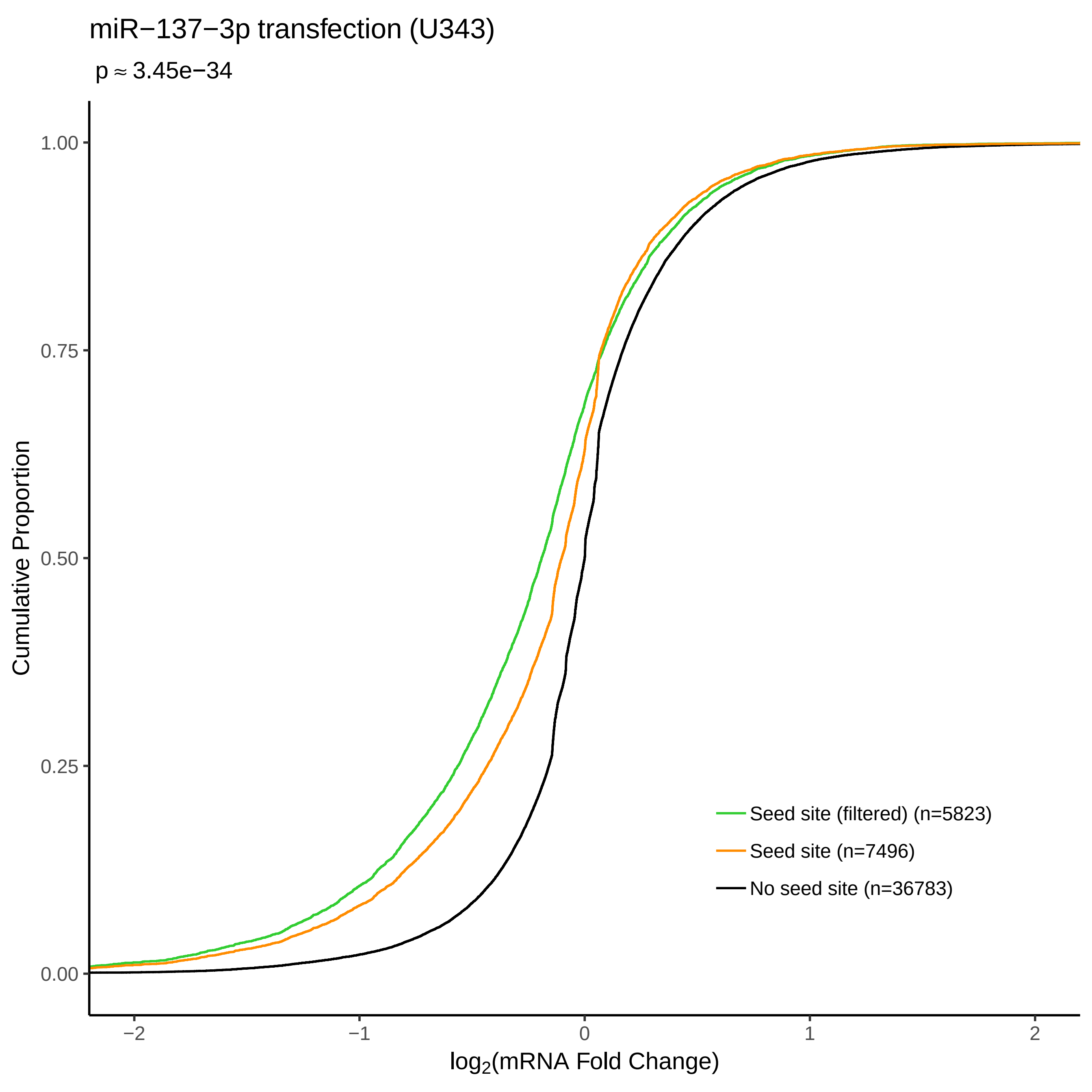

miR−137−3p transfection (U343)
p ≈ 3.45e−34
Cumulative Proportion
1.00
0.75
0.50
0.25
0.00
−2
−1
0
1
2
log2(mRNA Fold Change)
Seed site (filtered) (n=5823)
Seed site (n=7496)
No seed site (n=36783)

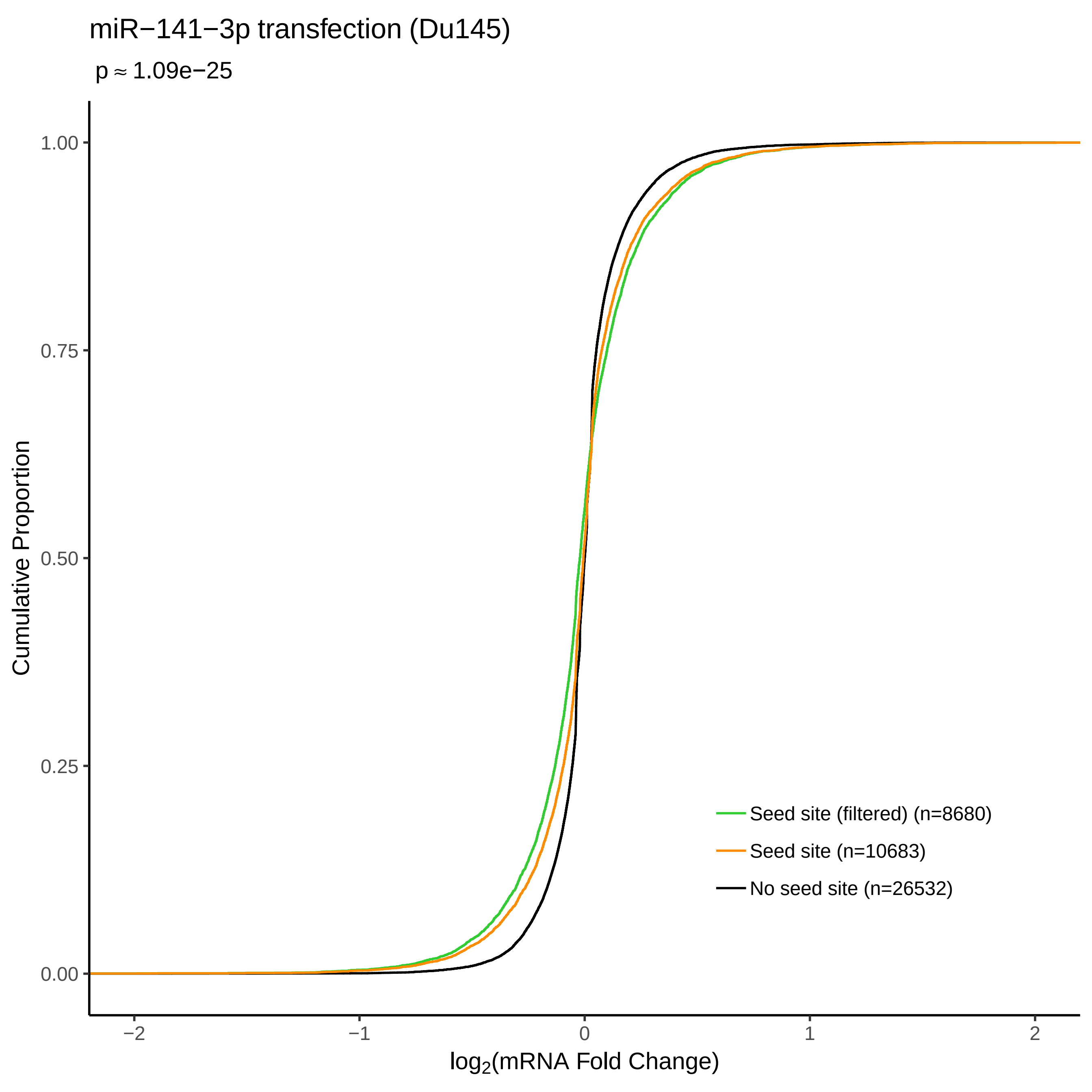
miR−141−3p transfection (Du145)
p ≈ 1.09e−25
Cumulative Proportion
1.00
0.75
0.50
0.25
0.00
−2
−1
0
1
2
log2(mRNA Fold Change)
Seed site (filtered) (n=8680)
Seed site (n=10683)
No seed site (n=26532)

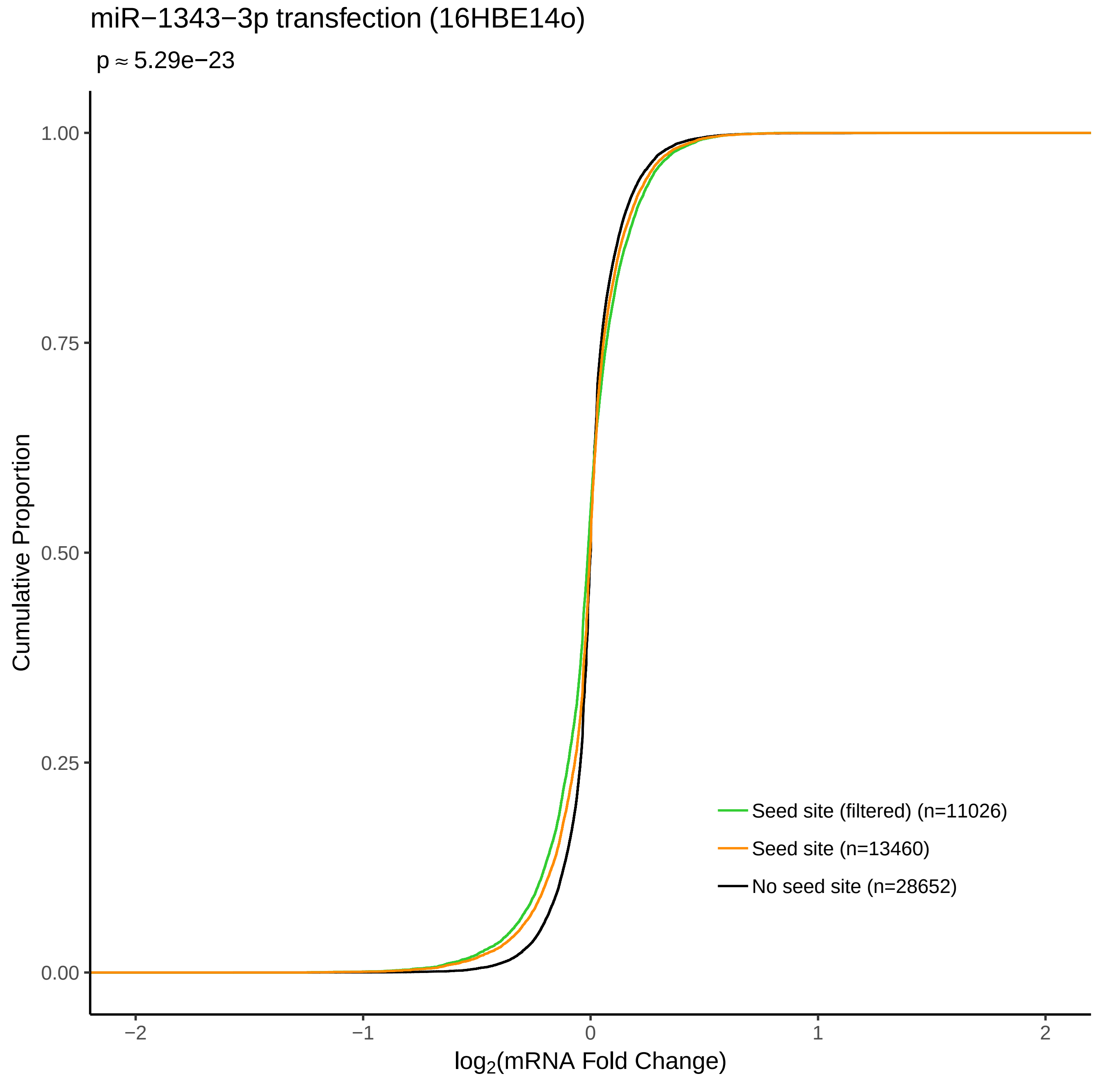
miR−1343−3p transfection (16HBE14o)
p ≈ 5.29e−23
Cumulative Proportion
1.00
0.75
0.50
0.25
0.00
−2
−1
0
1
2
log2(mRNA Fold Change)
Seed site (filtered) (n=11026)
Seed site (n=13460)
No seed site (n=28652)

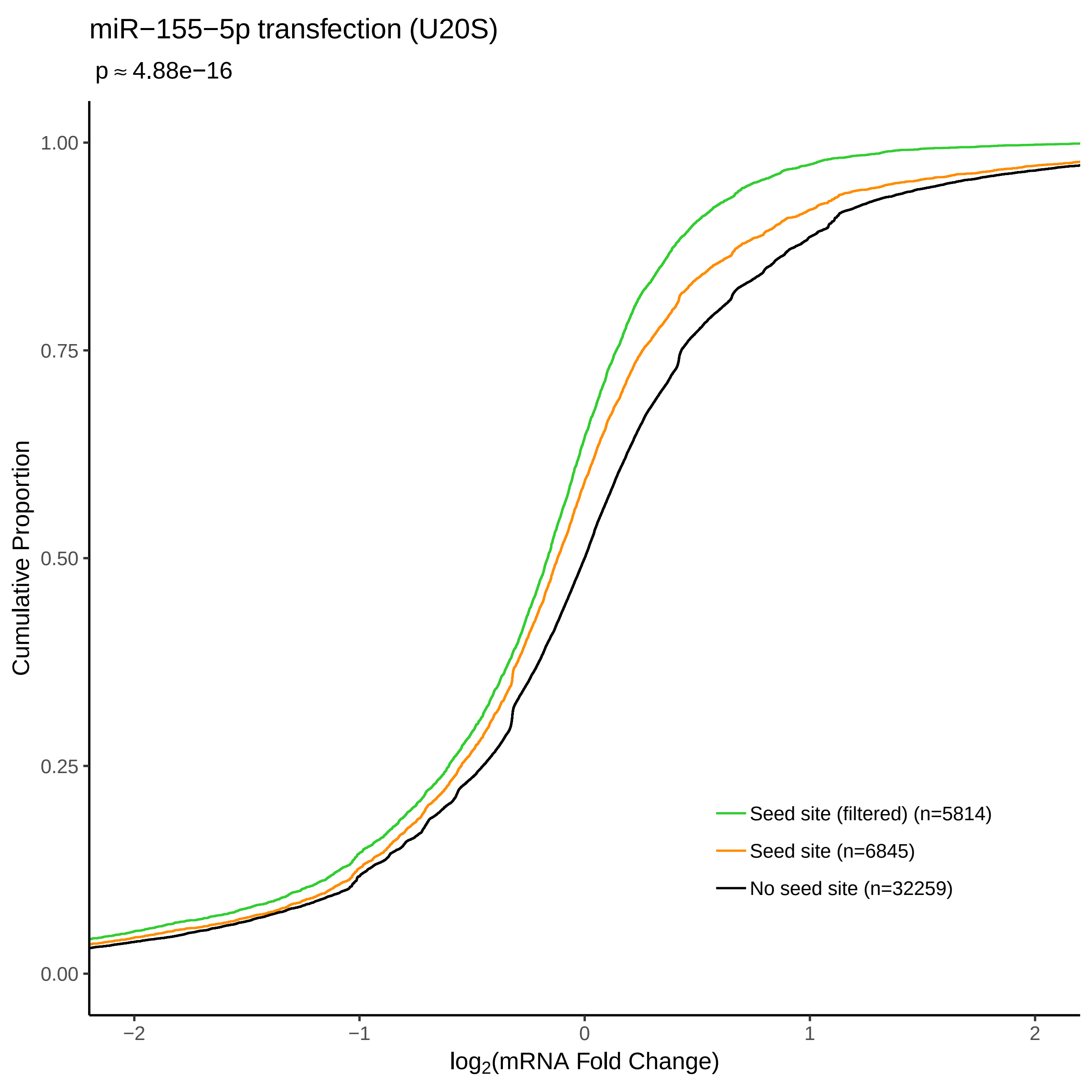

miR−155−5p transfection (U20S)
p ≈ 4.88e−16
Cumulative Proportion
1.00
0.75
0.50
0.25
0.00
−2
−1
0
1
2
log2(mRNA Fold Change)
Seed site (filtered) (n=5814)
Seed site (n=6845)
No seed site (n=32259)

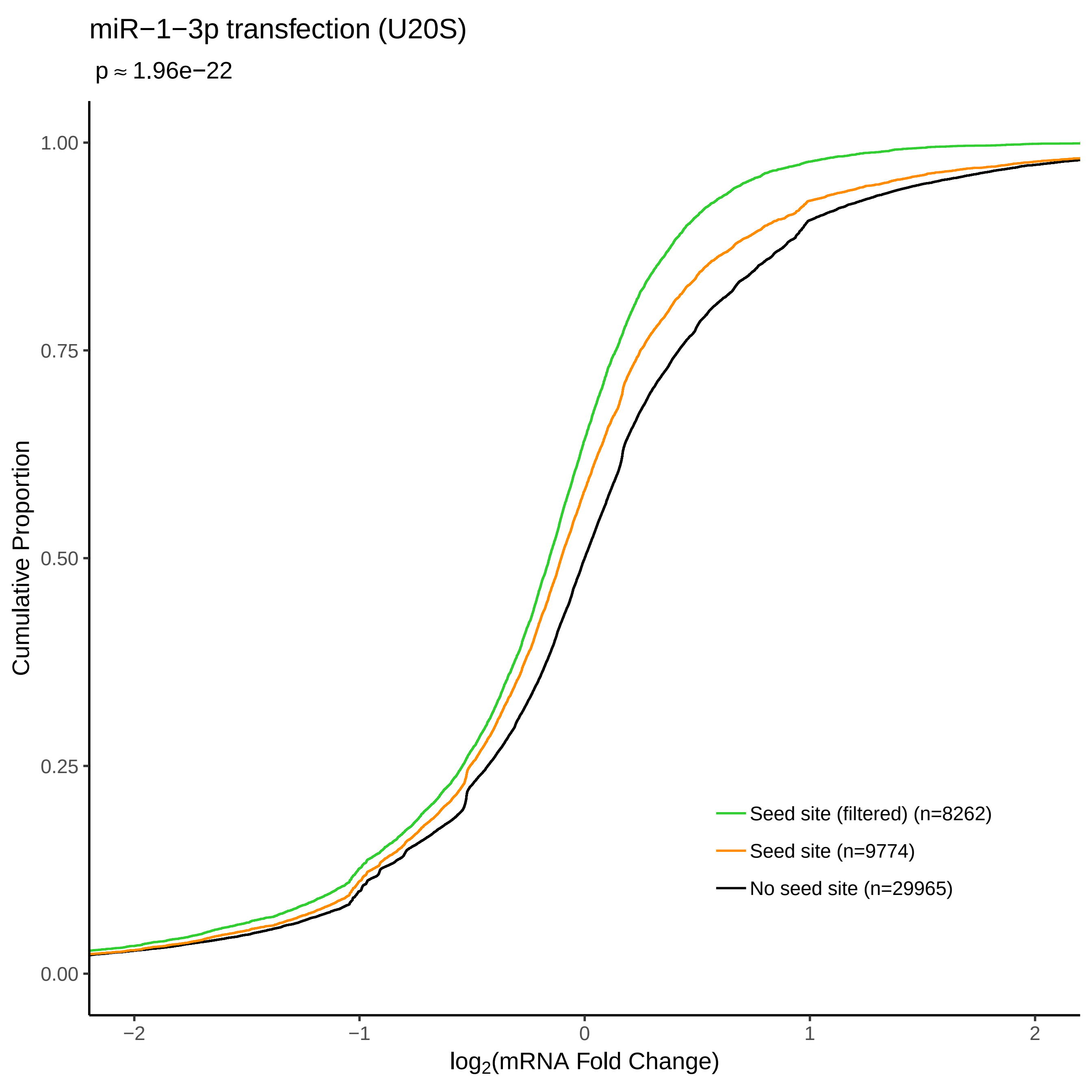

miR−1−3p transfection (U20S)
p ≈ 1.96e−22
Cumulative Proportion
1.00
0.75
0.50
0.25
0.00
−2
−1
0
1
2
log2(mRNA Fold Change)
Seed site (filtered) (n=8262)
Seed site (n=9774)
No seed site (n=29965)

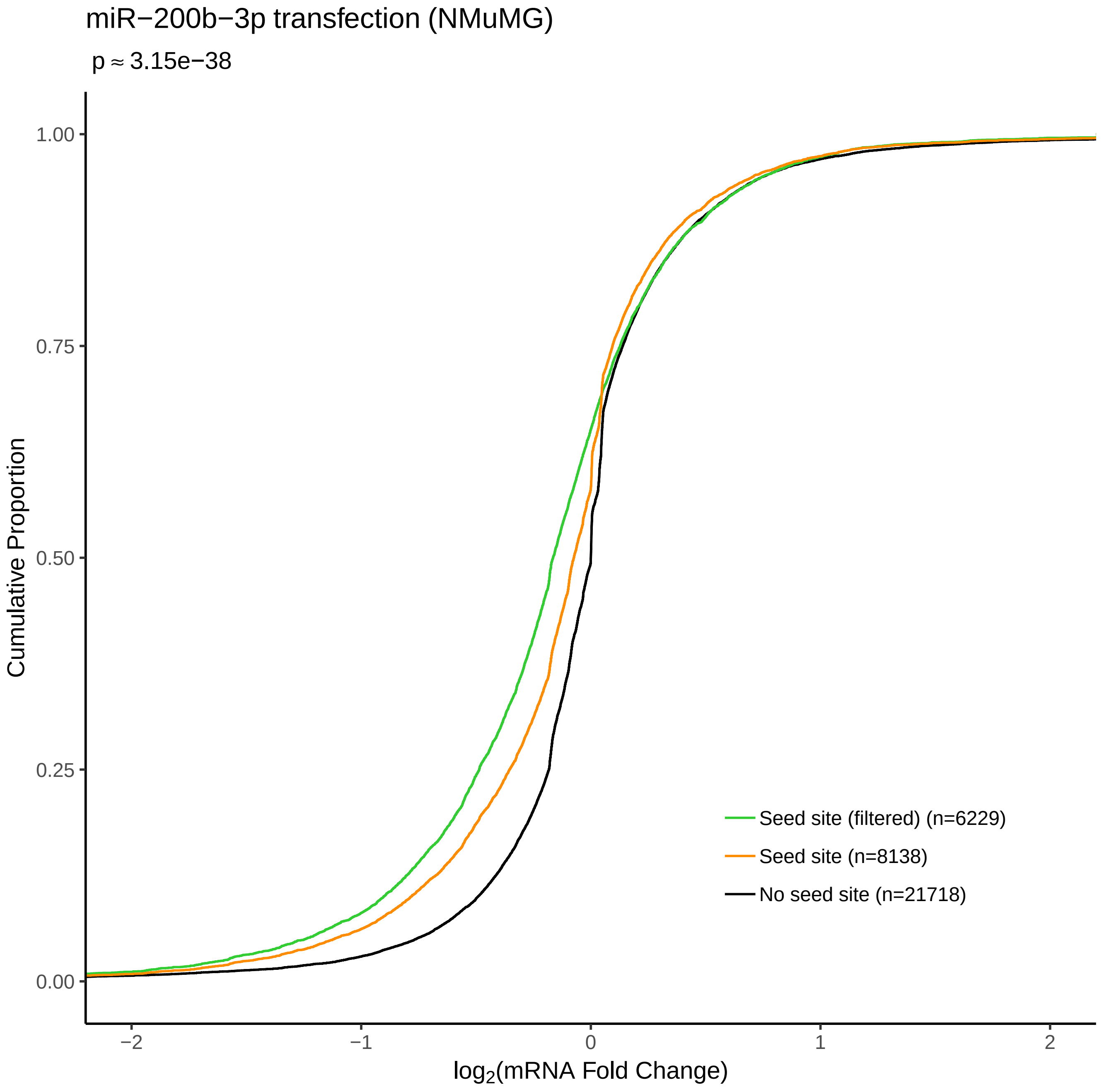
miR−200b−3p transfection (NMuMG)
p ≈ 3.15e−38
Cumulative Proportion
1.00
0.75
0.50
0.25
0.00
−2
−1
0
1
2
log2(mRNA Fold Change)
Seed site (filtered) (n=6229)
Seed site (n=8138)
No seed site (n=21718)

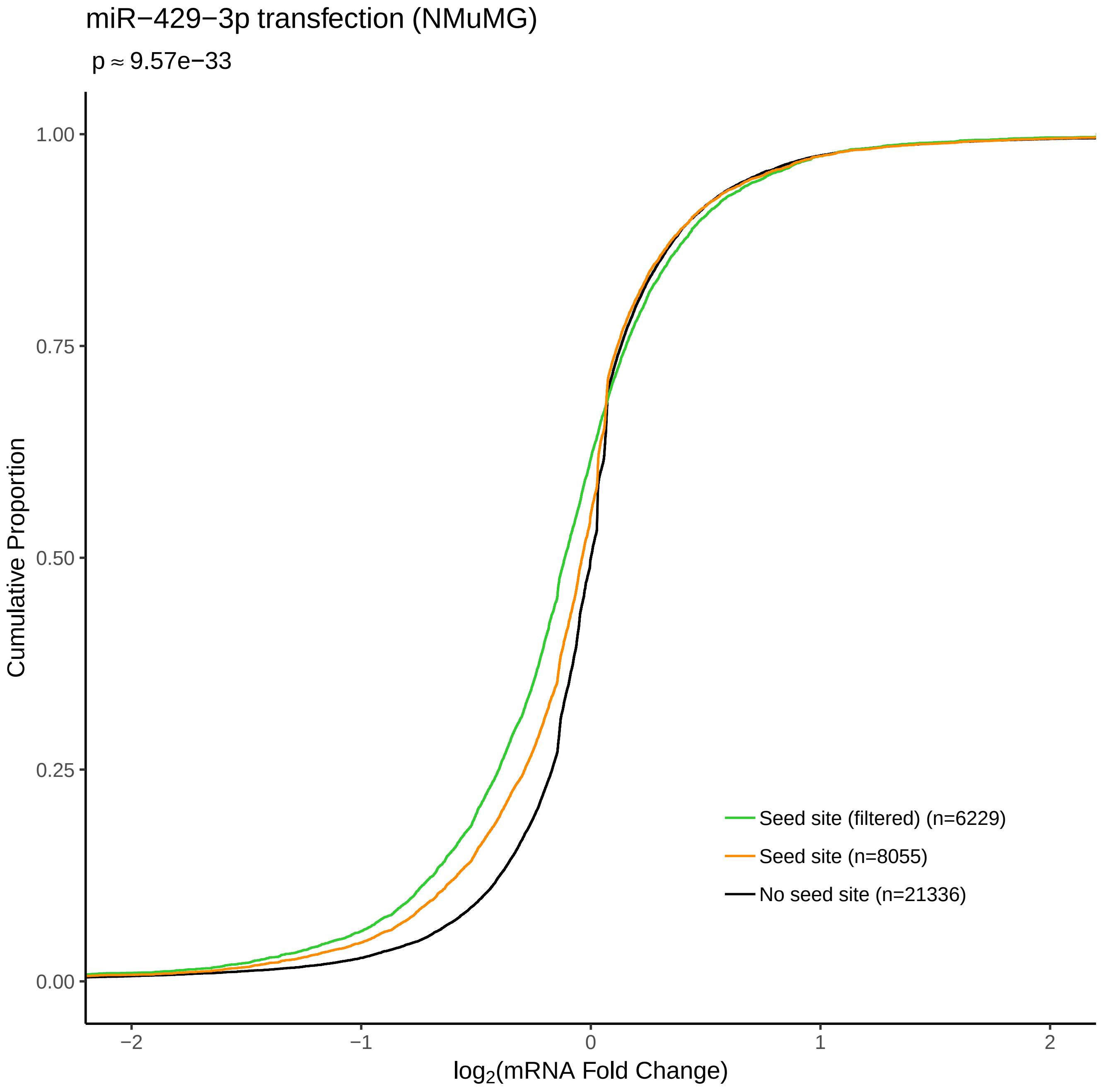
miR−429−3p transfection (NMuMG)
p ≈ 9.57e−33
Cumulative Proportion
1.00
0.75
0.50
0.25
0.00
−2
−1
0
1
2
log2(mRNA Fold Change)
Seed site (filtered) (n=6229)
Seed site (n=8055)
No seed site (n=21336)

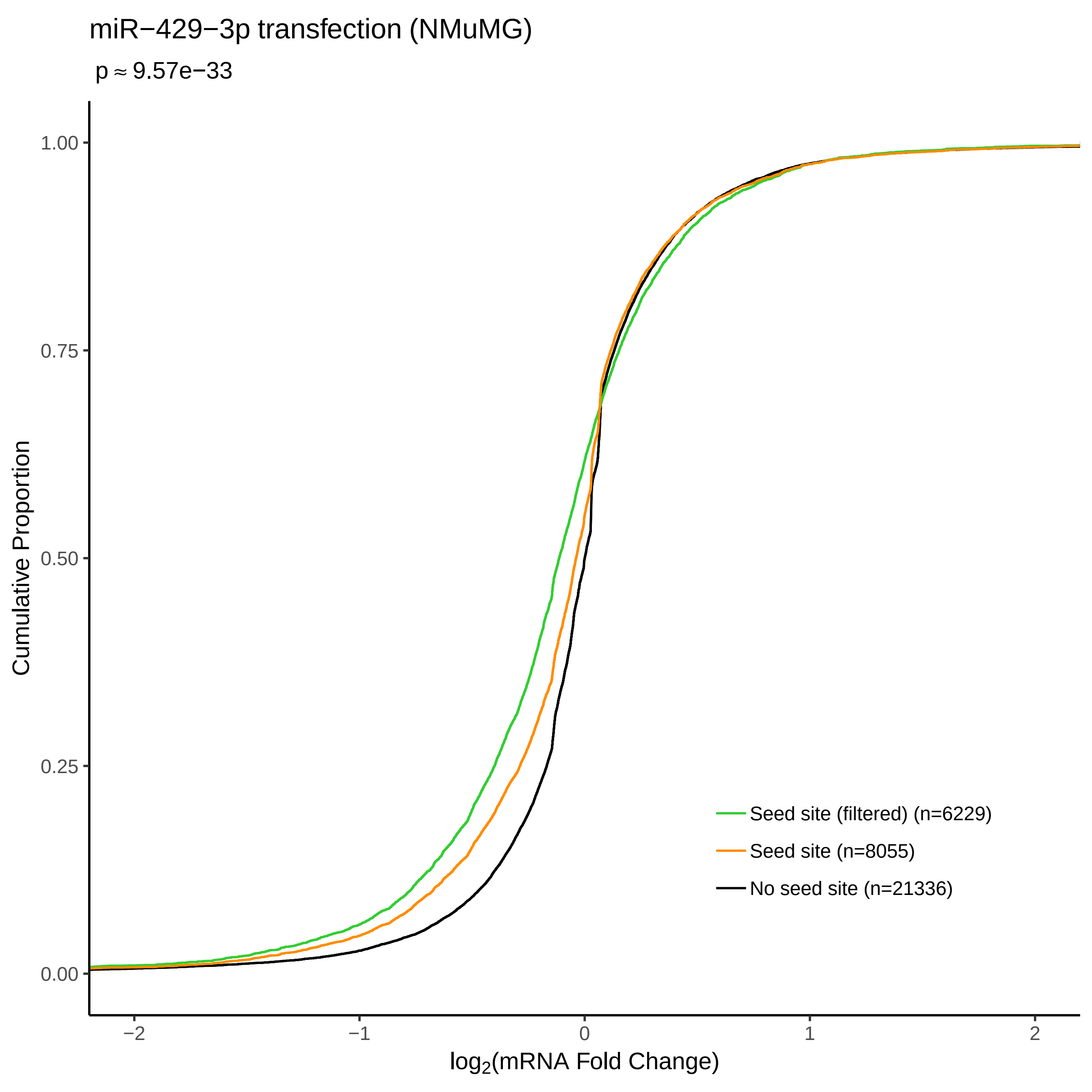

miR−429−3p transfection (NMuMG)
p ≈ 9.57e−33
Cumulative Proportion
1.00
0.75
0.50
0.25
0.00
−2
−1
0
1
2
log2(mRNA Fold Change)
Seed site (filtered) (n=6229)
Seed site (n=8055)
No seed site (n=21336)

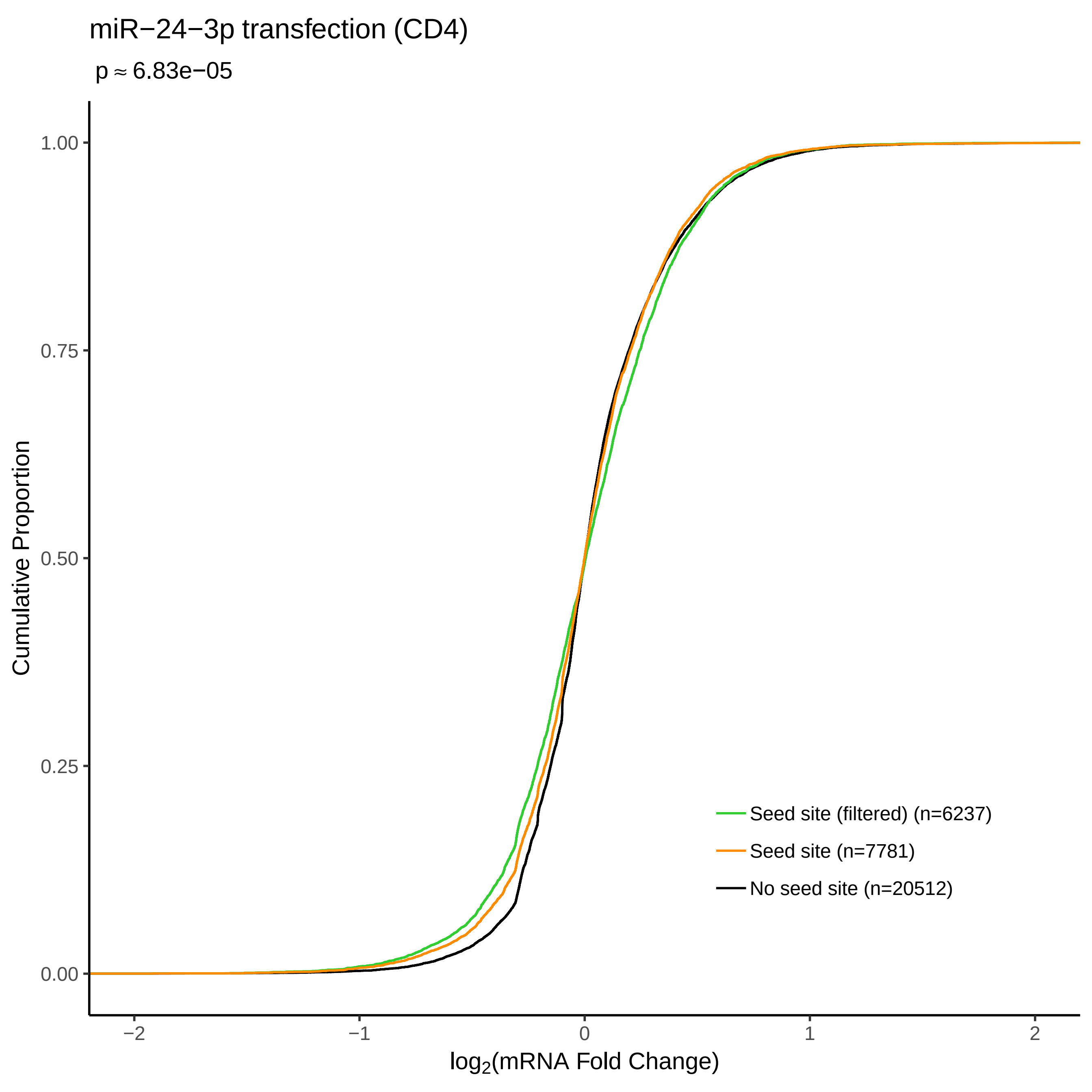

miR−24−3p transfection (CD4)
p ≈ 6.83e−05
Cumulative Proportion
1.00
0.75
0.50
0.25
0.00
−2
−1
0
1
2
log2(mRNA Fold Change)
Seed site (filtered) (n=6237)
Seed site (n=7781)
No seed site (n=20512)

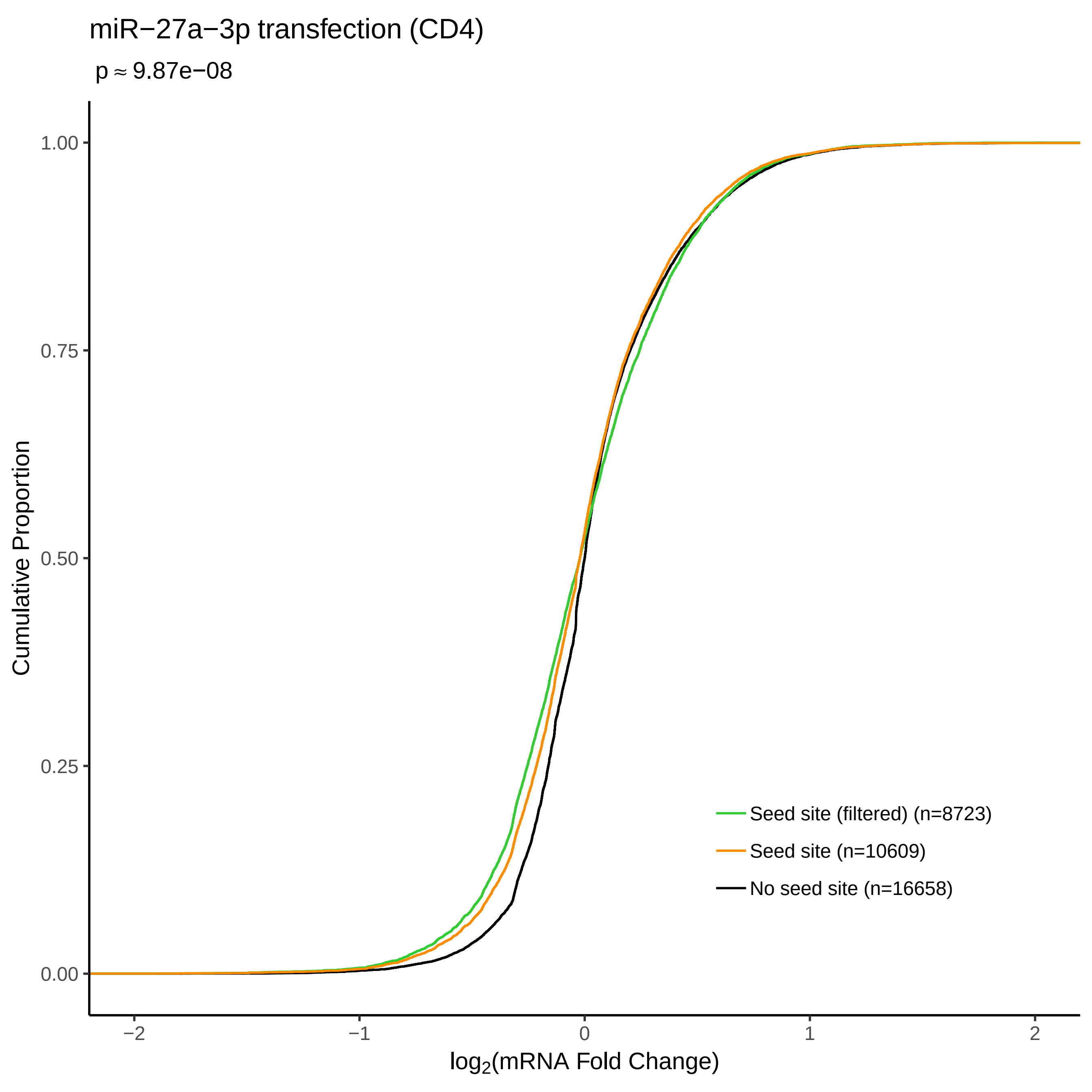
miR−27a−3p transfection (CD4)
p ≈ 9.87e−08
Cumulative Proportion
1.00
0.75
0.50
0.25
0.00
−2
−1
0
1
2
log2(mRNA Fold Change)
Seed site (filtered) (n=8723)
Seed site (n=10609)
No seed site (n=16658)

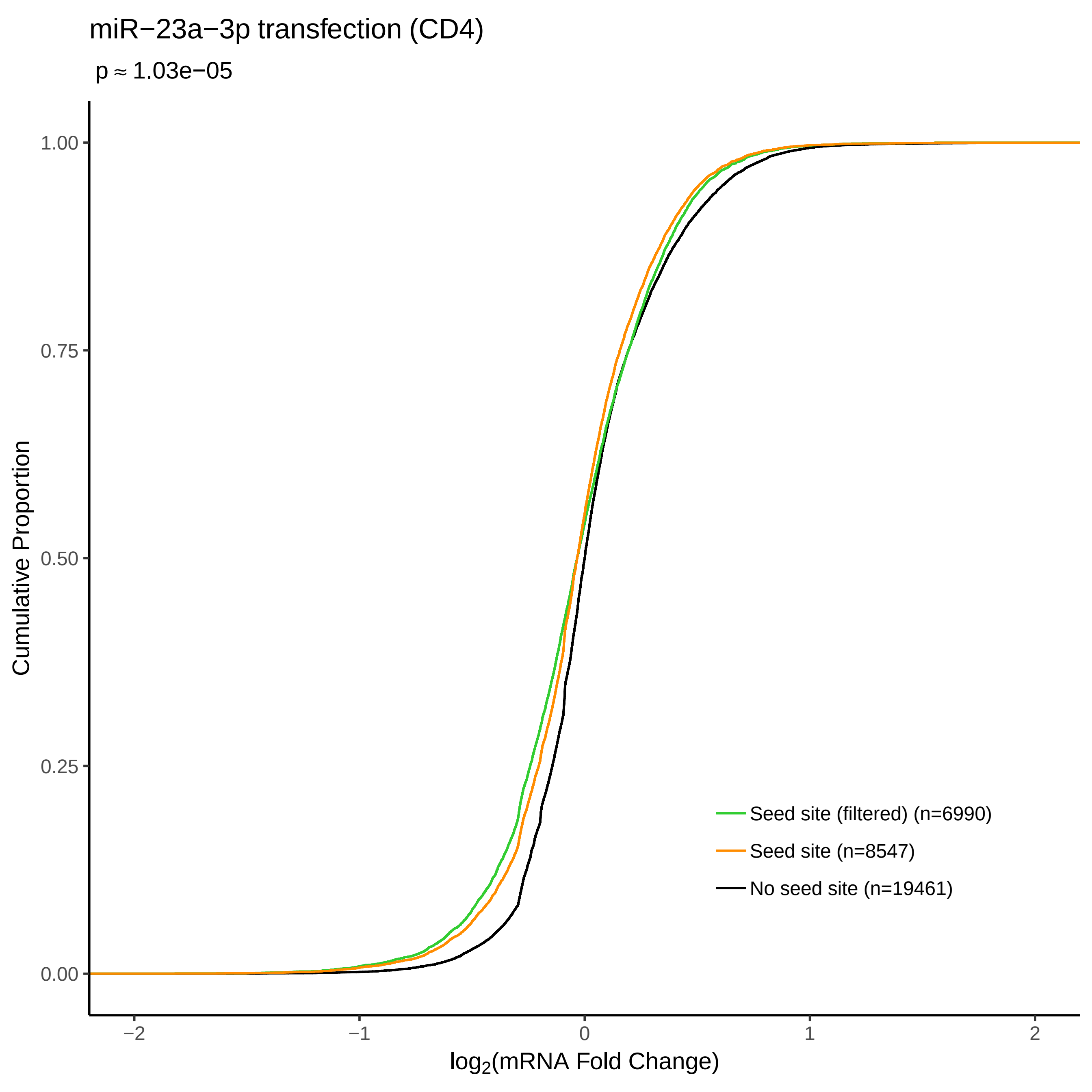
miR−23a−3p transfection (CD4)
p ≈ 1.03e−05
Cumulative Proportion
1.00
0.75
0.50
0.25
0.00
−2
−1
0
1
2
log2(mRNA Fold Change)
Seed site (filtered) (n=6990)
Seed site (n=8547)
No seed site (n=19461)

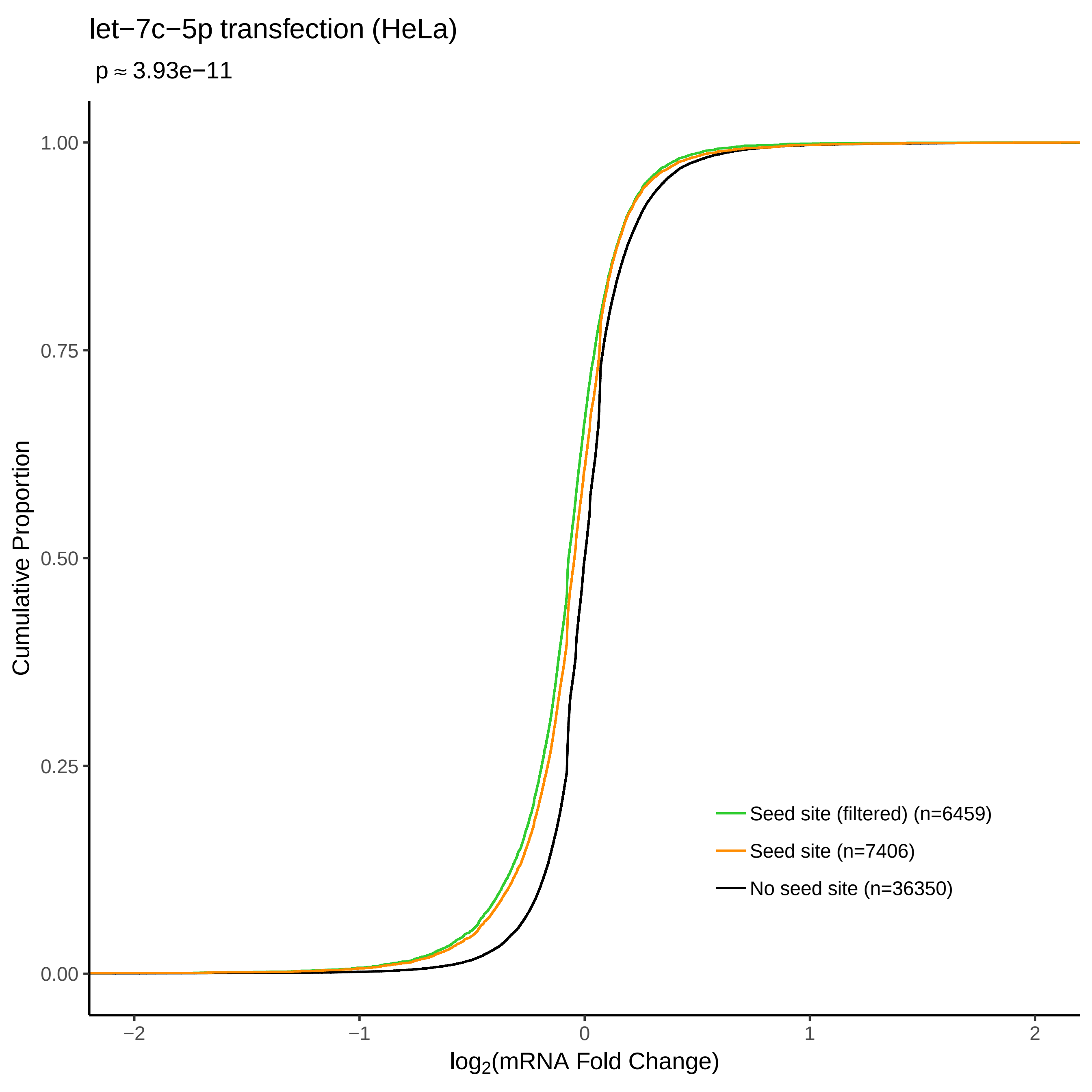

let−7c−5p transfection (HeLa)
p ≈ 3.93e−11
Cumulative Proportion
1.00
0.75
0.50
0.25
0.00
−2
−1
0
1
2
log2(mRNA Fold Change)
Seed site (filtered) (n=6459)
Seed site (n=7406)
No seed site (n=36350)

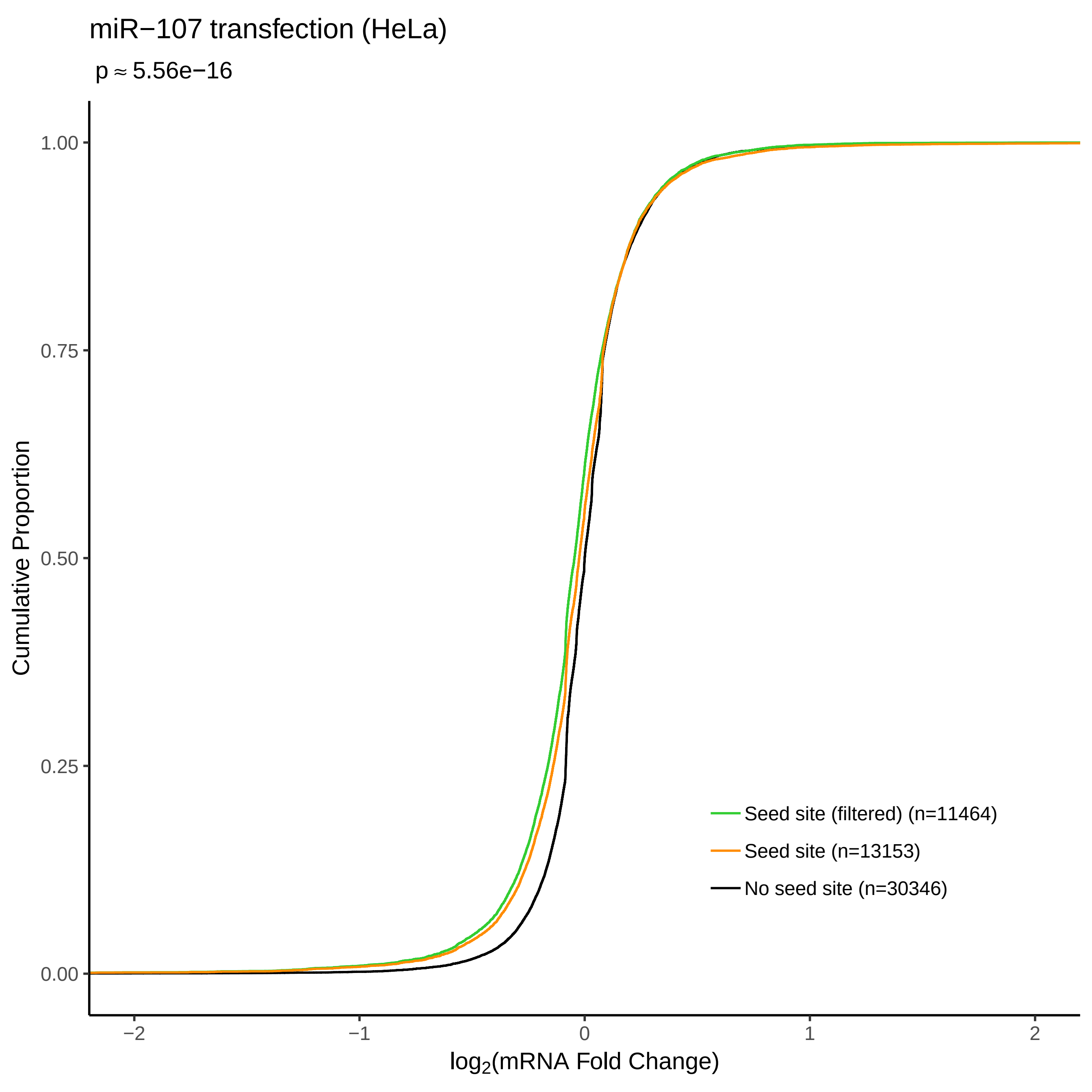

miR−107 transfection (HeLa)
p ≈ 5.56e−16
Cumulative Proportion
1.00
0.75
0.50
0.25
0.00
−2
−1
0
1
2
log2(mRNA Fold Change)
Seed site (filtered) (n=11464)
Seed site (n=13153)
No seed site (n=30346)

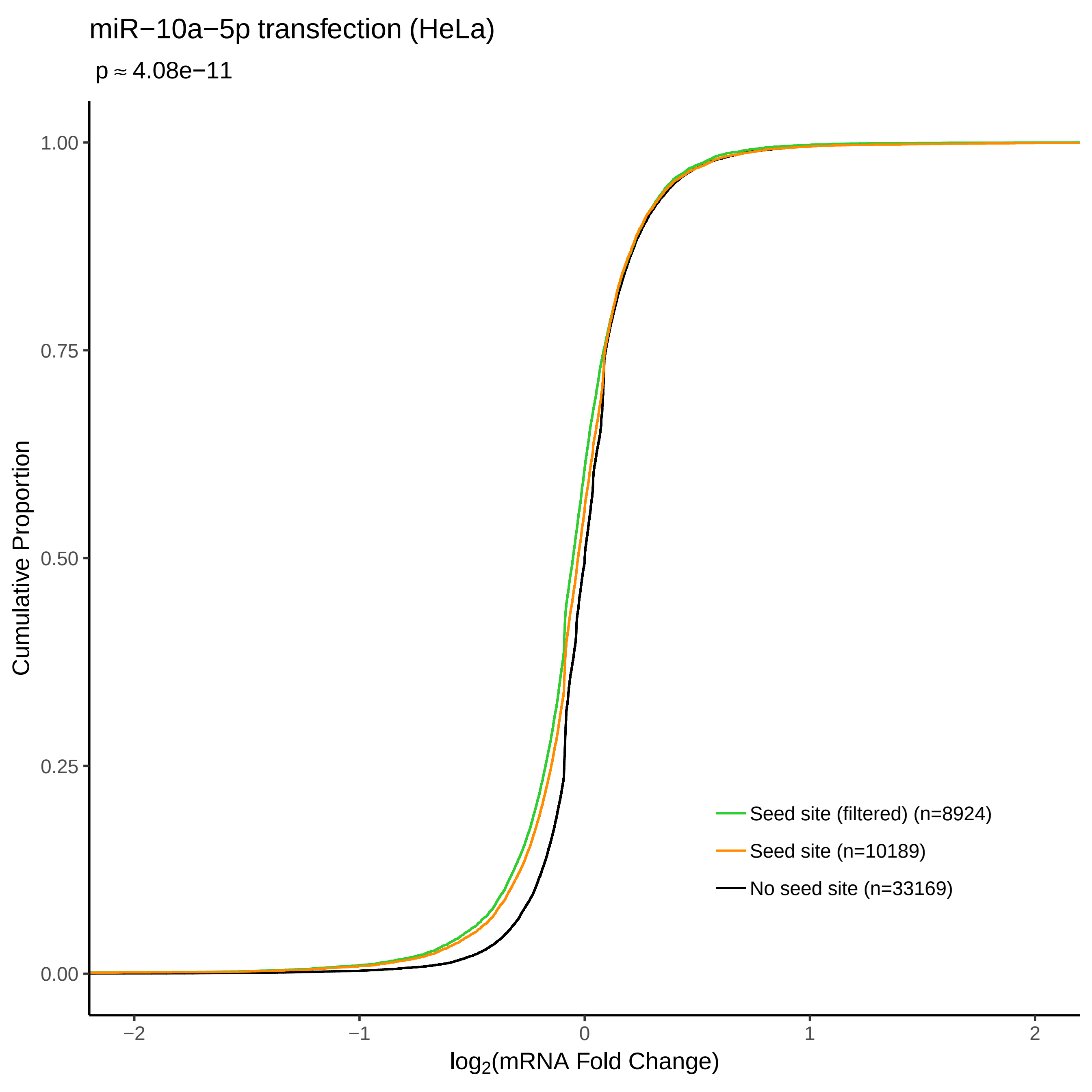

miR−10a−5p transfection (HeLa)
p ≈ 4.08e−11
Cumulative Proportion
1.00
0.75
0.50
0.25
0.00
−2
−1
0
1
2
log2(mRNA Fold Change)
Seed site (filtered) (n=8924)
Seed site (n=10189)
No seed site (n=33169)

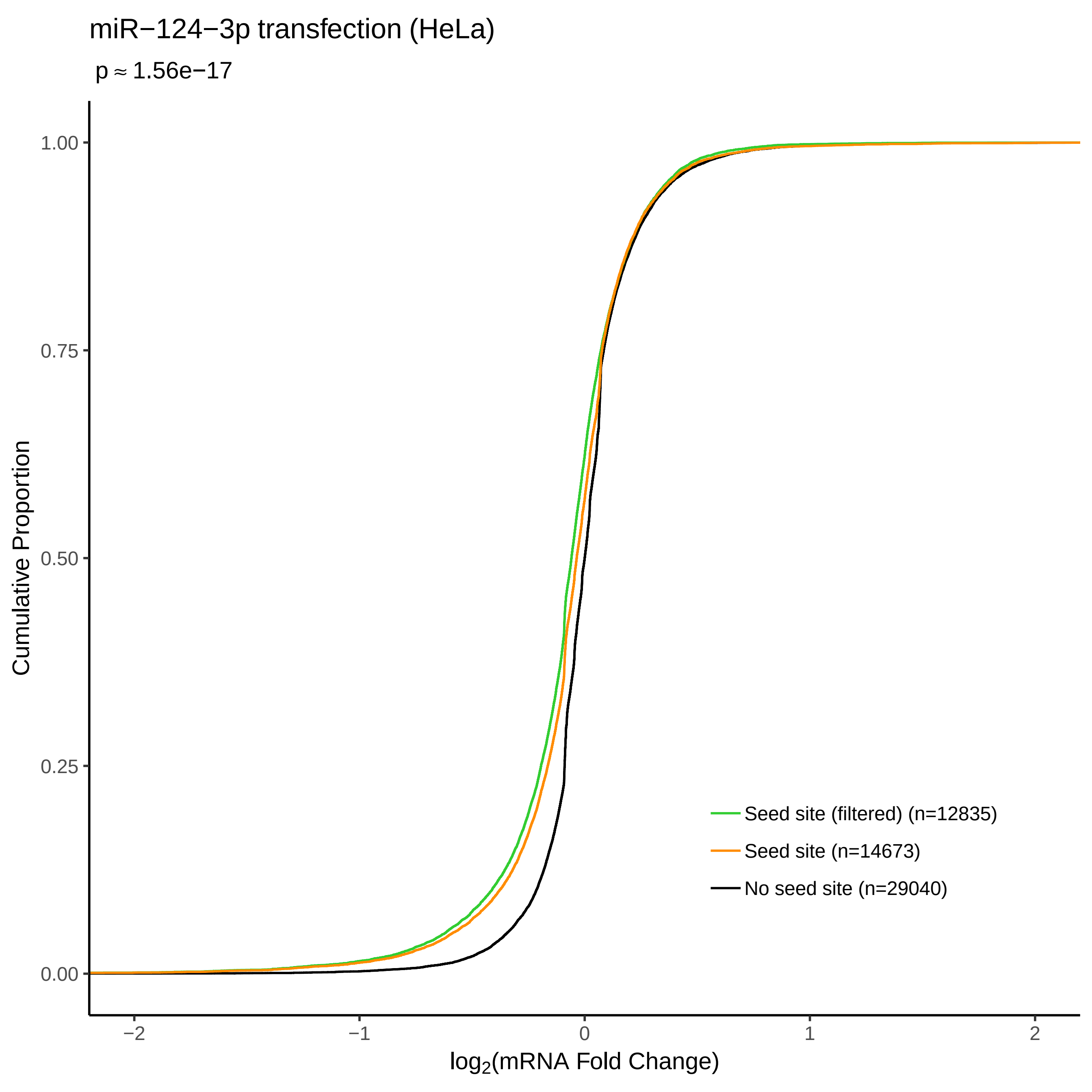

miR−124−3p transfection (HeLa)
p ≈ 1.56e−17
Cumulative Proportion
1.00
0.75
0.50
0.25
0.00
−2
−1
0
1
2
log2(mRNA Fold Change)
Seed site (filtered) (n=12835)
Seed site (n=14673)
No seed site (n=29040)

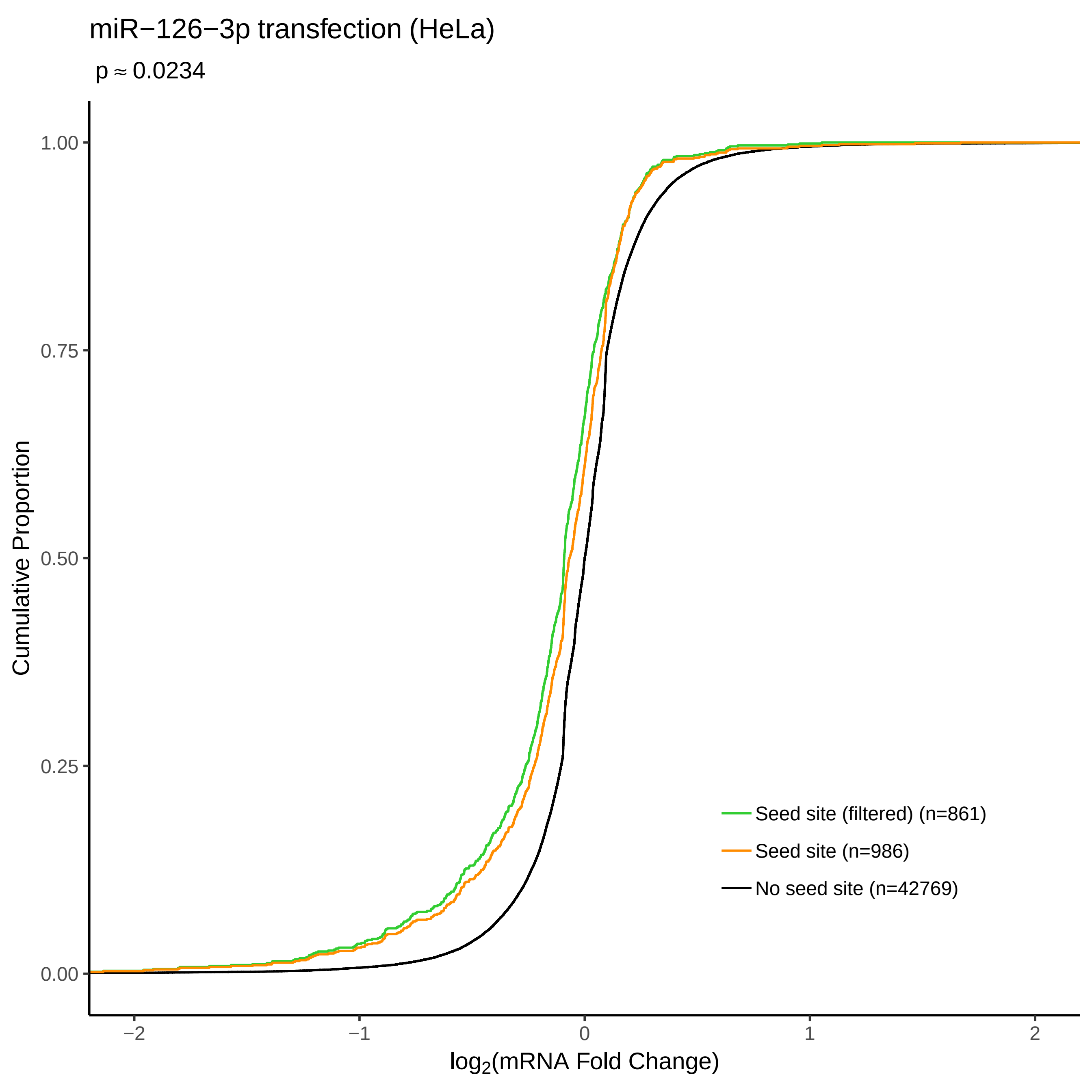

miR−126−3p transfection (HeLa)
p ≈ 0.0234
Cumulative Proportion
1.00
0.75
0.50
0.25
0.00
−2
−1
0
1
2
log2(mRNA Fold Change)
Seed site (filtered) (n=861)
Seed site (n=986)
No seed site (n=42769)

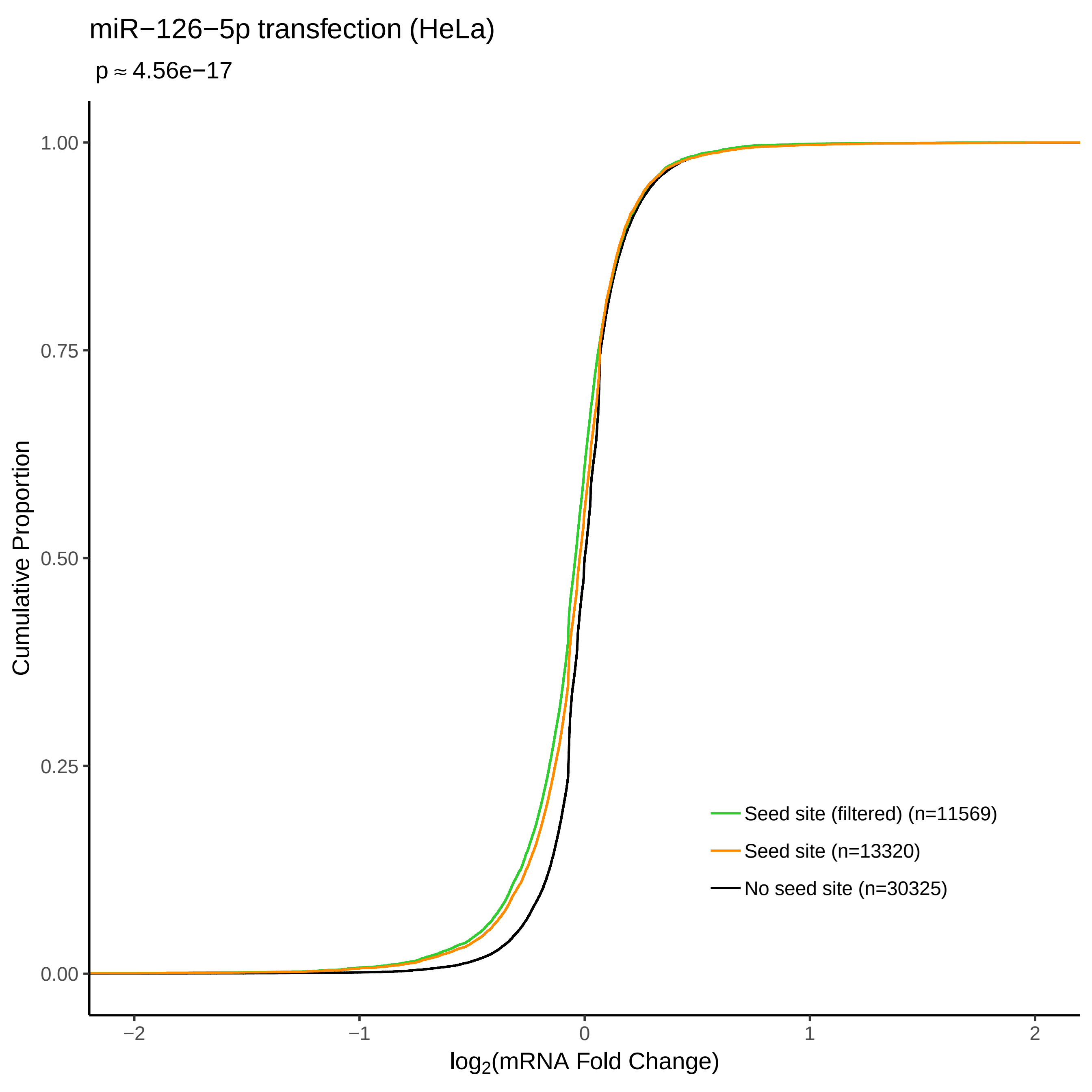

miR−126−5p transfection (HeLa)
p ≈ 4.56e−17
Cumulative Proportion
1.00
0.75
0.50
0.25
0.00
−2
−1
0
1
2
log2(mRNA Fold Change)
Seed site (filtered) (n=11569)
Seed site (n=13320)
No seed site (n=30325)

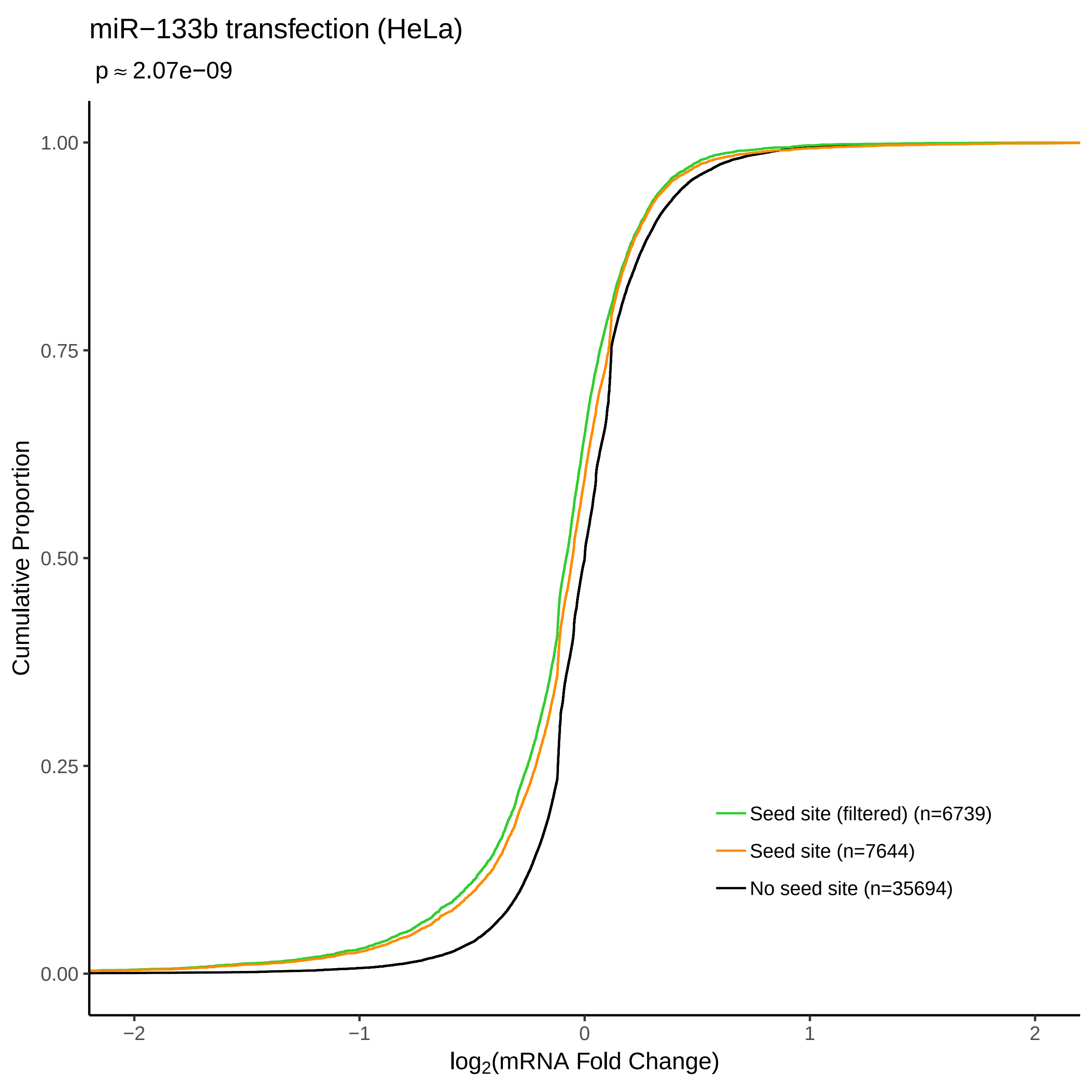

miR−133b transfection (HeLa)
p ≈ 2.07e−09
Cumulative Proportion
1.00
0.75
0.50
0.25
0.00
−2
−1
0
1
2
log2(mRNA Fold Change)
Seed site (filtered) (n=6739)
Seed site (n=7644)
No seed site (n=35694)

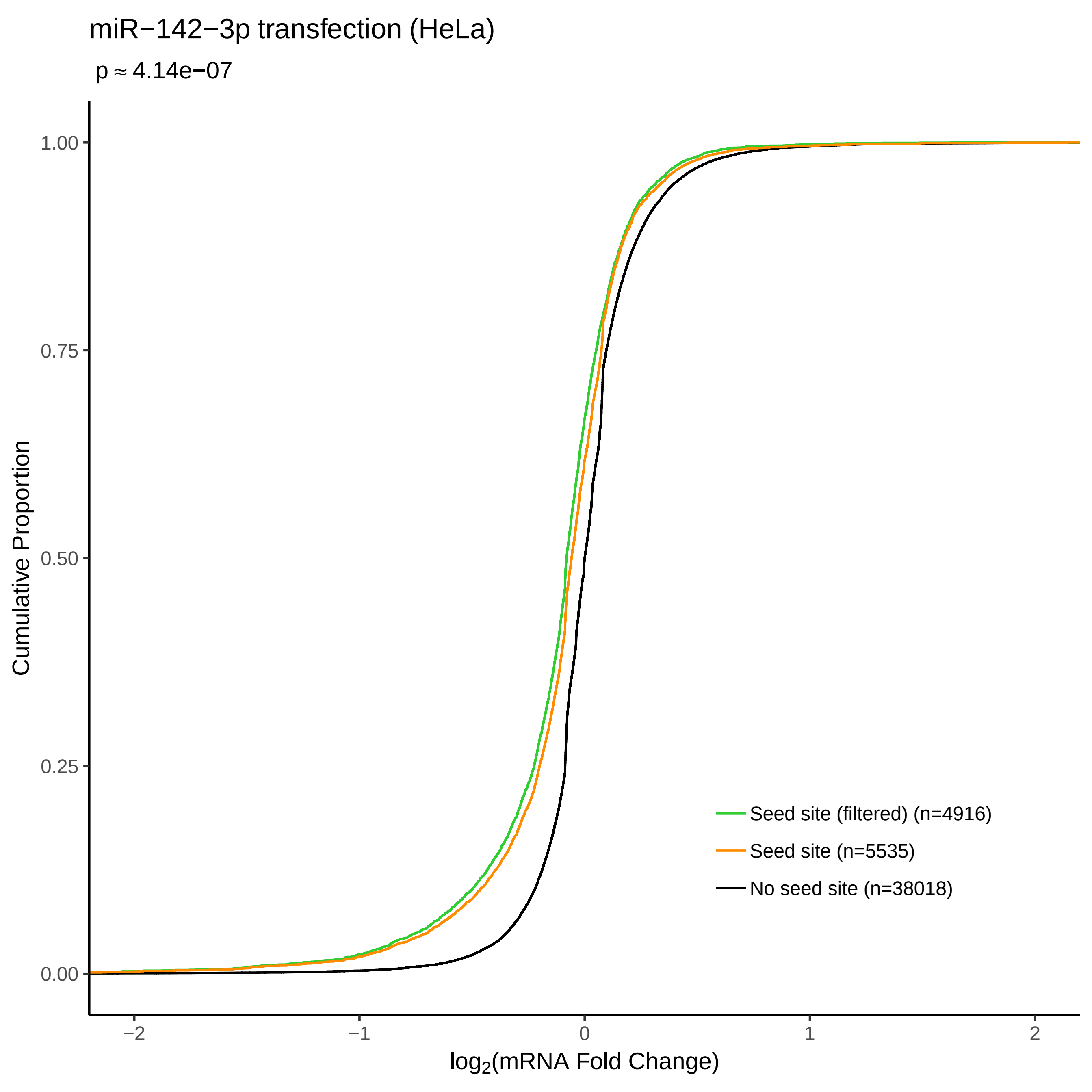

miR−142−3p transfection (HeLa)
p ≈ 4.14e−07
Cumulative Proportion
1.00
0.75
0.50
0.25
0.00
−2
−1
0
1
2
log2(mRNA Fold Change)
Seed site (filtered) (n=4916)
Seed site (n=5535)
No seed site (n=38018)

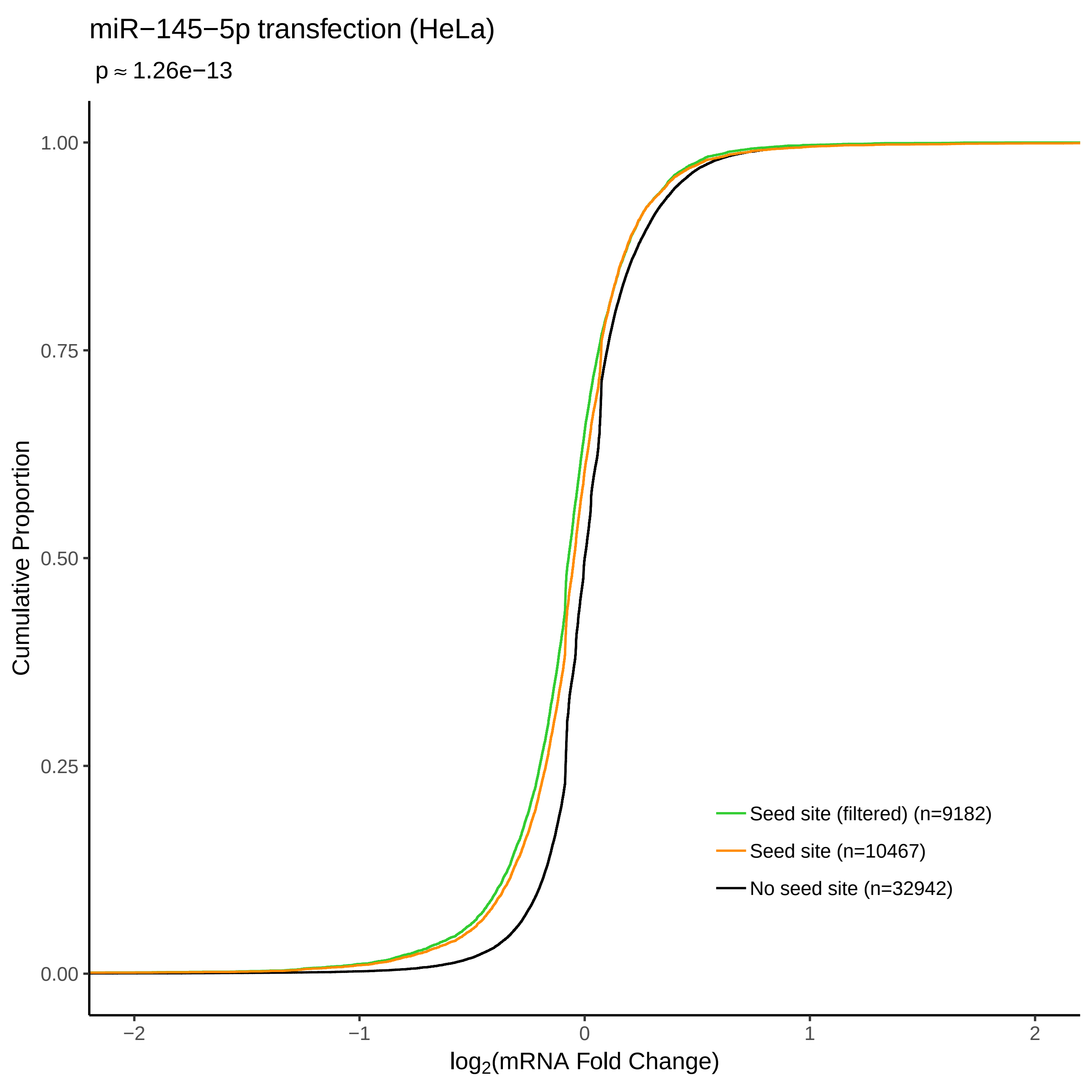

miR−145−5p transfection (HeLa)
p ≈ 1.26e−13
Cumulative Proportion
0.00
0.25
0.50
0.75
1.00
−2
−1
0
1
2
log2(mRNA Fold Change)
Seed site (filtered) (n=9182)
Seed site (n=10467)
No seed site (n=32942)

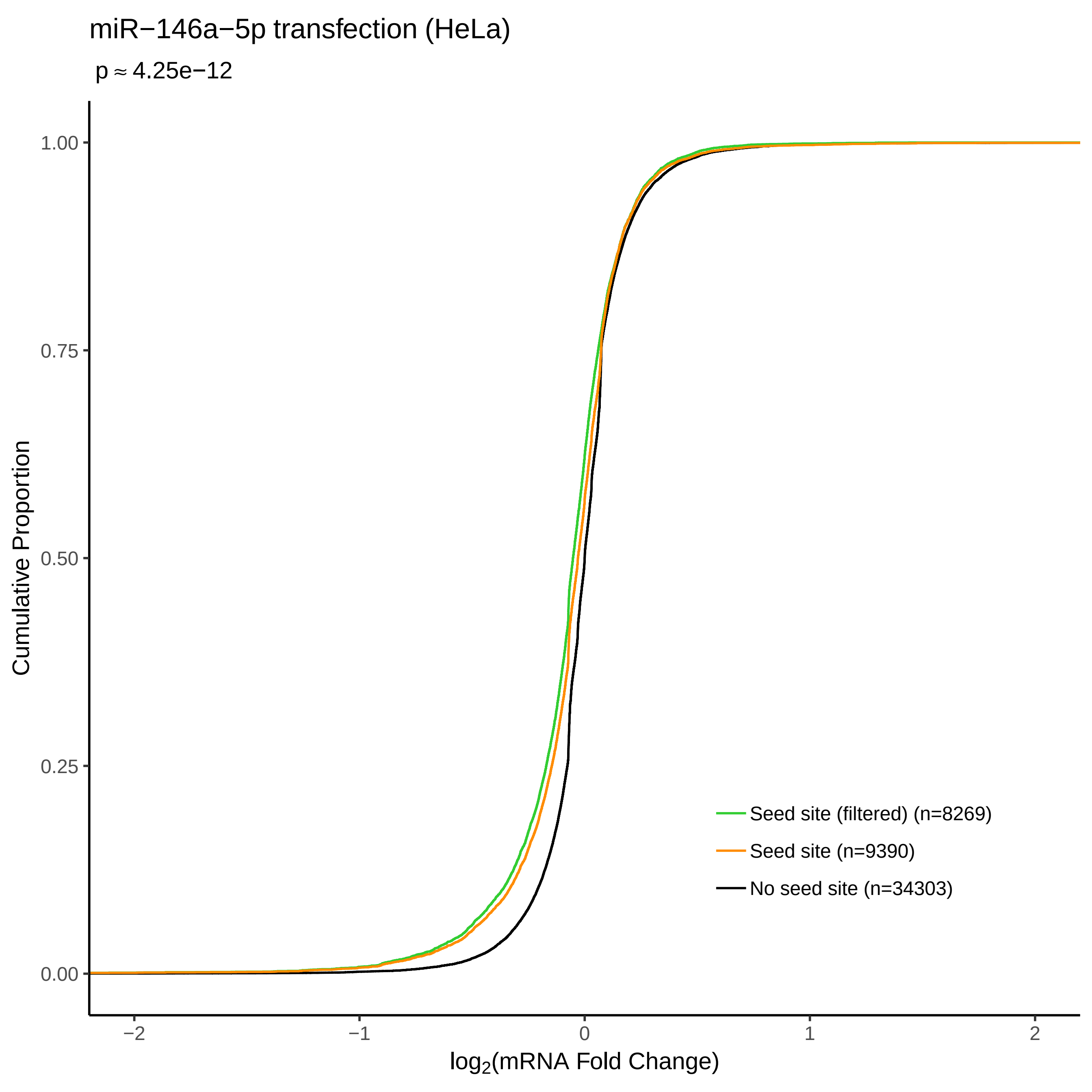

miR−146a−5p transfection (HeLa)
p ≈ 4.25e−12
Cumulative Proportion
1.00
0.75
0.50
0.25
0.00
−2
−1
0
1
2
log2(mRNA Fold Change)
Seed site (filtered) (n=8269)
Seed site (n=9390)
No seed site (n=34303)

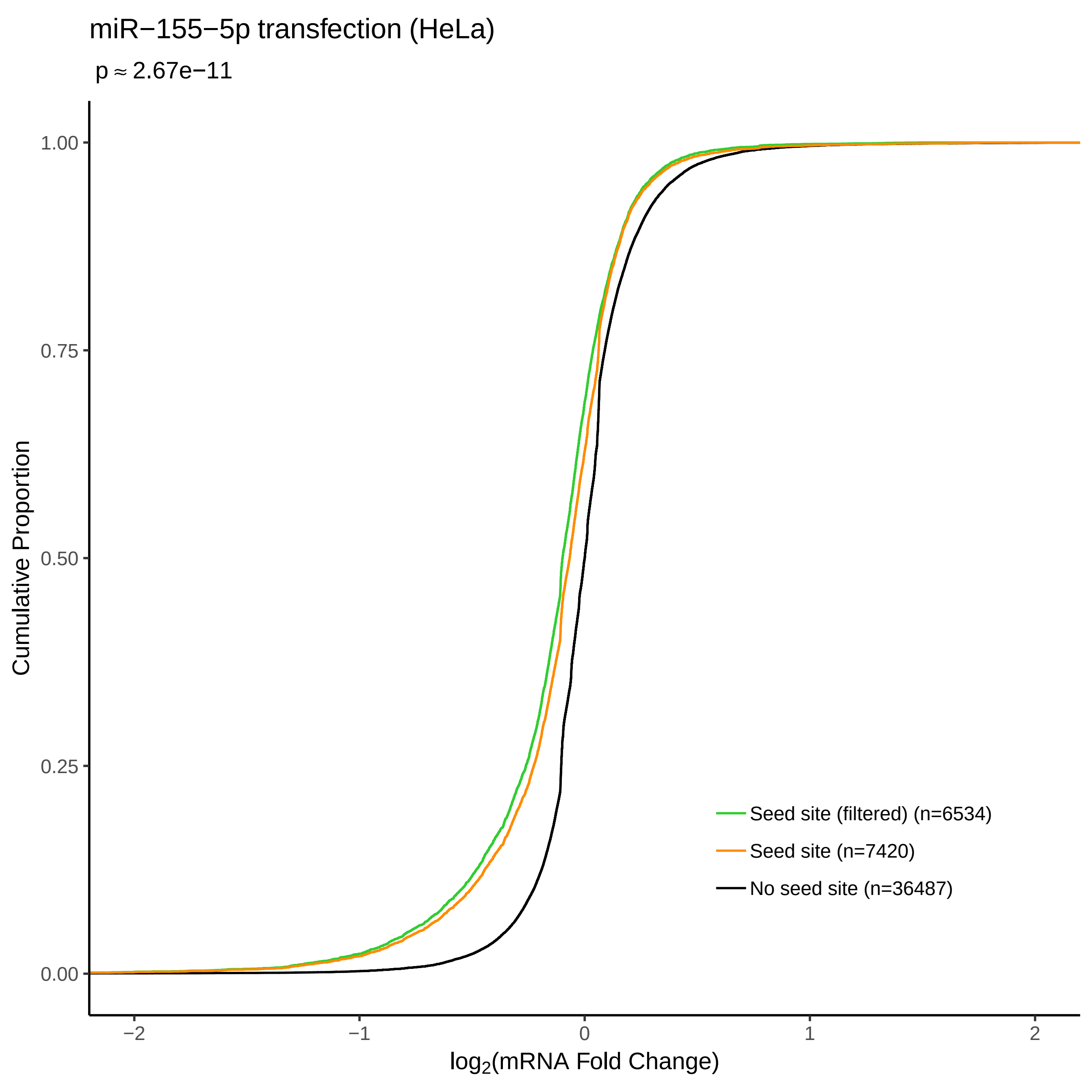
miR−155−5p transfection (HeLa)
p ≈ 2.67e−11
Cumulative Proportion
1.00
0.75
0.50
0.25
0.00
−2
−1
0
1
2
log2(mRNA Fold Change)
Seed site (filtered) (n=6534)
Seed site (n=7420)
No seed site (n=36487)

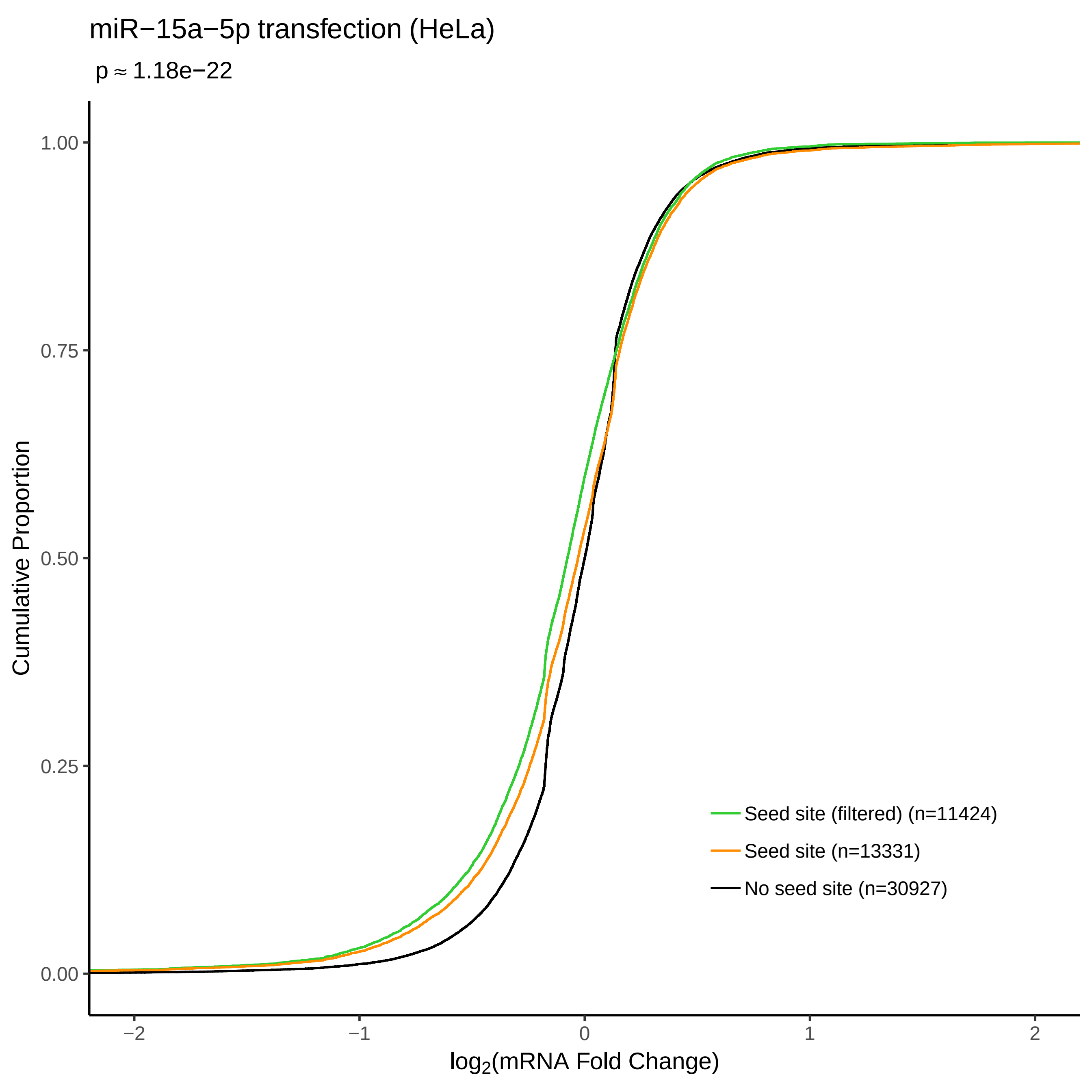

miR−15a−5p transfection (HeLa)
p ≈ 1.18e−22
Cumulative Proportion
1.00
0.75
0.50
0.25
0.00
−2
−1
0
1
2
log2(mRNA Fold Change)
Seed site (filtered) (n=11424)
Seed site (n=13331)
No seed site (n=30927)

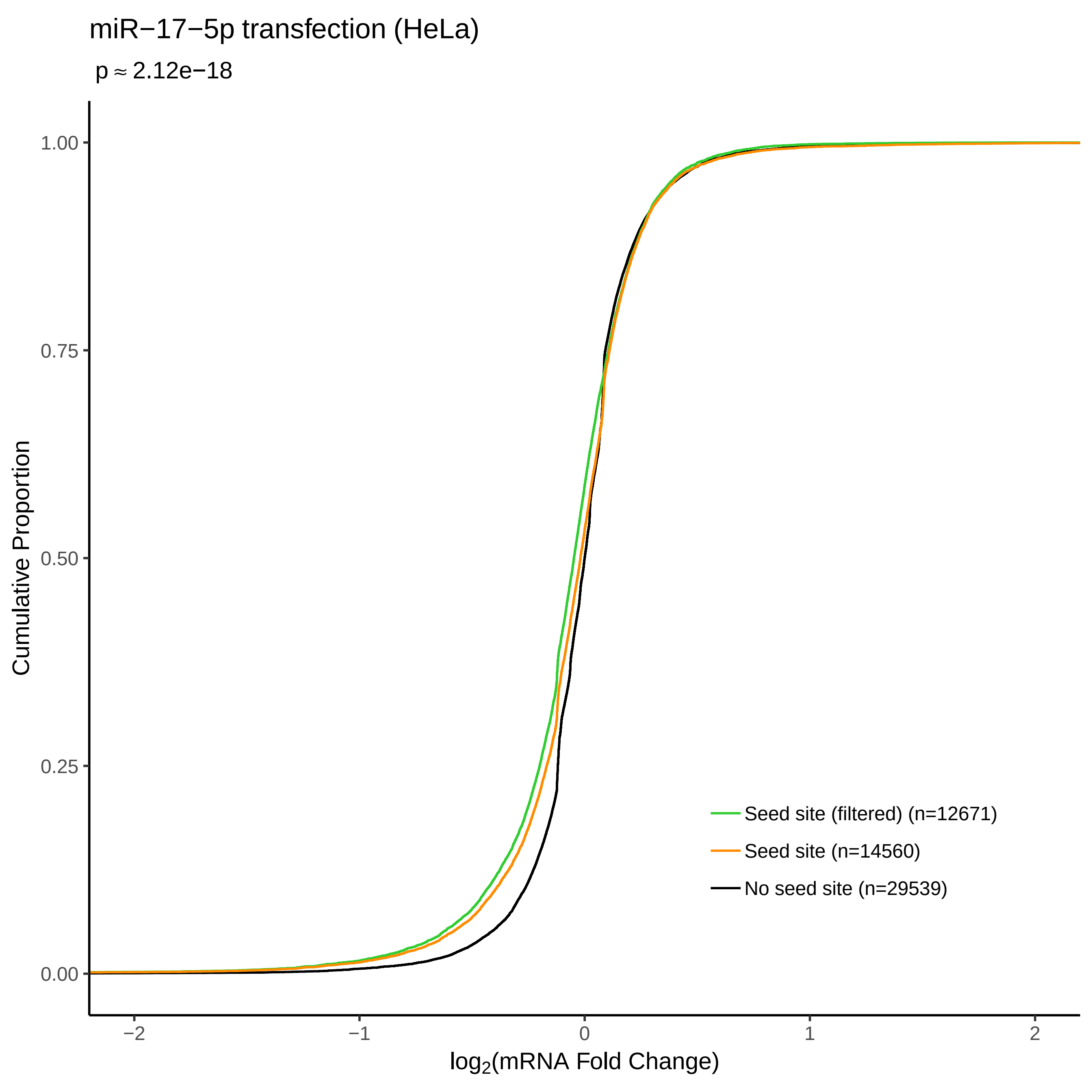

miR−17−5p transfection (HeLa)
p ≈ 2.12e−18
Cumulative Proportion
1.00
0.75
0.50
0.25
0.00
−2
−1
0
1
2
log2(mRNA Fold Change)
Seed site (filtered) (n=12671)
Seed site (n=14560)
No seed site (n=29539)

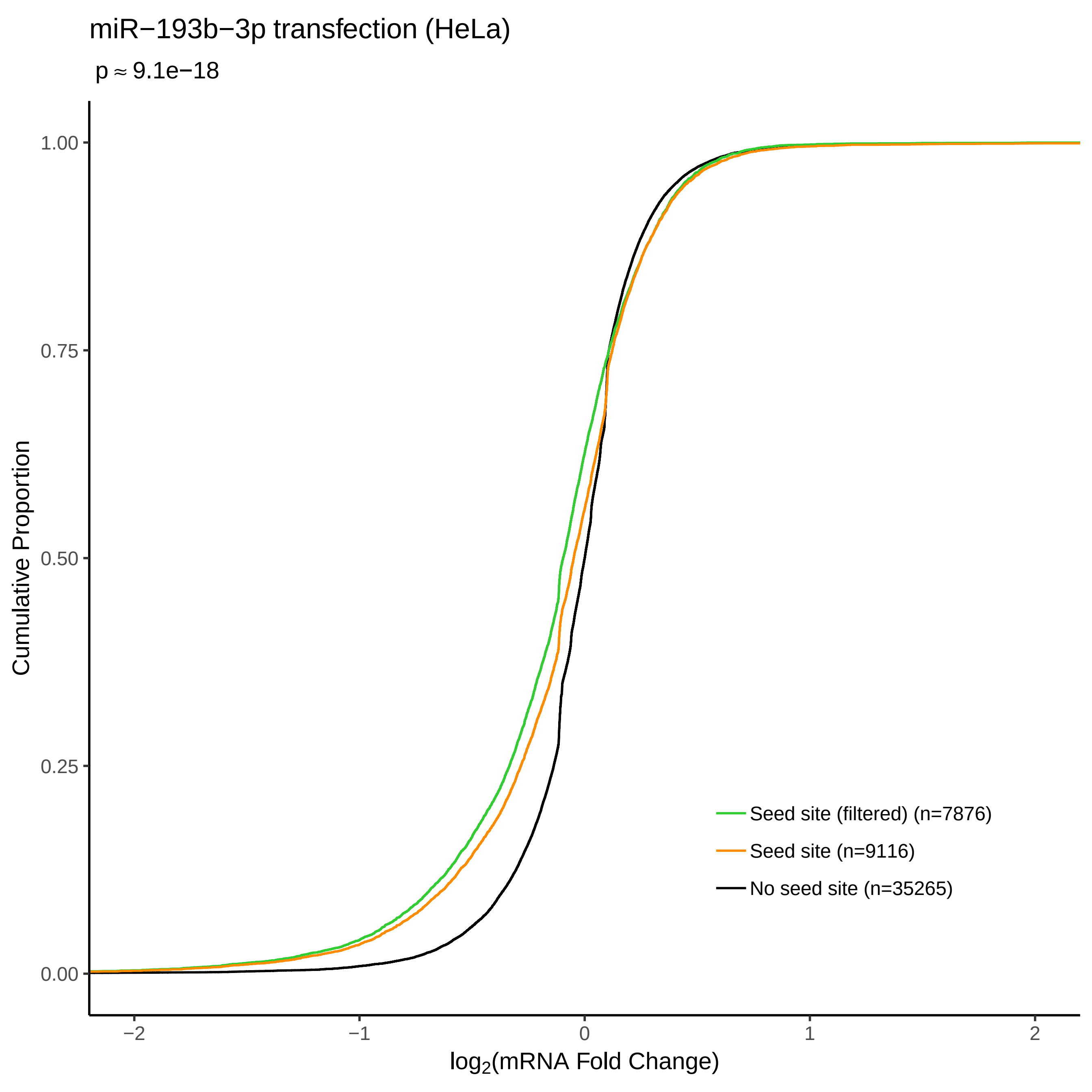

miR−193b−3p transfection (HeLa)
p ≈ 9.1e−18
Cumulative Proportion
1.00
0.75
0.50
0.25
0.00
−2
−1
0
1
2
log2(mRNA Fold Change)
Seed site (filtered) (n=7876)
Seed site (n=9116)
No seed site (n=35265)

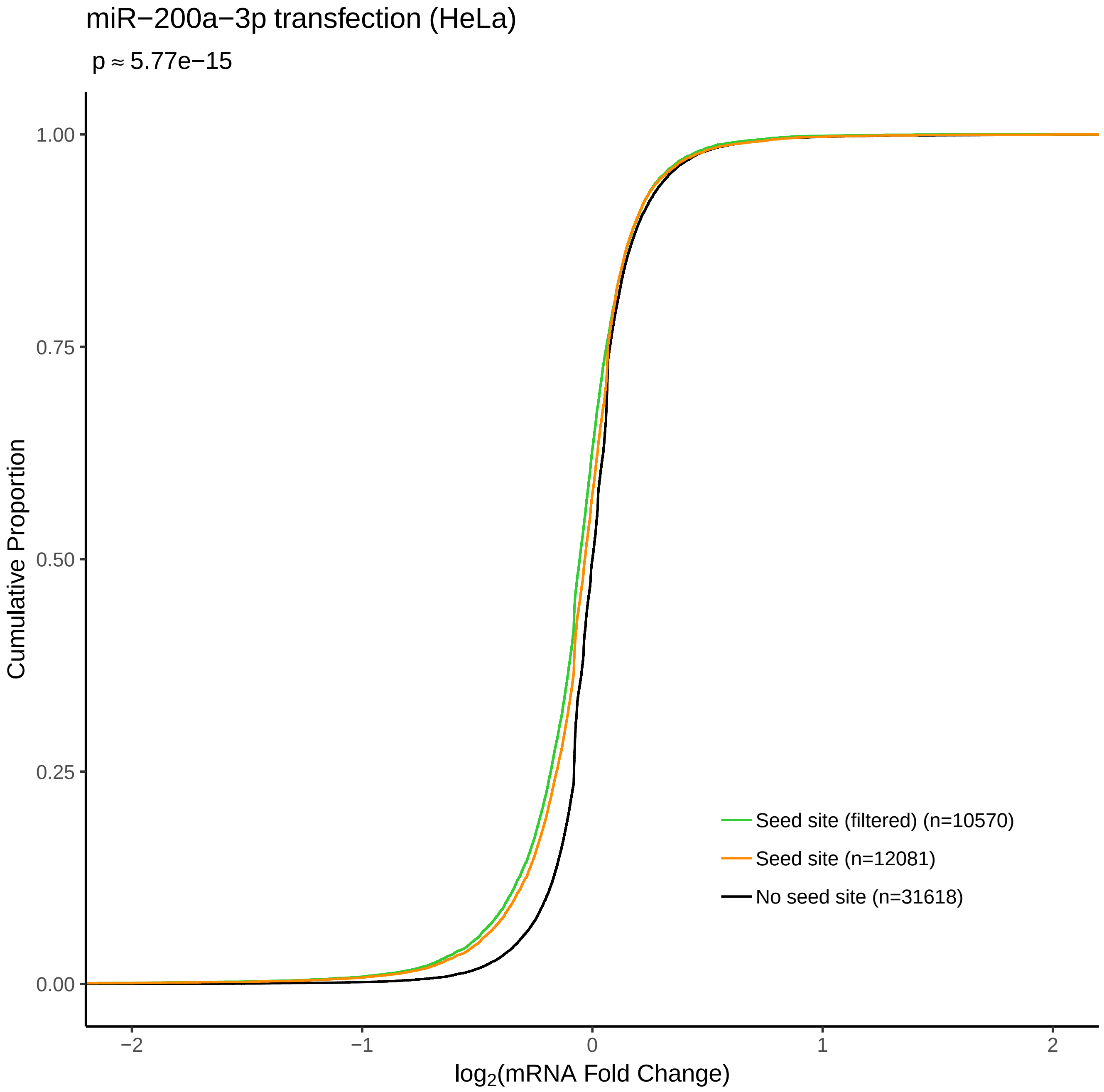

miR−200a−3p transfection (HeLa)
p ≈ 5.77e−15
Cumulative Proportion
1.00
0.75
0.50
0.25
0.00
Seed site (filtered) (n=10570)
Seed site (n=12081)
No seed site (n=31618)
−2
−1
0
1
2
log2(mRNA Fold Change)

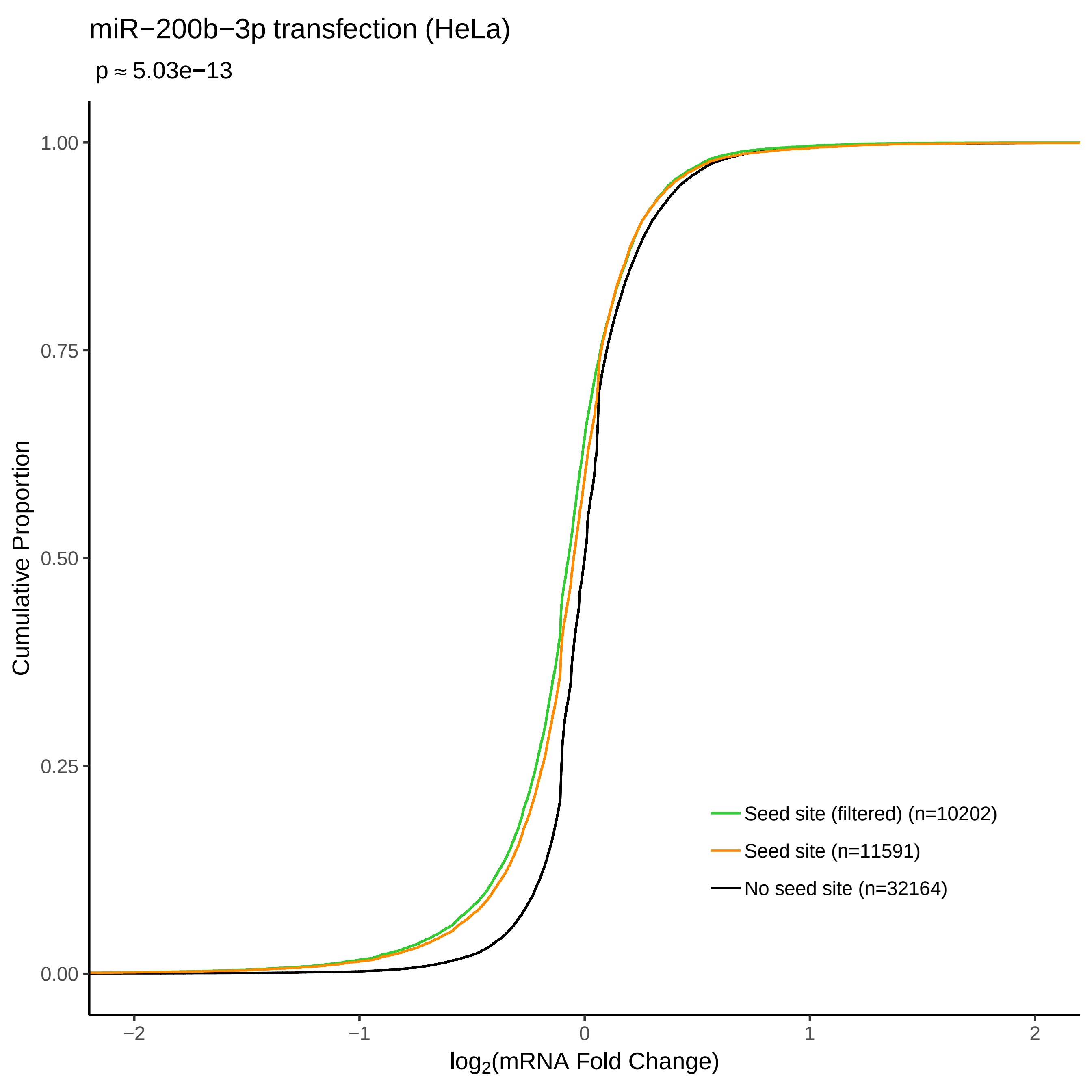

miR−200b−3p transfection (HeLa)
p ≈ 5.03e−13
Cumulative Proportion
1.00
0.75
0.50
0.25
0.00
−2
−1
0
1
2
log2(mRNA Fold Change)
Seed site (filtered) (n=10202)
Seed site (n=11591)
No seed site (n=32164)

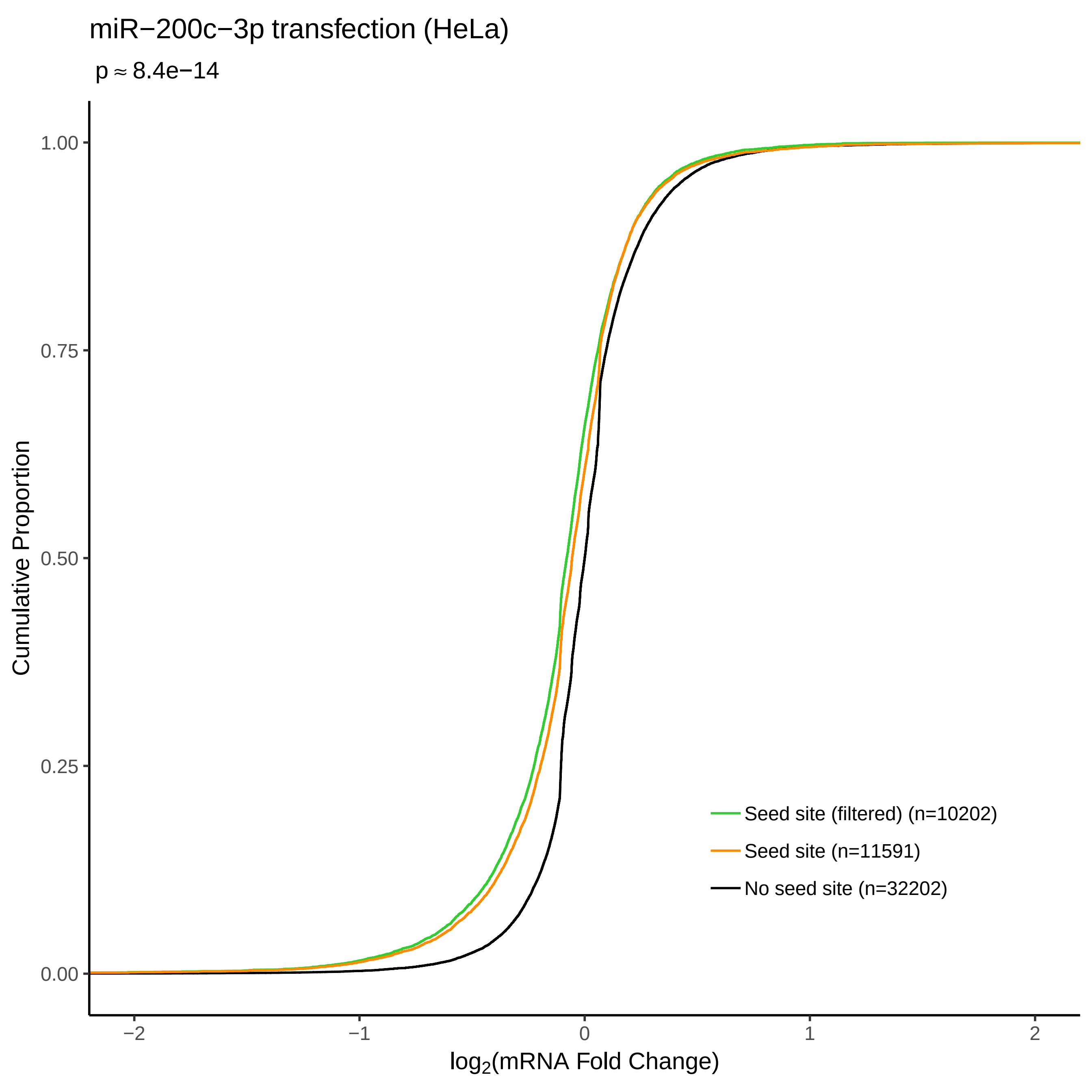

miR−200c−3p transfection (HeLa)
p ≈ 8.4e−14
Cumulative Proportion
1.00
0.75
0.50
0.25
0.00
−2
−1
0
1
2
log2(mRNA Fold Change)
Seed site (filtered) (n=10202)
Seed site (n=11591)
No seed site (n=32202)

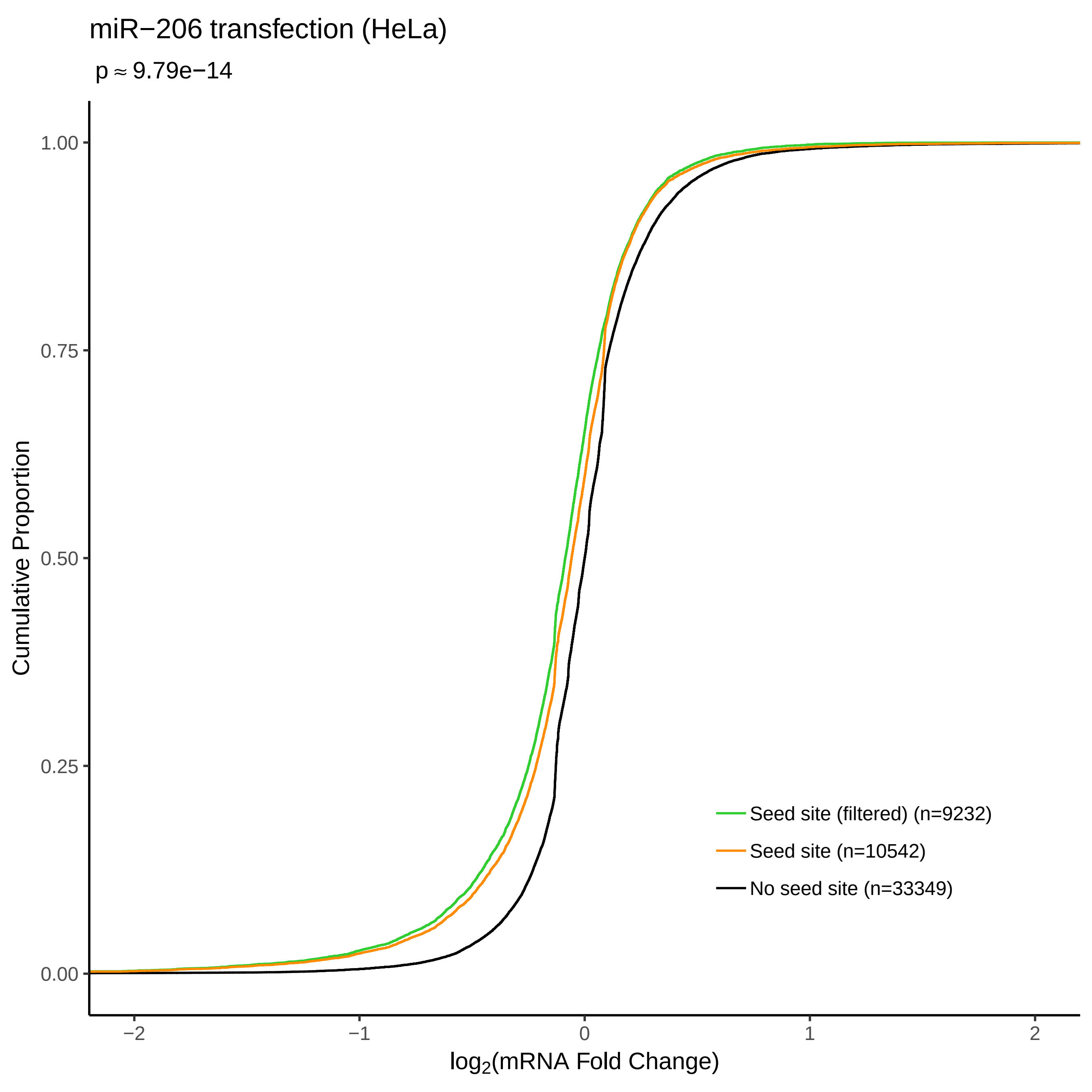

miR−206 transfection (HeLa)
p ≈ 9.79e−14
Cumulative Proportion
1.00
0.75
0.50
0.25
0.00
−2
−1
0
1
2
log2(mRNA Fold Change)
Seed site (filtered) (n=9232)
Seed site (n=10542)
No seed site (n=33349)

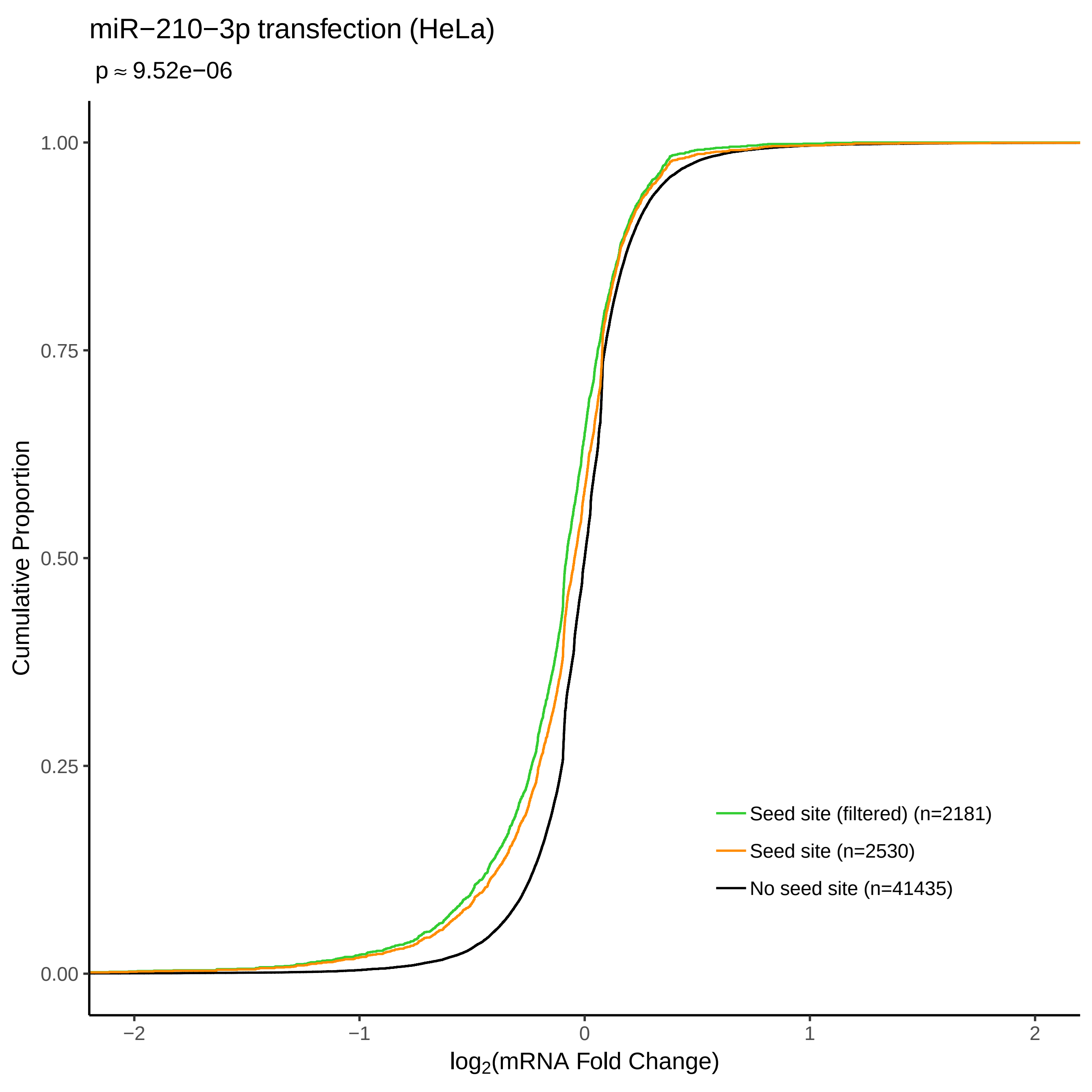

miR−210−3p transfection (HeLa)
p ≈ 9.52e−06
Cumulative Proportion
1.00
0.75
0.50
0.25
0.00
−2
−1
0
1
2
log2(mRNA Fold Change)
Seed site (filtered) (n=2181)
Seed site (n=2530)
No seed site (n=41435)

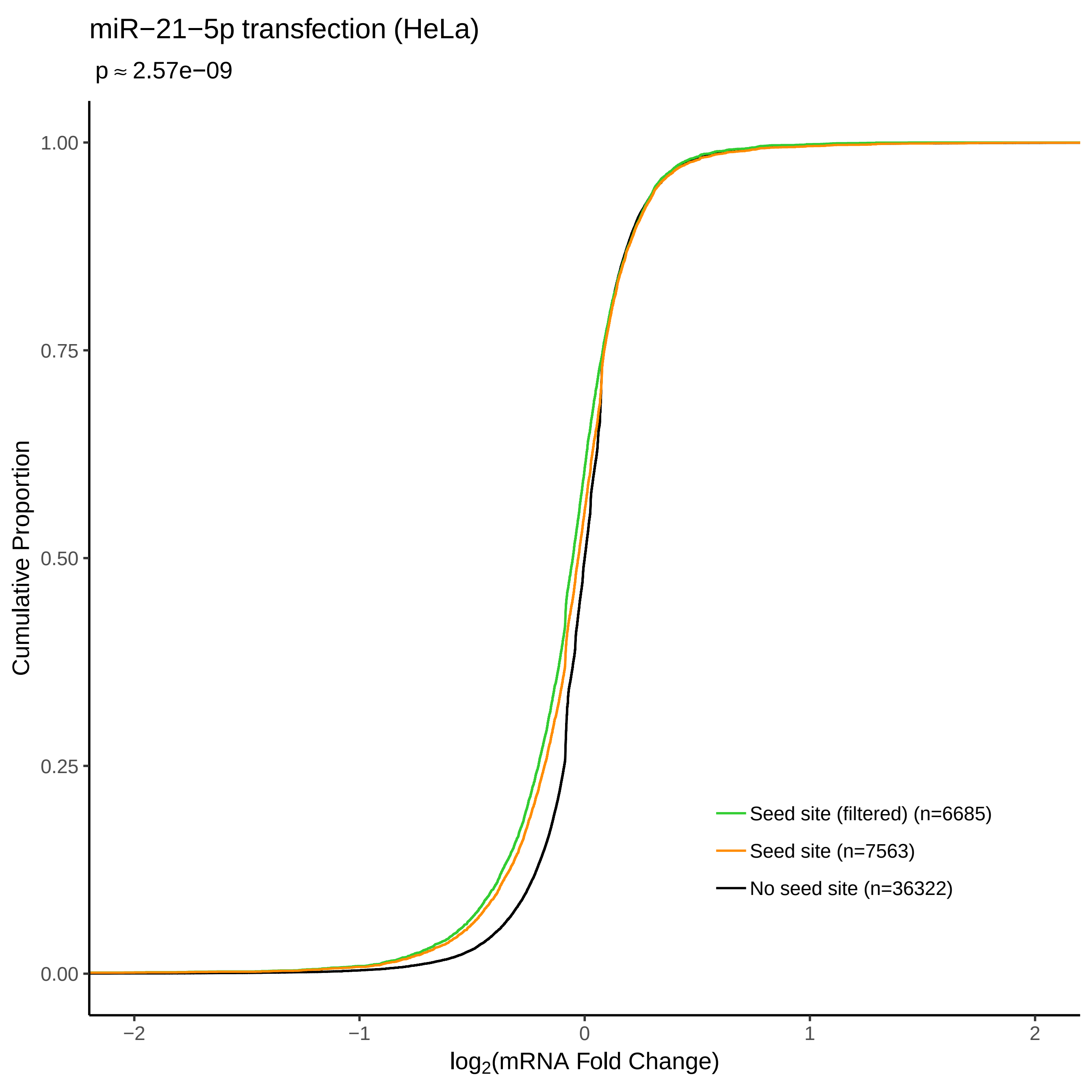
miR−21−5p transfection (HeLa)
p ≈ 2.57e−09
Cumulative Proportion
1.00
0.75
0.50
0.25
0.00
−2
−1
0
1
2
log2(mRNA Fold Change)
Seed site (filtered) (n=6685)
Seed site (n=7563)
No seed site (n=36322)

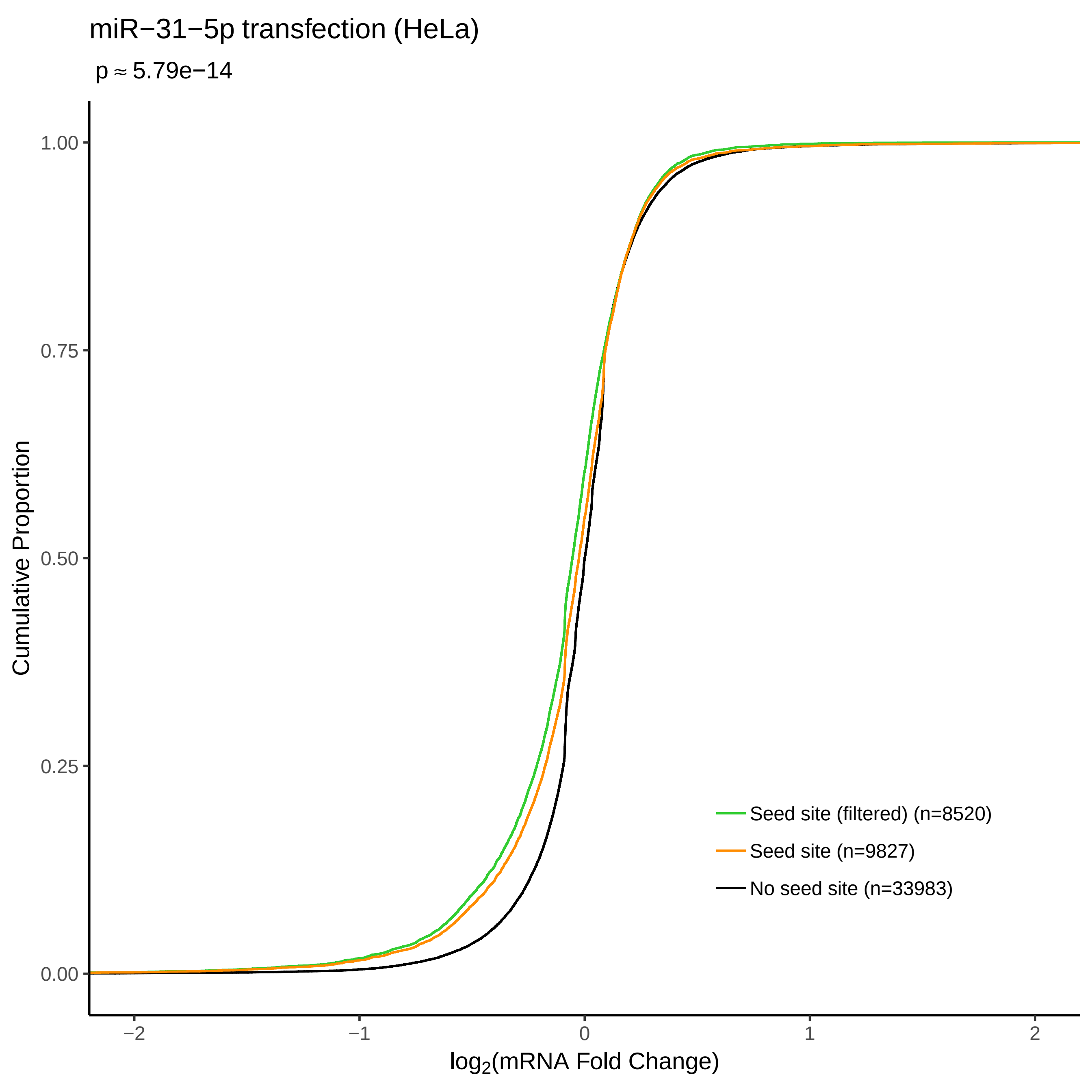

miR−31−5p transfection (HeLa)
p ≈ 5.79e−14
Cumulative Proportion
1.00
0.75
0.50
0.25
0.00
−2
−1
0
1
2
log2(mRNA Fold Change)
Seed site (filtered) (n=8520)
Seed site (n=9827)
No seed site (n=33983)

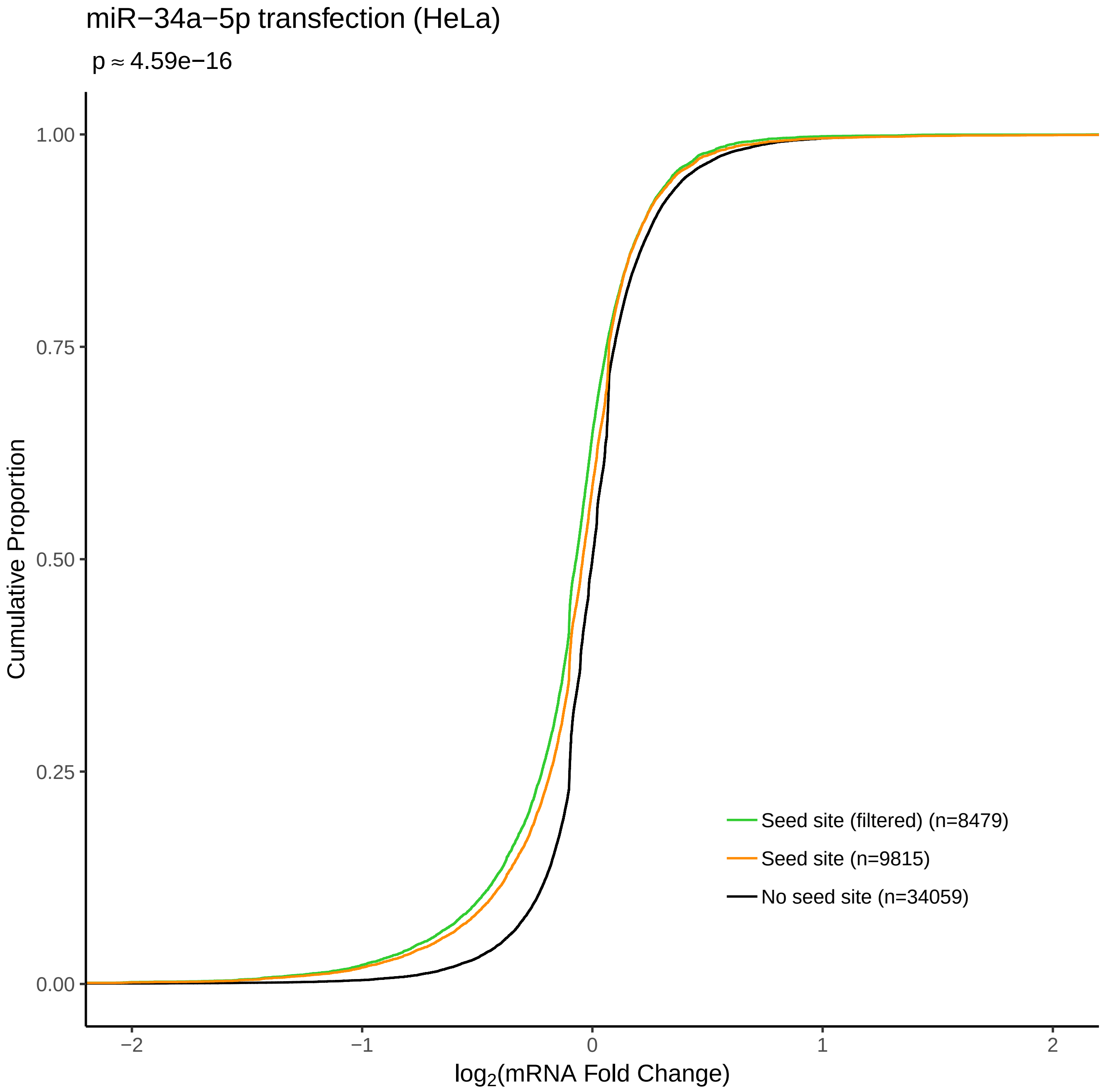

miR−34a−5p transfection (HeLa)
p ≈ 4.59e−16
Cumulative Proportion
1.00
0.75
0.50
0.25
0.00
−2
−1
0
1
2
log2(mRNA Fold Change)
Seed site (filtered) (n=8479)
Seed site (n=9815)
No seed site (n=34059)

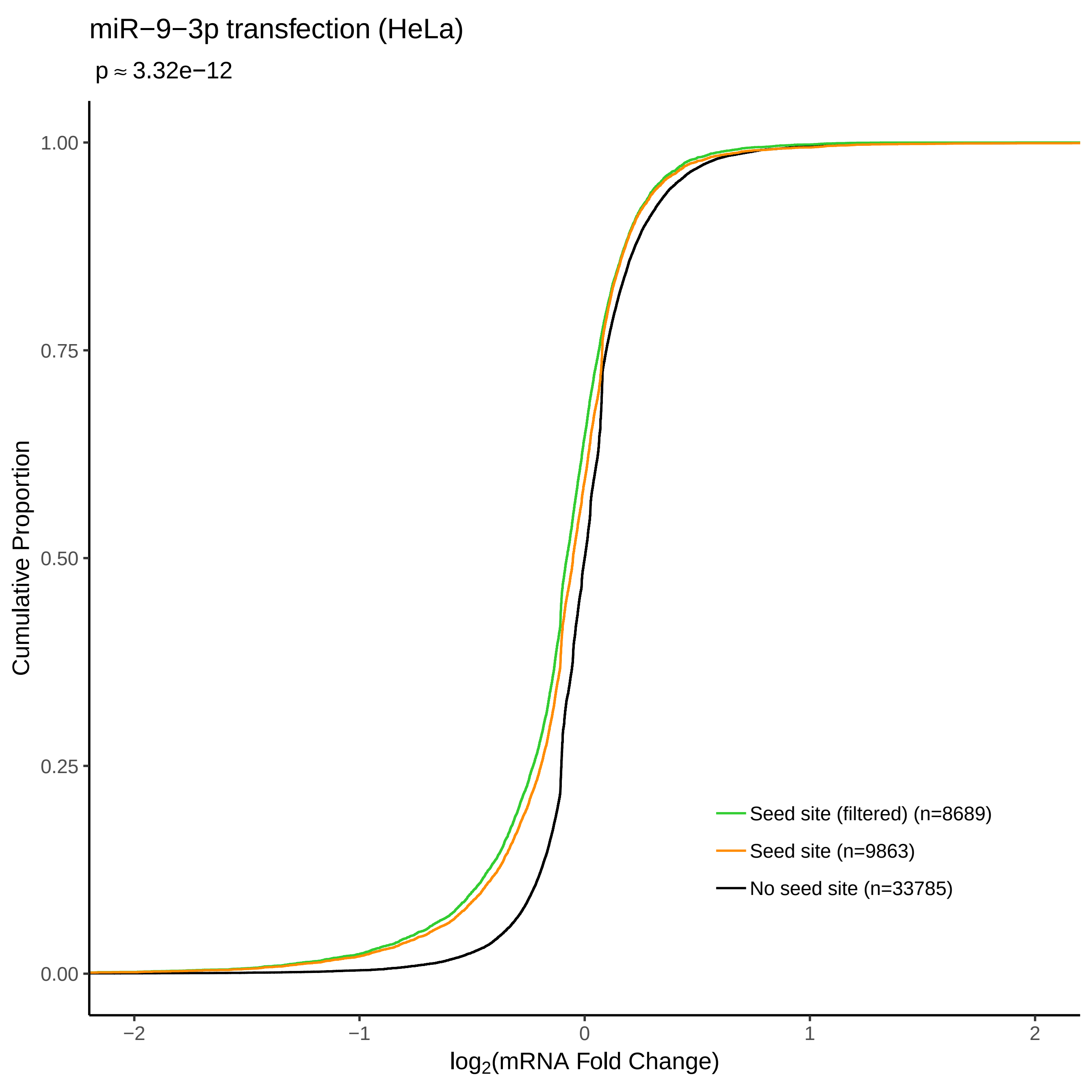
miR−9−3p transfection (HeLa)
p ≈ 3.32e−12
Cumulative Proportion
1.00
0.75
0.50
0.25
0.00
−2
−1
0
1
2
log2(mRNA Fold Change)
Seed site (filtered) (n=8689)
Seed site (n=9863)
No seed site (n=33785)

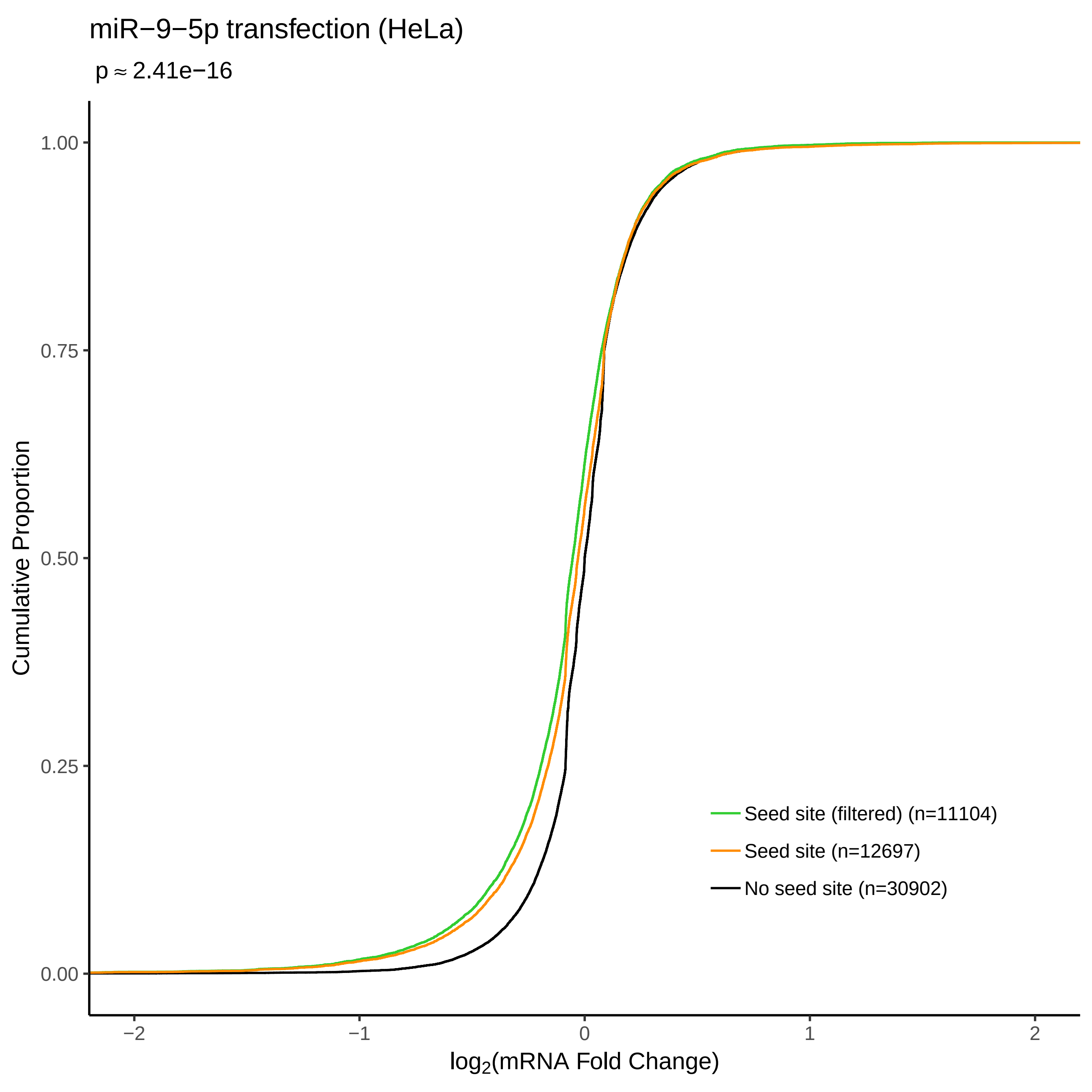

miR−9−5p transfection (HeLa)
p ≈ 2.41e−16
Cumulative Proportion
1.00
0.75
0.50
0.25
0.00
−2
−1
0
1
2
log2(mRNA Fold Change)
Seed site (filtered) (n=11104)
Seed site (n=12697)
No seed site (n=30902)
