## Supplementary figures and images for "FilTar: Using RNA-Seq data to improve microRNA target prediction accuracy in animals"

### Supplementary File 4

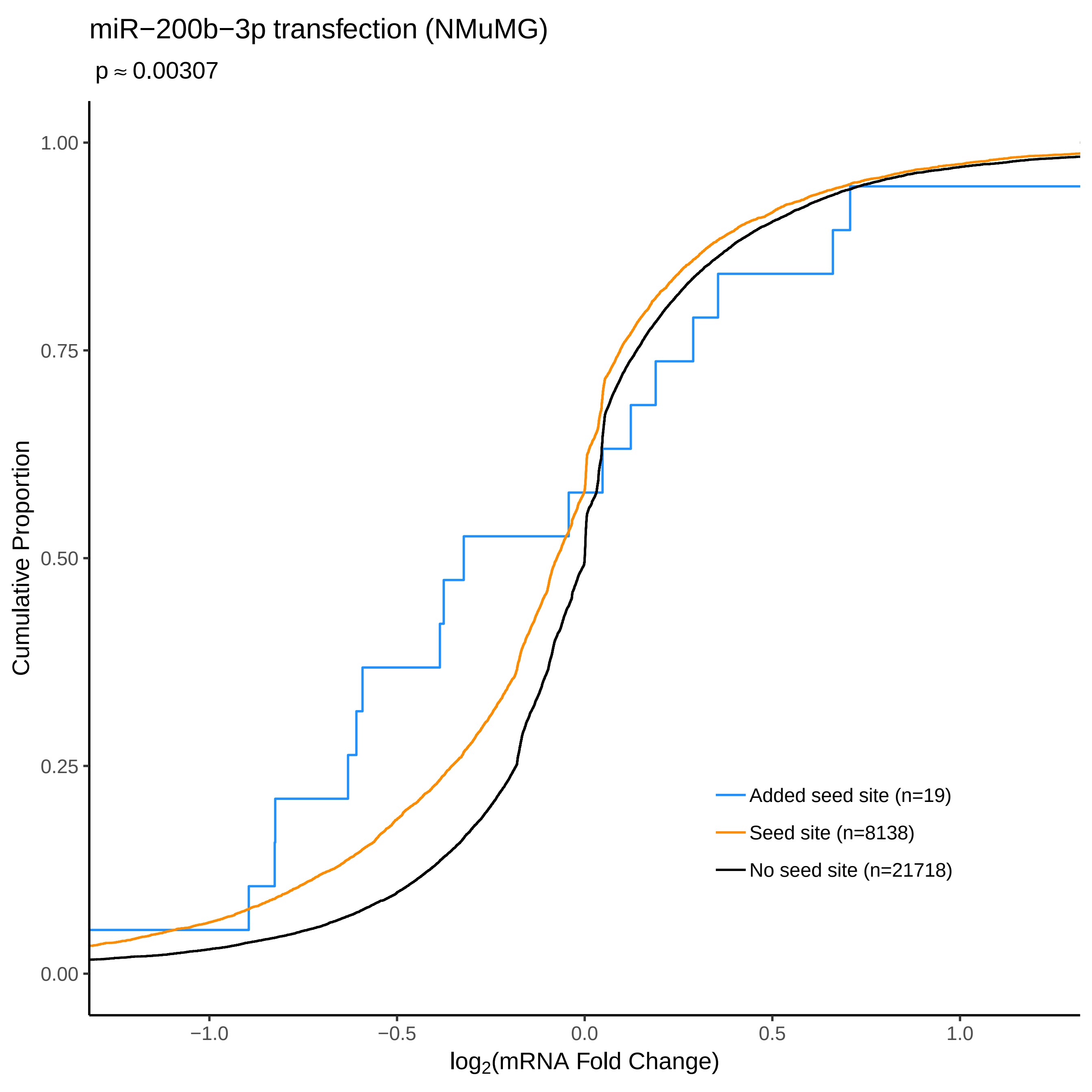

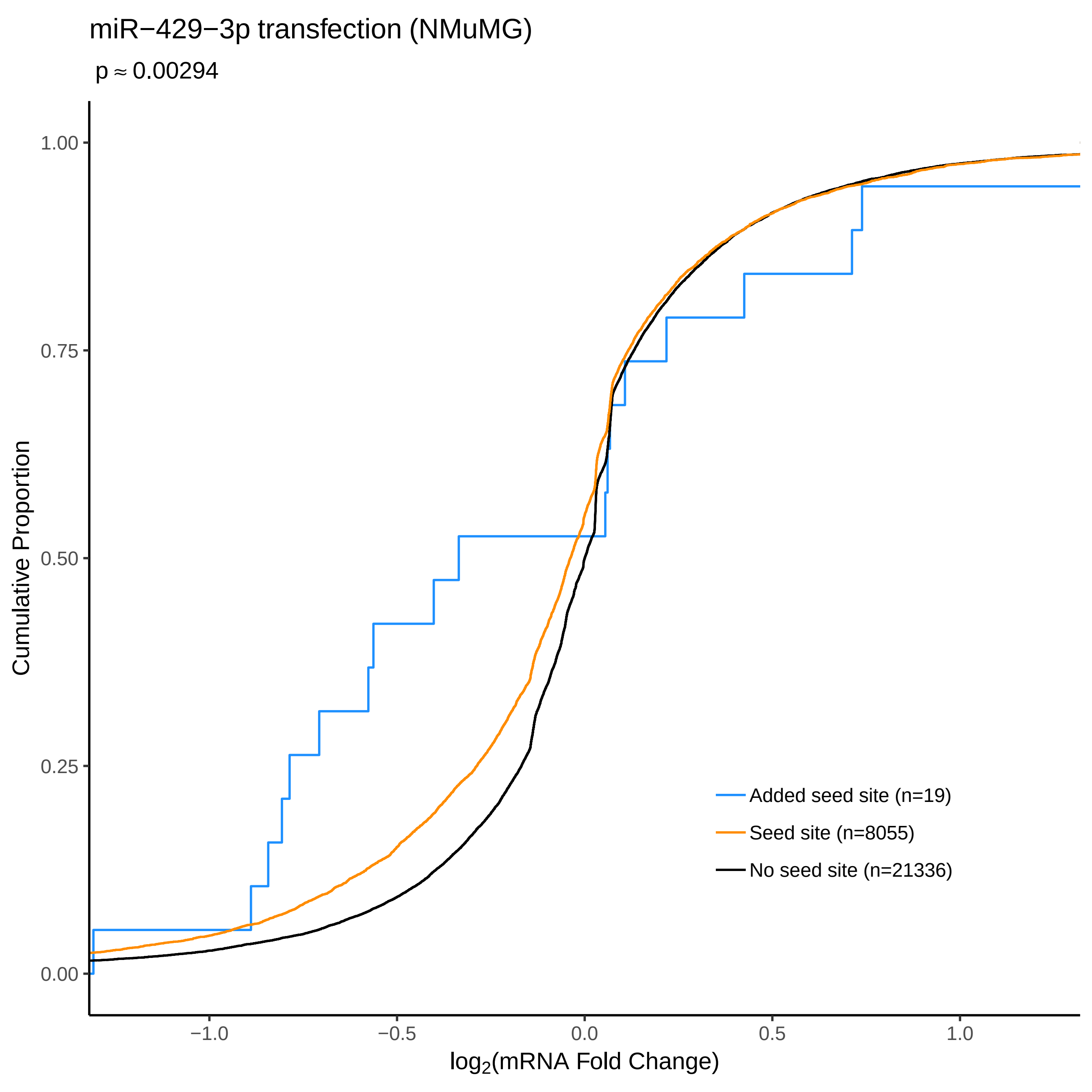

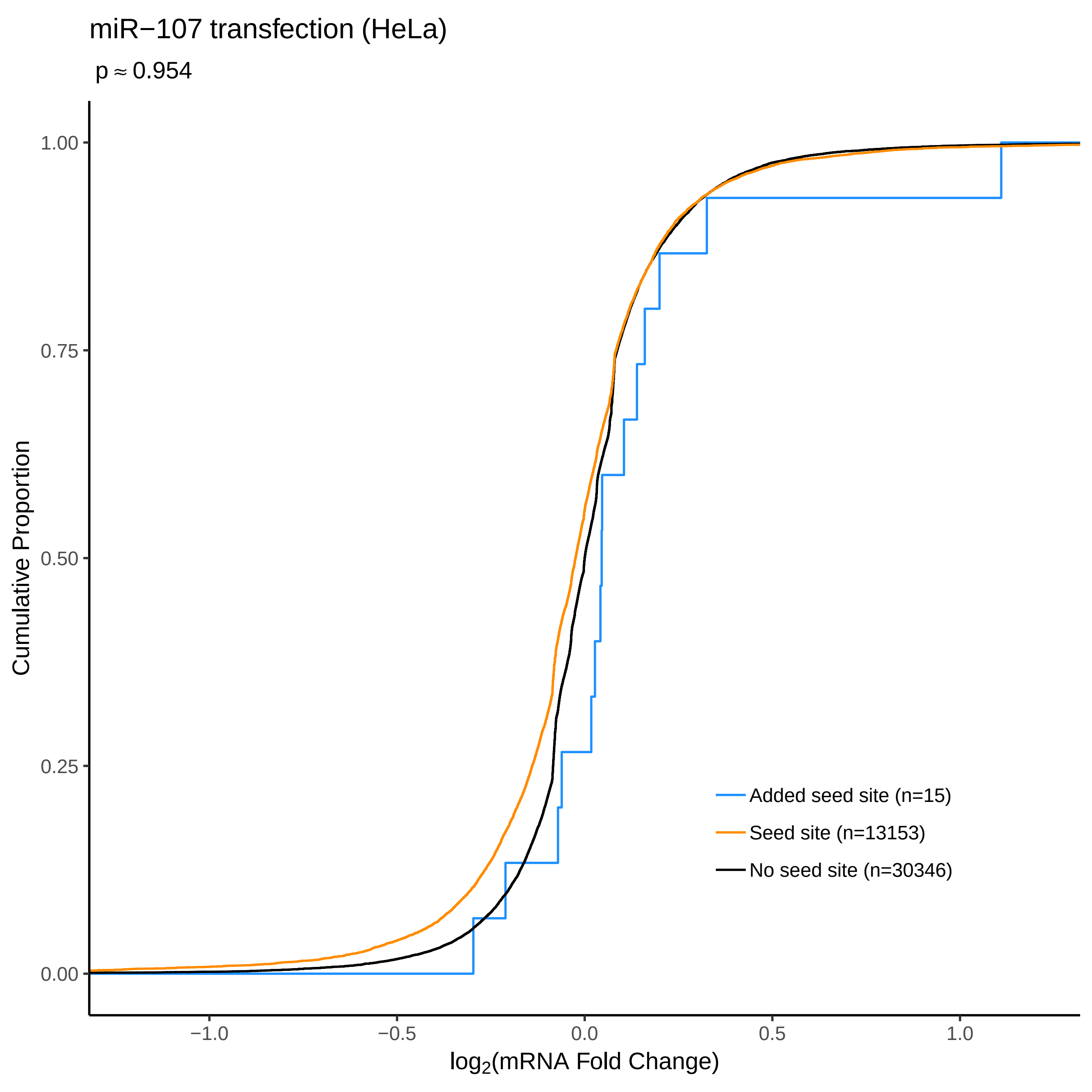

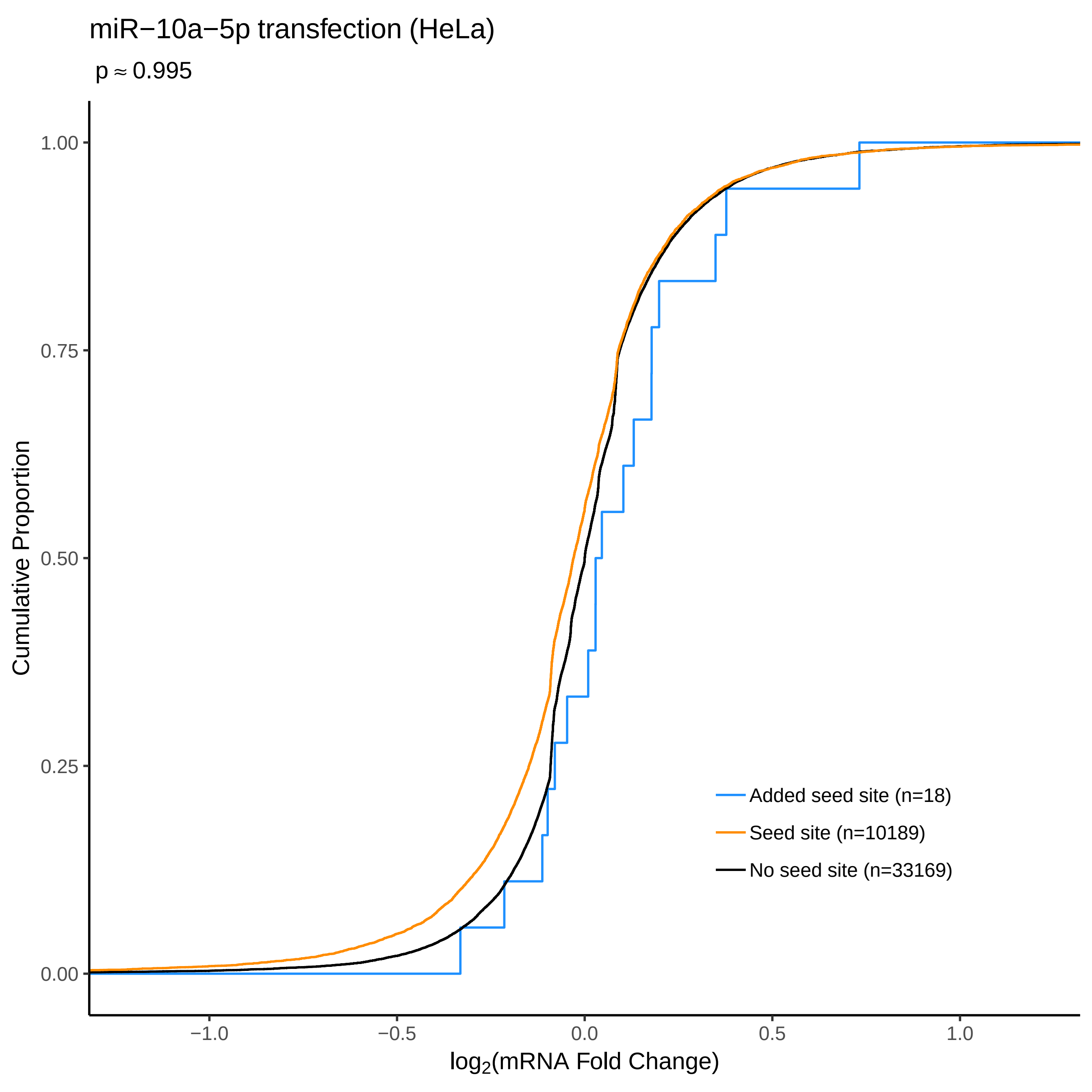

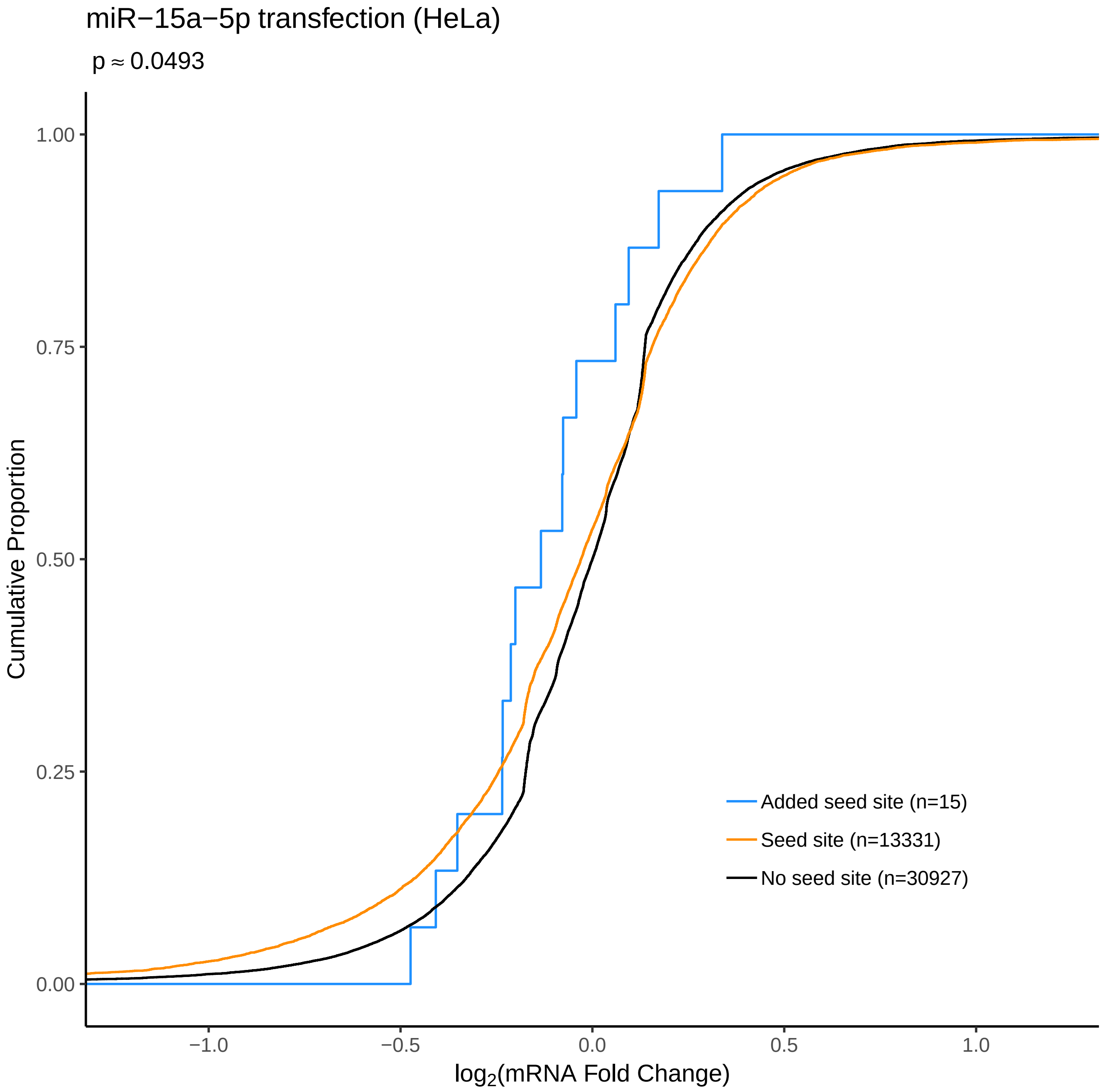
