## Supplementary File 5 for "FilTar: Using RNA-Seq data to improve microRNA target prediction accuracy in animals"

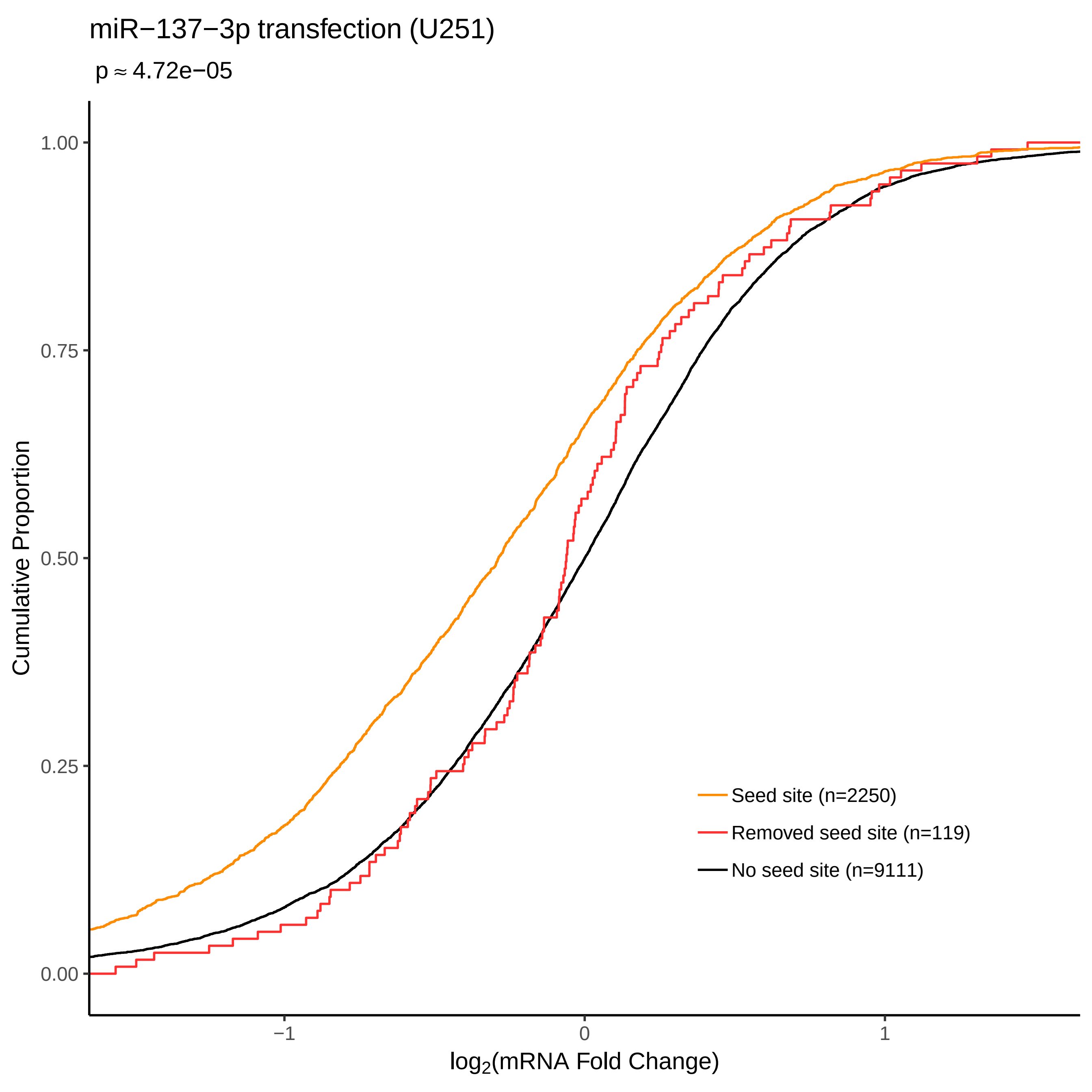
miR−137−3p transfection (U251)
p ≈ 4.72e−05
Cumulative Proportion
1.00
0.75
0.50
0.25
0.00
−1
0
1
log2(mRNA Fold Change)
Seed site (n=2250)
Removed seed site (n=119)
No seed site (n=9111)

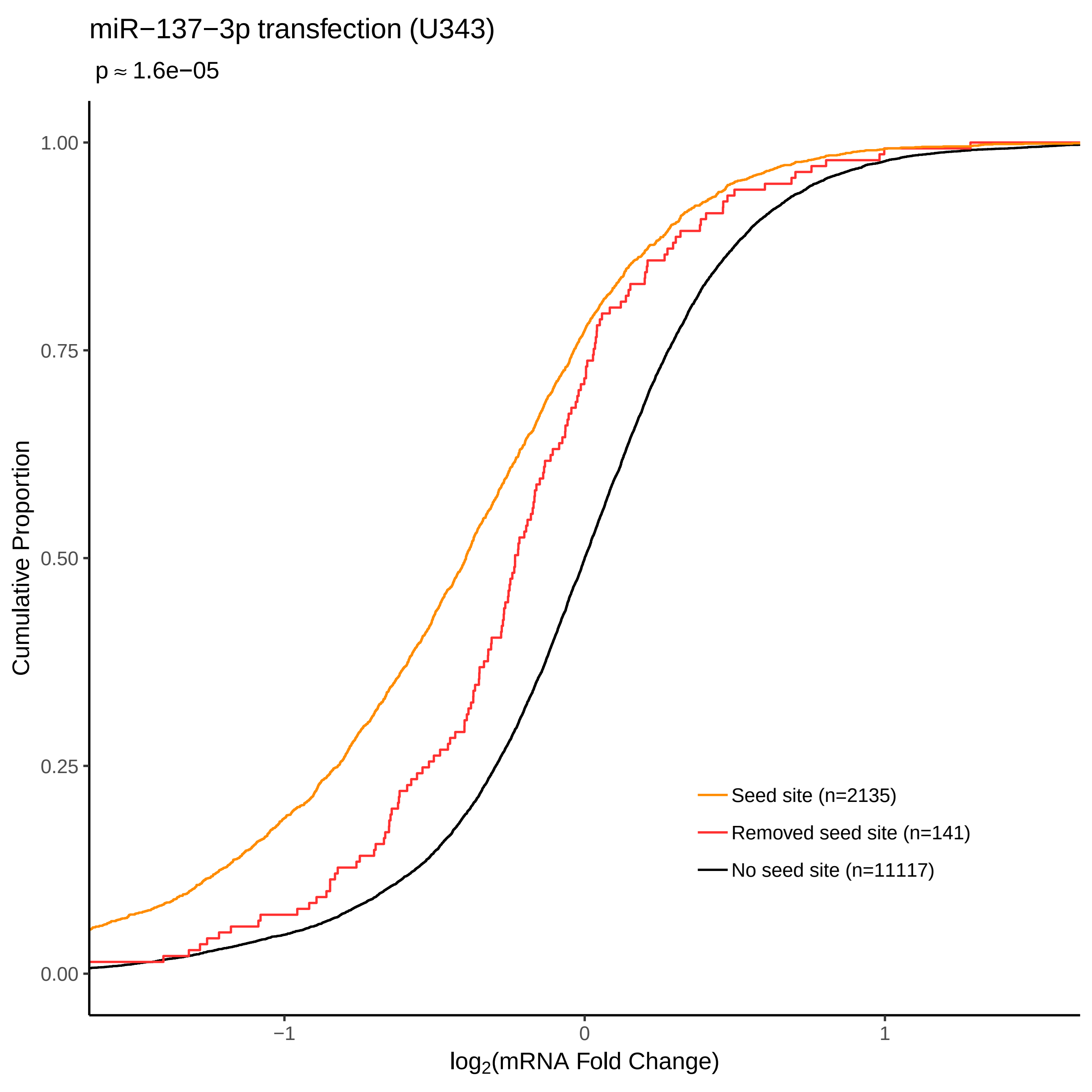
miR−137−3p transfection (U343)
p ≈ 1.6e−05
Cumulative Proportion
1.00
0.75
0.50
0.25
0.00
−1
0
1
log2(mRNA Fold Change)
Seed site (n=2135)
Removed seed site (n=141)
No seed site (n=11117)

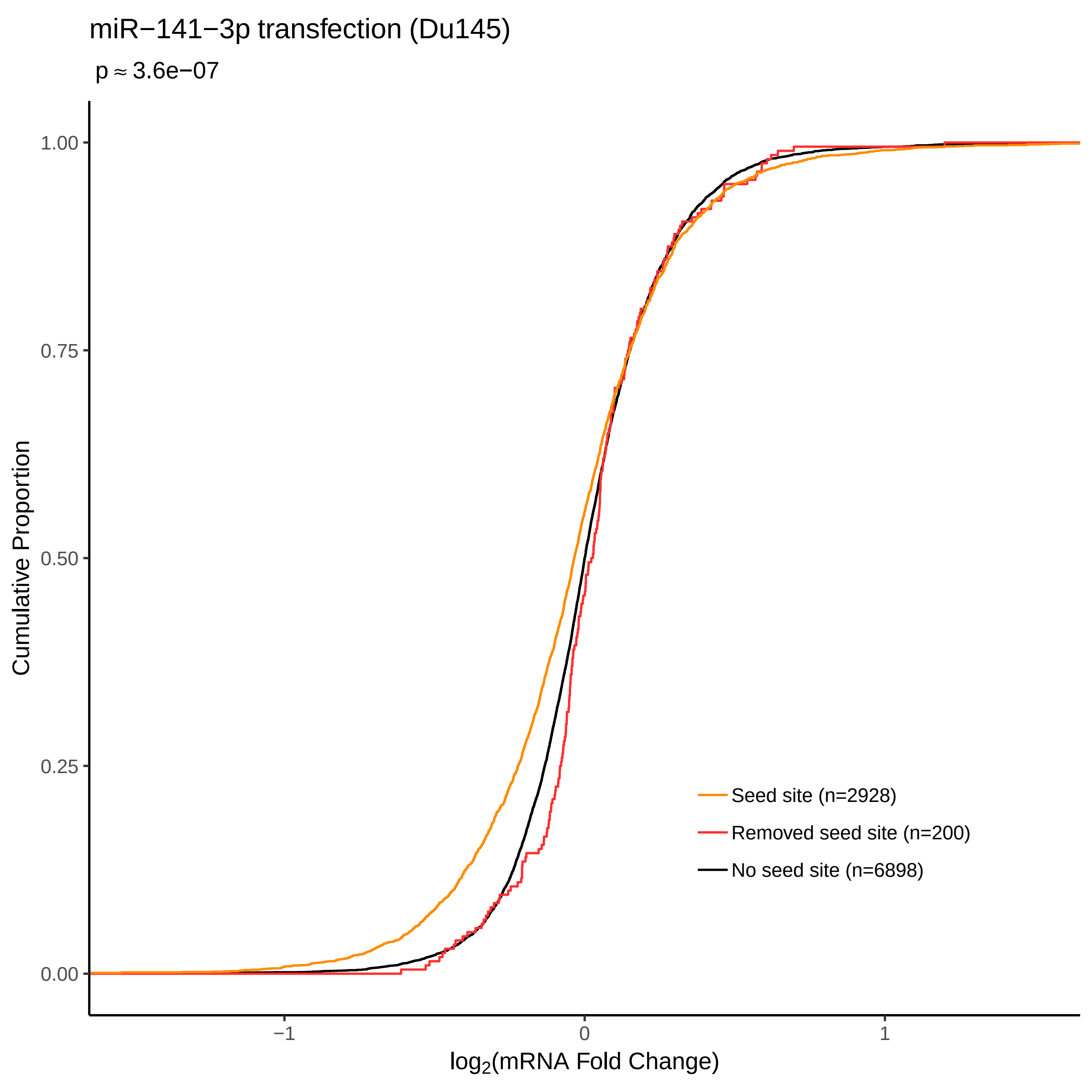

miR−141−3p transfection (Du145)
p ≈ 3.6e−07
Cumulative Proportion
1.00
0.75
0.50
0.25
0.00
−1
0
1
log2(mRNA Fold Change)
Seed site (n=2928)
Removed seed site (n=200)
No seed site (n=6898)

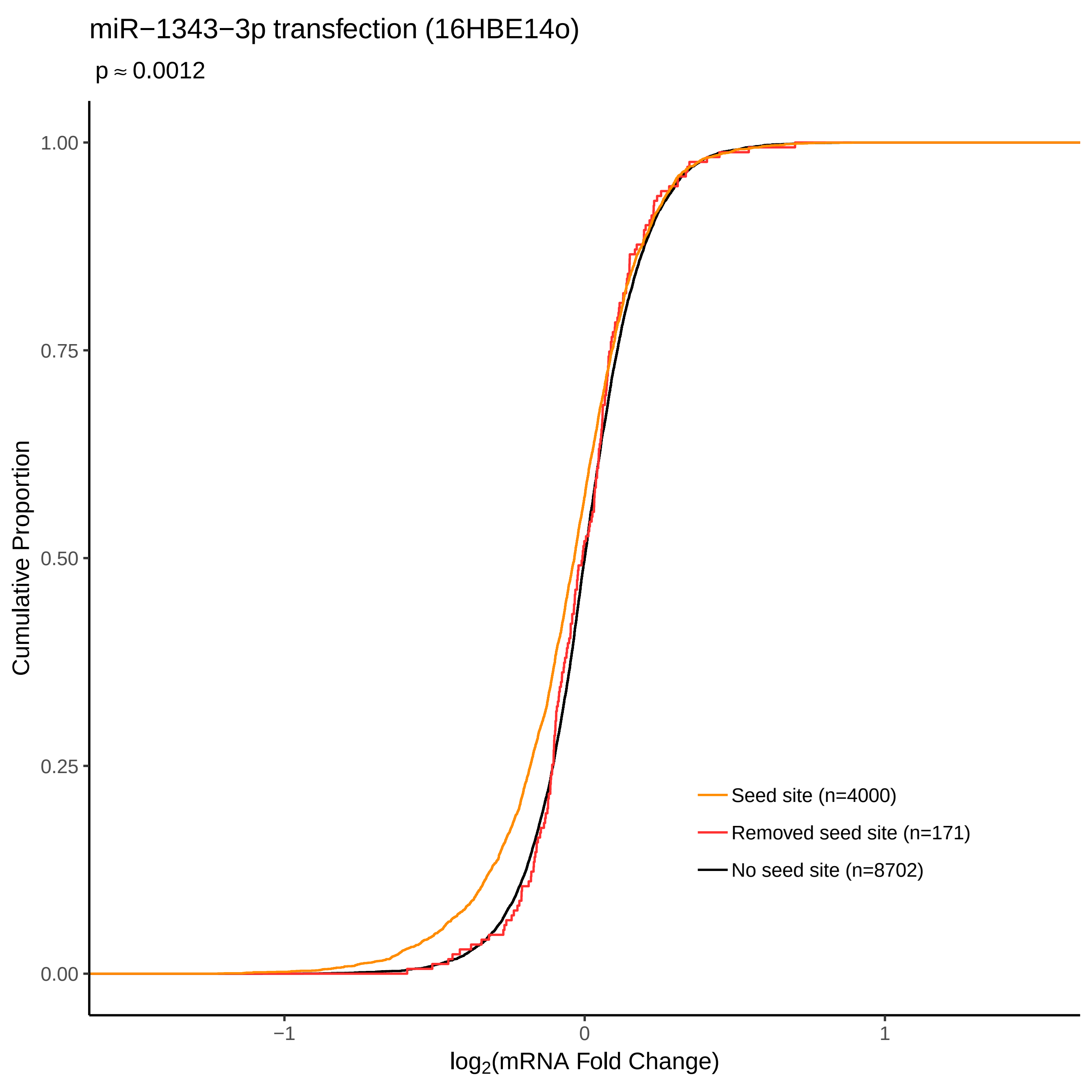

miR−1343−3p transfection (16HBE14o)
p ≈ 0.0012
Cumulative Proportion
1.00
0.75
0.50
0.25
0.00
−1
0
1
log2(mRNA Fold Change)
Seed site (n=4000)
Removed seed site (n=171)
No seed site (n=8702)

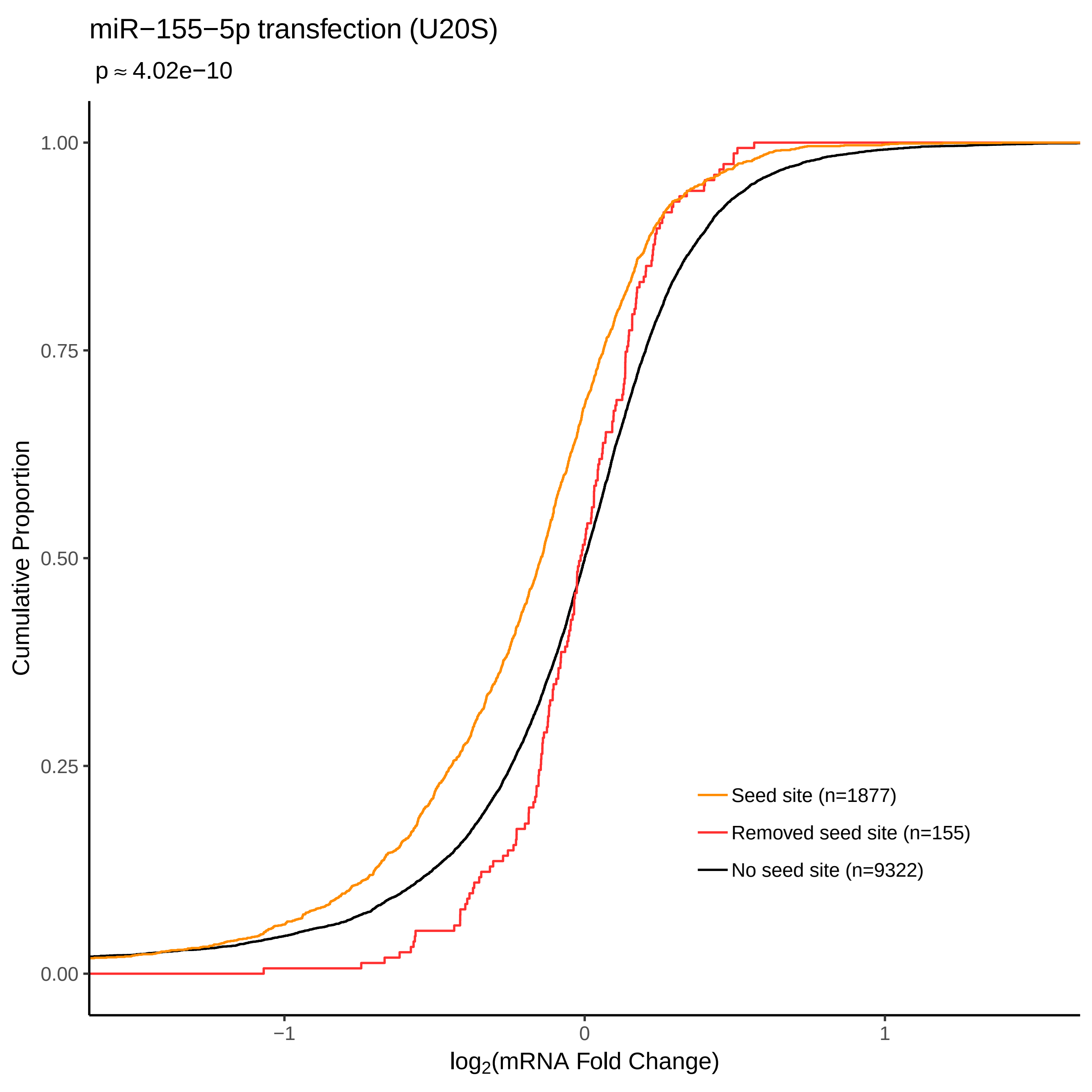
miR−155−5p transfection (U20S)
p ≈ 4.02e−10
Cumulative Proportion
1.00
0.75
0.50
0.25
0.00
−1
0
1
log2(mRNA Fold Change)
Seed site (n=1877)
Removed seed site (n=155)
No seed site (n=9322)

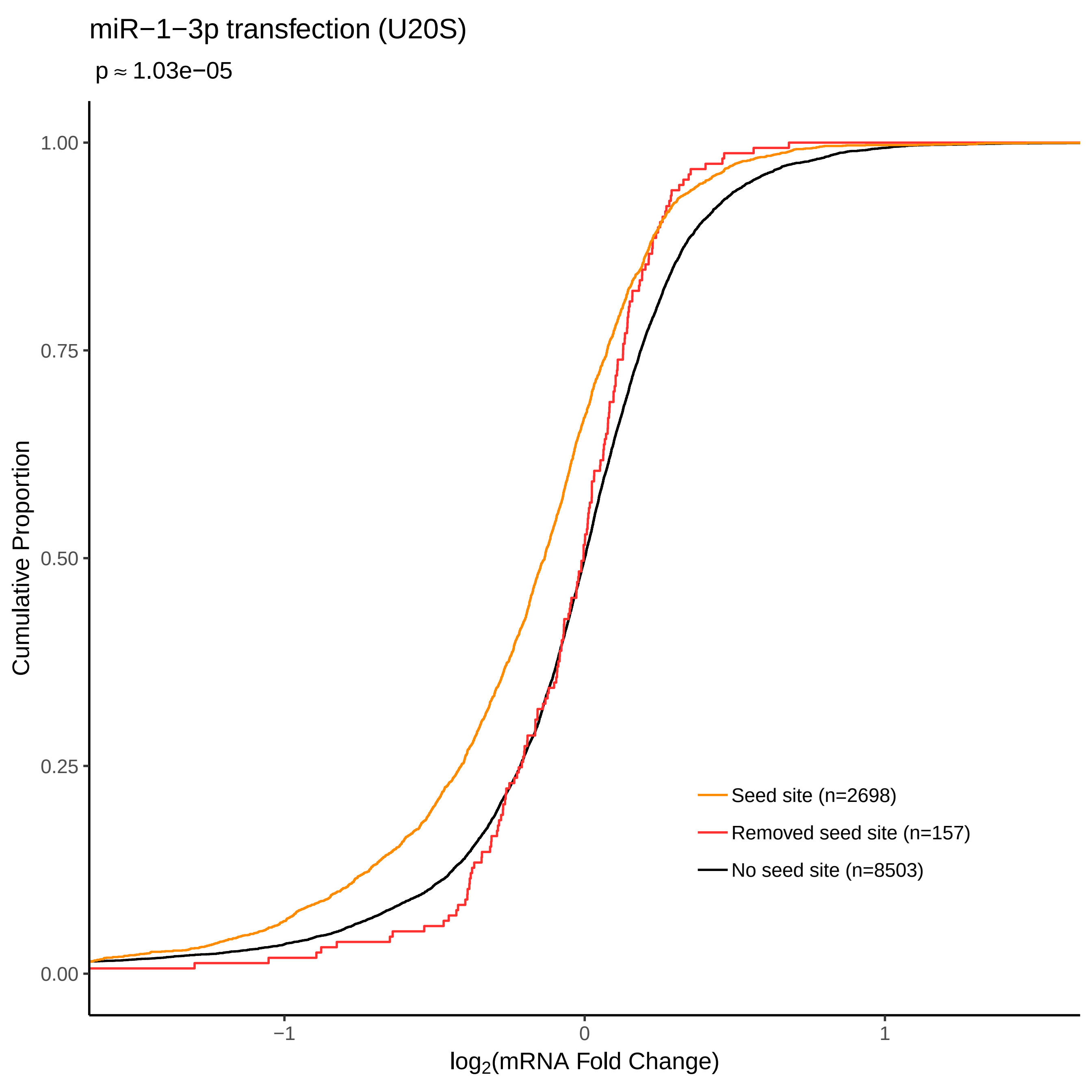
miR−1−3p transfection (U20S)
p ≈ 1.03e−05
Cumulative Proportion
1.00
0.75
0.50
0.25
0.00
−1
0
1
log2(mRNA Fold Change)
Seed site (n=2698)
Removed seed site (n=157)
No seed site (n=8503)

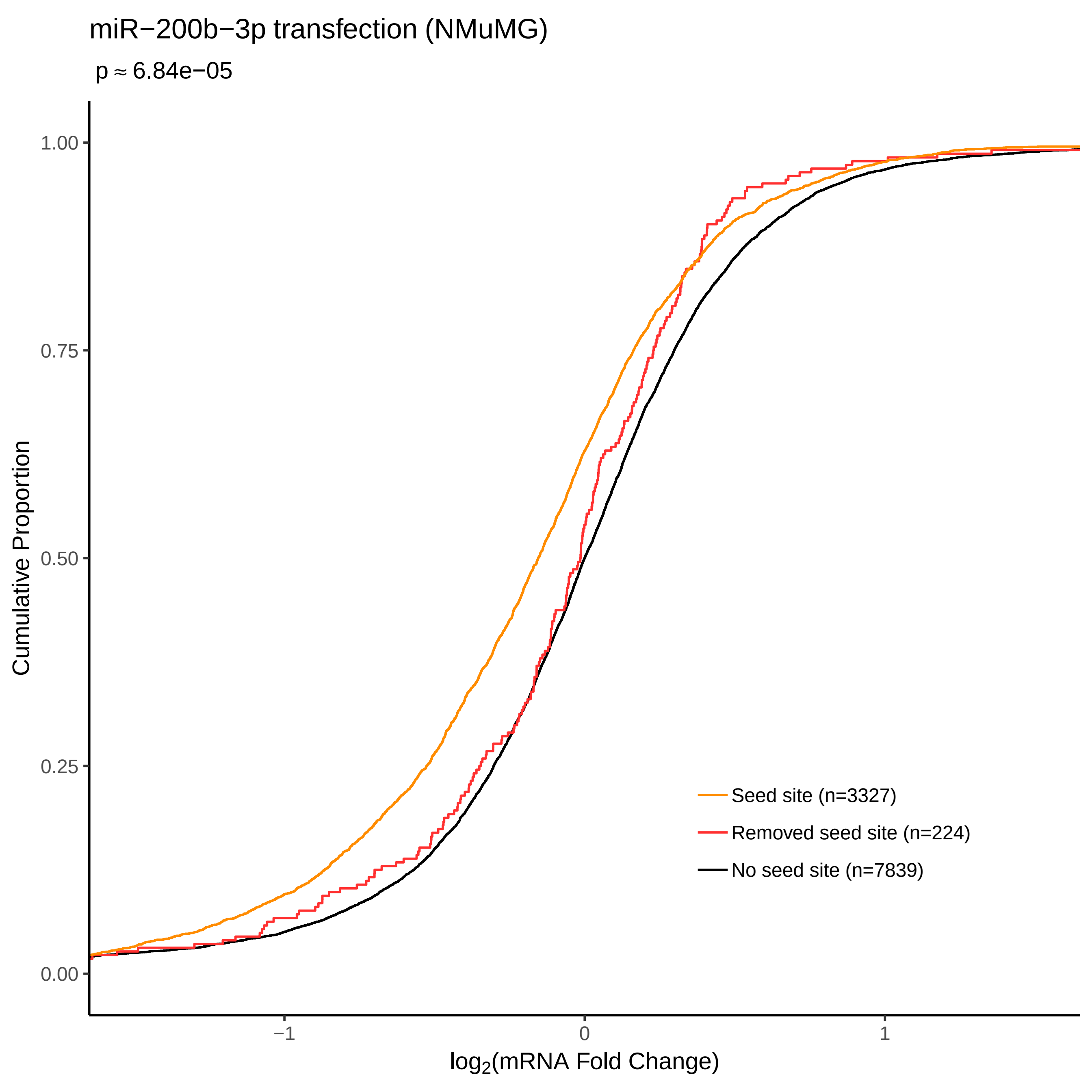

miR−200b−3p transfection (NMuMG)
p ≈ 6.84e−05
Cumulative Proportion
1.00
0.75
0.50
0.25
0.00
−1
0
1
log2(mRNA Fold Change)
Seed site (n=3327)
Removed seed site (n=224)
No seed site (n=7839)

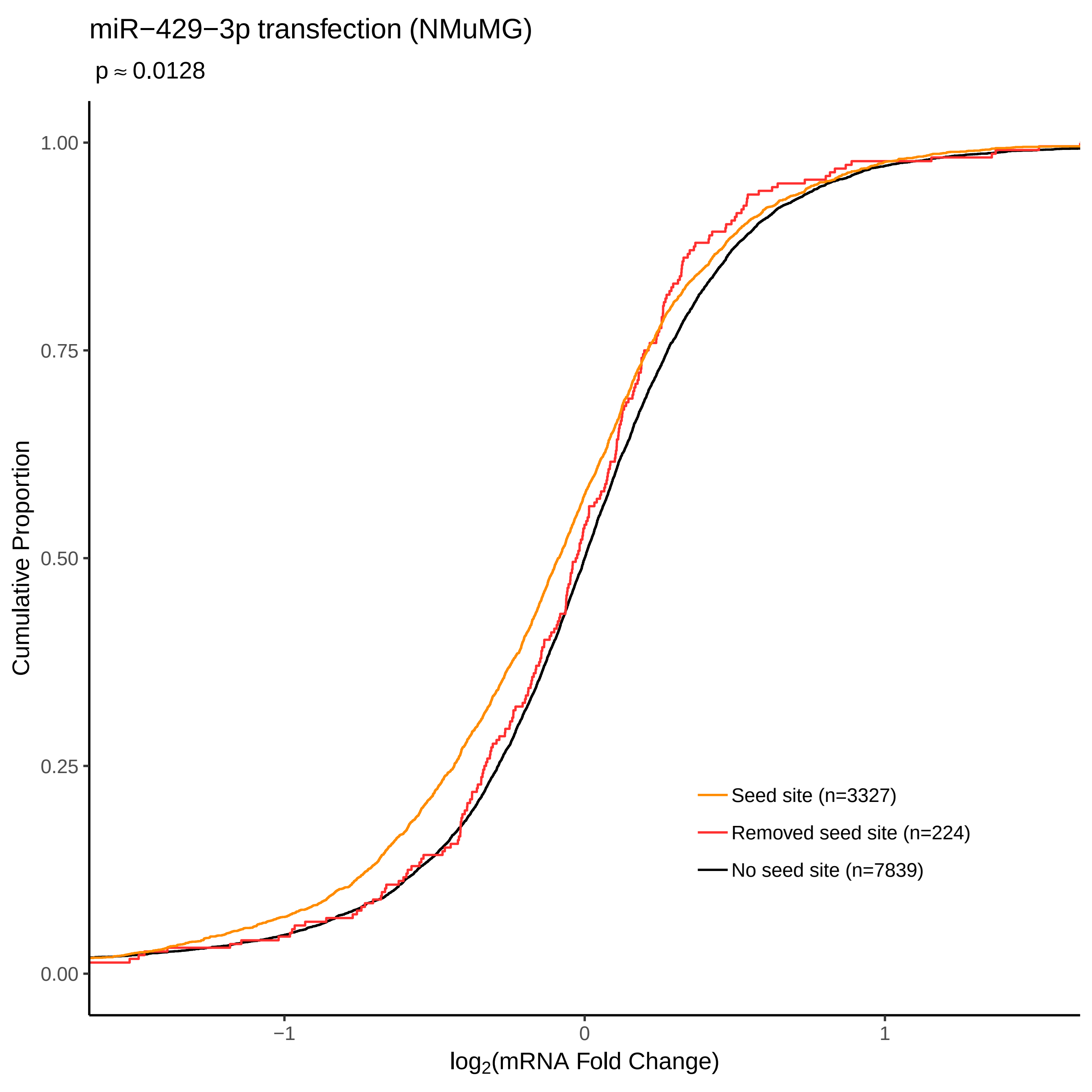
miR−429−3p transfection (NMuMG)
p ≈ 0.0128
Cumulative Proportion
1.00
0.75
0.50
0.25
0.00
−1
0
1
log2(mRNA Fold Change)
Seed site (n=3327)
Removed seed site (n=224)
No seed site (n=7839)

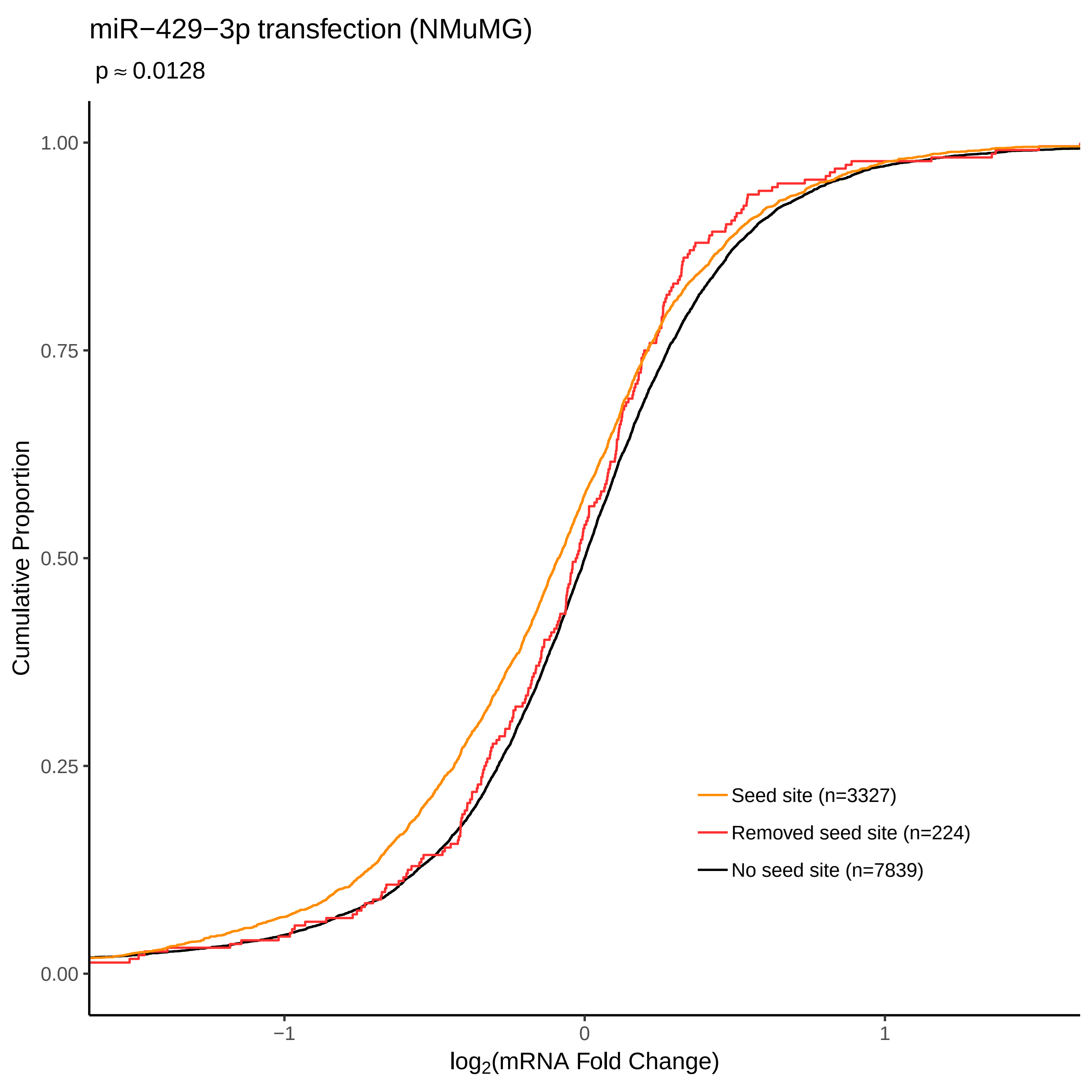
miR−429−3p transfection (NMuMG)
p ≈ 0.0128
Cumulative Proportion
1.00
0.75
0.50
0.25
0.00
−1
0
1
log2(mRNA Fold Change)
Seed site (n=3327)
Removed seed site (n=224)
No seed site (n=7839)

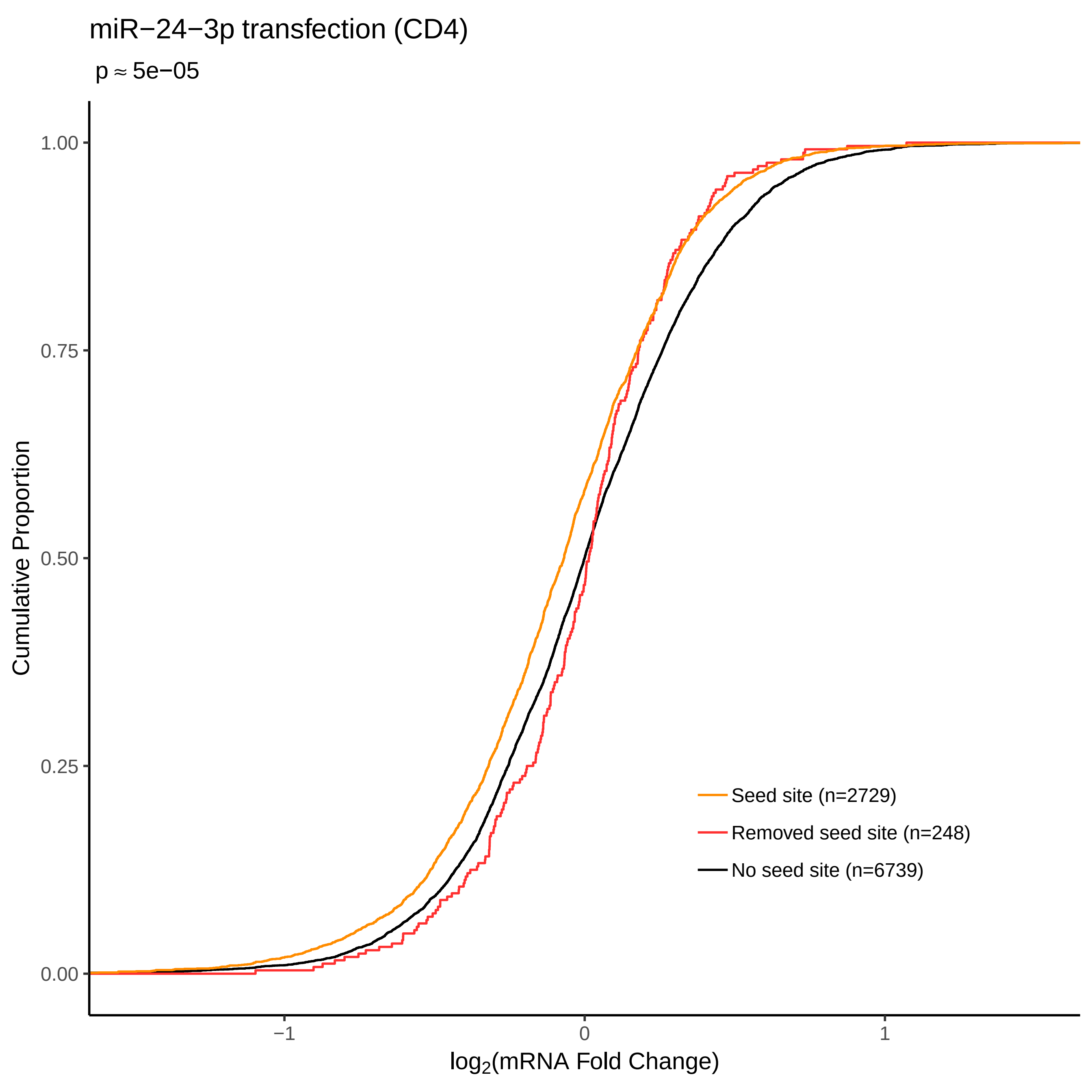

miR−24−3p transfection (CD4)
p ≈ 5e−05
Cumulative Proportion
1.00
0.75
0.50
0.25
0.00
−1
0
1
log2(mRNA Fold Change)
Seed site (n=2729)
Removed seed site (n=248)
No seed site (n=6739)

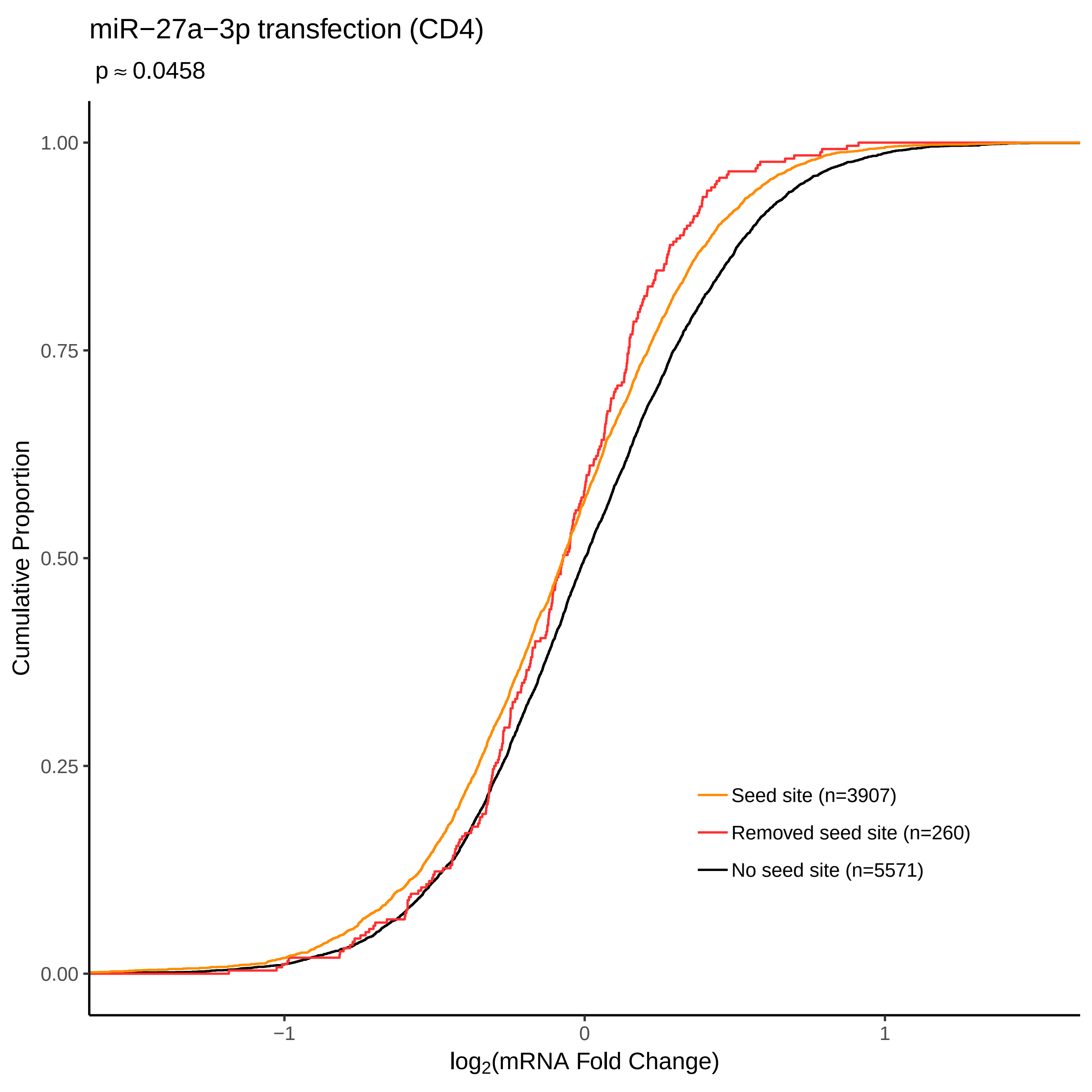
miR−27a−3p transfection (CD4)
p ≈ 0.0458
Cumulative Proportion
1.00
0.75
0.50
0.25
0.00
−1
0
1
log2(mRNA Fold Change)
Seed site (n=3907)
Removed seed site (n=260)
No seed site (n=5571)

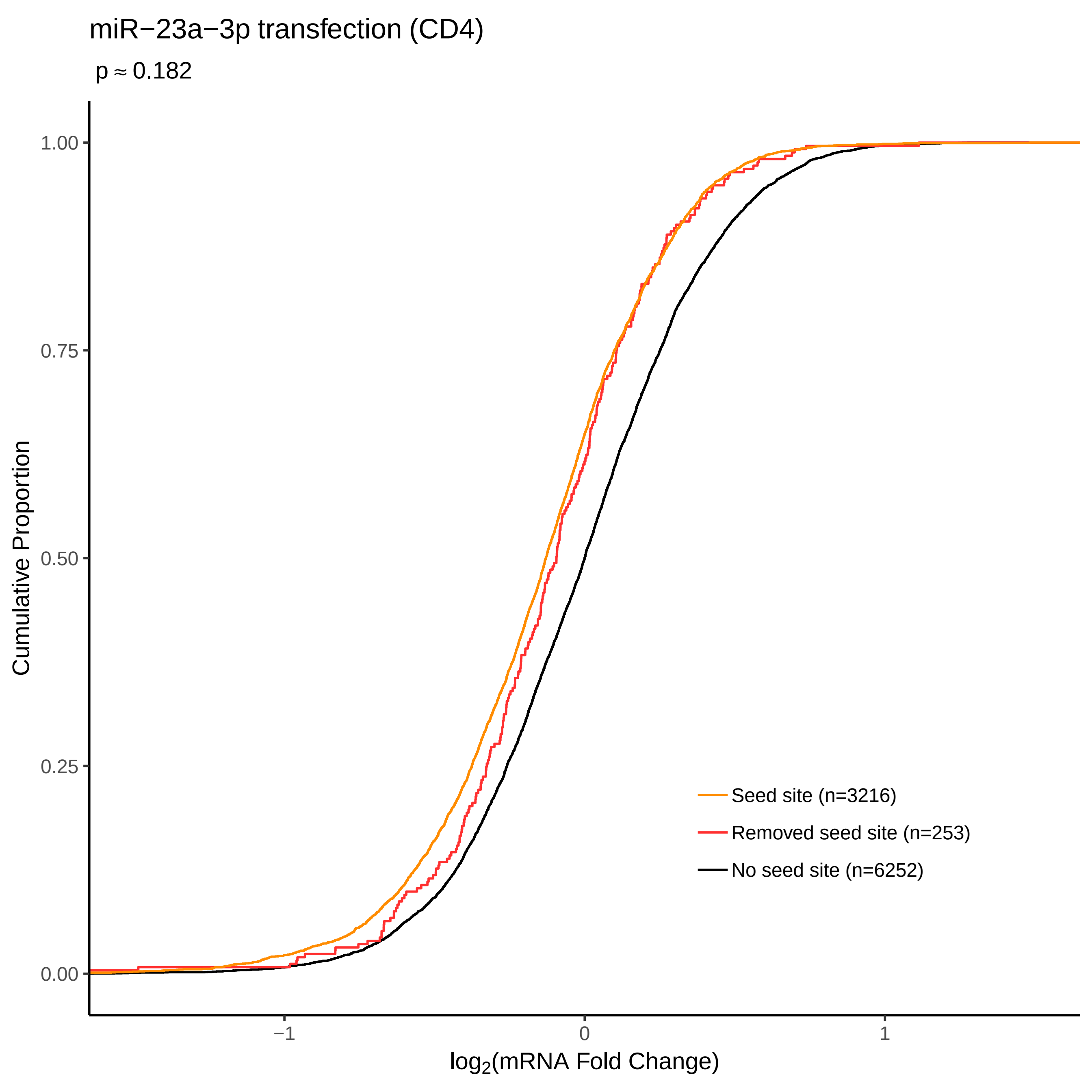

miR−23a−3p transfection (CD4)
p ≈ 0.182
Cumulative Proportion
1.00
0.75
0.50
0.25
0.00
−1
0
1
log2(mRNA Fold Change)
Seed site (n=3216)
Removed seed site (n=253)
No seed site (n=6252)

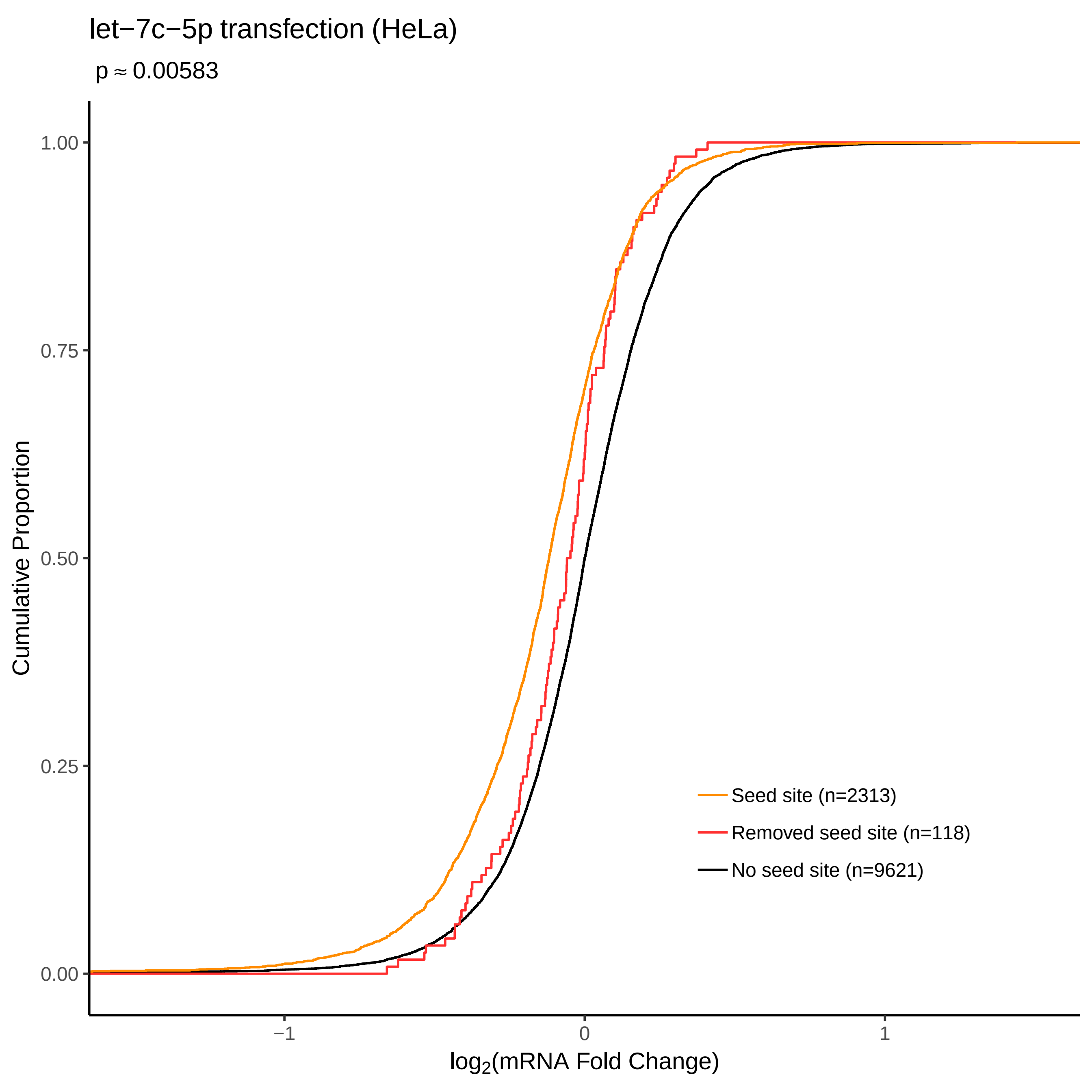
let−7c−5p transfection (HeLa)
p ≈ 0.00583
Cumulative Proportion
1.00
0.75
0.50
0.25
0.00
−1
0
1
log2(mRNA Fold Change)
Seed site (n=2313)
Removed seed site (n=118)
No seed site (n=9621)

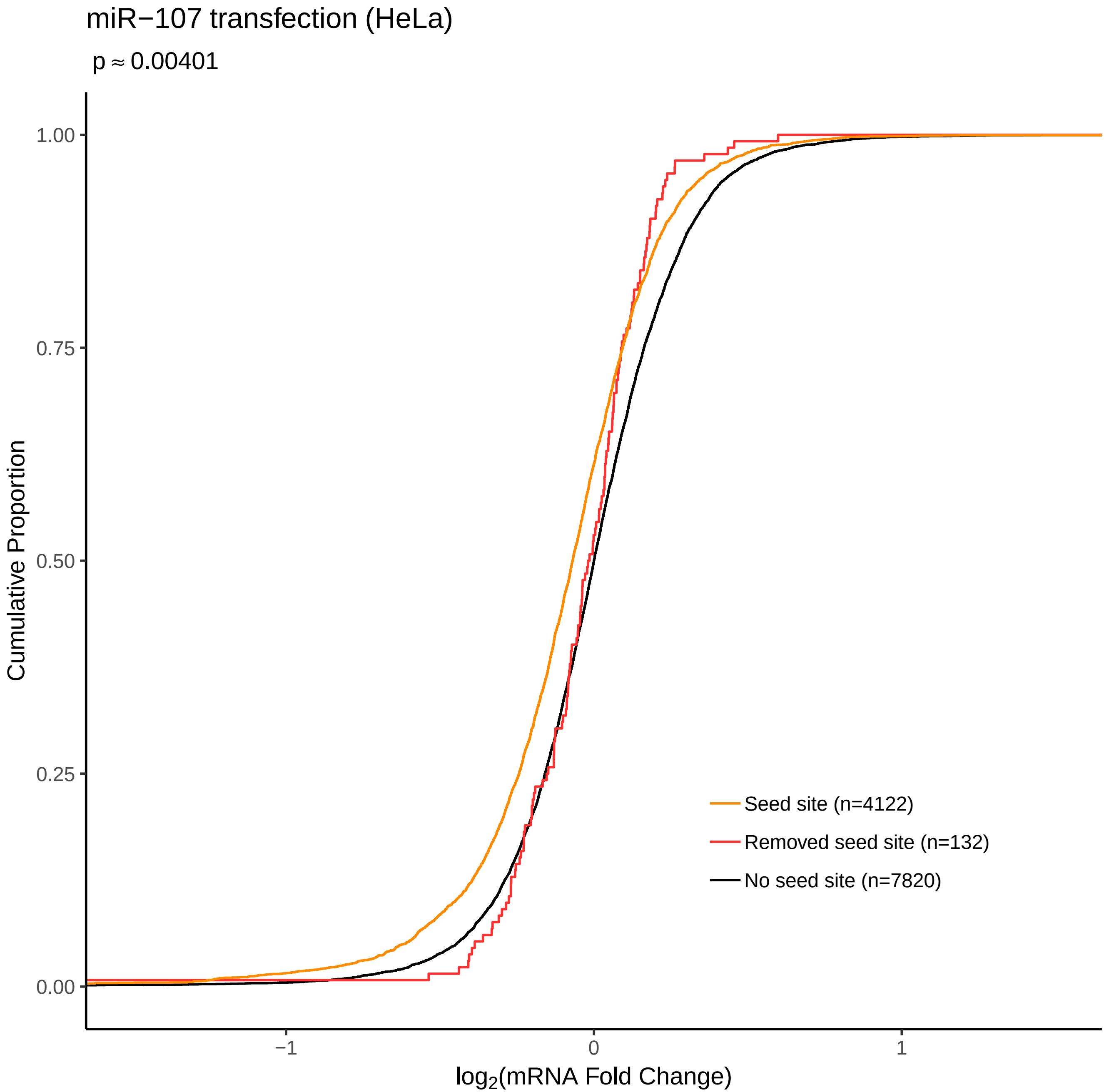

miR−107 transfection (HeLa)
p ≈ 0.00401
Cumulative Proportion
1.00
0.75
0.50
0.25
0.00
−1
0
1
log2(mRNA Fold Change)
Seed site (n=4122)
Removed seed site (n=132)
No seed site (n=7820)

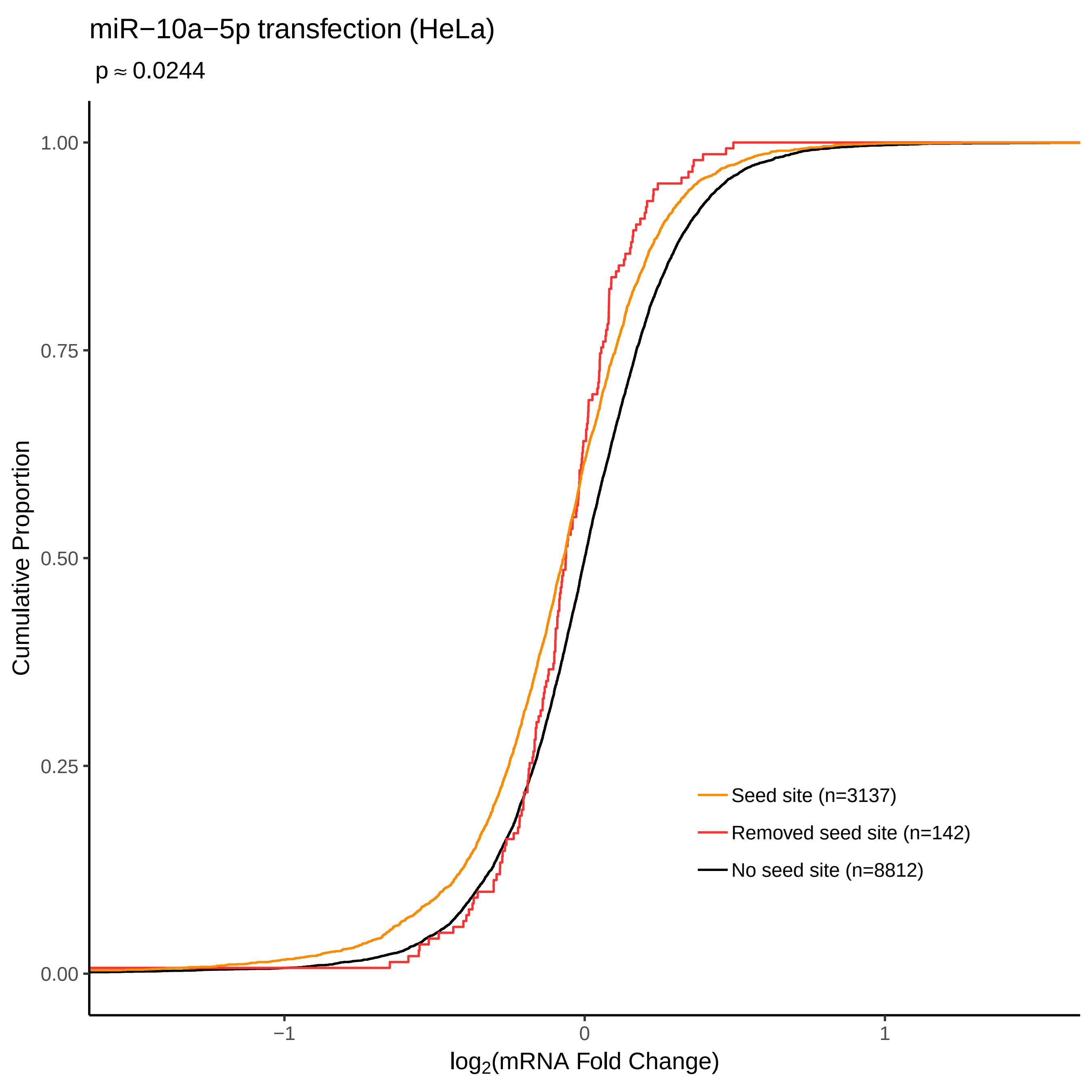
miR−10a−5p transfection (HeLa)
p ≈ 0.0244
Cumulative Proportion
1.00
0.75
0.50
0.25
0.00
−1
0
1
log2(mRNA Fold Change)
Seed site (n=3137)
Removed seed site (n=142)
No seed site (n=8812)

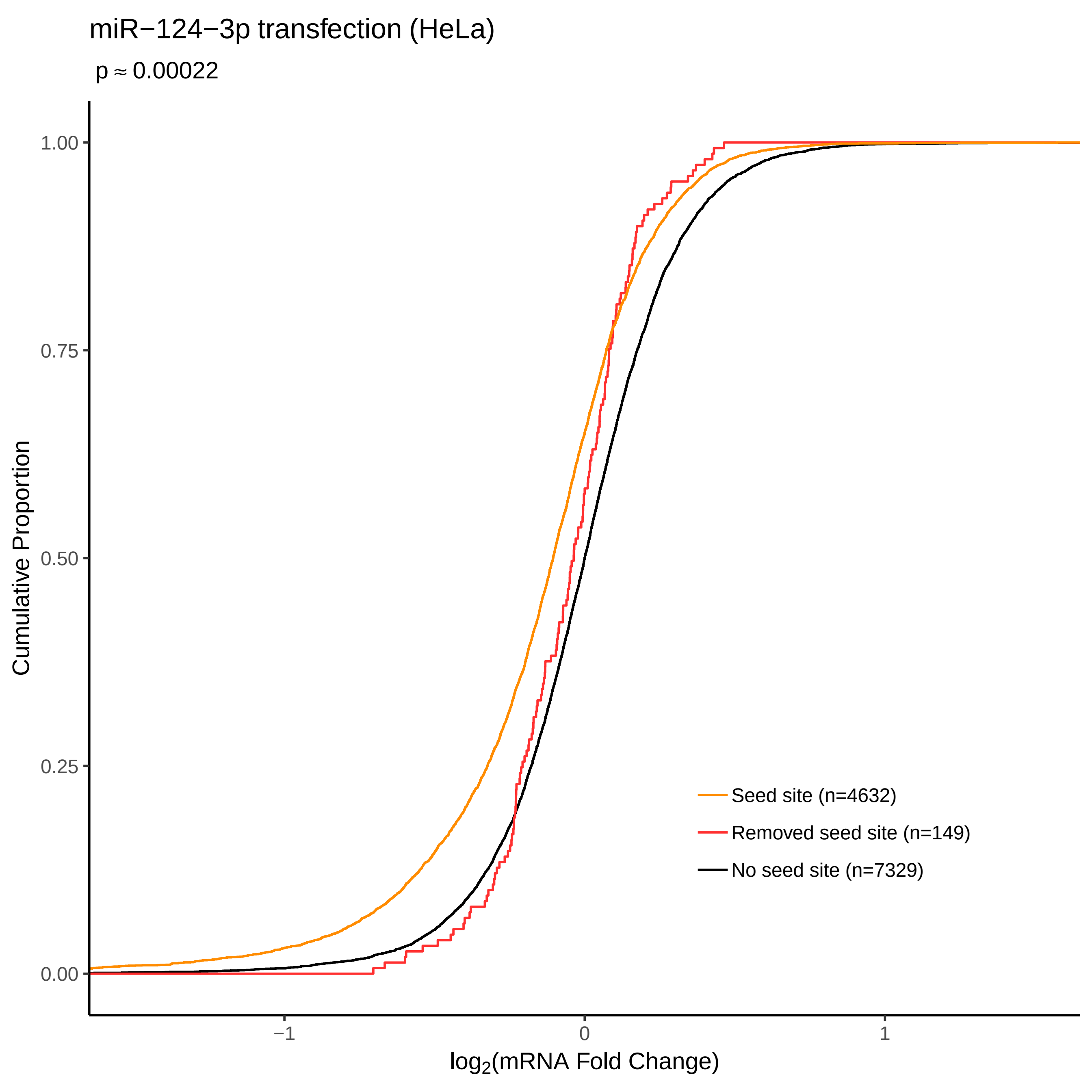

miR−124−3p transfection (HeLa)
p ≈ 0.00022
Cumulative Proportion
1.00
0.75
0.50
0.25
0.00
−1
0
1
log2(mRNA Fold Change)
Seed site (n=4632)
Removed seed site (n=149)
No seed site (n=7329)

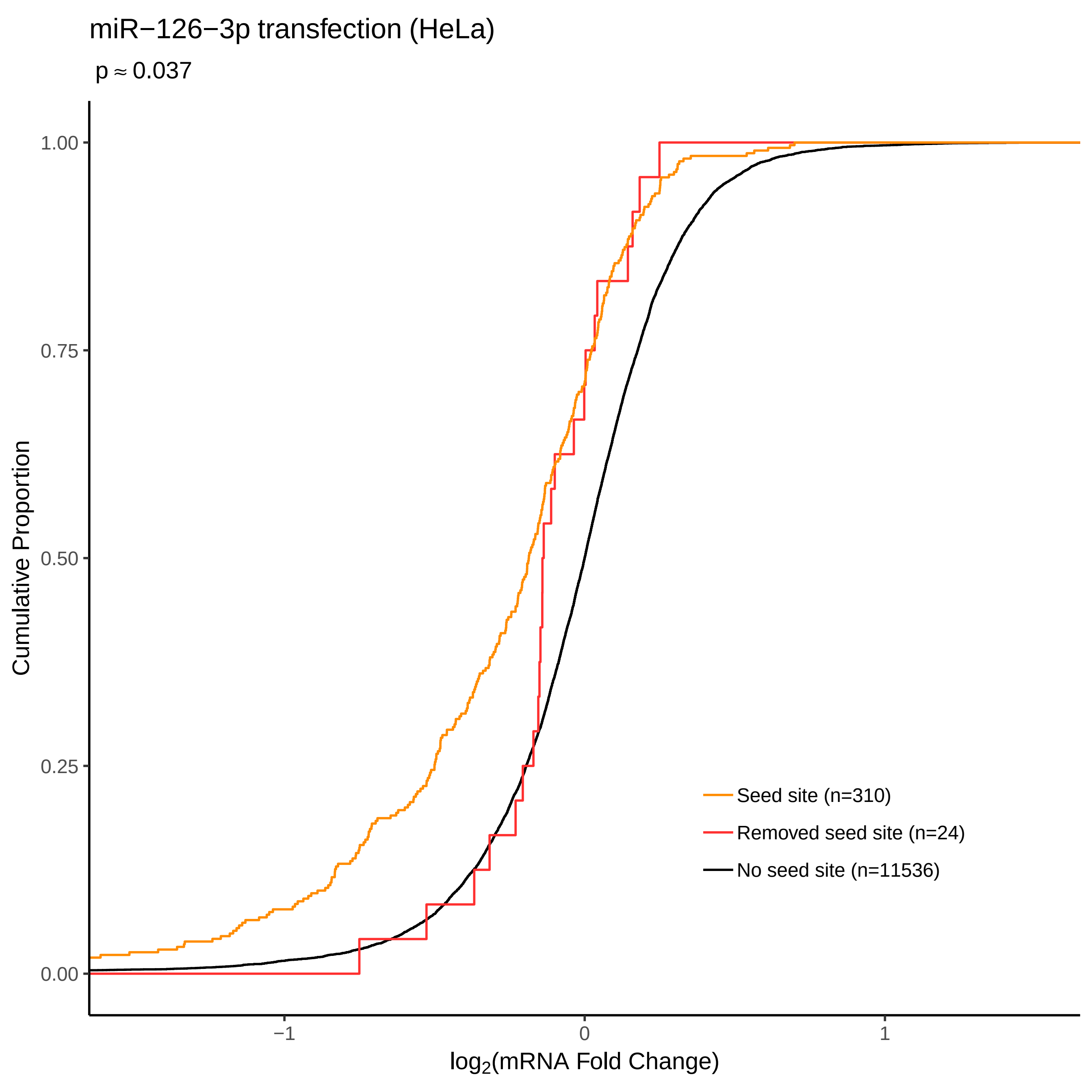
miR−126−3p transfection (HeLa)
p ≈ 0.037
Cumulative Proportion
1.00
0.75
0.50
0.25
0.00
−1
0
1
log2(mRNA Fold Change)
Seed site (n=310)
Removed seed site (n=24)
No seed site (n=11536)

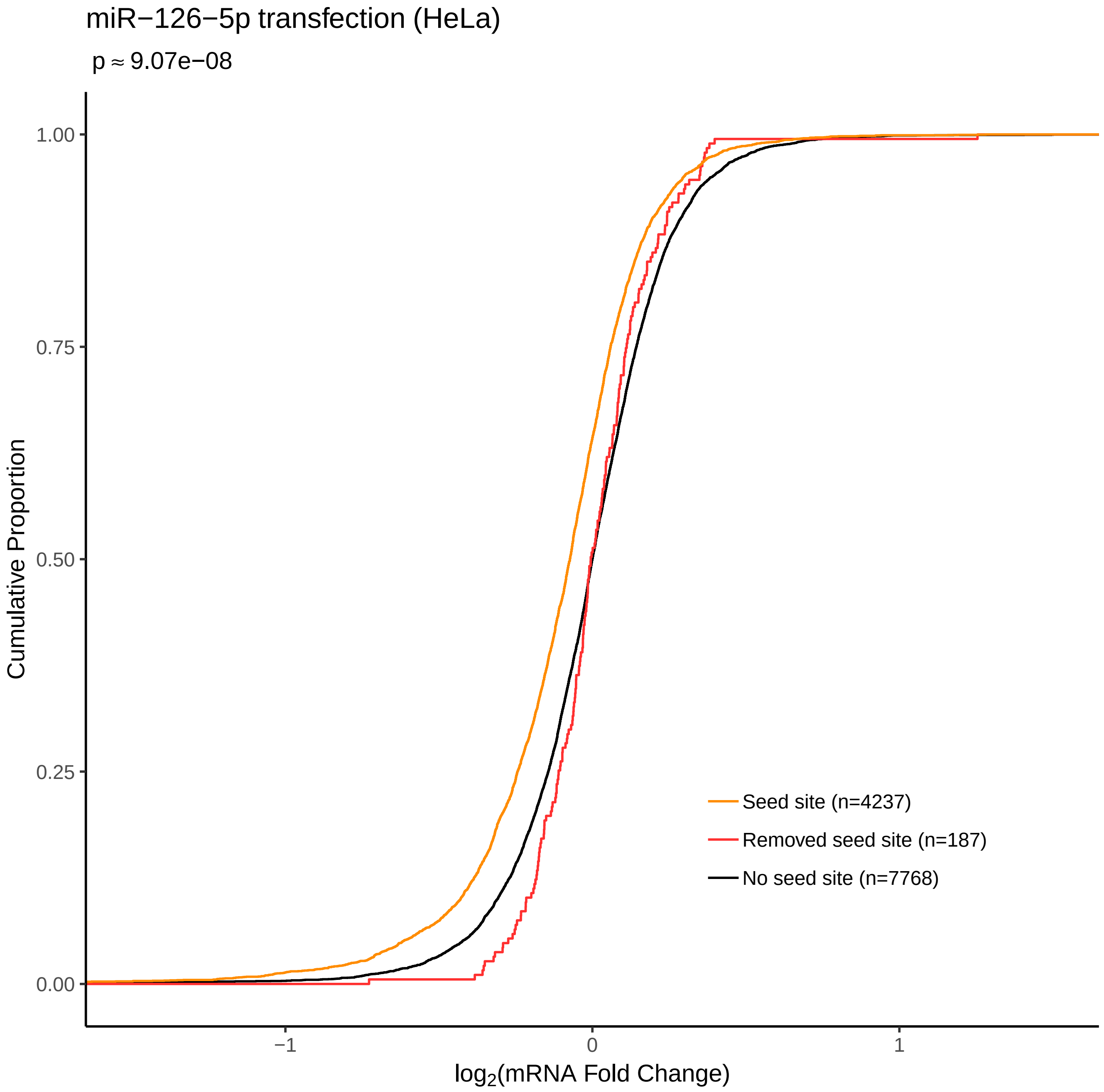
miR−126−5p transfection (HeLa)
p ≈ 9.07e−08
Cumulative Proportion
1.00
0.75
0.50
0.25
0.00
−1
0
1
log2(mRNA Fold Change)
Seed site (n=4237)
Removed seed site (n=187)
No seed site (n=7768)

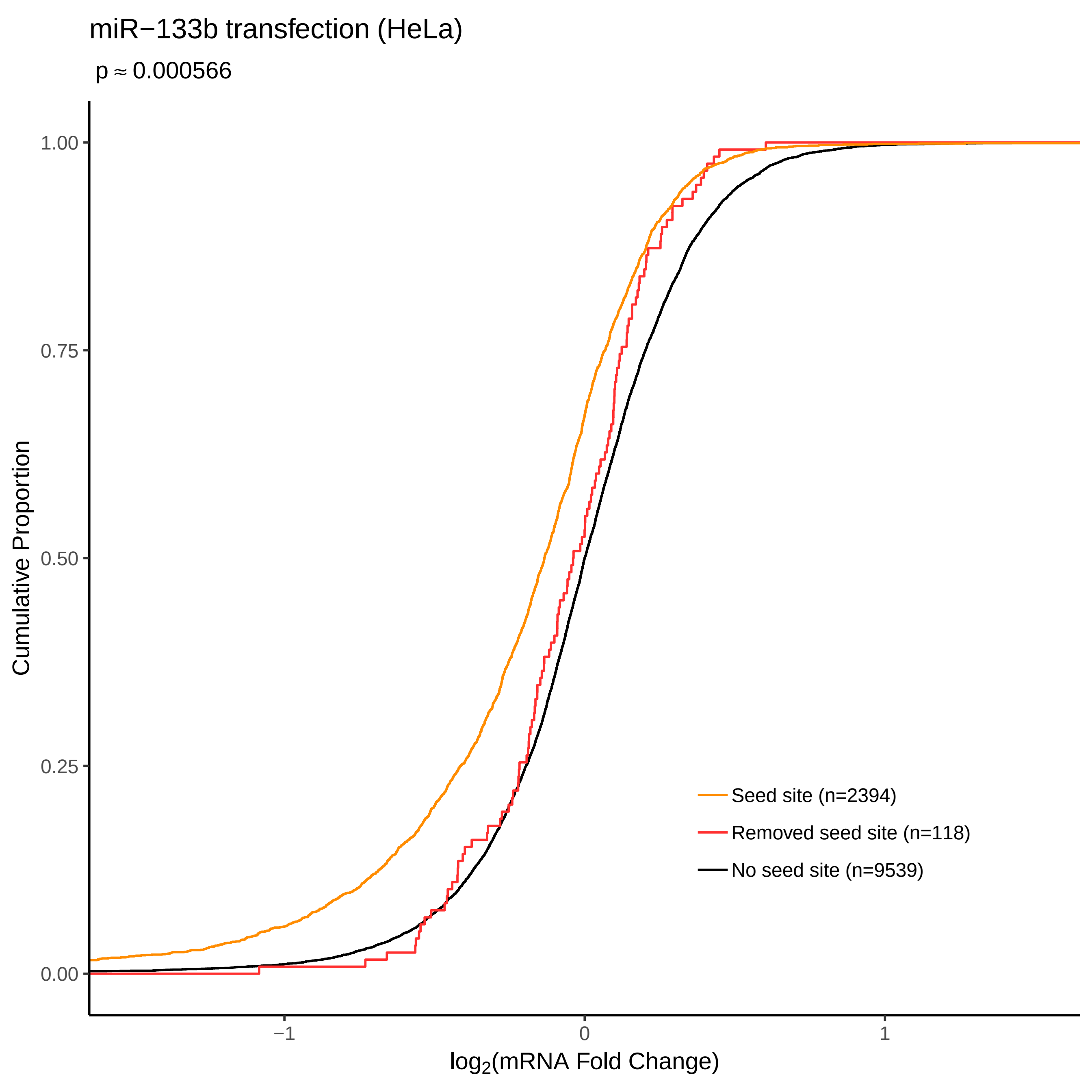
miR−133b transfection (HeLa)
p ≈ 0.000566
Cumulative Proportion
1.00
0.75
0.50
0.25
0.00
−1
0
1
log2(mRNA Fold Change)
Seed site (n=2394)
Removed seed site (n=118)
No seed site (n=9539)

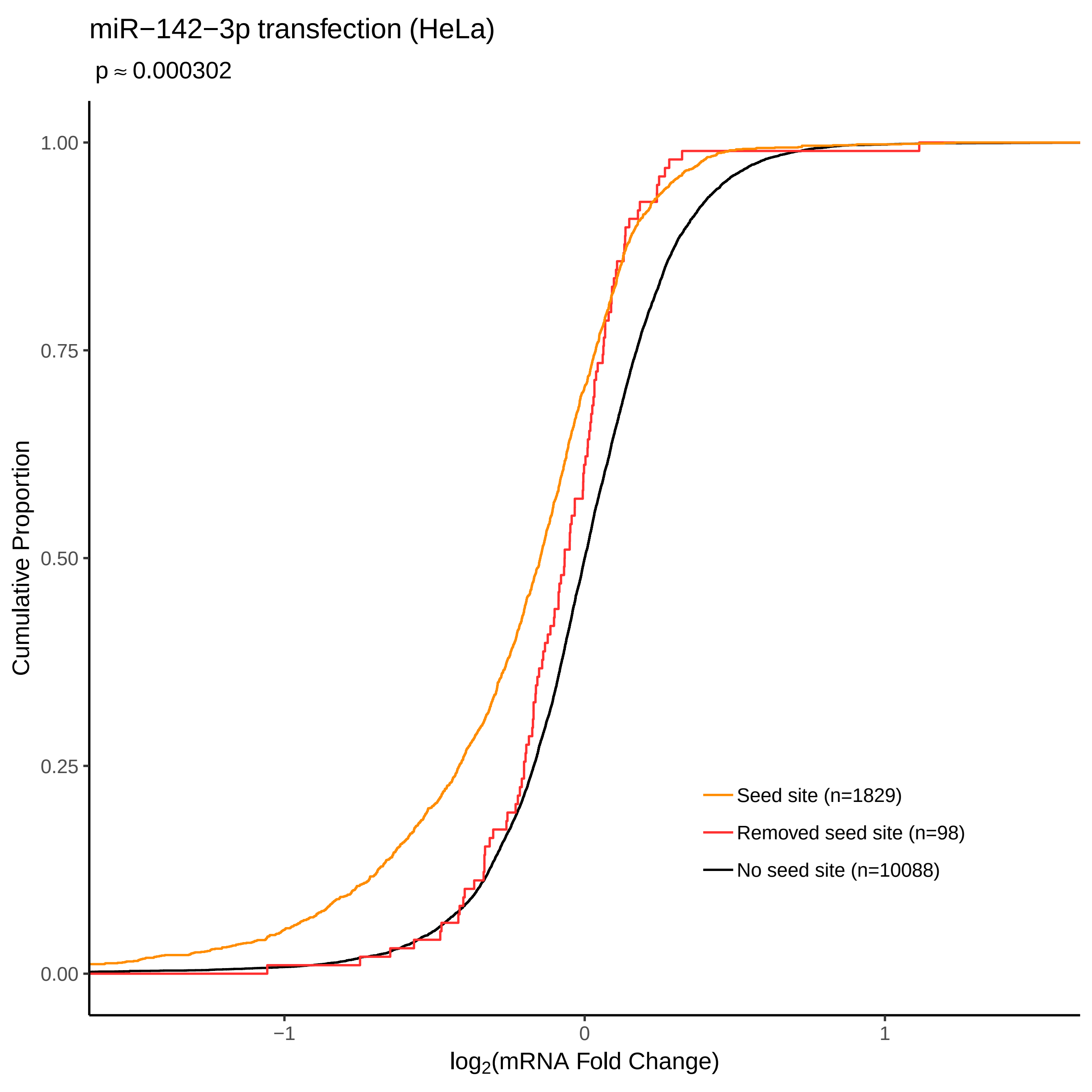
miR−142−3p transfection (HeLa)
p ≈ 0.000302
Cumulative Proportion
1.00
0.75
0.50
0.25
0.00
−1
0
1
log2(mRNA Fold Change)
Seed site (n=1829)
Removed seed site (n=98)
No seed site (n=10088)

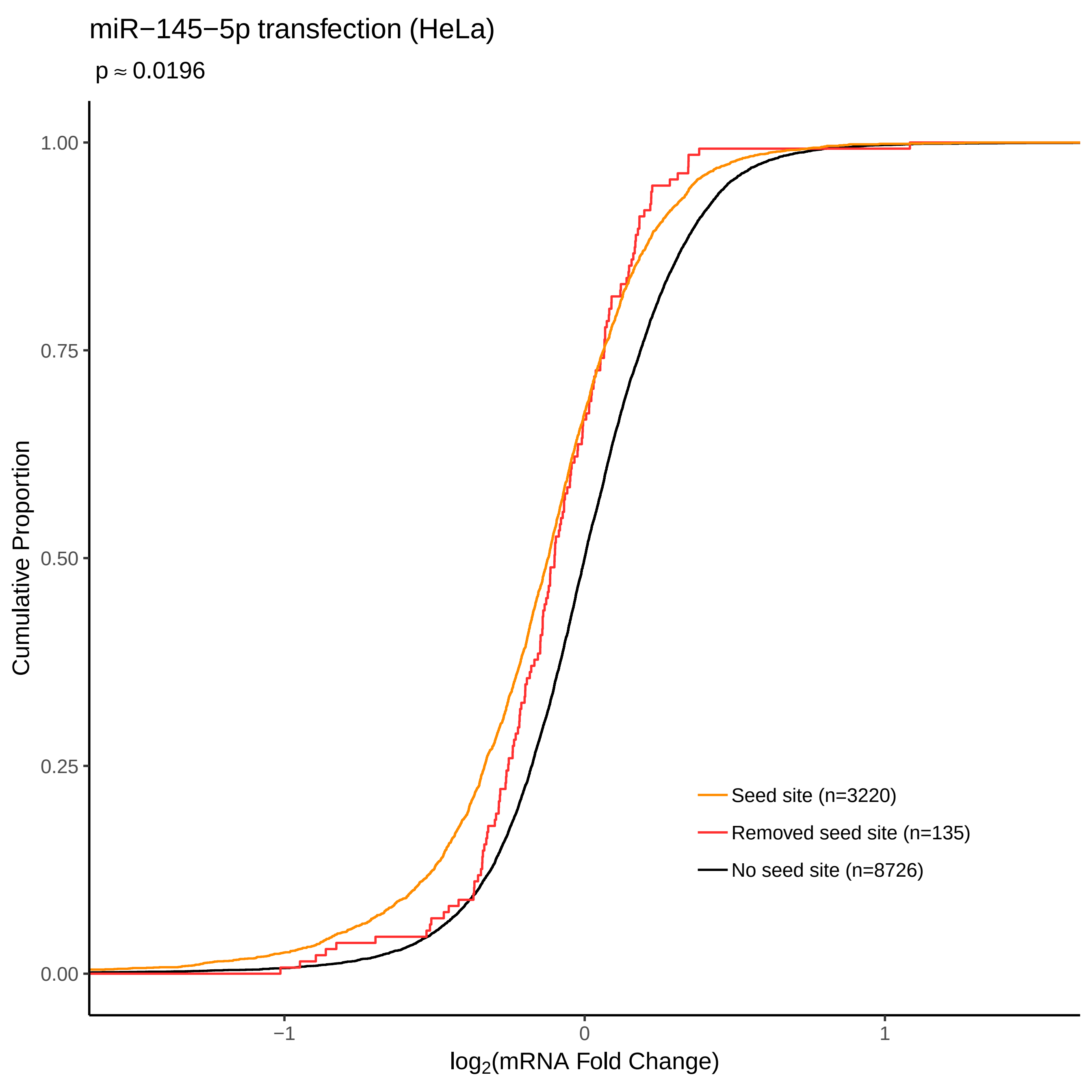

miR−145−5p transfection (HeLa)
p ≈ 0.0196
Cumulative Proportion
1.00
0.75
0.50
0.25
0.00
−1
0
1
log2(mRNA Fold Change)
Seed site (n=3220)
Removed seed site (n=135)
No seed site (n=8726)

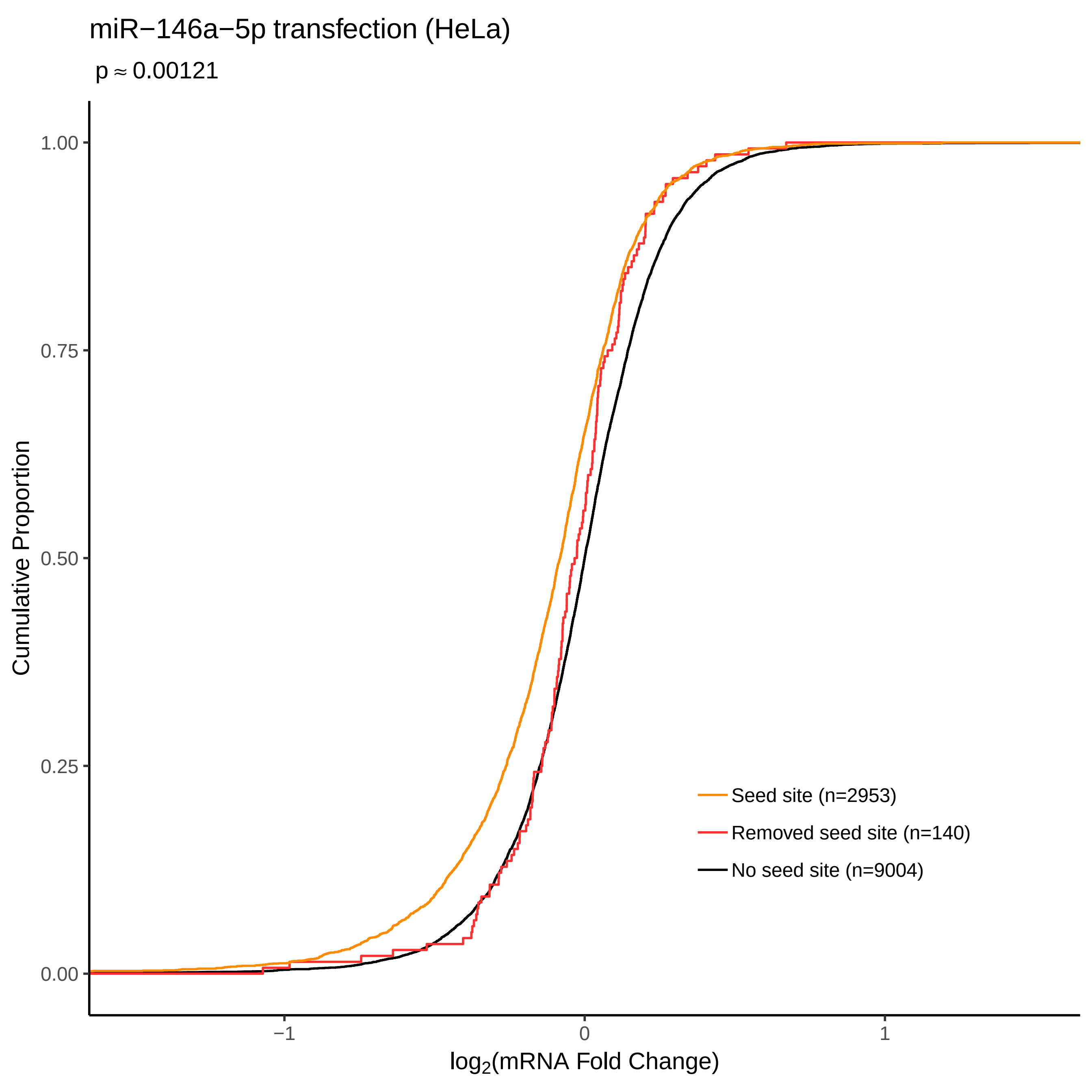

miR−146a−5p transfection (HeLa)
p ≈ 0.00121
Cumulative Proportion
1.00
0.75
0.50
0.25
0.00
−1
0
1
log2(mRNA Fold Change)
Seed site (n=2953)
Removed seed site (n=140)
No seed site (n=9004)

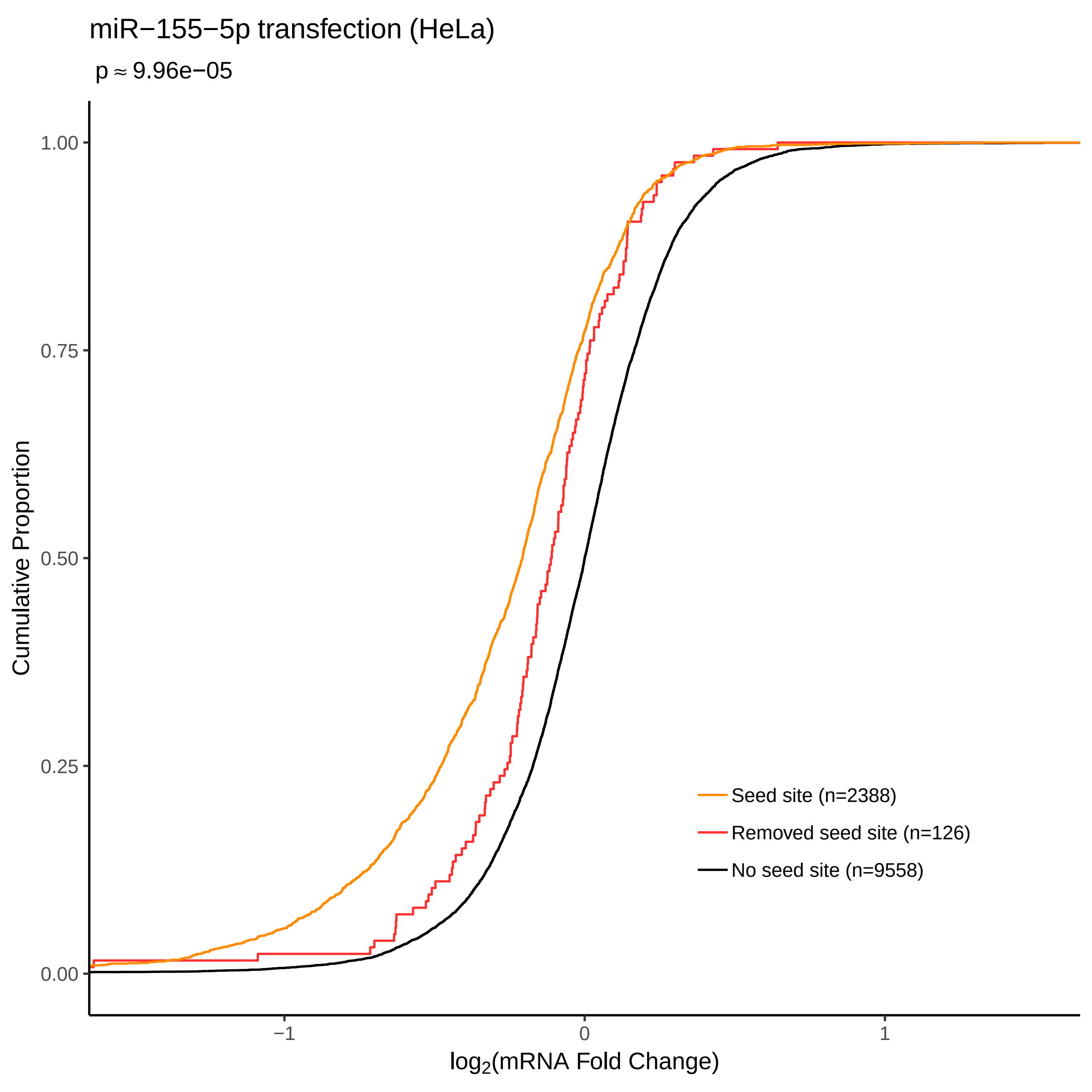

miR−155−5p transfection (HeLa)
p ≈ 9.96e−05
Cumulative Proportion
1.00
0.75
0.50
0.25
0.00
−1
0
1
log2(mRNA Fold Change)
Seed site (n=2388)
Removed seed site (n=126)
No seed site (n=9558)

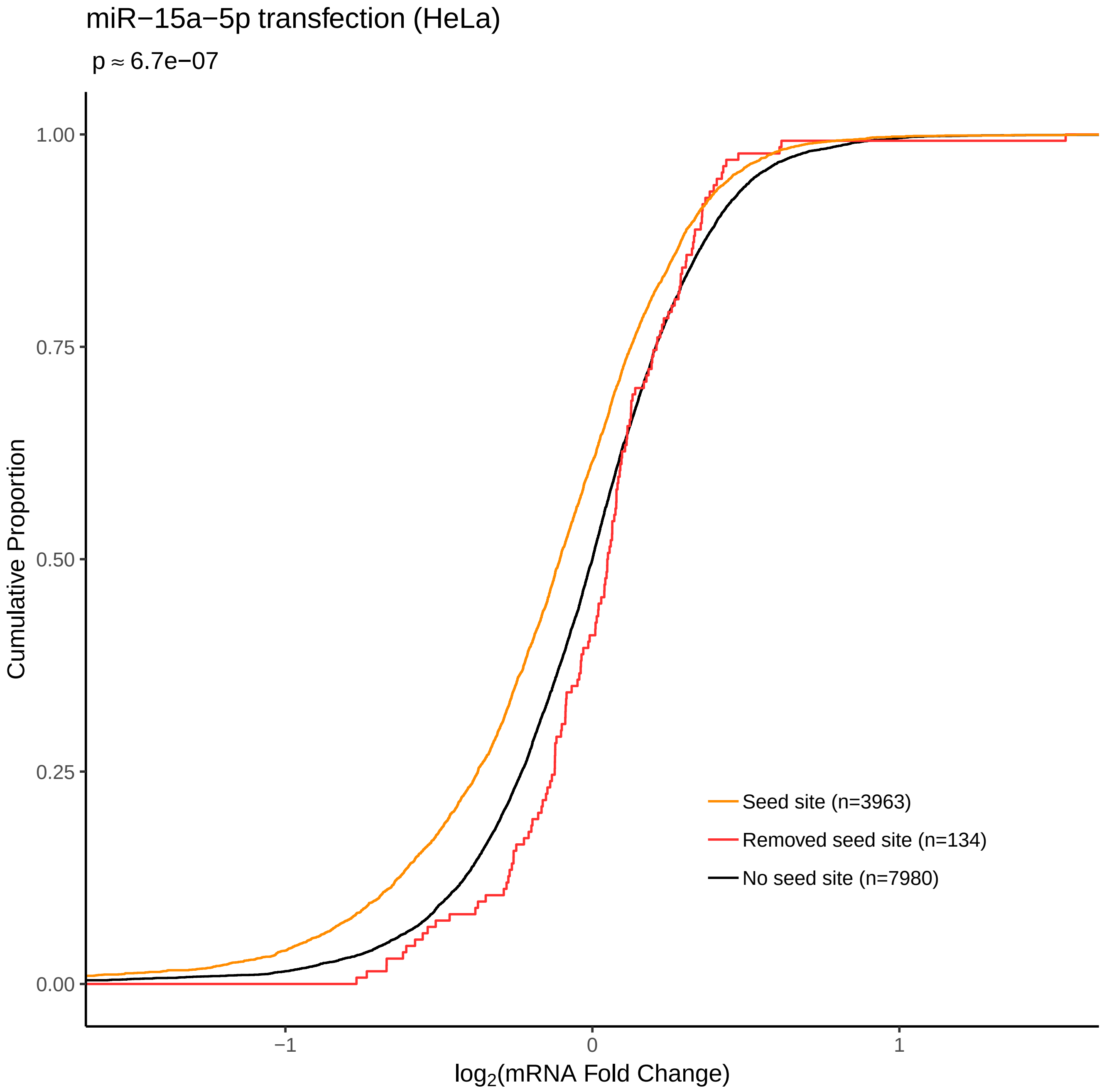
miR−15a−5p transfection (HeLa)
p ≈ 6.7e−07
Cumulative Proportion
1.00
0.75
0.50
0.25
0.00
−1
0
1
log2(mRNA Fold Change)
Seed site (n=3963)
Removed seed site (n=134)
No seed site (n=7980)

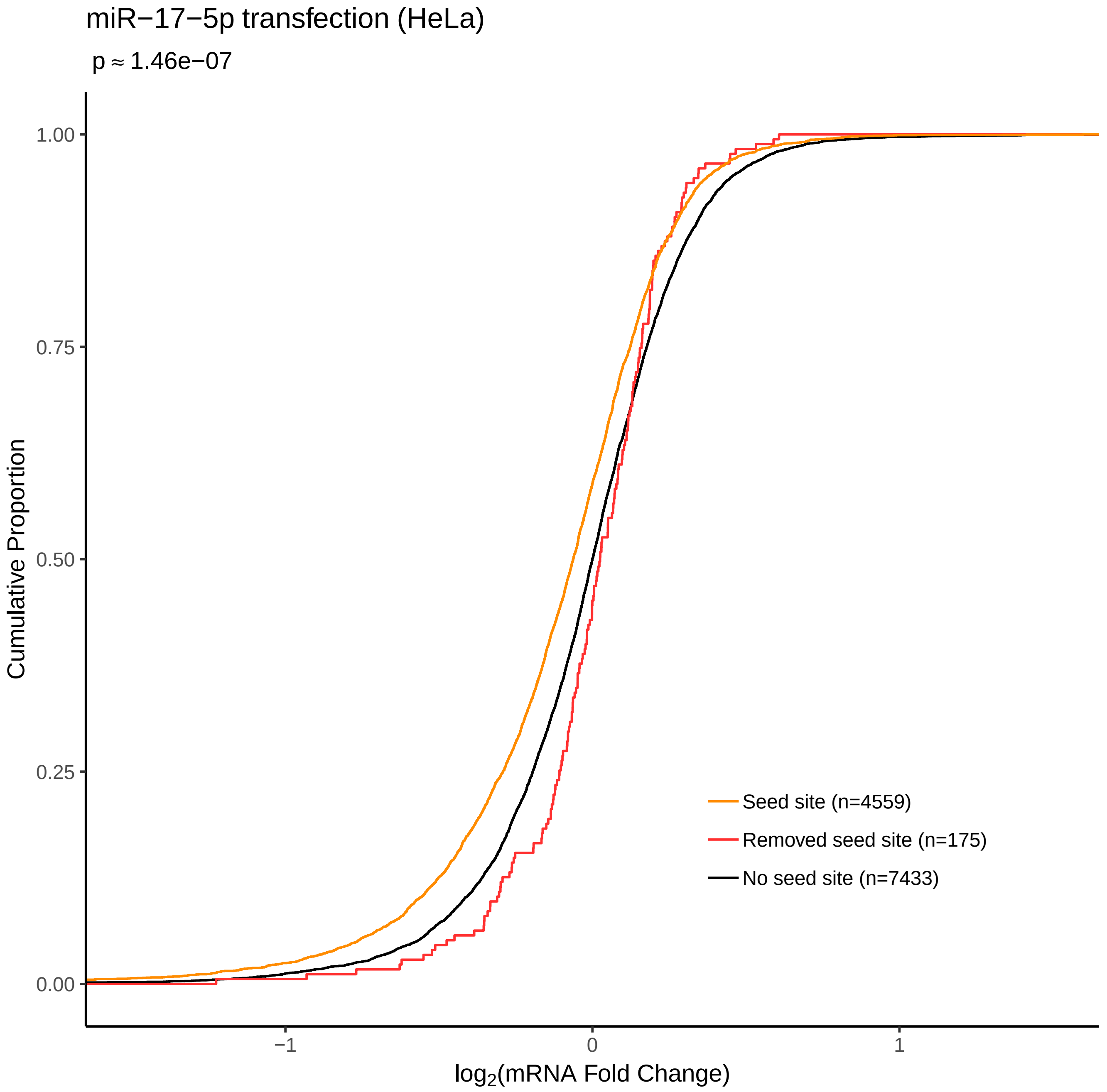

miR−17−5p transfection (HeLa)
p ≈ 1.46e−07
Cumulative Proportion
1.00
0.75
0.50
0.25
0.00
−1
0
1
log2(mRNA Fold Change)
Seed site (n=4559)
Removed seed site (n=175)
No seed site (n=7433)

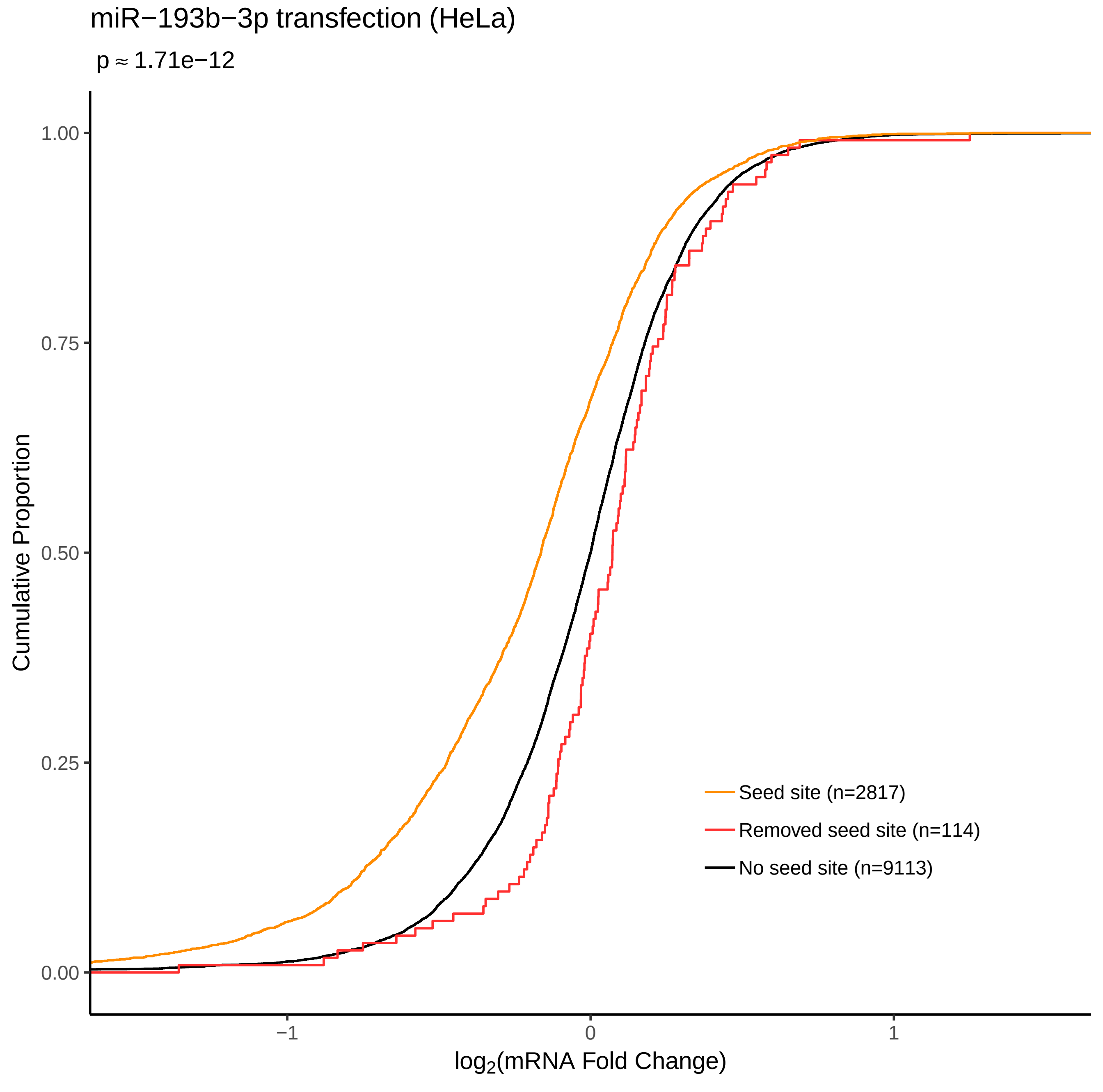

miR−193b−3p transfection (HeLa)
p ≈ 1.71e−12
Cumulative Proportion
1.00
0.75
0.50
0.25
0.00
−1
0
1
log2(mRNA Fold Change)
Seed site (n=2817)
Removed seed site (n=114)
No seed site (n=9113)

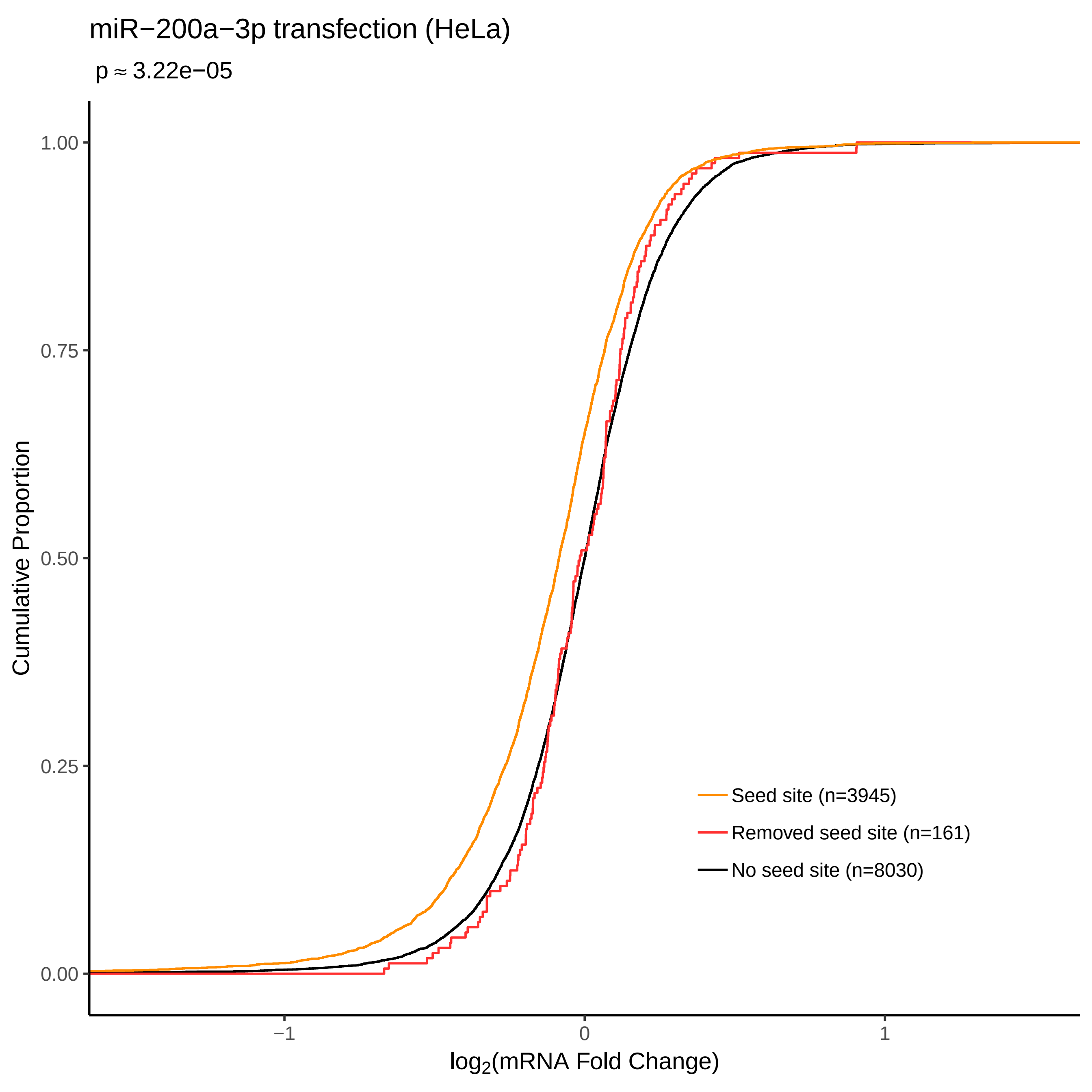
miR−200a−3p transfection (HeLa)
p ≈ 3.22e−05
Cumulative Proportion
1.00
0.75
0.50
0.25
0.00
−1
0
1
log2(mRNA Fold Change)
Seed site (n=3945)
Removed seed site (n=161)
No seed site (n=8030)

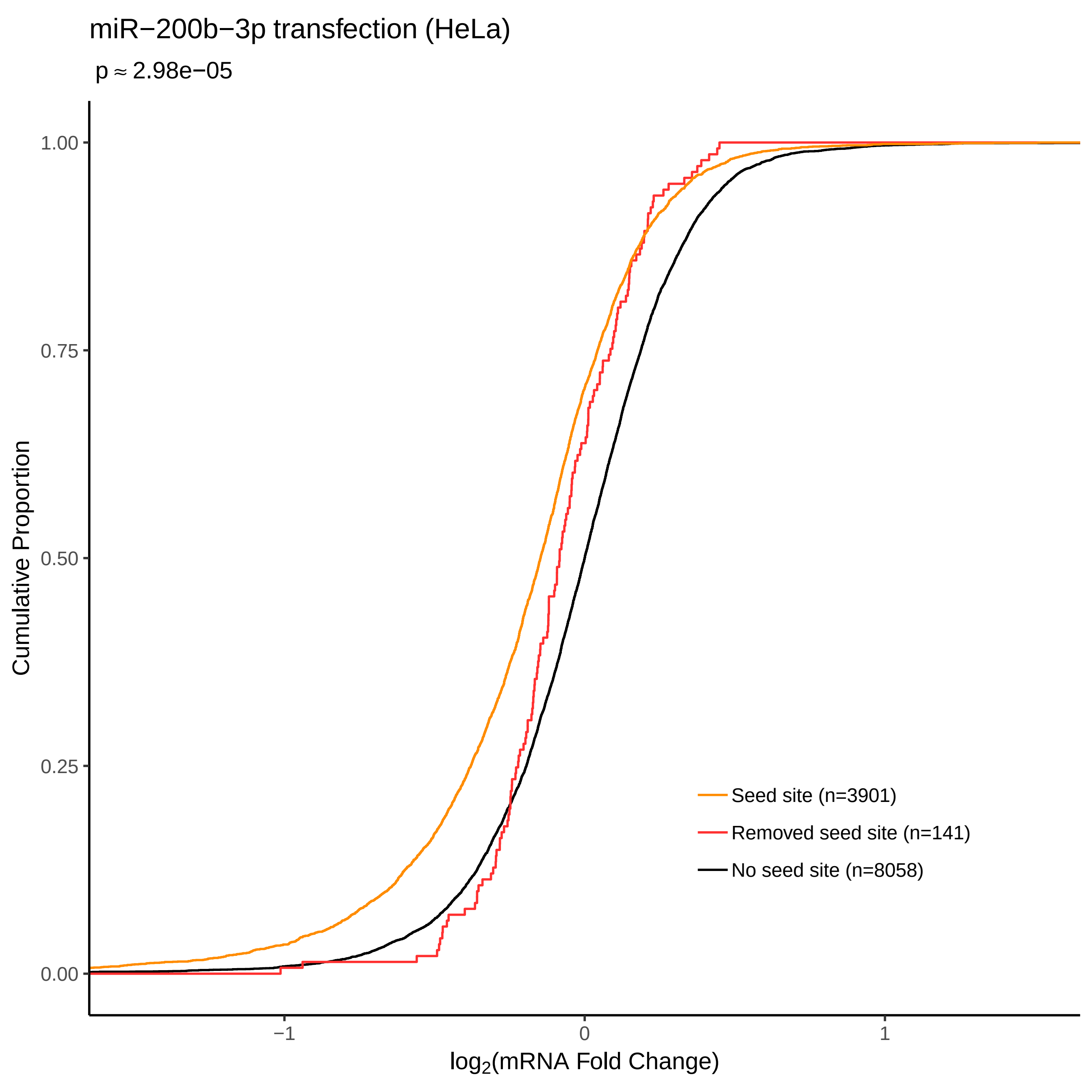
miR−200b−3p transfection (HeLa)
p ≈ 2.98e−05
Cumulative Proportion
1.00
0.75
0.50
0.25
0.00
−1
0
1
log2(mRNA Fold Change)
Seed site (n=3901)
Removed seed site (n=141)
No seed site (n=8058)

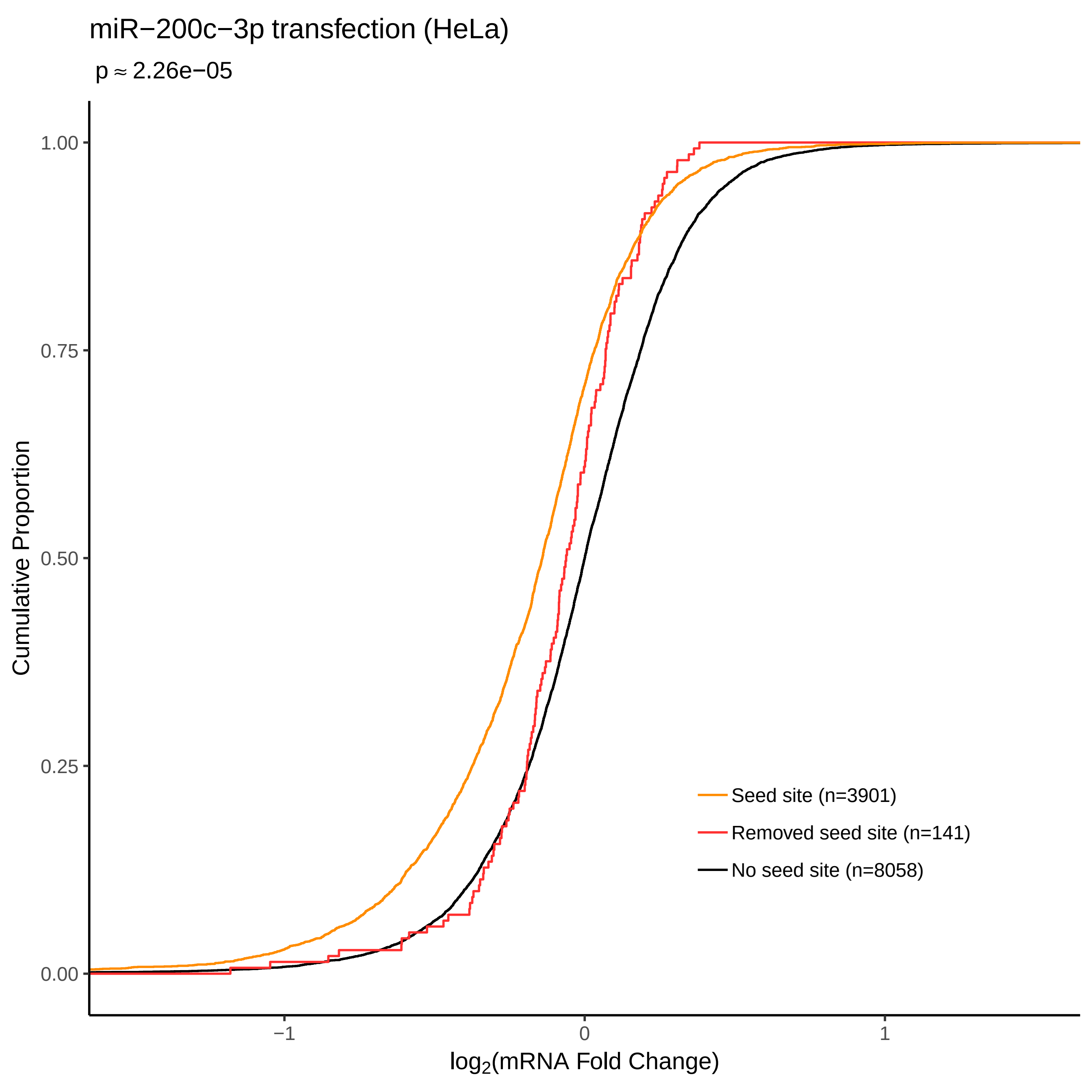
miR−200c−3p transfection (HeLa)
p ≈ 2.26e−05
Cumulative Proportion
1.00
0.75
0.50
0.25
0.00
−1
0
1
log2(mRNA Fold Change)
Seed site (n=3901)
Removed seed site (n=141)
No seed site (n=8058)

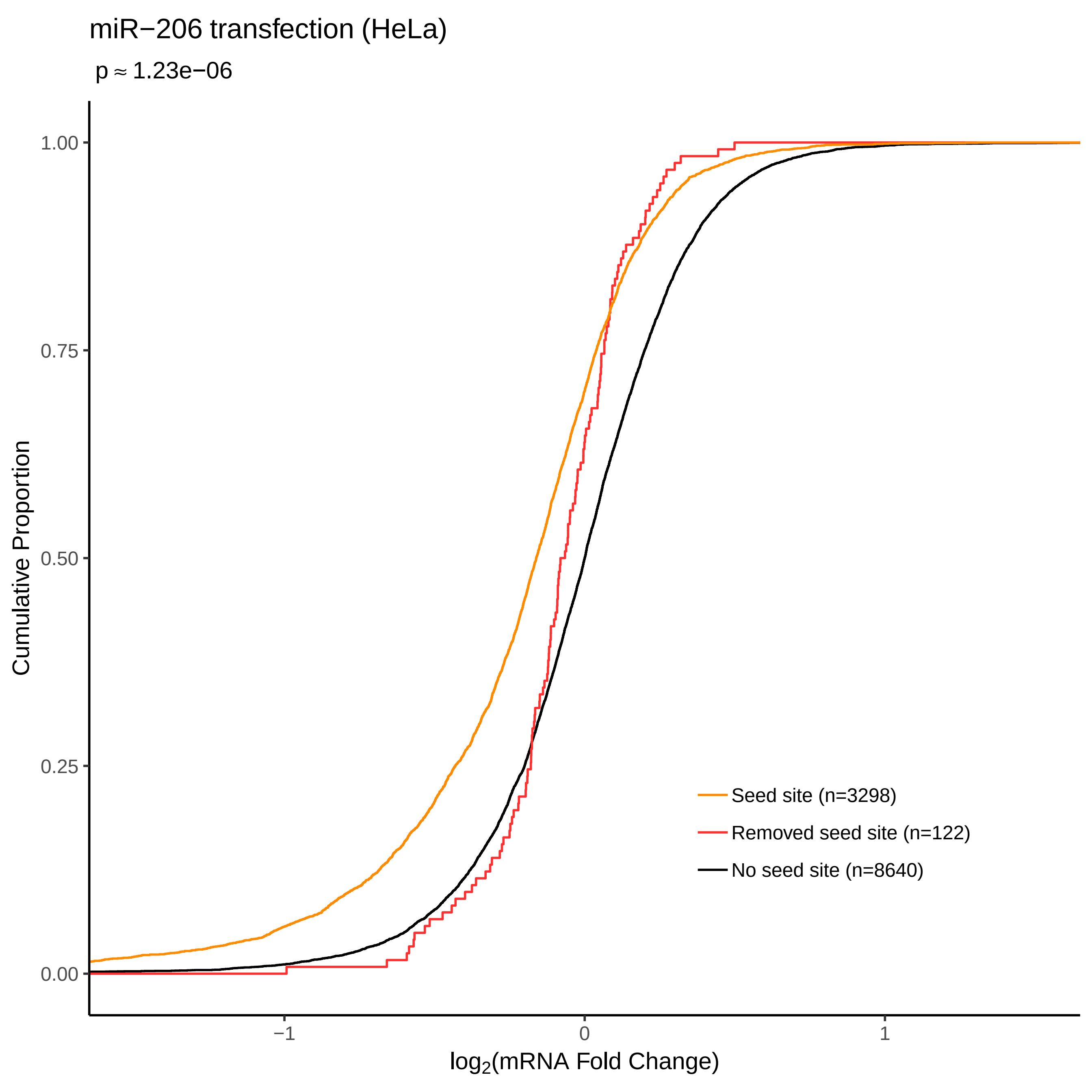
miR−206 transfection (HeLa)
p ≈ 1.23e−06
Cumulative Proportion
1.00
0.75
0.50
0.25
0.00
−1
0
1
log2(mRNA Fold Change)
Seed site (n=3298)
Removed seed site (n=122)
No seed site (n=8640)

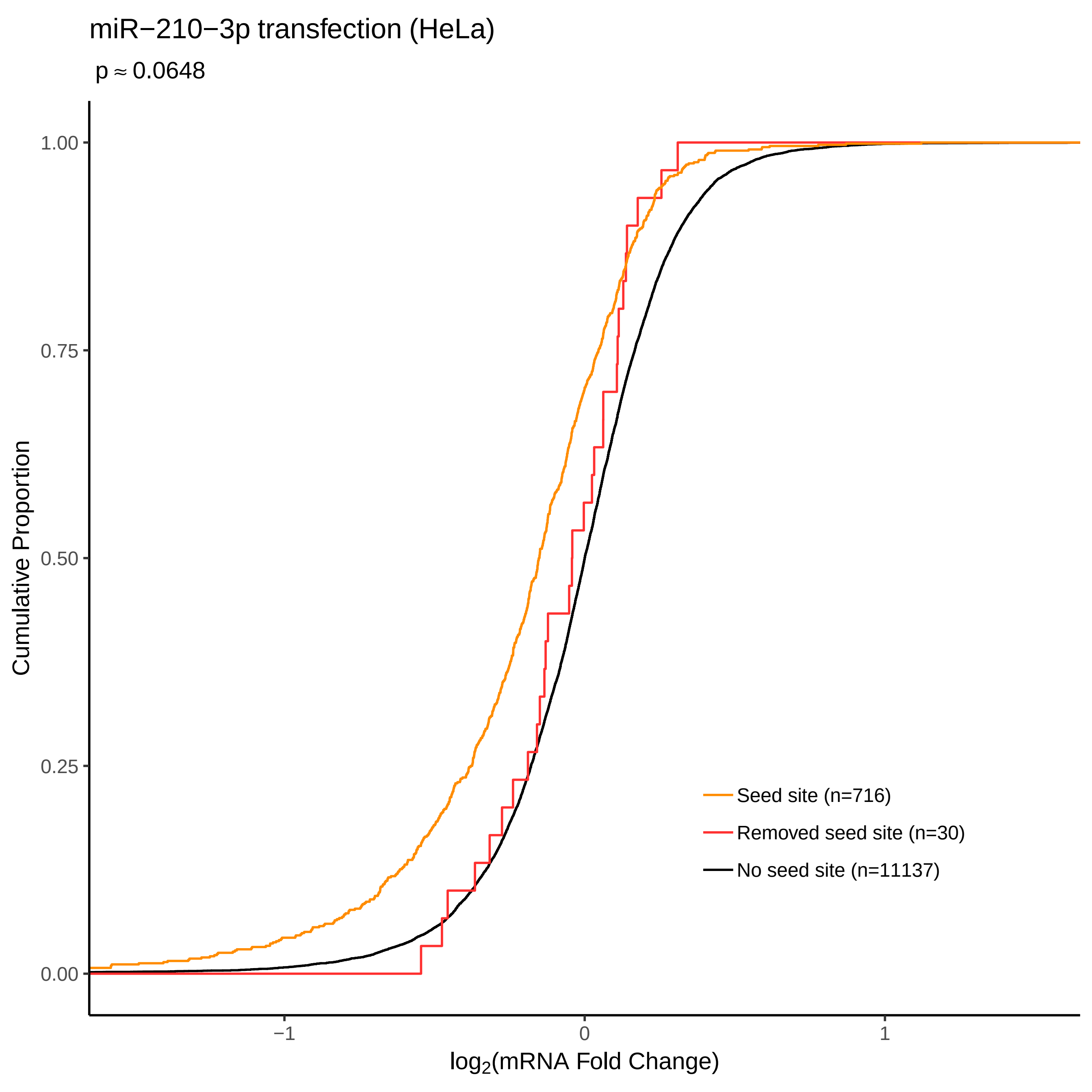
miR−210−3p transfection (HeLa)
p ≈ 0.0648
Cumulative Proportion
1.00
0.75
0.50
0.25
0.00
−1
0
1
log2(mRNA Fold Change)
Seed site (n=716)
Removed seed site (n=30)
No seed site (n=11137)

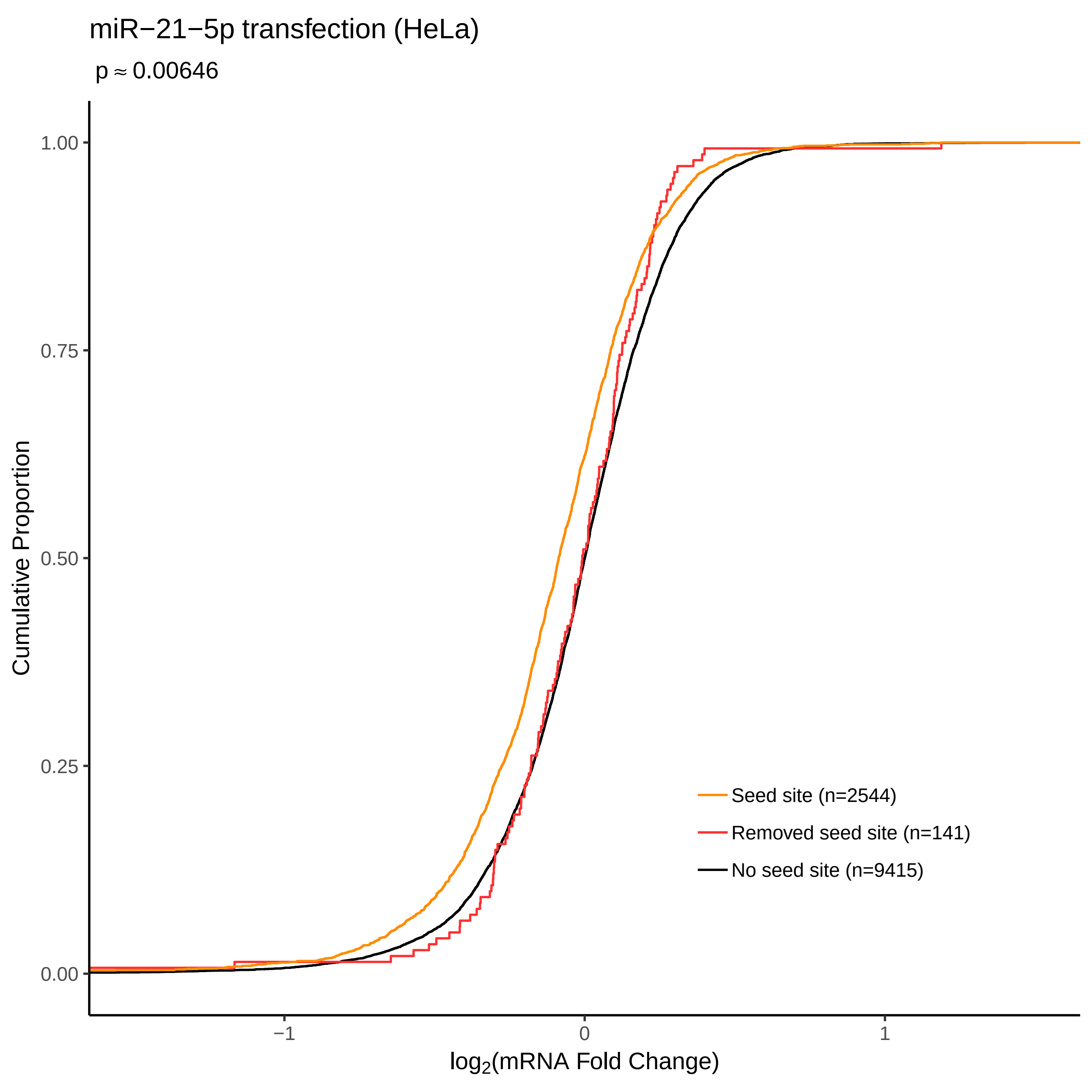
miR−21−5p transfection (HeLa)
p ≈ 0.00646
Cumulative Proportion
1.00
0.75
0.50
0.25
0.00
−1
0
1
log2(mRNA Fold Change)
Seed site (n=2544)
Removed seed site (n=141)
No seed site (n=9415)

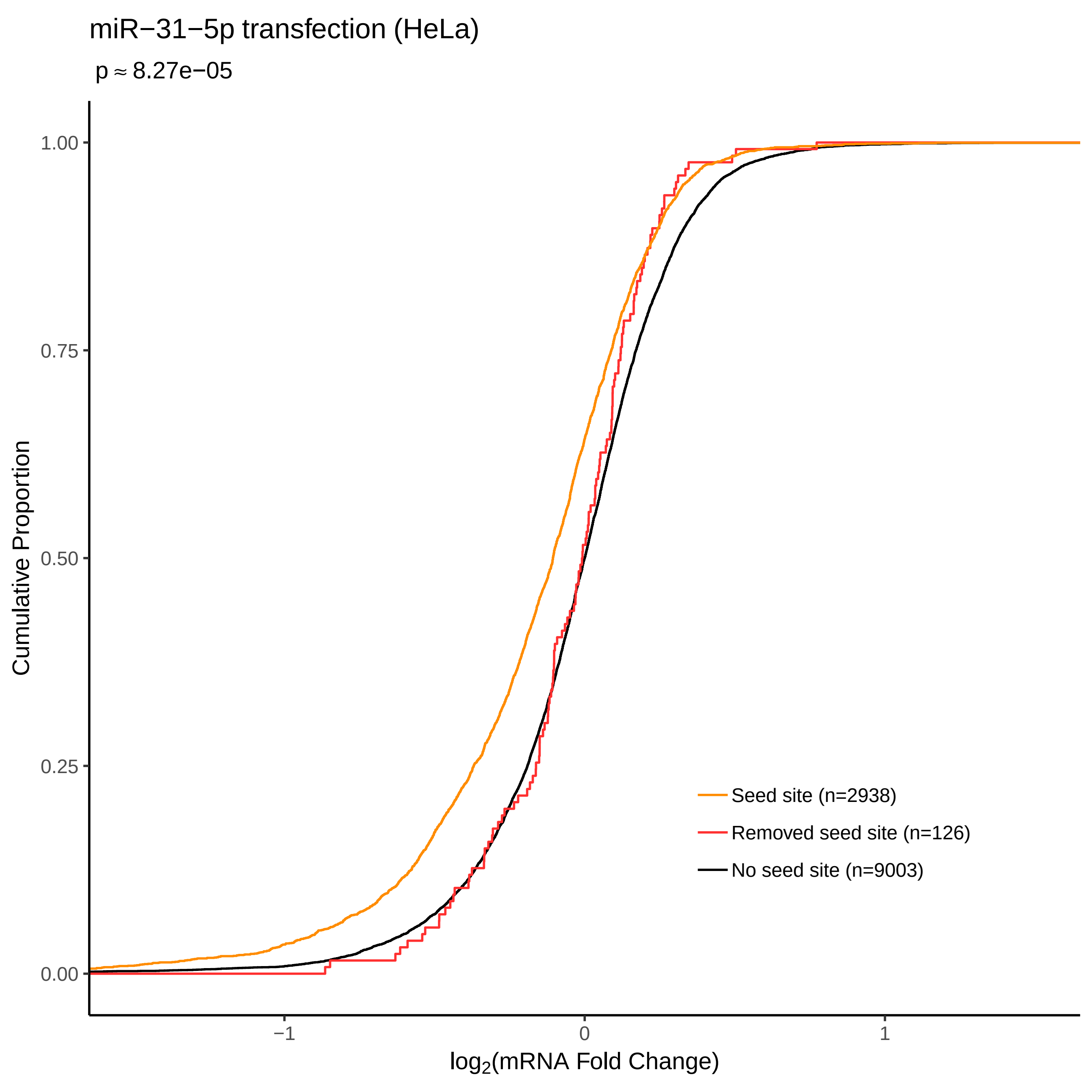

miR−31−5p transfection (HeLa)
p ≈ 8.27e−05
Cumulative Proportion
1.00
0.75
0.50
0.25
0.00
−1
0
1
log2(mRNA Fold Change)
Seed site (n=2938)
Removed seed site (n=126)
No seed site (n=9003)

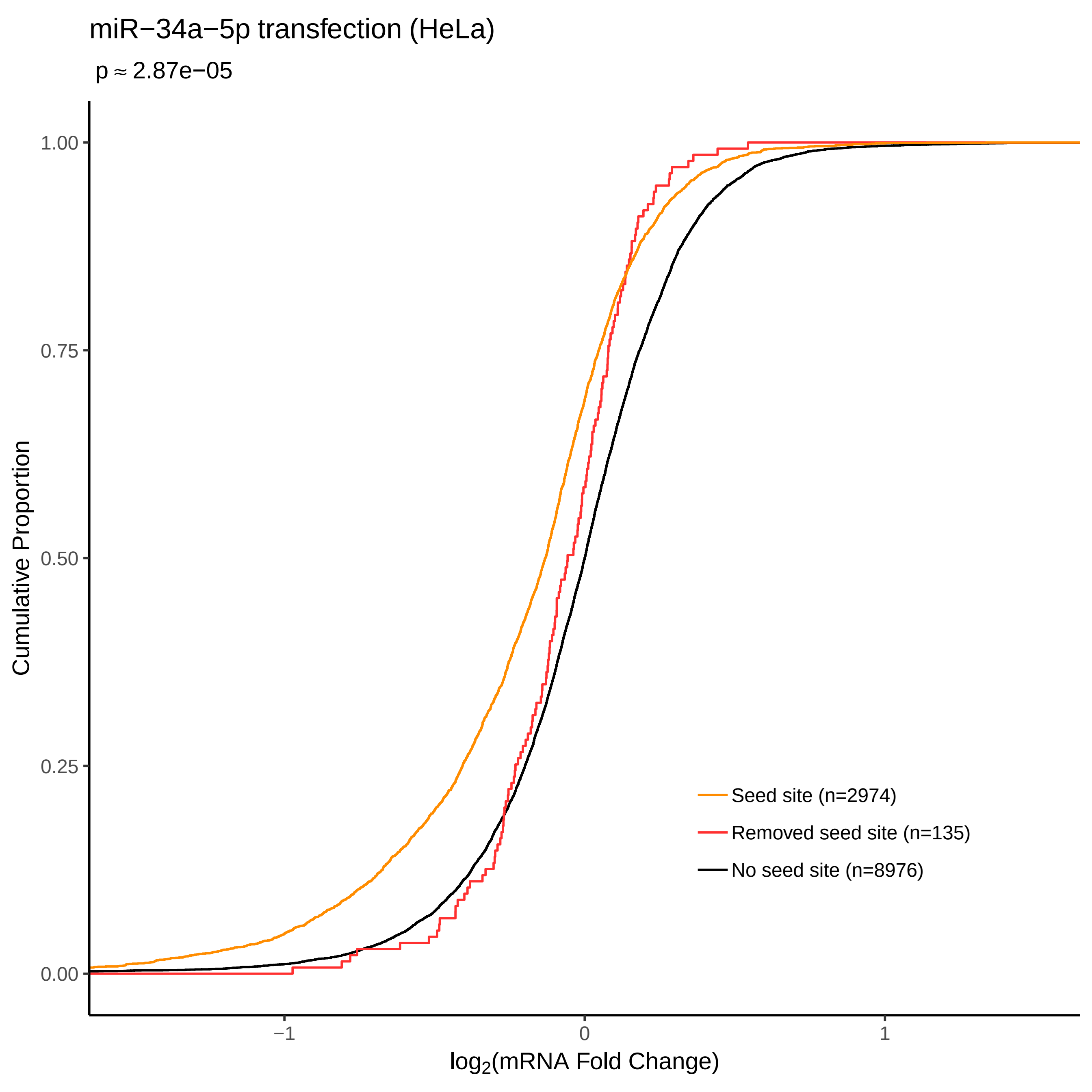

miR−34a−5p transfection (HeLa)
p ≈ 2.87e−05
Cumulative Proportion
1.00
0.75
0.50
0.25
0.00
−1
0
1
log2(mRNA Fold Change)
Seed site (n=2974)
Removed seed site (n=135)
No seed site (n=8976)

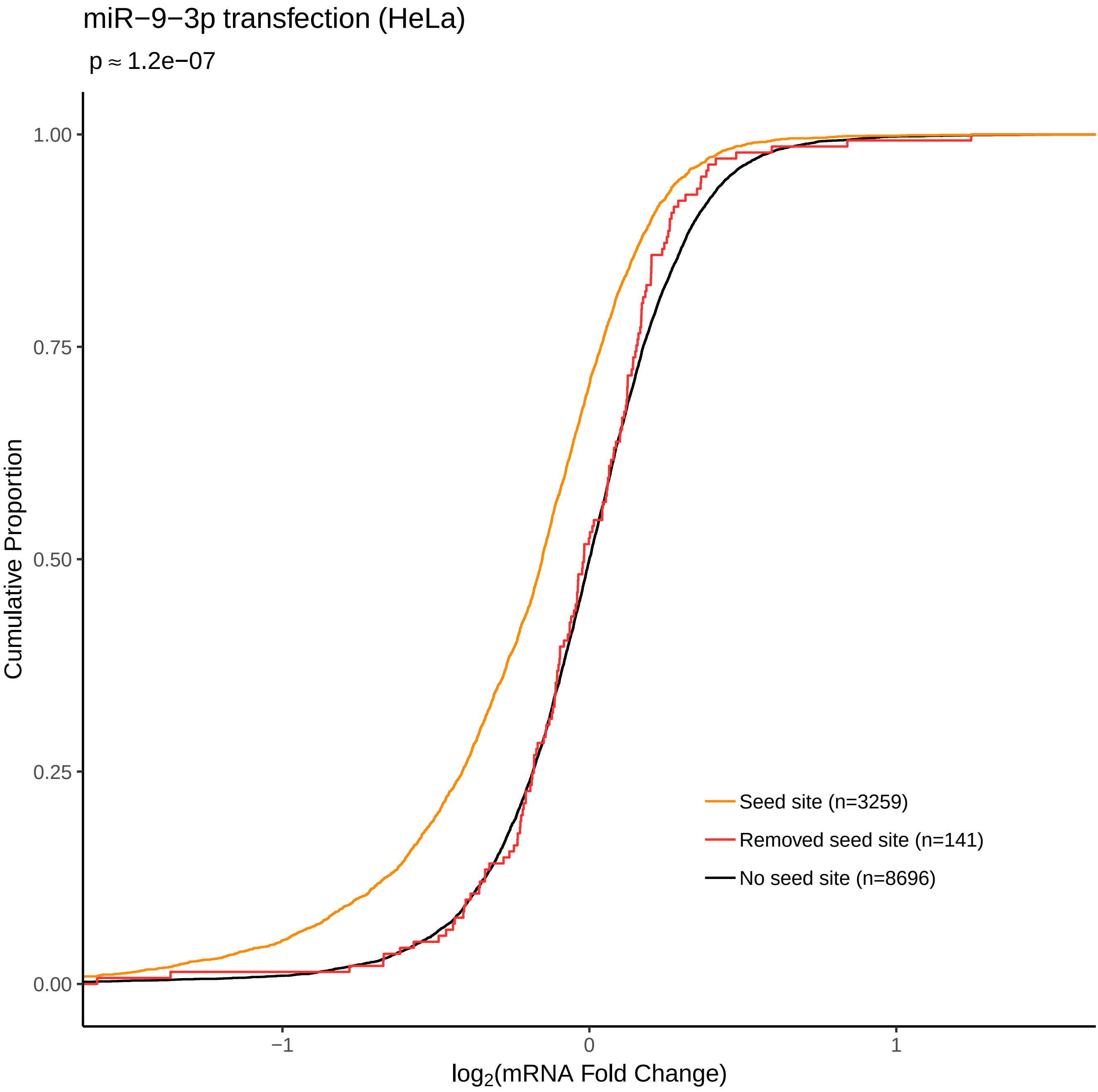
miR−9−3p transfection (HeLa)
p ≈ 1.2e−07
Cumulative Proportion
1.00
0.75
0.50
0.25
0.00
−1
0
1
log2(mRNA Fold Change)
Seed site (n=3259)
Removed seed site (n=141)
No seed site (n=8696)

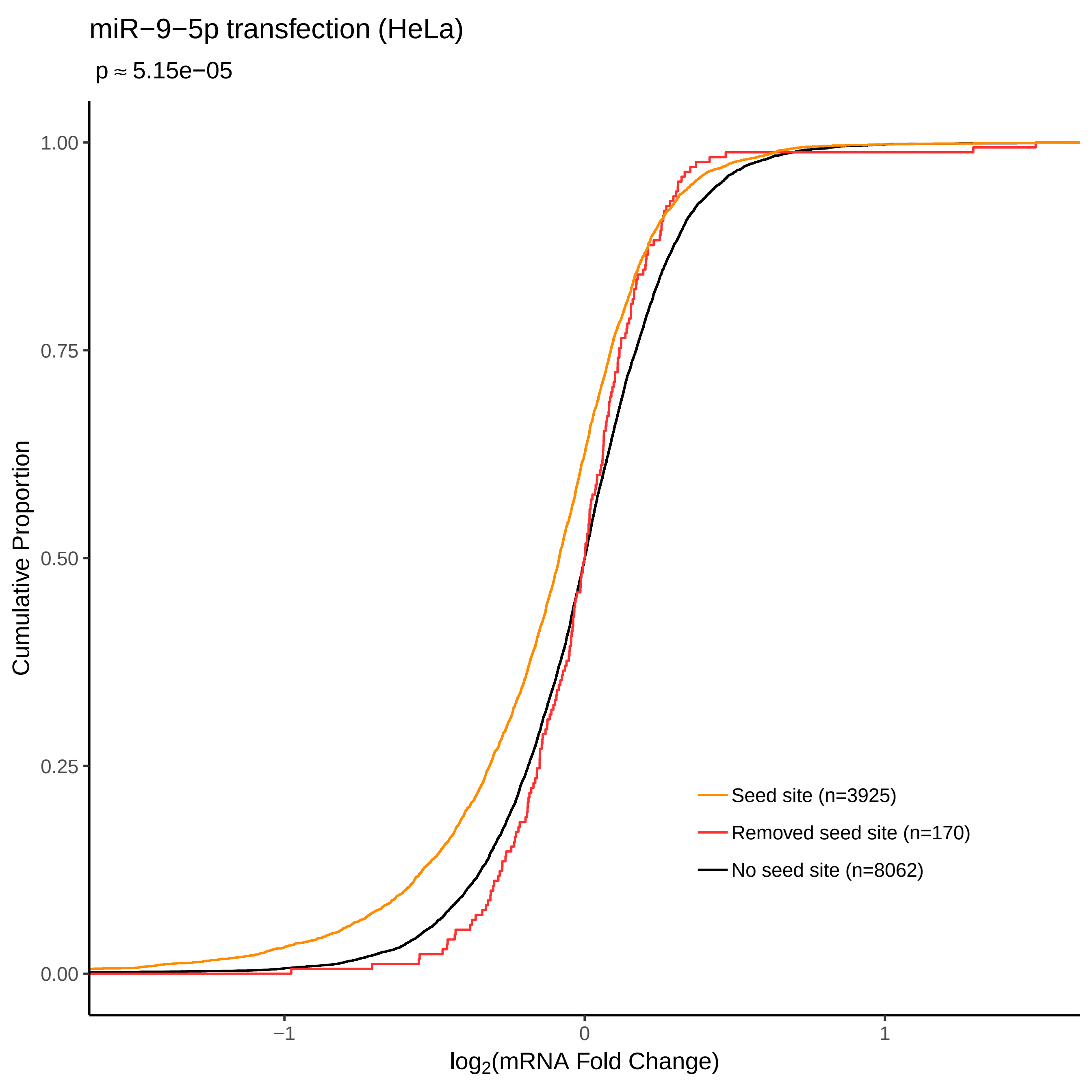

miR−9−5p transfection (HeLa)
p ≈ 5.15e−05
Cumulative Proportion
1.00
0.75
0.50
0.25
0.00
−1
0
1
log2(mRNA Fold Change)
Seed site (n=3925)
Removed seed site (n=170)
No seed site (n=8062)
